# Infrequent strong connections constrain connectomic predictions of neuronal function

**DOI:** 10.1101/2025.03.06.641774

**Authors:** Timothy A. Currier, Thomas R. Clandinin

## Abstract

How does circuit wiring constrain neural computation? Recent work has leveraged connectomic datasets to predict the function of cells and circuits in the brains of many species. However, many of these hypotheses have not been compared with physiological measurements, obscuring the limits of connectome-based functional predictions. To explore these limits, we characterized the visual responses of 91 cell types in the fruit fly and quantitatively compared them to connectomic predictions. We show that these predictions are accurate for some response properties, such as orientation tuning, but are surprisingly poor for other properties, such as receptive field size. Importantly, strong synaptic inputs are more functionally homogeneous than expected by chance, and exert an outsized influence on postsynaptic responses, providing a powerful modeling constraint. Finally, we show that physiology is a stronger predictor of wiring than wiring is of physiology, revising our understanding of the structure-function relationship in the brain.

## Introduction

Brains are structurally complex, with dramatic variation in architecture, cellular morphology, and synaptic connectivity (Shepherd, 2003). Understanding how these structural features shape functional properties has drawn interest for a thousand years, beginning with work relating the structure of the visual system to the optics of the eye (al-Haytham, ca. 1020). More recent work using physiological approaches has defined organizing principles that relate structure to function in many contexts, spanning scales that range from topographic maps of sensory systems (Kaas, 1997), to morphological asymmetries that allow specific neurons to respond to particular directions of visual motion (Kim et al., 2008). While these and many other studies demonstrate that structure and function are correlated, the limits of this relationship are unclear. To what extent can functional properties of cells and circuits be predicted based on their morphology and synaptic connectivity?

Ultrastructural imaging via electron microscopy (EM) has become a foundational tool for exploring brain structure at the level of individual cells and synapses (Williams & Carter, 1996). In the four decades since pioneering work in *C. elegans*, the reconstruction of synaptic connectivity maps, or “connectomes,” has expanded to include organisms ranging from ctenophores to humans (White et al., 1986; Meinertzhagen & O’Neil, 1991; Mishchenko et al., 2010; Rivera-Alba et al., 2011; Anderson et al., 2011; Takemura et al., 2013; Helmstaedter et al., 2013; Kim et al., 2014; Kasthuri et al., 2015; Motta et al., 2019; Verasztó et al., 2020; Scheffer et al., 2020; Phelps et al., 2021; Winding et al., 2023; Burkhardt et al., 2023; Chua et al., 2023; Bidel et al., 2023; Schneider-Mizell et al., 2023; MICrONS Consortium et al., 2023; Shapson-Coe et al., 2024; Azevedo et al., 2024; Lesser et al., 2024; Schoofs et al., 2024; Dorkenwald et al., 2024; Nern et al., 2024). These datasets cover different sensory systems, motor systems, cortical regions, and enteric ganglia, with some spanning the entire brain. Analyses of connectome data have produced insights into synaptic organization. For example, studies in rodent cortex (Kasthuri et al., 2015) and hippocampus (Mishchenko et al., 2010) revealed a uniform density of postsynapses along dendrites, and demonstrated that axo-dendritic contact was not sufficient for synapse formation. Analyses of connectome data have also provided insights into circuit function. For example, specific spatial patterns of synaptic inputs onto the dendrites of “elementary motion detector” neurons in both the fruit fly visual system and the vertebrate retina have informed models of motion processing (Takemura et al., 2013; Takemura et al., 2017; Briggman, Helmstaedter & Denk, 2011; Kim et al., 2014).

The promise of connectomic data lies in the generation of new hypotheses through granular anatomical descriptions of many cells and circuits. Recent work with the connectome of the adult fruit fly has revealed general principles of neural network organization (Lin et al., 2024; Dorkenwald et al., 2024; Schlegel et al., 2024) and has allowed researchers to predict the functional properties of neurons, including neurotransmitter identity, through anatomical analyses and network simulation (Eckstein et al., 2024; Zhao et al., 2023; Dombrovski et al., 2023; Pospisil et al., 2024; Shiu et al., 2024; Lappalainen et al., 2024; Garner et al., 2024; Seung, 2024a; Seung, 2024b; Ganguly et al., 2024; Schretter et al., 2024). Functional verifications of such predictions have typically focused on a small number of specific cell types (Briggman, Helmstaedter & Denk, 2011; Takemura et al., 2017; Zhao et al., 2023; Dombrovski et al., 2023; Shiu et al., 2024; Turner et al., 2024; Vishwanathan et al., 2024), or relied on artificial activation (Liu et al., 2022; Randi et al., 2023; Pospisil et al., 2024; Creamer, Leifer & Pillow, 2024). However, it has been challenging to obtain physiological measurements from a large number of cell types that can be identified in the connectome, making it difficult to assess the broad predictive capacity of connectome data.

The *Drosophila* visual system has proven to be a fertile ground both for functional dissection and connectomic characterization (reviewed in Currier, Pang & Clandinin, 2023; Meinertzhagen & O’Neill, 1991; Rivera-Alba et al., 2011, Takemura et al., 2013; Nern et al., 2024; Matsliah et al., 2024). The visual system comprises the retina, and four optic ganglia, the lamina, the medulla, the lobula, and the lobula plate that collectively include approximately 100,000 neurons of a few hundred types. As a whole, the structure is organized retinotopically, and is characterized by a high degree of morphological stereotypy across flies, such that many individual cell types can be identified by their distinctive shapes and synaptic connections. Both the lamina and the medulla receive direct input from photoreceptors, and the medulla is the last neuropil that all eye-derived visual signals must pass through. The medulla includes more than 100 cell types that each preferentially respond to specific visual features, including light and dark contrasts, colors, and directions of motion. Furthermore, the anatomical structure of the medulla has been described in detail, beginning first with light level studies, and culminating in several connectomes (Fishbach & Dittrich, 1989; Morante & Desplan, 2008; Nern et al., 2015; Meinertzhagen & O’Neill, 1991; Rivera-Alba et al., 2011, Takemura et al., 2013; Nern et al., 2024; Matsliah et al., 2024). Finally, connectome studies have predicted the visual response properties of many medulla cell types, offering a large set of testable hypotheses (Kind et al., 2021; Shinomiya et al., 2022; Seung, 2024a; Seung, 2024b).

Here we assess the power of connectomic data to predict functional properties of many cell types in the medulla. Our results confirm and challenge intuition in equal measure, and reveal which prediction types and modeling strategies are more likely to be accurate. Finally, we discover an unexpected asymmetry in the predictive relationship between circuit structure and cellular function, revising our understanding of how the connectome constrains brain activity.

## Results

### Efficient recording of many cell types with SPARC-L

To broadly assess the predictive power of connectome data, we sought to directly compare measured physiological properties to connectomic predictions of those properties across cell types. To do this, we needed to efficiently sample the response properties of many different cells. We chose to target the optic lobe, where many connectomic predictions of cellular responses have been made (Kind et al., 2021; Shinomiya et al., 2022; Zhao et al., 2023; Dombrovski et al., 2023; Seung, 2024a; Seung, 2024b; Garner et al., 2024; Lappalainen et al., 2024; Ganguly et al., 2024; Schretter et al., 2024). Specifically, we focused on the medulla, a region with substantial cell type diversity and several connectomic datasets (Meinertzhagen & O’Neill, 1991; Rivera-Alba et al., 2011, Takemura et al., 2013; Nern et al., 2024; Matsliah et al., 2024). To limit genetic variation across flies and capture responses from many neurons with single cell precision, we extended the SPARC method (Sparse Predictive Activity through Recombinase Competition; Isaacman-Beck et al., 2020) (Fig. 1A-B). In SPARC, two mutually exclusive recombination reactions compete. One reaction excises a STOP cassette, comprised of transcriptional and translational terminators, to turn on a transgene of interest. The other reaction leaves the STOP cassette intact, preventing transgene expression. The relative probability of these two outcomes can be tuned by varying the length of the attP sites that guide ΦC31-mediated recombination, allowing the fraction of transgene-expressing cells to be precisely controlled. We combined a pair of “layered” SPARC elements (SPARC-L), such that both cassettes must activate in the same cell. This strategy allowed us to target 0.1% to 1% of cells in a population of interest. Here we used neurotransmitter-specific GAL4 lines (Deng et al., 2019) to drive sparse expression of LexA (layer 1), which in turn drove sparse expression of GCaMP8m (layer 2, see Fig. 1A; Zhang et al., 2020). With this combination of tools, SPARC-L results in GCaMP expression in an extremely sparse, random subset of cells that express a specific neurotransmitter (acetylcholine, glutamate, or GABA). We identified GCaMP-labeled neurons by comparing their morphology to existing cell type atlases (Fischbach & Dittrich, 1989; Nern, Pfeiffer & Rubin, 2015) and connectomes (Matsliah et al., 2024; Nern et al., 2024) (Fig 1C). This generalizable approach allowed us to efficiently sample many cell types, without cell-type specific genetic reagents, at single-neuron resolution.

**Figure 1.**
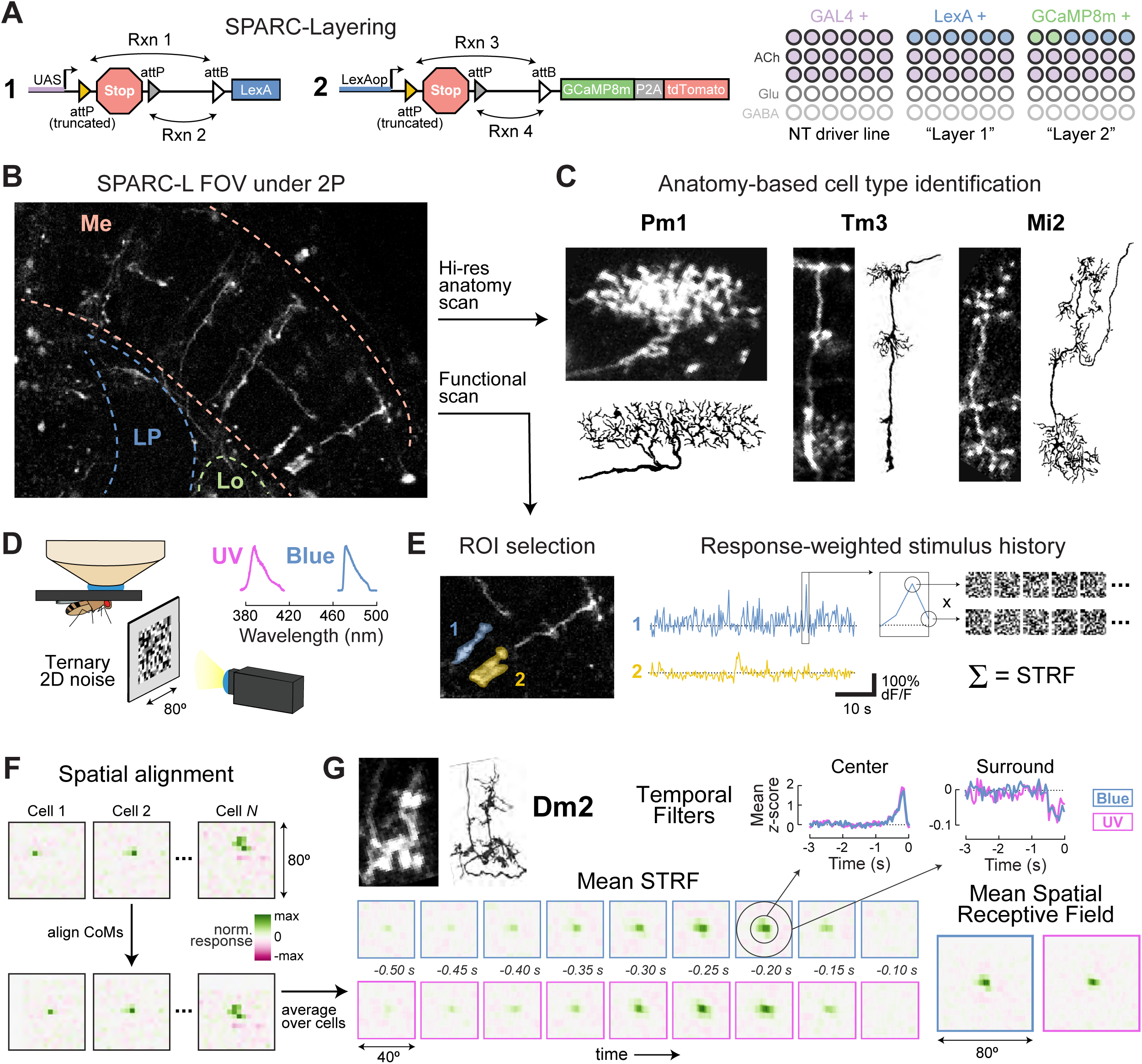
Efficient recording of many cell types with SPARC-L. (A) Left: Schematic illustration of SPARC-L. Two SPARC (Isaacman-Beck et al., 2020) cassettes are layered “in series,” with each layer sparsening expression via competing ФC31-mediated recombination reactions. Rxn 1 and Rxn 3 excise a STOP element and allow expression, while Rxn 2 and Rxn 4 do not. Right: Scematic illustration of expression patterns. GAL4 is expressed via a neurotransmitter-specific driver line. LexA is sparsely expressed in GAL4+ cells. GCaMP8m-P2A-myr::tdTomato is sparsely expressed in LexA+ cells. (B) An example field of view (FOV) under 2P imaging, showing sparse expression of GCaMP8m in the optic lobe. Neuropil borders are outlined. Me: medulla, Lo: lobula, LP: lobula plate. (C) Anatomy-based cell type identification. Images of three example neurons are shown next to traces of the same cell types (Fishbach & Dittrich, 1989). (D) Experimental setup. SPARC-L flies were imaged using a 2P microscope during 32 minutes of 2D ternary noise presentation. Separate noise trials were presented with either blue or UV light. Inset shows the spectral content of each stimulus. (E) Data analysis procedure. Left: ROIs were drawn over areas with high fluorescence variance that unambiguously contain the neurites of a single cell (Methods). Two example ROIs are shown from (B). Middle: 45 s of noise responses for the two example ROIs. Right: For each imaging sample, we weighted the prior 3 s of stimulus history by fluorescence intensity at that sample. Summing over weighted stimulus histories produces a spatiotemporal receptive field, which defines the stimulus that maximally excites that ROI. (F) Prior to averaging, all cells of the same type were spatially aligned. We shifted each STRF so that the center of mass of the strongest spatially contiguous element was centered (Methods). This procedure is illustrated for 3 example Dm2 neurons, with “raw” responses on the top, and “centered” responses on the bottom. (G) Visual selectivity profile for an example cell type (Dm2). Comparable data is available for all 91 cell types in our dataset (Fig. S1-S3). Top left: an imaged Dm2 neuron and a connectome reconstruction of Dm2 (Matsliah et al., 2024). Bottom left: the last 0.5 s and the central 40° of the mean STRF for Dm2. Top right: the full (3 s) temporal filters for Dm2 center and surround (see Fig. S3A). Bottom right: the full (80°) spatial receptive field for Dm2. For all plots, the response to blue (blue) and UV (pink) noise stimuli are shown separately. See also Fig. S4-S5.

We recorded neuronal activity in medulla neurons in SPARC-L flies under 2-photon excitation while presenting a 2-dimensional ternary noise stimulus (Fig. 1D). In this stimulus, each 5° patch of the display randomly adopts a dark, light, or gray intensity, updated at 20Hz, parameters that were chosen to match previously recorded responses of medulla neurons (Currier et al., 2024). We presented this stimulus in two color ranges, one in blue (peak λ = 475 nm), and one in ultraviolet (UV, peak λ = 385 nm), and tuned the intensity of both stimuli to equally excite R1-R6 photoreceptors. Regions of interest (ROIs) containing the neurites of single neurons were identified as areas with high variance over the course of noise presentation, allowing semi-automated identification of responding cells (Fig. 1E and Methods). For every point in each ROI’s response time-course, we weighted the prior 3 sec of stimulus history by the amplitude of the calcium response, then summed over time to produce a response-weighted stimulus history (Clark et al., 2011; Leong et al., 2016). These weighted histories capture the spatial and temporal stimulus features that drove the strongest activity for each ROI, defining a spatiotemporal receptive field (STRF). We performed this weighting separately for the blue and UV noise stimuli, yielding distinct STRFs for each color.

We recorded single neuron STRFs in 54 SPARC-L flies, yielding 571 ROIs from 510 unique neurons (hereafter referred to as the “full” dataset). High resolution anatomical scans allowed us to assign cell type identities to 365 of these recordings, yielding STRFs for 91 unique cell types (Table S1; hereafter referred to as the “labelled” dataset). The remaining 145 cells could not be uniquely identified due to unfavorable labeling density. Of the 91 cell types in the labelled dataset, approximately 55 have not been previously recorded. Because SPARC-L labels neurons randomly, cell types that are less numerous are infrequently labelled: of the 91 recorded cell types, 43 were sampled three or more times. Unless otherwise noted, we restricted all subsequent analyses to these cell types (Fig. S1, hereafter referred to as the “main dataset”). For completeness, the visual responses of “low *N*” cell types (sampled only once or twice) are shown in Fig. S2.

We first summarized the visual selectivity of each cell type in our labelled dataset. Because the spatial receptive field of each neuron was slightly different, we shifted each STRF in x and y to align their centers of mass (Fig. 1F). We then z-scored spatially aligned STRFs and took the average across independent samples (cells) of each cell type (Fig. 1G). We then defined temporal receptive fields (TRFs) for each cell type: the center TRF was the time-course of the 5° patch with the strongest response, and the surround TRF was the average time-course of the largest oppositely signed and spatially contiguous set of 5° patches above the noise floor (Fig. S3, Methods). Finally, we calculated a mean spatial receptive field for each cell type (Fig. S3A-B, Methods).

To validate SPARC-L, we compared our results to noise recordings derived from cell-type specific targeting (Arenz et al., 2017; Drews et al., 2020). Over 12 previously characterized cell types, our results largely recapitulated earlier work (Fig. S4). One notable exception is that our STRFs, measured with GCaMP8m, had consistently faster dynamics than prior recordings using GCaMP6f, reflecting the faster kinetics of the newer sensor. We also note that our 2D noise did not drive spatial surrounds as strongly as the 1D noise used previously, again consistent with expectations. We also observed within-cell type variability, even when the same cell type was unambiguously recorded multiple times in a single fly (Fig. S5), perhaps reflecting biological variation. Similar variation has been previously observed in some medulla neurons using cell-type specific genetic tools (Fisher et al., 2015; Cornean et al., 2024). Overall, our results suggest that recording with SPARC-L, followed by anatomy-based cell type identification, is an efficient strategy for recording many cell types with single cell precision.

### Distinct patterns of visual representation across medulla cell types

We first sought to holistically assess visual representation across our full set of recorded cells. We performed a principal components analysis (PCA) using single neuron responses as samples, revealing a set of common spatiotemporal motifs, ranked by the fraction of cross-neuron variance they explain (Fig. 2A-B and Fig. S6). The six highest-ranking PCs in our analysis correspond to previously observed visual features (Fig. 2A). We found that the dominant pair of PCs represented temporal motifs: one monophasic PC that integrates over light intensity, and one biphasic PC that detects changes in light intensity over time. The next pair of PCs each describe half of the canonical center-surround spatial organization that is common in both fly and vertebrate visual systems (Kuffler, 1953; Freifeld et al., 2013), while a third pair of PCs show oppositely signed weights for blue and UV light, a hallmark of spectral opponency (Wagner, Macnichol & Wolbarsht, 1960; Heath et al., 2020; Li et al., 2021). Plotting variance explained as a function of PC rank reveals less explanatory power for PCs beyond these first six (Fig. 2B).

**Figure 2.**
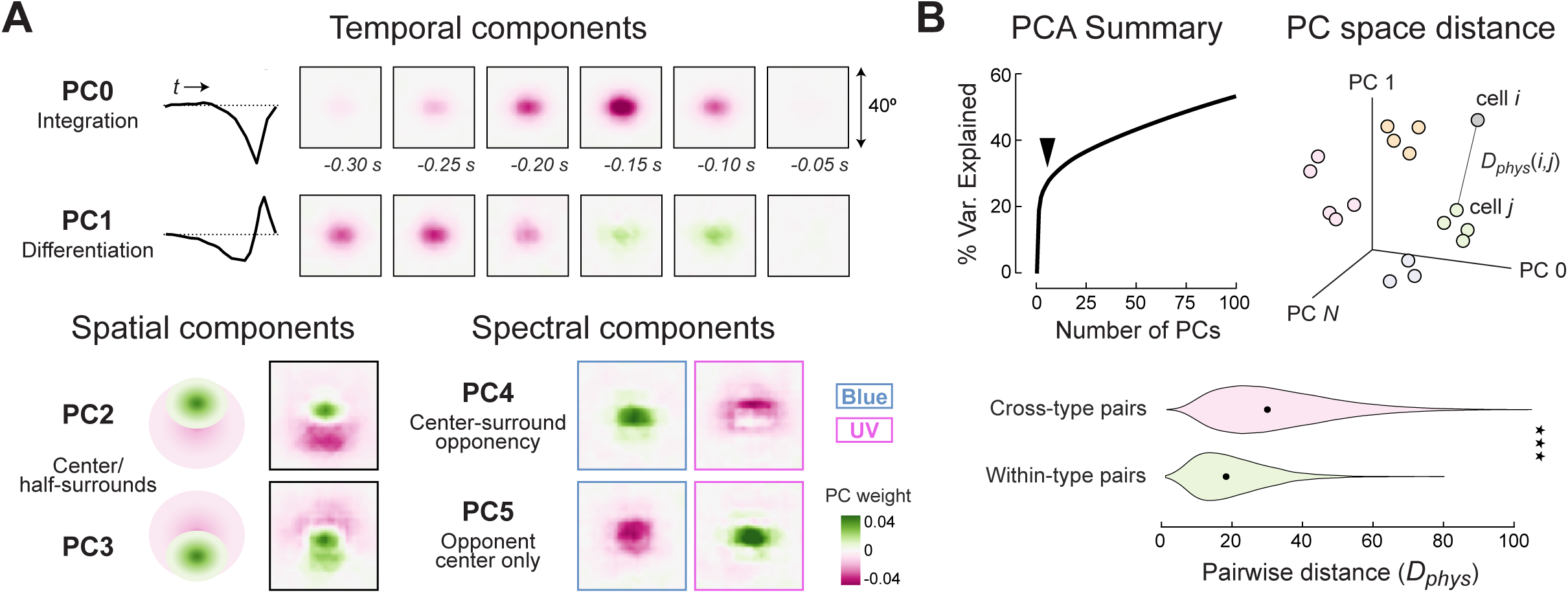
Distinct patterns of visual representation across medulla cell types. (A) Principal components analysis. Specific aspects of the first six PCs are highlighted. Top: the last 0.3 s and central 40° of PC0 and PC1, with a schematic time-course of PC weights. Bottom left: single temporal snapshots of PC2 and PC3, with schematic illustrations of their spatial arrangements. The spatiotemporal details are similar for both blue and UV components of PCs 0-3. Bottom right: single temporal snapshots of PC4 and PC5, illustrating the blue and UV components separately. PCA model details are in Fig. S6. (B) Top left: Cumulative % variance explained as a function of the number of PCs. Arrowhead indicates the dramatic change in slope that occurs after the first six PCs. Top right: illustration of the principal component space into which single neuron responses are projected. We calculate the Euclidean distance (*D_phys_*) between all pairs of neurons. Bottom: violin plots of *D_phys_* for different categories of cell pairs. Black dots indicate the medians of each distribution. Within-type *D_phys_* is significantly lower than cross-type *D_phys_* by rank-sum test (*p* < 7.1 x 10^-304^).

We projected the responses of individual neurons into a space defined by the first 100 PCs (Fig. 2B). This dimensionality reduction strategy provides an efficient way to summarize and compare the functional properties of cells and cell types. In this “PC space”, we measured the Euclidean distance (*D_phys_*) between every pair of neurons, allowing us to quantify their functional similarity. As an additional check on the accuracy of our morphology-based cell type identification strategy, we asked whether *D_phys_* was smaller for pairs of cells of the same morphological type compared to pairs of cells of different morphological types. We found that the distribution of “within-type” distances was significantly smaller than the “across-type” distribution (*p* < 7.1 x 10^-304^). These results further validate our recording strategy and establish *D_phys_* as a quantitative measure of similarity between cells and cell types, which we will later use to define functional sub-types.

### Predicting feature selectivity from connectomes

As an initial exploration of the predictive power of connectome data, we examined predictions relating to three fundamental elements of visual processing: orientation selectivity (OS), directional motion selectivity (DS), and spectral preference. For example, prior work described asymmetrically elongated neurites in Dm3 and TmY9 neurons, and posited that these cell types might possess OS (Seung, 2024b). To calculate an orientation selectivity index (OSI) across our main dataset, we first merged each cell’s blue and UV STRFs into a single “joint-color” STRF (Methods). We then thresholded this joint-color STRF (Fig. S3D) and integrated over the interval defined by the final lobe in the temporal filter (Fig. 3A; Fig. S3B). By fitting an ellipse to this thresholded spatial receptive field, we could calculate OSI as the ratio of the long axis to the short axis. This metric has a minimum value of 1, with larger values indicating stronger OS (Fig. 3A). Importantly, previously recorded cell types that lack orientation tuning, such as L1 (Drews et al., 2020), displayed little OS in our data. Conversely, we detected strong OS in T4, in line with previous measurements (Maisak et al., 2013; Fisher, Silies & Clandinin, 2015). In agreement with connectome predictions, we also found that subtypes of Dm3 and TmY9 did indeed exhibit moderate to strong OS (Fig. 3B-C). Intriguingly, we also observed moderate to strong OS in nine other cell types that had not been previously predicted to have OS. Many of these cell types, including Dm13, Dm16, TmY16, and Pm1, have pronounced arbor elongation, consistent with de novo computation of OS in their dendrites. Conversely, a small number of other cell types are OS, but do not have elongated arbors, including Mi1, T2, and TmY5a. These cells might therefore inherit orientation tuning from their upstream synaptic partners.

**Figure 3.**
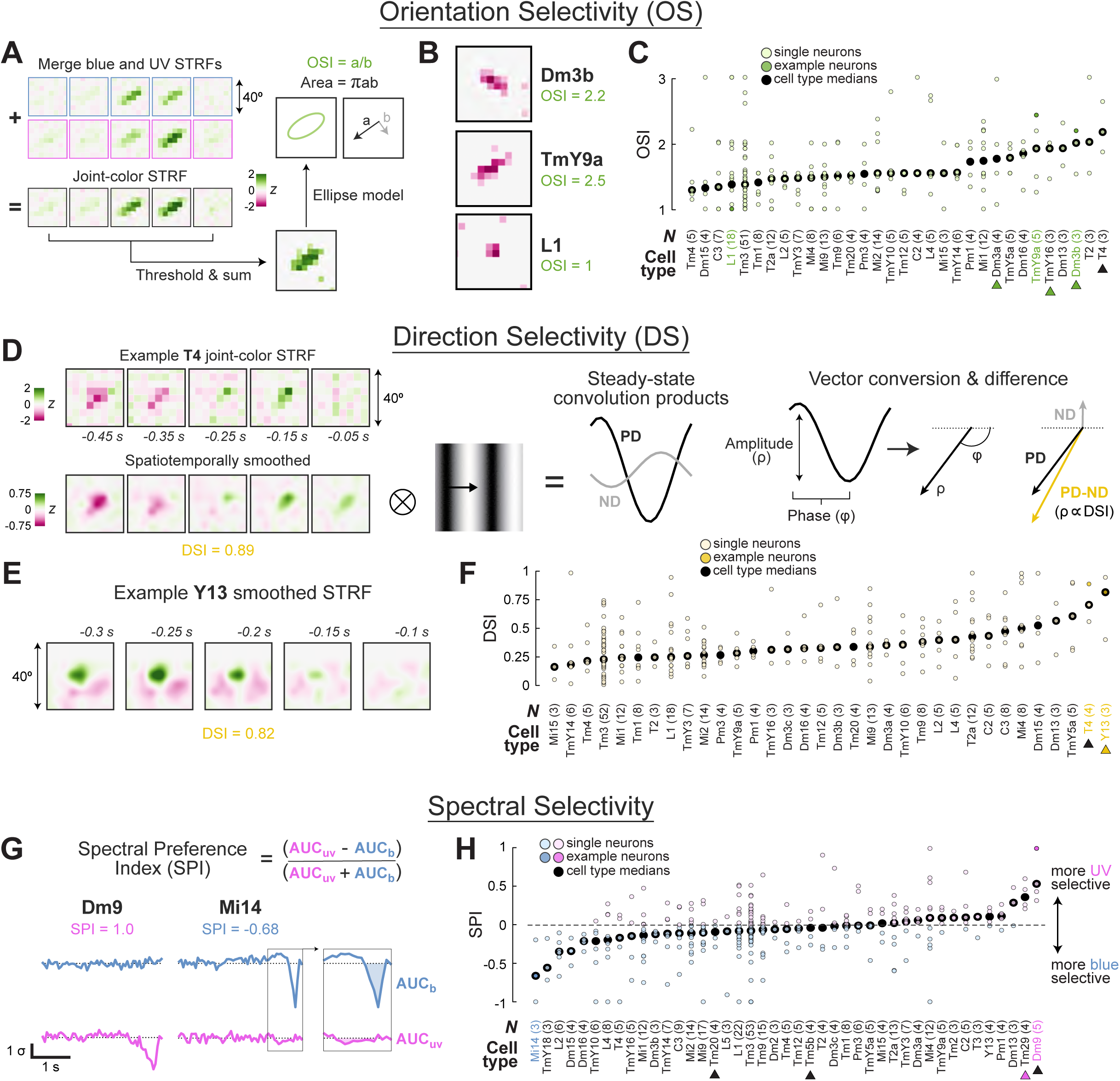
Specific response properties match qualitative connectome-based predictions. (A) Procedure for calculating an orientation selectivity index (OSI). Blue and UV STRFs for each cell are merged, thresholded, and integrated over the last temporal lobe (Methods). An ellipse is fit to this thresholded receptive field, and OSI is calculated as the ratio of the ellipse long axis to the ellipse short axis. (B) Thresholded receptive fields for three example neurons and the corresponding OSI for each cell. (C) OSI by cell type for the main dataset, excluding neurons with strong color preference or small spatial receptive fields (Methods). Small green dots are individual neurons, large black dots are cell type medians. Example cells shown in (B) are highlighted (darker green dots; and cell type names in green). The number of neurons measured from each cell type are shown in parentheses. Green arrows indicate cell types predicted to be OS based on connectome data (Seung, 2024b). Black arrow indicates a cell type previously shown to be OS. (D) Procedure for calculating direction selectivity index (DSI). The joint-color STRF was smoothed in space and time, then convolved with sinusoidal grating “stimuli” that moved in one of eight directions (Methods). Steady-state convolution products were also sinusoidal, with an amplitude and phase that was converted to a vector length and angle, respectively. DSI is computed as the vector difference between the largest-amplitude response (preferred direction, PD) and the response to motion 180° opposite the PD (null direction, ND). A smoothed STRF is also shown for one example T4 neuron. (E) Smoothed STRFs for two example neurons and the corresponding DSI for each cell. (F) DSI by cell type for cells in the main dataset, excluding neurons with strong color preference or small spatial receptive fields (Methods). Yellow arrow: cell type predicted to be DS based on connectome data (Shinomiya et al., 2022). Black arrow: cell type previously shown to be DS. (G) Procedure for calculating spectral preference index (SPI). For each cell, we integrated under the last lobe of the blue and UV center TRFs. SPI was defined as the difference between these AUCs, over their sum. Opponent responses with oppositely signed blue and UV AUCs were manually set to 1 (UV preference) or -1 (blue preference, see Methods). The example Dm9 neuron shown is one such case. (H) SPI by cell type for the main dataset. Magenta arrow: cell type predicted to be color selective based on connectome data (Kind et al., 2021). Black arrows: cell types previously shown to be color selective. Note that Tm20 and Tm5b are not expected to differentiate between the wavelengths of light used in this study.

We next performed an analogous analysis for DS, a prominent feature of T4/T5 responses and a subject of extensive study (Currier, Pang & Clandinin, 2023). A connectomic reconstruction of the lobula complex, the primary output neuropil of medulla projection neurons, revealed significant T4/T5 input to the “Y” family of cell types, including Y13 (Shinomiya et al., 2022). In that work, the authors suggested that these cell types might also show strong DS, having inherited it from T4/T5. To test this prediction and identify other DS cell types in the medulla, we simulated responses to drifting sinusoidal gratings for each cell type in our main dataset (Fig. 3D). We spatiotemporally smoothed joint-color STRFs, then convolved them with gratings moving in 8 directions (Methods). We defined the preferred direction (PD) for each cell as the movement direction eliciting the largest response. The null direction (ND) was defined as 180° opposed to the PD. By converting the PD and ND convolution products to vectors, we could define DSI as the relative length of the difference vector (PD-ND). These simulations revealed only two cell types with strong DS (Fig. 3E-F). Consistent with previous work, we found that T4 neurons show strong DS (Fig. 3D,F). Strikingly, Y13 also displayed strong DS (Fig. 3E), in agreement with connectomic predictions. These data support the notion that the *de novo* computation of DS is rare in the visual system, and may be unique to T4/T5 (Shinomiya et al., 2022; Zhao et al., 2023).

Finally, we examined blue vs. UV selectivity across cell types (Fig. 3G-H). R7 photoreceptors respond strongly to light in the UV range but not to light in the blue range, and directly innervate a small number of medulla cell types (Morante & Desplan, 2008; Karuppudurai et al., 2014; Kind et al., 2021; Nern et al., 2024). One simple prediction is that cells receiving such input might display a chromatic preference. To calculate a spectral preference index (SPI), we integrated under the last lobe of each neuron’s blue and UV center TRFs. We then set SPI to equal the difference between these integrated values divided by their sum (Fig. 3G). The resulting index ranges from -1 (strong blue preference) to 1 (strong UV preference), with values around 0 indicating no spectral preference. We found cell types with strong selectivity for either blue or UV (Fig. 3H). In agreement with prior studies, Dm9 displayed robust UV selectivity, while Tm20 and Tm5b had very little preference for the wavelengths used here (Heath et al., 2020; Christenson et al., 2024). Of the other cell types receiving direct R7 input, only Tm29 showed robust spectral selectivity. L1, C2, Dm2, Mi9, and Mi15, all of which receive only modest R7 input (Kind et al., 2021; Nern et al., 2024), did not show selectivity. Intriguingly, none of the cell types we found to be blue selective were identified as part of a predicted “color processing subnetwork” (Matsliah et al., 2024), suggesting that it may be easier to accurately predict UV selectivity from connectome data. In addition, many cell types we recorded at low *N* (in the labelled dataset) also displayed chromatic preference according to connectomic predictions, including Tm5a, Tm5Y, Tm34, Tm37, Tm38, Dm8, and Dm11 (Fig. S2). Collectively, these results suggest that important visual features, like OS, DS and spectral preference, can often be predicted by connectivity.

### Arbor size does not predict spatial receptive field area

One seemingly intuitive assumption is that visual neurons with larger arbors, receiving input from more medulla columns, will have larger spatial receptive fields (Seung, 2024a; Garner et al., 2024; Ganguly et al., 2024). However, prior work has also revealed that neuronal arbors can be functionally compartmentalized (Yang et al., 2016; Meier & Borst, 2019), and that both passive and active membrane properties can restrict the spread of postsynaptic currents (Jack, Noble & Tsien, 1975; Liu et al., 2022). As a result, how well arbor size correlates with receptive field size across cell types is unclear.

To address this question, we fit an ellipse to the receptive field center for each neuron in our main dataset (as in Fig. 3A). We calculated the area of this ellipse, then found the median for each cell type (Fig. 4A-B). The fly’s interommatidial angle is approximately 4.6° (Stavenga, 2003), meaning that a single medulla column receives feedforward input covering as little as πr^2^ = 17 deg^2^ of visual space. Recent connectomic models have assumed that each additional column innervated by a given cell increases its receptive field size by this amount (Zhao et al., 2023; Seung, 2024a; Garner et al., 2024; Ganguly et al., 2024). Under this assumption, receptive field areas for the cell types in our main dataset, which receive input from one to 62 columns, should range from 17 deg^2^ to 1,054 deg^2^. Consistent with prior observations, individual neurons had center areas as small as a single stimulus patch (20 deg^2^) and as large as half the screen (800 deg^2^), with median areas for each cell type ranging from 100 deg^2^ up to 368 deg^2^. Small neurons (like L1) often possessed small receptive fields, and large neurons (like Pm5) often possessed larger receptive fields (Fig. 4A). However, many counter examples were also observed: Mi2 is a small, columnar neuron with some of the largest receptive fields in the dataset. Conversely, Dm13, the largest cell in our main dataset, had a receptive field of only average size.

**Figure 4.**
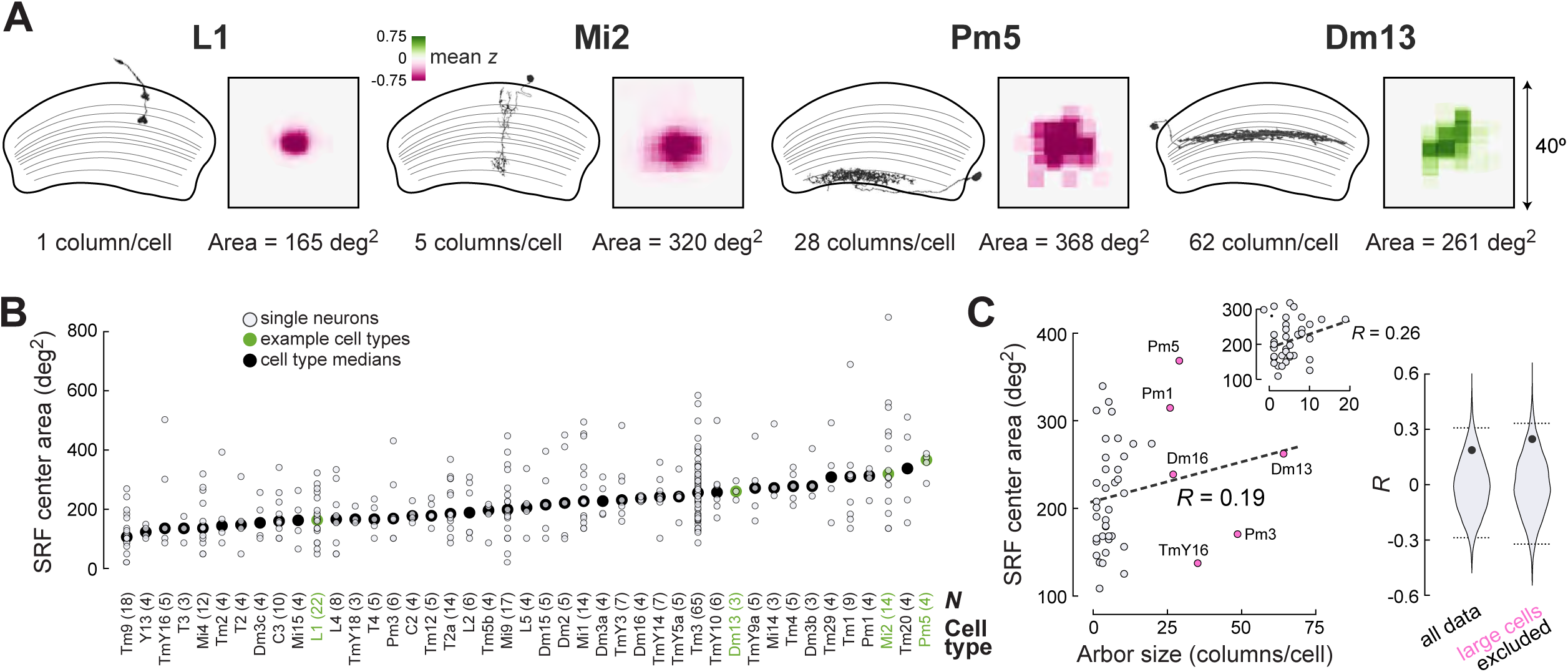
Arbor size does not predict spatial receptive field area. (A) Morphology cartoons and mean thresholded receptive field centers (as in Fig. 3B) for four example cell types, with arbor size (in columns/cell) and RF area (in deg^2^). RF area is calculated as shown in Fig. 3A. (B) Spatial receptive field center area by cell type for the main dataset. (C) Left: median spatial receptive field center area as a function of arbor size for each cell type. Dashed line indicates the best linear fit to the data, and the correlation coefficient (*R*) illustrates the quality of this linear fit. Inset shows the same scatter and best linear fit with large cells (pink) removed. Anatomical data from Nern et al., 2024. Right: shuffle-based bootstrapping (Methods). The distribution of *R* values for 10,000 resampled datasets is shown, with dashed lines indicating the 95% confidence interval. The *R* value of the real data, marked as a black dot, does not cross the 95% CI for either correlation.

These observations suggested that arbor size may not strongly predict receptive field size. When we correlated receptive field area and arbor size, the relationship was not significant (*R* = 0.19) and the slope of the best-fit line was only 0.93 deg^2^/column (Fig. 4C). Because large neurons innervating more than 20 columns were rare in our main dataset, we wondered if excluding these potential outliers might recover the assumed relationship. However, even with these large cells removed from the correlation, arbor size did not significantly predict receptive field area (*R* = 0.26), and the best-fit slope remained far below 17 deg^2^/column (Fig. 4C, inset). These results demonstrate that receptive field size cannot be estimated based solely on arbor size, contradicting a common assumption of predictive connectomics.

### Presynapse number and postsynapse density predict response amplitude

We next sought to evaluate connectomic predictions of more general functional properties. A core assumption when building connectome-inspired models is that more synapses lead to a stronger response. To test this hypothesis, we explored which morphological and synaptic parameters correlate with response strength across cell types (Fig. 5A-B). We began with the center TRF for each neuron in our main dataset, and used the peak *z*-score of the calcium response as a measure of response strength (Fig. 5C). We then found the median peak z-score for each cell type (Fig. 5D), and correlated the absolute value of these medians with a variety of connectome-derived variables (Fig. 5E-F and Fig. S7). We first asked whether cell types with more inputs (postsynapses) have larger responses. This relationship was not significant (*R* = -0.19, Fig. S7A); however, this simple model does not consider passive membrane properties, which spatially dampen incoming postsynaptic currents (Jack, Noble & Tsien, 1975; Liu et al., 2022). In other words, synapses that are physically separated (at low density) would be expected to interact less than synapses that are close together (at high density). We therefore asked whether response amplitude correlated with input *density* (i.e., postsynapses per unit area) (Turner et al., 2024; Liu et al., 2022). To compute input density, we normalized each cell type’s postsynapse counts by the average number of columns that cell type innervates. Under these conditions, we observed a significant correlation between postsynapse density and peak z-score (*R* = 0.55, *p* < 0.001, Fig. 5E). This relationship supports the intuition that more inputs, closer together, predict greater response strength.

**Figure 5.**
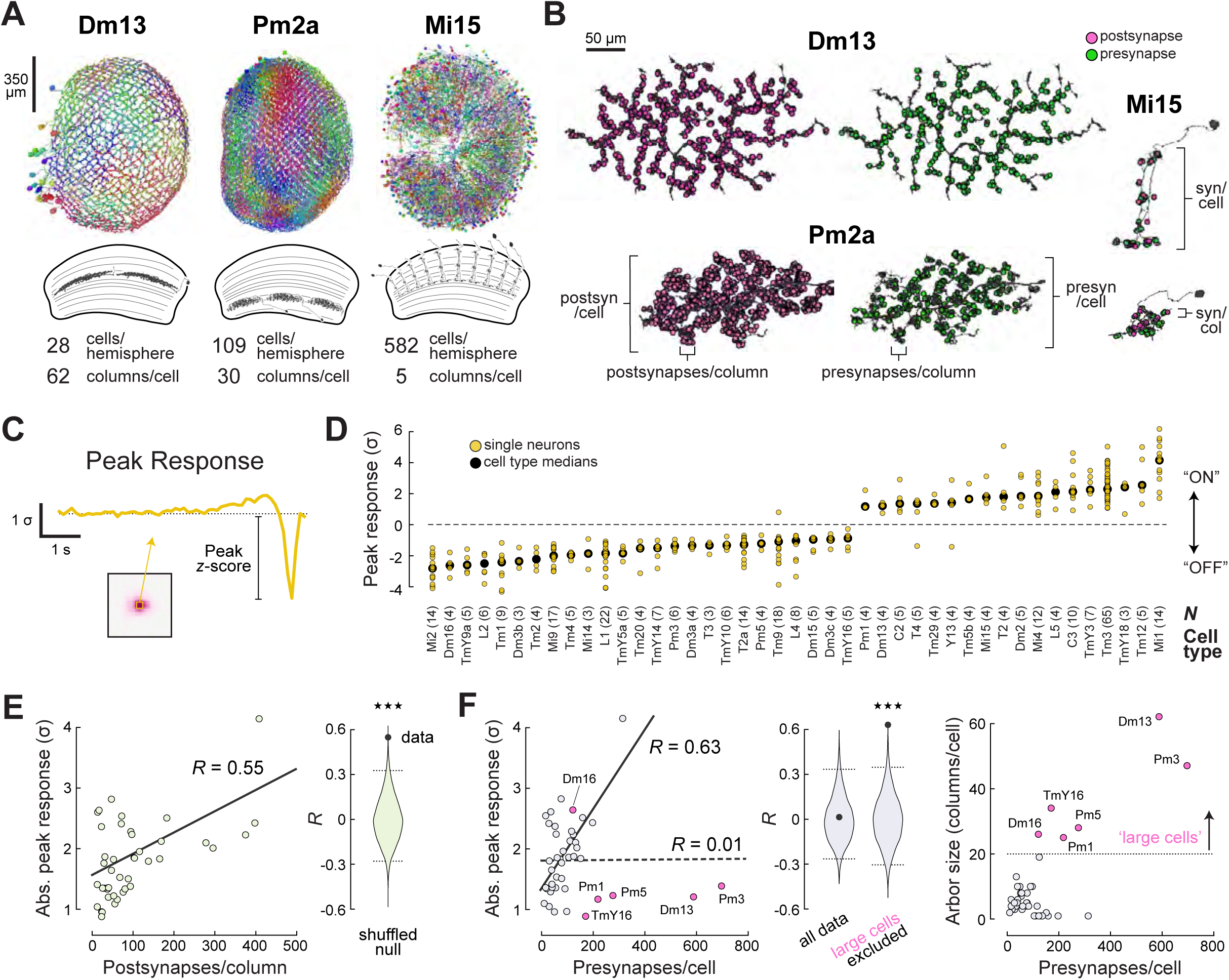
Presynapse number and postsynapse density predict response amplitude. (A) Morphological variation in three example cell types, highlighting connectome-derived variables. Top: whole-optic lobe reconstructions of each cell type, with individual neurons drawn in different colors (Matsliah et al., 2024). The medulla is oriented on-face in each image. Bottom: cartoon depictions of horizontal slices through part of the medulla, illustrating the different sizes of each cell type. (B) Synaptic variation in three example cell types, highlighting connectome-derived variables. One example reconstruction is shown twice for each cell type, with postsynapses (inputs) in magenta and presynapses (outputs) in green. All images are oriented on-face, except for the top Mi15 image, which simulates the horizontal slices cartooned in (A). (C) Procedure for calculating peak response. The center TRF, drawn from the max-responding patch of the overall STRF (Fig. S3A), is used to find the largest absolute z-score for each neuron. (D) Peak response by cell type for cells in the main dataset. Positive values are generally associated with ON (contrast increment) preferences, while negative values are generally associated with OFF (contrast decrement) preferences. (E) Left: median absolute peak response as a function of postsynapse density for each cell type. Solid line indicates the best linear fit. Right: shuffle analysis, as in Fig. 4C. The *R* value of the real data lies outside the 99.9% CI of the resampled *R* distribution. See Figure S7 for additional correlations. (F) Left: same as E, but for presynapse number. Dashed line is the non-significant correlation that considers all cell types. Solid line is the significant correlation when large cell types (pink) are excluded. Middle: shuffled null distributions for the “all data” and “large cells excluded” correlations. The real data lies outside the 99.9% CI for the latter correlation. Right: Arbor size as a function of presynapse number, highlighting that cell types larger than 20 columns (dotted line) are the outliers in the plot on the left.

We next wondered whether more *outputs* (presynapses) might also be predictive of stronger responses. For instance, we reasoned that neurons with higher presynapse densities might require elevated calcium signals to drive coincident synaptic release. Across all cell types, presynapse number was uncorrelated with peak z-score (*R* = 0.01, Fig. 5F). However, we noticed that smaller cells, innervating fewer than 20 columns, displayed a strong relationship between presynapse number and response strength. When we excluded the six cell types in our main dataset that innervate more than 20 columns, we observed a robust correlation (*R* = 0.63, *p* < 0.001). Curiously, output *density* only predicted response strength for a handful of “highly presynaptic” cell types that innervate only one column, and have many synaptic outputs (*R* = 0.62, *p* < 0.001, Fig. S7B). Together, these results suggest that smaller neurons with higher presynaptic densities indeed do have stronger calcium responses.

Finally, we wondered whether the previously noted within-cell type variance (Fig. S5) might be a “feature” of the visual system (biological noise). One naïve hypothesis is that more variation might be observed in cell types that could have greater functional redundancy, either through a larger population size (more cells per hemisphere), or through greater spatial overlap (more cells per column). We quantified within-type variance as the median distance in PC space (*D_phys_*) among members of each cell type (Fig. S7C). Neither population size nor spatial overlap correlated with median *D_phys_* (*R* = 0.2 and -0.04, respectively, Fig. S7D-E). However, we did observe a modest relationship between postsynapse density and within-type variability (*R* = 0.39, *p* < 0.01, Fig. S7F). This result is consistent with the intuition that a cell type that draws from a larger pool of potentially noisy inputs can display a greater diversity of responses.

### Strong inputs are highly correlated with their postsynaptic partners

Having demonstrated that, across all inputs to a cell, higher synaptic density predicts larger responses (Fig. 5E), we wondered whether this relationship held true for individual connections. Specifically, we wondered if a strong input comprised of many synapses will have a greater influence on a postsynaptic neuron’s temporal dynamics than a weak input comprised of few synapses. We first asked whether the correlation coefficient between pre- and post-synaptic TRFs was larger for strong connections than for weak connections (Fig. 6A). We began by calling a connection “strong” when it accounted for 5% or more of the postsynaptic partner’s total input (i.e., input fraction [%_in_] > 5%), and calling a connection “weak” when it was below this threshold. Depending on the total synaptic density of each cell type, an input fraction of 5% corresponded to approximately 15-30 synapses. Consistent with recent work in human cortex (Shapson-Coe et al., 2024), strong connections are relatively rare in the fly: across connected cell type pairs in our main dataset, this >5%_in_ threshold captured the strongest 10% of connections (Fig. S8A). To account for inhibitory connections, we multiplied the TRF cross-correlation by -1 when the presynaptic neuron in a connected pair was glutamatergic or GABAergic (Eckstein et al., 2024). While both strong and weak connections covered the full range of correlation values, strong positive relationships were much more common for strong inputs (Fig. 6A, *p* < 0.01). This significant difference persisted across a range of %_in_ thresholds, ranging from a cutoff fraction of 10% down to 2% (∼6-12 synapses; Fig. S8A). These results are consistent with another fundamental assumption of predictive connectomics: that strong inputs have a larger effect on postsynaptic neurons than weak inputs.

**Figure 6.**
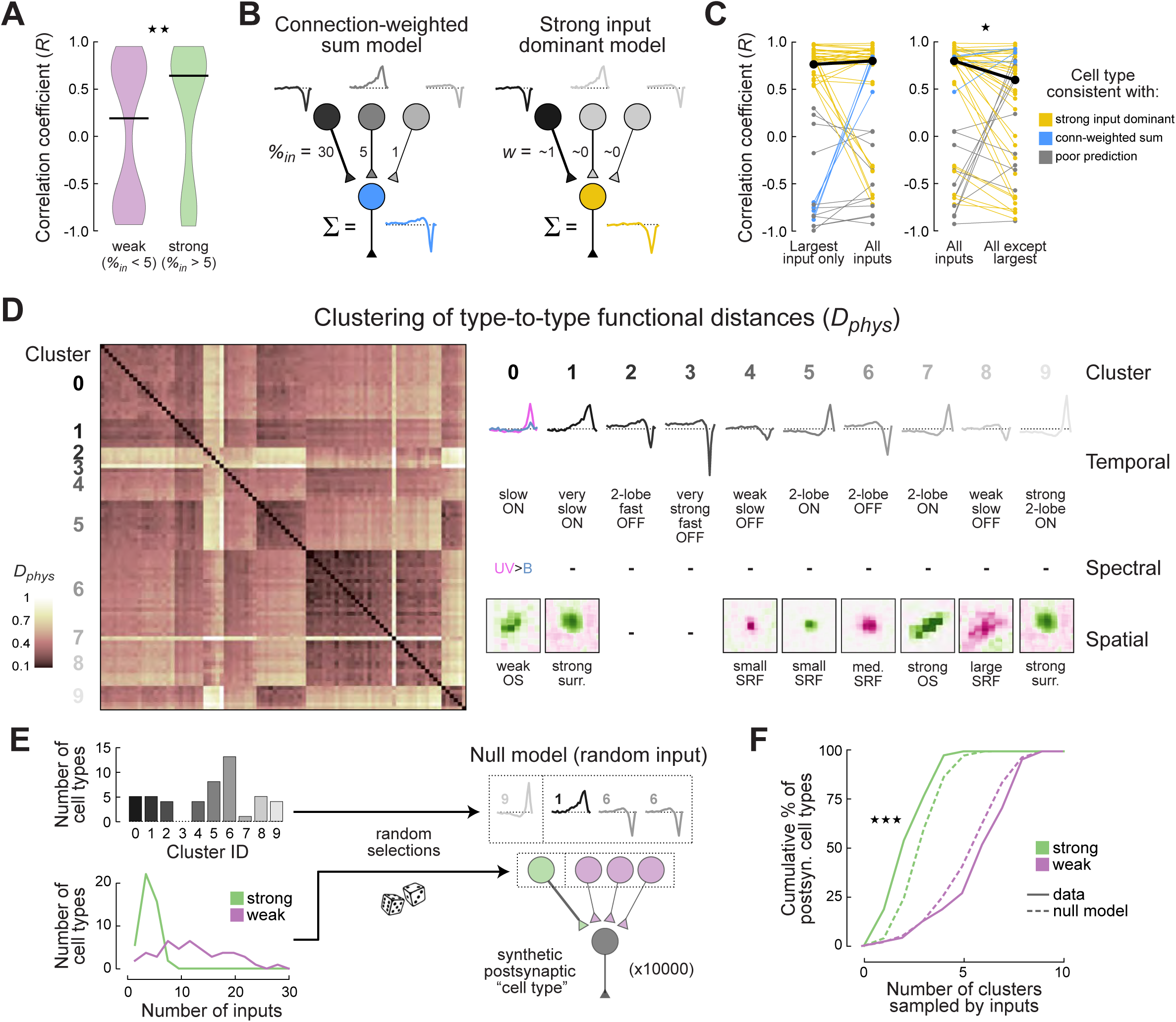
Strong inputs are functionally homogeneous and highly correlated with their postsynaptic partners. (A) Distributions of cross-correlation coefficients between the center TRFs of all connected cell type pairs. Connections are grouped according to the fraction of total input onto the postsynaptic partner that is provided by the presynaptic partner. Strong connections (green) have an input fraction greater than 5%, while weak connections (purple) are below this threshold. Black lines show distribution medians. Strong connections show significantly stronger pre-/postsynaptic correlations (rank-sum test, *p* < 0.01), and this difference persists over a range of strong/weak cutoff values (Fig. S8A). (B) Two models of synaptic inheritance. Left: a linear model that weights presynaptic responses by the fraction of total input each presynaptic cell type provides. Weighted presynaptic responses are summed to produce the predicted postsynaptic activity. Right: a nonlinear model that over-weights strong inputs and under-weights weak inputs. The predicted postsynaptic response strongly resembles the strongest input’s activity. (C) Correlation coefficients between measured responses and a linear connection-weighted sum prediction that considers different subsets of inputs. Each line is one cell type, colored by the model that cell type is most consistent with: yellow for the nonlinear “strong inputs dominant” model, and blue for the linear “connection-weighted sum of inputs” model. Gray lines are not consistent with either model. Black is the median across cell types. Left: Predictions are not improved by adding more inputs beyond the single largest input (see also Fig. S8B). Right: removing the single largest input from the full model significantly reduces prediction quality (signed-rank test, *p* < 0.05). (D) Functional clustering of all 91 cell types in the dataset via affinity propagation. Cell types are clustered according to their Euclidean distance (*D_phys_*) to all other cell types in PC space (as defined in Fig. 2). Left: normalized *D_phys_* between each cell type and all other cell types. Individual rows are labeled in Fig. S8. Right: functional elements common to cell types in each cluster, including a schematized center TRF and spatial receptive field. Entries are dashed (–) when members of a cluster do not show a common motif for a functional element. (E) Null model construction for tests of functional redundancy among inputs. The number of cell types in each cluster and the distributions of strong and weak input counts across cell types are used to simulate 10,000 postsynaptic cell types. Random draws from these distributions define the inputs to each synthetic cell type. (F) Cumulative probability density functions describing the number of functional clusters represented by strong (green) and weak (purple) inputs across real (solid lines) and synthetic (dashed lines) postsynaptic cell types. Strong inputs to real neurons come from significantly fewer functional clusters than expected by chance (K-S test, *p* < 0.001), and this difference persists over a range of strong/weak cutoff values (Fig. S8F). (G) Strong vs. weak input bias among functional types. For each cluster, the difference between the probabilities that a cell type in that cluster will be a strong or weak input is plotted.

To explore the privileged position of strong inputs in more detail, we considered two alternative models that quantitatively predict postsynaptic responses from presynaptic input (Fig. 6B). The first model uses the connection-weighted sum of all inputs to construct the post-synaptic TRF (“connection-weighted sum” model), while the second model only uses the single strongest input as the best predictor (“strong input dominant” model). To assess the prediction quality of the connection-weighted sum model, we calculated a connection-weighted sum of input TRFs for each cell type in the main dataset. We then found the correlation coefficient between this prediction and the actual postsynaptic TRF that we measured. Out of necessity, these predictions only considered presynaptic cell types that were also part of the main dataset, but those cell types collectively accounted for more than half of the total input synapses in most cases (50% confidence interval = 45–77%; Fig. S8B). On average, these connectome-weighted predictions performed no better than a simple correlation between a postsynaptic neuron and its single strongest input, the strong input dominant model (Fig. 6C, left). Indeed, 28 out of 43 cell types were well described by the strong input dominant model, while only four out of 43 cell types were better described by the connection-weighted sum model (Fig. 6B). Finally, we tested each strongest input’s unique contribution by removing it from the connection-weighted sum and re-calculating the cross-correlation (Fig. 6C, right). Under these conditions, prediction quality was significantly lower across cell types (*p* < 0.05), with 28 out of 43 types displaying a decrease in *R* value. Notably, removal of the strongest input did not reduce correlations to 0, consistent with a degree of functional redundancy among inputs. These results argue that the strongest input plays an outsized role in shaping postsynaptic responses for most cell types.

### Strong inputs to a given cell type tend to be functionally homogeneous

One simple reason why strong inputs to a given cell could be functionally redundant is because those inputs are physiologically similar to one another. To assess this model, we first computed the median position of each cell type in the functional PC space (Fig. 2), and found *D_phys_* between all pairs of cell types in the labeled dataset (Methods). We then used these distances to functionally cluster cell types. To remain agnostic to the number of functional subtypes (i.e., clusters), we opted for an affinity propagation algorithm, which determines an appropriate number of clusters based on sample variance (Frey & Dueck, 2007). This approach yielded 10 clusters, each containing cell types with relatively similar response properties (Fig. 6D and Fig. S8D-E). For example, cell types in cluster 0 tended to be slow, ON-selective neurons with some degree of UV selectivity, while cell types in cluster 8 tended to be OFF-selective neurons with large spatial receptive fields (Fig. 6D). To quantitatively assess the degree of functional redundancy among inputs, we defined a null hypothesis based on artificial “cell types” that stochastically receive input from functional clusters based on the relative size of each cluster (Fig. 6E). For the main dataset, we assessed the number of cell types in each functional cluster (Fig. S8E), as well as the number of strong (%_in_ > 5) and weak inputs (%_in_ < 5) onto each cell type. These two distributions form the statistical basis of the null model (Fig. 6E). For each synthetic postsynaptic neuron, we drew strong and weak inputs based on these distributions. Then, for each drawn input to the synthetic neuron, a cluster was probabilistically assigned based on the number of cell types in each cluster. We performed this sampling process for 10,000 synthetic cell types, forming the null distribution.

We then asked, for both the actual data and the null distribution, how many functional clusters are sampled by strong and weak inputs (Fig. 6F). As expected, the number of functional subtypes represented by strong inputs was generally smaller than that of weak inputs, consistent with the fact that neurons receive many more weak inputs than strong ones (Fig. 6E). However, even when the difference in the number of strong inputs was taken into account by the null model, inputs to real neurons were members of significantly fewer functional subtypes than expected by chance (Fig. 6F, *p* < 0.001). This functional homogeneity of strong inputs persisted across a wide range of strong/weak %_in_ thresholds (Fig. S8F). In contrast, the functional diversity of weak inputs was consistent with chance (*p* = 0.089). Thus, strong and weak inputs are categorically different in their degree of functional homogeneity.

### Similar connectivity does not predict similar physiology

One common intuition is that functionally similar cell types might share at least some synaptic partners. If this were true, connectivity patterns and physiological properties should predict one another. To quantitatively assess this bidirectional relationship, we began with measures of connectomic and functional similarity. As previously described by Matsliah and colleagues (2024), Jaccard distance (*D_conn_*) can quantify the connection similarity between two cell types. Cell types with identical inputs and outputs would have a *D_conn_* of 0, while types with completely different inputs and outputs would have a *D_conn_* of 1. As described above (Fig. 6), *D_phys_* can capture the Euclidean distance between cell types in a high dimensional space representing neuronal function. As with *D_conn_*, small *D_phys_* values correspond to functionally similar cell types, while dissimilar cell types have large *D_phys_* values. To directly compare *D_conn_* to *D_phys_*, we divided *D_phys_* values by the maximum type-to-type *D_phys_* we observed across the data, producing a normalized metric that ranged from 0 (most similar) to 1 (most different).

We first asked whether similar patterns of connectivity predict similar physiological responses (Fig. 7A-B, top row). We performed K-means clustering on *D_conn_* and sorted cell types within each cluster by median *D_conn_* to all other cell types in our main dataset (Methods). We then plotted *D_phys_* for these anatomically clustered and sorted cell types (Fig. 7B, top). If neurons with similar connectivity also tend to possess similar physiological responses, “blocks” of relatively low *D_phys_* should extend from the diagonal of the matrix. Instead, the resultant matrix displayed no apparent structure, indicating that similar connectivity is not associated with similar physiology.

**Figure 7.**
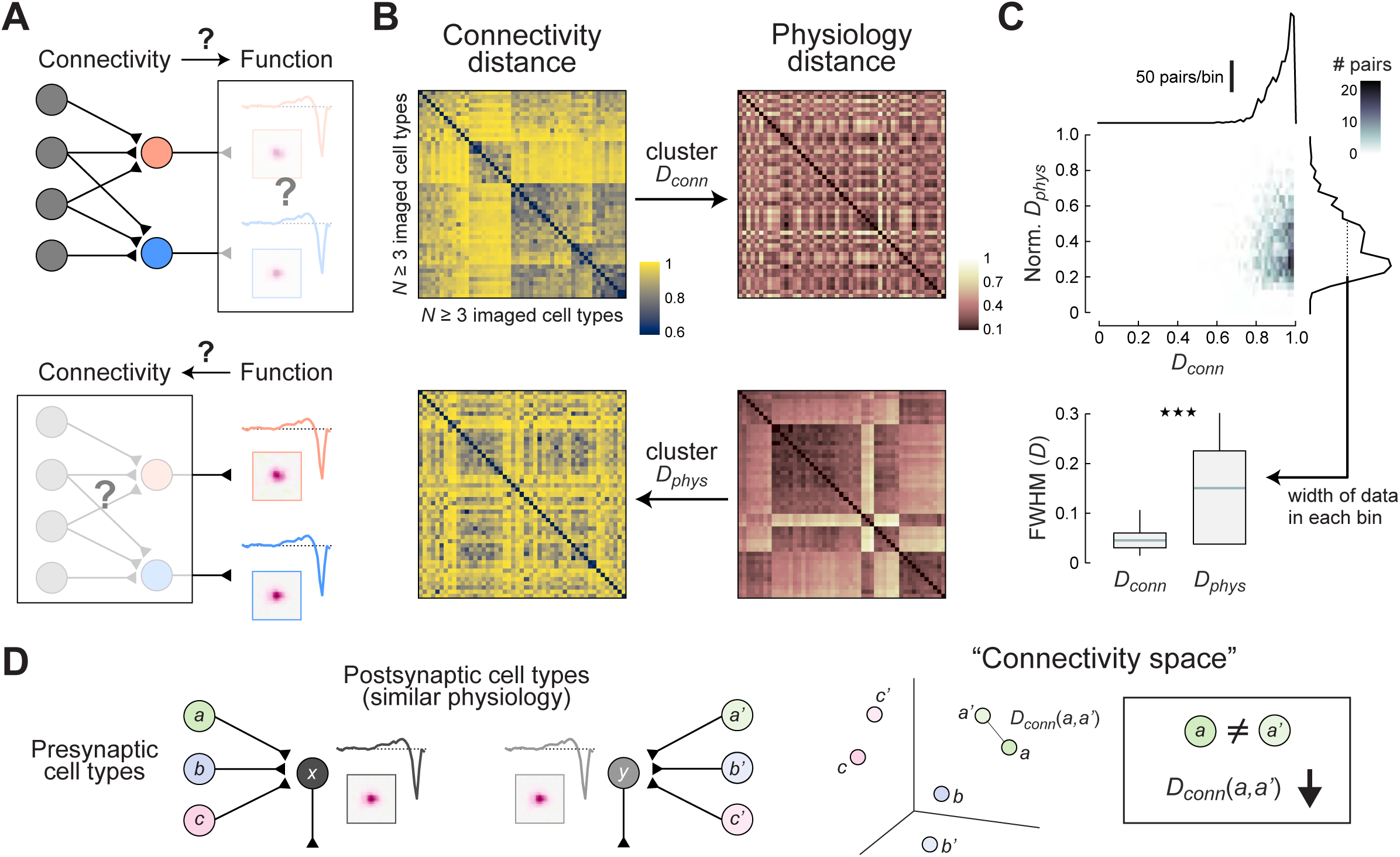
Similar connectivity does not predict similar physiology. (A) Connectivity-function hypotheses. Top: does similar connectivity predict similar physiology? Bottom: does similar physiology predict similar connectivity? (B) Tests of these models using the main dataset. Left column: matrices of type-to-type connectivity distances (*D_conn_*), as calculated in a recent connectome analysis (Matsliah et al., 2024). Right column: matrices of normalized *D_phys_*, as in Fig. 6D. Top row: K-means clustering performed on the *D_conn_* matrix is mirrored onto the *D_phys_* matrix. Bottom row: K-means clustering performed on the *D_phys_* matrix is mirrored onto the *D_conn_* matrix. (C) Top: *D_phys_* - *D_conn_* joint probability distribution. The marginal distributions of each distance metric are shown on the top and right of the joint distribution plot. Bottom: distributions of full-width at half-max values for marginals of each distance metric. Distributions are comprised of FWHM values evaluated at each *D_phys_* or *D_conn_* bin (Methods). Boxes illustrate the 75% CI, whiskers cover the full range of values. The *D_phys_* distribution covers a significantly wider range of values across bins (rank-sum test, *p* < 0.001). (D) Updated model of the connectivity-function relationship. In contrast to the model shown in (A), cell types with similar physiology (x, y) do not necessarily share input cell types (a, b, c and a’, b’, c’). Instead, inputs to cell types with similar physiology tend to be close to one another in a “connectivity space,” such that *D_conn_*(a,a’), *D_conn_*(b,b’), and *D_conn_*(c,c’) are low. However, no cell types are particularly close to one another in connectivity space, preventing physiology from being predicted from connectivity.

We next performed the inverse analysis: clustering and sorting based on *D_phys_*, then plotting *D_conn_* for physiologically clustered and sorted cell types (Fig. 7B, bottom). If neurons with similar physiological responses also tend to possess similar connectivity patterns, “blocks” of relatively high *D_conn_* should extend from the diagonal of the matrix. Strikingly, we observed considerable block structure, indicating that similar physiology is indeed associated with relatively similar connectivity.

To understand how function can predict connectivity, but connectivity cannot predict function, we examined the distributions of *D_conn_* and *D_phys_* (Fig. 7C). Critically, the *D_conn_* distribution is tightly clumped around 0.9, meaning that most cells have highly dissimilar connectivity patterns. In contrast, normalized *D_phys_* covers a broad range from 0.2 to 0.8, reflecting the mixture of functionally similar and functionally dissimilar cell types in this part of the visual system. We next sought to quantify the range of *D_phys_* at different *D_conn_* values and vice-versa. We calculated the full-width at half-max of *D_phys_* marginal distributions evaluated at different *D_conn_* values and *D_conn_* marginal distributions evaluated at different *D_phys_* values. As expected, the *D_phys_* distributions were significantly wider that the *D_conn_* distributions (*p* < 0.001). In order for bidirectional predictability to exist, the variance of these two distributions would have to be similar. The fact that *D_conn_* has much less variance than *D_phys_* explains the relative lack of predictive power for *D_conn_* – the connectivity of any given cell type is equally unlike that of all other cell types, so *D_conn_* has little potentially explanatory variance. Thus, structure is surprisingly incapable of predicting function at this level of resolution, while function can indeed predict structure across cell types.

## Discussion

Using a novel genetic strategy that does not rely on cell-type specific genetic access, we record the visual responses of 510 neurons from at least 91 anatomical cell types, establishing a new resource for vision science (Fig. 1-2, Fig. S1-S3). We show that some important visual features, such as orientation tuning, direction selectivity, and spectral preference, can be predicted from the connectome (Fig.3). Conversely, receptive field size cannot be predicted by a cell-type’s retinotopic coverage (Fig. 4). We also find that postsynapse density predicts both response amplitude, and the temporal correlation between pre- and postsynaptic partners (Fig. 5-6). Interestingly, postsynaptic responses are poorly described by an intuitive connection-weighted sum of input activity; instead, strong inputs have an outsized influence on postsynaptic activity (Fig. 6). Surprisingly, a clustering analysis reveals that similar connectivity cannot predict similar physiology, but similar physiology can predict similar connectivity (Fig. 7). This non-invertible relationship provides a new outlook on how wiring constrains function.

### A new approach for rapid physiological profiling

We demonstrate that SPARC-L is an efficient method for recording from a large variety of cell types. By sparsening indicator expression through two consecutive competitive recombination cassettes, we achieve labeling densities amenable to single cell recordings using genetic tools that target broad classes of cells. By adjusting the recombination efficiency of each SPARC element (Isaacman-Beck et al., 2020), we can achieve a wide range of expression densities, ranging from 2.5%-0.25% of cells, allowing labeling to be aligned with the density and diversity of a target neuropil. This tunability allowed us to achieve single-neuron resolution while also maximizing the number of neurons recorded in each fly. With around 35,000 neurons per hemisphere, the medulla is one of the densest neuropils in the fly brain (Matsliah et al., 2024; Nern et al., 2024); our success in this region implies that this approach will be highly generalizable. Finally, as this approach does not require the identification of driver lines that target each cell type of interest, it is particularly useful for exploring as-yet undescribed circuits.

### Observing and predicting feature selectivity across cell types

The medulla is the last neuropil that all eye-derived visual signals pass through, and our imaging data represent the most comprehensive description to date of signal processing in this region. Intriguingly, our principal components analysis suggests that cell types are not evenly distributed across function space (Fig. 2). This observation raises the possibility that specific combinations of features might systematically co-vary as a prelude to downstream processing. In any case, our data covers a substantial fraction of the cellular diversity of the medulla and future models of higher visual representation will benefit from our description (Currier, Pang & Clandinin, 2023).

Our characterization of visual encoding across many cell types also allowed us to test several connectome-based hypotheses pertaining to feature selectivity (Fig. 3). For example, Dm3 and TmY9 have elongated arbors that draw feedforward input across columns, and were predicted to be OS (Seung, 2024b). In our data, Dm3 and TmY9 were indeed OS, as were several other cell types. Similarly, the Y family of neurons receive substantial input from motion-selective cells, and were therefore predicted to be DS (Shinomiya et al., 2022). We recorded from one such cell type, Y13, and observed strong DS. Finally, a connectome analysis focusing on cell types downstream of spectrally selective photoreceptors suggested cell types that might display color preference (Kind et al., 2021). In our main dataset, Dm9 and Tm29 both displayed spectral preference, as predicted. However, several other cell types, including L1, C2, Dm2, Mi9, and Mi15 did not display spectral preferences, perhaps reflecting the particular wavelengths of light we used. These results show that particular visual response properties largely follow connectomic predictions.

At the same time, several recent studies have used the retinotopic coverage of a cell type’s neurites as a proxy for receptive field size (Zhao et al., 2023; Seung, 2024a; Garner et al., 2024; Ganguly et al., 2024). Our results conflict with this assumption, showing no significant relationship between arbor size and the area of the receptive field center (Fig. 4). In our data, neurons with small arbors sometimes had larger than predicted receptive fields, and neurons with large arbors often had smaller than predicted receptive fields. These two types of prediction errors might arise from two distinct mechanisms. In the former case, neurons with small arbors could draw input from cells covering larger portions of retinotopic space, enlarging the observed receptive field. In the latter case, neurons with large arbors could be compartmentalized (Yang et al., 2016; Meier & Borst, 2019). In these cases, input signals fail to spread uniformly across the entire dendrite, leading to calcium activity reflecting only local input. For example, the ability of electrical signals to spread can depend on neurite morphology, as electrical signals spreading more readily along “lower resistance” (thicker) processes (Jack, Noble & Tsien, 1975). Indeed, this phenomenon has been predicted for cell types with extremely thin neurites connecting different visual columns (Seung, 2024a). As our measurements were designed to capture local changes in intracellular calcium, our data reveals these processing compartments. As each output synapse responds only to local calcium signals, these compartmentalized receptive fields are the relevant descriptor of the spatial information passed to downstream neurons.

### Improving future connectome-inspired models

One of the goals of our work was to improve the accuracy of future connectome-based models, further enhancing the value of these important resources. Toward this end, we observed a categorical difference between strong and weak inputs, establishing a “strong input dominant” model as a better framework for generating quantitative connectome-inspired predictions (Fig. 6). We also found that strong inputs to a given cell type tend to be more functionally homogeneous than expected by chance. Collectively, these results paint a cohesive picture of functional subtype membership by presynaptic cell types – if strong inputs to a cell type are likely to be functionally similar, a large percentage of total input can be described by the single strongest input. Under these circumstances, the strongest input would be expected to be highly correlated with the postsynaptic response. Strong input homogeneity might also be described as functional redundancy, consistent with the relatively modest effects that excluding the single strongest input had on our correlation analysis. The “strong input dominant” model represents a new quantitative framework for predictive connectomics, describing synaptic inheritance better than a simple connection-weighted linear sum of inputs.

Viewed from another angle, the predictive power of the “strong input dominant” model suggests that weak inputs have a sub-linear impact on postsynaptic responses. Many recent connectome analyses have established 5-10 synapses as a “cutoff” for excluding the brain’s numerous weak connections (Zhao et al., 2023; Dorkenwald et al., 2024; Lin et al., 2024; Dombrovski et al., 2023; Garner et al., 2024; Ganguly et al., 2024; Schretter et al., 2024). Surprisingly, our results imply that this cutoff is conservative. At least in the medulla, we found that key elements of computation are not influenced by weak connections (%_in_ < 5), neither individually nor collectively. Utilizing a higher, input fraction-based thresholding process might further improve the reliability, and reduce the complexity, of future connectomic predictions. This sub-linear functional influence of weak connections is consistent with the greater variability seen in such connections across connectomes (Takemura et al., 2015; Cornean et al., 2024). Given that a very large fraction of total connections are “weak” (Matsliah et al., 2024; Nern et al., 2024; Fig. S8A), the role that these weak connections might play remains mysterious, and an important subject for future research.

### A revised model of connectomic constraints on physiology

We also considered the common assumption that neurons with similar connectivity are likely to have similar functions (Fig. 7). However, after clustering cell types by their connectivity, and examining their function, we found that similar connectivity patterns were not correlated with similar functions. Prior work used the same kind of clustering to define computational subnetworks, groups of cell types that process motion, color, and object features (Matsliah et al., 2024). Integrating these findings with ours, we infer that within each subnetwork, the details of a cell type’s physiology, such as ON/OFF selectivity, response speed, spatial organization, and receptive field size, cannot be accurately judged from connectivity alone.

Surprisingly, after clustering cell types by their function and examining their connectivity, we found that similar functional properties were indeed associated with relatively similar wiring patterns. This relationship relies on a high degree of informative variance in the distribution of physiological distances between cell types. At the same time, connectomic studies have revealed that the connectivity matrix of the brain is sparse (White et al., 1986; Helmstaeder et al., 2013; Motta et al., 2019; Winding et al., 2023; Vishwanathan et al., 2024; Dorkenwald et al., 2024; Nern et al., 2024; Fig. S8). This sparsity presents challenges to connectome-based predictions, because the number of shared presynaptic elements between any pair of cell types is often small. Stated differently, no two cell types are particularly close to one another in “connectivity space” – instead, all cell types are similarly distant from one another. So how can functional similarities predict wiring similarities?

Common intuition argues that neurons with similar physiological properties derive them from sharing very similar sets of inputs. However, the ubiquity of dissimilar connectivity suggests that, within a particular functional cluster, there are few (if any) shared presynaptic cell types (Fig. 7D). Rather than neurons with similar function sharing input from the same cell types (as schematized in Fig. 7A), our data suggest that inputs to functionally similar cells are instead distinct cell types that lie relatively close to one another in a high-dimensional “connectivity space” (Fig. 7D). Consider two postsynaptic cell types x and y with similar physiology that draw input from cell types a, b, c and a’, b’, c’, respectively. While a and a’ are not the same cell type, a is closer to a’ in connectivity space than it is to b (Fig. 7D, right). In this way, two cell types can have similar physiology and relatively similar connectivity without sharing input cell types. Overall, this analysis suggests that defining connection similarity by cell types may overly discretize the network, obscuring structure-function relationships.

### Specificity and generality in connectomic predictions

Connectomes are powerful sources of functional hypotheses that may or may not incorporate additional information, such as prior physiological recordings. When functional data is available, connectomes can provide mechanistic insights by examining granular circuit properties, such as the precise spatial position, identity, and variability of presynapses (Briggman, Helmstaedter & Denk, 2011; Kim et al., 2014; Takemura et al., 2017; Liu et al., 2022; Dombrovski et al., 2023; Cornean et al., 2024). At the other extreme are studies that exclusively utilize connectome data, or draw minimally from outside constraints, to model circuits, systems, or even whole brains (Zhao et al., 2023; Lappalainen et al., 2024; Shiu et al., 2024; Lin et al., 2024; Pospisil et al., 2024). While this range of uses robustly demonstrates the versatility of connectomic data, it also raises the question of how a model could best incorporate all of the information available in a connectome. Our work argues that large-scale connectome models that draw sparingly from physiological constraints are likely to miss many important functional properties, while capturing a gestalt. From this perspective, the relationship between structure and function in a large-scale connectome model is perhaps best appreciated as a pointillist painting, obscuring functional details but capturing a unified whole.

## Acknowledgements

We would like to thank Jesse Isaacman-Beck, Ina Anreiter, and Carl Wienecke for fly resources; Michael Reiser and members of the Reiser Lab for providing early access to male optic lobe connectome data and for sharing summary metrics from their analyses; Sebastian Seung for sharing type-to-type Jaccard distances for FAFB optic lobe cell types; Janina Troper for python code to scrape connectome data from websites; and Marion Silies and members of the Clandinin Lab for thoughtful feedback and discussion on the manuscript. This work was supported by T32EY027816 and F32EY035135 to TAC and R01E022638 and P30EY026877 to TRC. TRC is an Investigator of the Chan-Zuckerberg Biohub.

## Resource availability

Calcium imaging data will be made available on Dryad at the time of publication. The new lexAop-SPARC plasmid backbone will be deposited at Addgene, and all new fly strains will be deposited at the Bloomington *Drosophila* Stock Center. All software and analysis code can be found on github: stimulus presentation and experimental protocols were implemented with flystim (https://github.com/ClandininLab/flystim) and visprotocol (https://github.com/ClandininLab/visprotocol). The functionality of these software packages has also been integrated into the stimpack software suite (https://github.com/ClandininLab/stimpack). Initial curation of calcium imaging data, including motion correction, ROI definition, and dF/F computation, was performed with visanalysis (https://github.com/ClandininLab/visanalysis) and brainsss (https://github.com/ClandininLab/brainsss). All other analyses, including STRF calculation, visual response summary generation, and connectome-physiology correlations, can be found on TAC’s personal GitHub page (https://github.com/tacbox3/CurrierClandinin/tree/main/CurrierClandinin). This repository includes top-level scripts that interact with visanalysis and brainsss.

## Author Contributions

Conceptualization, TAC; Methodology, TAC and TRC; Investigation, TAC; Software, TAC; Formal Analysis, TAC; Visualization, TAC; Writing – Original Draft, TAC; Writing – Review & Editing, TAC and TRC; Funding Acquisition, TAC and TRC; Supervision, TRC.

## Supplemental Informtion

Document S1. Supplemental Figure S1.

Document S2. Supplemental Figure S2.

Document S3. Table S2 and Supplemental Figures S3-S8.

Table S1. Excel file, related to Figure 1. Genotypes from which all cell types were recorded.

## STAR Methods

### Experimental model and subject details

All flies were raised on a standard molasses-based food at 25° C and 40% relative humidity, under a strict 12:12 light:dark cycle. Experimental flies were aged 7-11 days before imaging. All strains used in this study are listed in the Key Resources Table.

For new fly strains, all plasmids were generated through synthesis and cloning (GenScript Biotech), and were sequenced via single primer extension (Sequetech). GCaMP8m and GCaMP8m-P2A-myr::tdTomato were cloned into SPARC2-I or SPARC2-S near-VK00005 donor backbones (Isaacman-Beck, 2020) via SalI restriction digest or Genscript’s CloneEZ method. Transgenic flies were created through CRISPR-HDR by co-injecting a SPARC donor plasmid (100-500 ng) and a pCFD5 backbone carrying near-VK00005 gRNA (75-250 ng) into *nos*-Cas9 embryos (BestGene). We created the following lines:

*w-; +; 13xLexAop-SPARC2-I-GCaMP8m*

*w-; +; 13xLexAop-SPARC2-S-GCaMP8m*

*w-; +; 13xLexAop-SPARC2-I-GCaMP8m-P2A-myr::tdTomato*

*w-; +; 13xLexAop-SPARC2-S-GCaMP8m-P2A-myr::tdTomato*

These lines were combined with existing *+; 20xUAS-SPARC2-I-LexA* or *+; 20xUAS-SPARC2-S-LexA* lines to create SPARC-L base stocks, which we will deposit at the Bloomington *Drosophila* Stock Center. We then crossed SPARC-L base stocks to T2A knock-in drivers of neurotransmitter synthesis or vesicular transport proteins (Deng et al., 2019) with pan-neuronal ΦC31 recombinase (i.e., *nsyb-ΦC31; VGluT-T2A-GAL4; +*) for imaging. For a small number of flies, a *GAL4* enhancer trap insert at the *apterous* locus was used in place of a neurotransmitter-specific driver (Calleja et al., 1996). Table S1 lists the full genotypes from which each cell type in the dataset were recorded.

### Method details

#### Animal preparation and 2-photon calcium imaging

Imaging experiments followed protocols used previously in the Clandinin Lab (Turner et al., 2022; Brezovec et al., 2024). Female flies were cold anesthetized and mounted with a ∼20° roll-axis tilt into a custom-cut hole in a thin aluminum shim at the bottom of a round imaging dish. This tilt positions the antero-ventral aspect of the left eye toward the stimulus screen.

Mounted flies were fixed in place with two small drops of UV-curing glue: one around the head, and one at the notum. The cuticle above the left optic lobe on the posterior side of the head was removed with a fine-gauge needle, and the fat and trachea covering the brain were cleared away with forceps. After preparation, imaging saline (103 mM NaCl, 3 mM KCl, 5 mM TES, 1 mM NaH_2_PO_4_, 4 mM MgCl_2_, 1.5 mM CaCl_2_, 10 mM trehalose, 10 mM glucose, 7 mM sucrose, and 26 mM NaHCO_3_) was bubbled with carbogen (95% O_2_, 5% CO_2_) and continuously perfused at room temperature into the imaging dish.

We imaged the left optic lobe (Fig. 1B) of flies under a Bruker Ultima IV resonant scanning 2-photon microscope with a 20x HCX APO 1.0 NA water immersion objective (Leica). A fast piezo-controlled Z drive manipulated the focal plane during volumetric scanning. The tissue was stimulated with a Mai Tai titanium sapphire laser (Spectra Physics) at 920 nm and 15-20 mW, and emitted photons were collected in two GaAsP-type photomultiplier tubes (PMTs) after filtering into red (595/50) and green (525/50) channels, allowing us to capture tdTomato and GCaMP8m fluorescence, respectively. Each fly had a slightly different imaging volume depending on its pattern of GCaMP expression. For functional scans, we performed volumetric imaging of the superficial-most 60-70 nm of the posterior medulla with a 222.8 μm x ∼120 μm field of view, which contains neurons downstream of photoreceptors on the antero-ventral aspect of the left eye. An imaging rate of 3.5-4.2 Hz was achieved with a voxel resolution of 0.87 x 0.435 μm for z-slices 3-4 μm apart; we recovered X-Y pixel isotropy during postprocessing (see below). For each fly, we used the same imaging volume and stimulation parameters for all three visual stimulation blocks described below.

After all functional scans were completed, the perfusion temperature of the saline was lowered with an in-line chiller to approximately 10° C. After a 3 min equilibration period, this reduced temperature had two effects: first, it dramatically amplified GCaMP8m fluorescence due to increased intracellular calcium; and second, it greatly reduced brain movement. The greater tissue stability and stronger baseline fluorescence allowed us to collect a high-resolution volume of the posterior two-thirds of the whole optic lobe in galvanometer scan mode with a long (3 μs) dwell time. Voxel resolution in these images was increased to 0.435 x 0.435 for z-slices 0.7 μm apart. A full experiment of three functional scans and one anatomical scan lasted 45 min.

#### Visual stimulation

We back-projected visual stimuli onto a cloth screen with a LightCrafter 4500 Pro that was modified to replace the factory red LED with a UV LED (Wintech). Stimulus light was passed through a 0.6 A-coated absorptive neutral density filter (Thorlabs) and a 390/482/587-25 multi-notch filter (Chroma) to reduce stimulus intensity and limit bleed-through into the PMTs, respectively. The stimulus-containing portion of the screen subtended approximately 80° of visual angle in elevation and azimuth. As previously described (Turner et al., 2022), visual stimuli were presented with flystim, a python and openGL-based software package that renders perspective-corrected images in real time. Throughout each stimulus presentation block, a photodiode recorded precise frame timing relative to imaging samples.

The command voltage to each LED was adjusted so that activation of Rh1-containing photoreceptors (R1-R6) was matched across stimulus colors. To do this, we multiplied the Rh1 absorption spectrum (Sharkey et al., 2020) by the LED emission spectrum, as measured at the eye during equal drive with a USB4000 Ocean Optics spectrophotometer. We then found the scale factor that equalized the areas under these curves. This scale factor was applied to the command voltages to normalize total Rh1 drive during blue and UV stimulus presentation. Under these conditions, peak photon flux was found to be 0.9 x 10^15^ photons/(cm^2^/s) for UV stimuli and 1.8 x 10^15^ photons/(cm^2^/s) for blue stimuli.

Each recording session consisted of two blocks of noise presentation followed by one block of full-screen flicker presentation. Each 16 min noise session was presented with a single LED: first blue, then UV (Fig. 1D). For the 6 min full-field flicker session, both Rh1 activation-matched LEDs were driven together.

For noise stimuli, we presented ternary 2-dimensional noise, divided into 20 “trials” of 45 s, each separated by 3 s of mean gray. The first 3 s of each trial was discarded during analysis to account for these gray interludes. Each patch in the noise grid subtended 5° of visual angle, approximating the coverage of a single medulla column. The full grid was 16 x 16 patches, for the full 80° x 80° screen coverage. With ternary noise, each patch displays one of three intensity values at each time point: mean gray (0.5), full brightness (1), or dark (0.01). The spatial and temporal sequence of patch contrasts is drawn randomly. Noise was uniquely generated for each trial in each fly, and updated at a rate of 20 Hz.

In each flicker trial, 2 s of mean gray covering the full 80° x 80° screen preceded sinusoidal intensity fluctuations, which began at mean gray and increased to full brightness, then decreasing to 0.1% brightness, before returning again to mean gray (Fig. S3B). 10 s of sinusoidal fluctuations were presented, followed by 2 s of post-stimulus mean gray. Four intensity oscillation frequencies (0.1 Hz / 1 cycle; 0.5 Hz / 5 cycles; 1 Hz / 10 cycles; 2 Hz / 20 cycles) were pseudo-randomly presented, with five repetitions of each frequency.

Across the dataset, we found neurons covering the full 80° of screen azimuth. However, the 10-15° uppermost portions of the screen were not observed to be a part of any cell’s receptive field, suggesting that the “working” dimensions of our stimulus was 80° azimuth by 65-70° elevation. From the fly’s perspective, the top part of the screen may have been occluded by the shim or mounting dish.

#### ROI selection

Functional scan data was preprocessed as follows: first, with ANTs-based motion correction (Avants, Tustison & Song, 2009; Brezovec et al., 2024). Second, X-Y pixels were made isotropic by averaging over every two pixels in Y, maximizing functional scan signal-to-noise ratio. Finally, we calculated the Fano factor of each voxel in motion-corrected volumes, then thresholded these “Fano factor maps” to select proto-regions of interest (ROIs). The Fano factor is a measure of variance over mean, with “hot spots” in the Fano map corresponding to pixels with high variation in fluorescence intensity over the course of stimulus presentation. Usually, a high Fano factor was associated with stimulus evoked calcium activity.

Using this Fano map as a starting point, ROIs were manually curated to maximize response strength. All ROIs were defined in single Z-planes (ROIs were not volumetric). We used the following guidelines when defining ROIS: (1) the ROI must unambiguously contain the neurites of only one cell. (2) The ROI should not cover more than ∼3 columns of retinotopic space, unless a larger ROI is required to recover a signal from a given neuron. This exception was invoked for fewer than 3% of ROIs in the dataset. (3) If a neuron shows discrete proximal-distal “compartments,” ROIs should contain only one compartment, unless a larger ROI was required to recover a signal from a given neuron. This exception was invoked in fewer than 1% of ROIs. Due to the 20° roll-axis tilt of flies in our experiments, there was often a slight shadow cast by the posterior-most part of the eye over the medulla, impacting distal layers more than proximal ones. For this reason, the best-responding compartment was usually the most proximal portion of each cell. (4) If a single neuron contains multiple discrete proto-ROIs in the Fano brain, multiple ROIs are only kept if there was a qualitative difference between them. Otherwise, only the single best-responding ROI was retained.

Stimulus-responsive ROIs were usually spatially clustered, consistent with the medulla’s retinotopic arrangement. Instances of non-responsive neurons co-mingled with responsive neurons were rare, but did happen; we believe these neurons were not well-driven by our stimulus. Generally, unresponsive neurons were large, covering many medulla columns. In at least 10 noise-unresponsive neurons, we did find a measurable response to full-field flashes. This was only formally tracked while analyzing the final ∼200 neurons collected in the dataset. By extrapolation, we can estimate that cells poorly driven by our noise stimulus make up approximately 5% of the medulla population.

#### Anatomy-based cell type identification and functional proofreading

For each ROI, we manually identified its cell type based on morphological characteristics. We compared the high-resolution anatomy scan to morphological cell type atlases (Fischbach & Dittrich, 1989; Morante & Desplan, 2008; Nern et al., 2015) and connectome datasets (Matsliah et al., 2024; Nern et al., 2024). We used the baseline-enhanced GCaMP8m signal (see above) along with the myristoylated tdTomato signal, when it was available. We also used the Fano map to assist in identification when non-ROI cells were dense in the vicinity of the ROI, because it made clear which neurites belong to the ROI-containing cell. Key properties that aided identification, beyond the medulla innervation pattern, were: (1) neurotransmitter identity (based on the Gal4 driver used), (2) soma location, when it could be determined, (3) projection patterns of the cell toward the lamina, or in the lobular complex. We also noted that, if a cell type was labeled in a particular brain, there tended to be multiple copies of that type labeled in that same brain; this is likely a consequence of SPARC recombination timing relative to the medulla development timeline. Cell type labels for each ROI were only retained when they met a high level of qualitative confidence.

After all ROIs had been assigned a cell type, we performed a functional proofreading process to eliminate bad IDs. For cell types with *N* = 3 or more, we could identify outlier responses in PC space (i.e., Fig. 2B and Fig. S7C, see below). These outliers represented approximately 5% of the ROIs, and most were removed from the labelled dataset, while the rest were re-examined morphologically and reassigned. Because cell types with *N* < 3 did not benefit from functional proofreading, we excluded these cell types from group analyses. This exclusion produced a “main” dataset of 43 cell types, which is a subset of the “labeled” 91-cell type dataset.

### Quantification and statistical analysis

#### Analysis of visually evoked calcium signals

For each ROI, raw fluorescence intensity was converted into dF/F using visanalysis (Turner et al., 2022). We regenerated the presented noise stimulus, and used the photodiode data to tether each imaging sample to the closest frame update time. For each imaging frame, we weighted the prior 3 s of stimulus history by fluorescence intensity at that frame. We then summed over all imaging frames to yield an STRF in units of dF/F. Because stimulus frames were updated at 20 Hz (see above), the Nyquist resolution limit of our STRFs was 10 Hz. We *z*-scored STRFs patch-by-patch by mean-subtracting and dividing by response variance in the first 1 s of a patch’s temporal filter. No neuron had signal in this early portion of the STRF, so *z*-scores are in units of baseline variance. Throughout this pipeline, blue and UV noise responses were handled separately.

To center each cell’s STRFs, we first identified the max-responding patch across blue and UV noise responses. We then thresholded the STRF for the max-responding color at *z* = 0.5 or -0.5, based on the sign of the max responding patch. *Z*-scores between -0.5 and 0.5 represent the noise floor for this dataset (Fig. S3C). We integrated the thresholded STRF over the period defined by the last temporal lobe of the max responding patch, yielding a thresholded SRF. We identified the strongest spatially contiguous feature within this SRF, then found its center of mass (CoM), which we centered by spatially shifting a 0-padded version of the full STRF. The same shift was mirrored onto the STRF of the non-max-responding color (that is, centering was performed on a cell-by-cell basis, rather than an STRF-by-STRF basis).

For each color’s STRF for each neuron, we define the center TRF as the time-course of the max-responding pixel. The largest spatially contiguous portion of the non-thresholded mean SRF (with a sign opposite the max-responding patch) is defined as the surround for each neuron. We calculate the surround TRF as the mean response of surround patches over time. Finally, radial slices through the mean SRF are averaged to produce a radial RF (Fig. S3A).

For flicker data, we calculated a stimulus-triggered average over repeated presentations of each flicker frequency. CoM-aligned STRFs and mean flicker responses were averaged across cells to yield a unique response profile for each cell type, as in Fig. 1G and Figs. S1 & S2.

#### Principal components analysis and D_phys_

Our principal components analysis utilized the “full” dataset. For each neuron, we vectorized (flattened) and concatenated the center 40° and last 1 s of the blue and UV STRFs. This spacetime period contains nearly all signal present in STRFs across cells. We also concatenated the mean dF/F response to each flicker stimulus. The flattened and concatenated STRFs and flicker responses defined one sample for the PCA. After fitting the PCA model, resulting eigenvectors had this same flattened and concatenated structure. To visualize eigenvectors (Fig. 2A), we de-vectorized and de-concatenated them (Fig. S6). Each PC therefore had a blue STRF portion, a UV STRF portion, and a set of flicker portions.

To calculate Euclidean distance in PC space (*D_phys_*), we first projected each neuron into the space defined by the first 100 PCs. We then found the length of the vector connecting a pair of cells (Fig. 2B) or a pair of cell type medians (Fig. 6&7) in this space. For the clustering analysis in Fig. 6&7, we normalized *D_phys_* by dividing all type-to-type *D_phys_* by the largest observed type-to-type *D_phys_*.

#### OS and DS estimation

Complex spatiotemporal features were not necessarily densely sampled by our 2D noise stimulus. In cases where single-color STRFs might not capture a response feature (OS and DS), we calculated a joint-color STRF by averaging over blue and UV STRFs. In these cases, individual neurons with strong spectral preference, as calculated by SPI (Fig. 3G) were excluded.

Similarly, we reasoned that OS and DS are ill-defined in neurons representing small visual areas. Our stimulus patches were each 5° x 5°; aliasing along both dimensions means that, in the worst case, a neuron representing a single ∼5° column could show a receptive field center area as large as 100 deg^2^ (corresponding to a 10° x 10° receptive field). For this reason, individual neurons with SRF center areas smaller than 110 deg^2^ were removed when calculating OSI or DSI. SRF center areas were calculated based on an ellipse fitted to the thresholded SRF using the skimage package (see Fig. 3A). Cell types that failed to meet the N ≧ 3 criterion with spectrally selective or small SRF neurons omitted were not considered in group analyses (Fig. 3C&F).

For the OSI calculation, we processed the joint-color STRF the same as the single color STRFs (Fig. S3A), defining a thresholded SRF. We fit an ellipse to the central spatially contiguous feature of the thresholded SRF, as described above for SRF center area calculation. OSI was defined as the ratio of the fitted ellipse’s long axis over its short axis.

Because DS cannot necessarily be inferred from the spatiotemporal structure of a receptive field, we simulated responses to moving sinusoidal gratings. As an additional correction for potential under-sampling of complex spatiotemporal features, we smoothed our STRFs before simulation: temporally with a 5-sample boxcar (down-sampled to 4 Hz), and spatially with a 2D Gaussian (σ = 2°). We convolved these spatiotemporally smoothed joint-color STRFs with moving sinusoidal gratings (spatial period = 80°, temporal period = 1 s). Grating parameters were chosen based on T4/T5 tuning (Maisak et al., 2013; Fisher, Silies & Clandinin, 2015). Each grating moved in one of eight directions (cardinal and off-cardinal axes), with the grating’s isointensity lines oriented perpendicular to the direction of motion. Each artificial stimulus lasted 10 s (or 10 cycles at 1 Hz), with the steady-state convolution product (see Fig. 3D) defined as the 6^th^ cycle response – convolutional edge effects are observed on cycles 1-3 and 7-10 due to the 3 sec duration of the STRF. Steady-state responses were vectorized and the PD was identified as shown in Fig. 3D. DSI was formally defined as the length of the PD-ND difference vector divided by the sum of the lengths of the individual PD and ND vectors.

#### Connectome analyses

Unless otherwise specified, all connectome summary variables used in this study were originally calculated in a male optic lobe connectome (Nern et al., 2024) using FAMB data version 1.0. Input fraction data used for connectome-weighted sum and functional redundancy modeling (Fig. 6 and Fig. S8) were scraped from the Reiser lab cell type explorer (https://reiserlab.github.io/male-drosophila-visual-system-connectome/), drawing from FAMB. Type-to-type Jaccard distances in connectivity space (Fig. 7) were calculated as part of a recent female optic lobe analysis (Matsliah et al., 2024) using FAFB data version 783 (Dorkenwald et al., 2024; Shlegel et al., 2024). All single neuron reconstructions (Fig. 1G, 4A, 5A-B) were rendered in 3D with Neuroglancer using FAFB data version 783 on <u>codex.flywire.ai</u>. Table S2 lists the individual cell IDs for each example neuron shown.

To test for relationships between a physiological variable and a connectome property, we found the correlation coefficient and best linear fit between these metrics across cell types. For all physiology-connectome correlations, we utilized the “main” dataset. While we use the within-type mean for connectome properties, which are well sampled in the FAMB dataset, we opted for the within-type median for physiological variables, as they were sampled as few as 3 times for some cell types. To test each correlation for significance, we resampled median functional variables 10,000 times with replacement, recalculating the correlation coefficient each time. Correlations were considered significant when the data’s true *R* value was above the 97.5^th^ percentile of this resampled *R* distribution. Resampling with replacement reflects a conservative strategy, as it creates a wider range of possible *R* values than a simple shuffle without replacement.

#### Clustering of type-to-type physiological and connectivity distances

For our initial classification of functional subtypes, we utilized an affinity propagation clustering algorithm, which does not take the number of desired clusters as an input, but rather determines an appropriate number of clusters based on sample variance (Frey & Dueck, 2007). We clustered all 91 cell types in the “full” dataset using type-to-type distance in PC space (*D_phys_*), giving an overall picture of functional variation. However, we only use the 43 “main” dataset cell types for subsequent analysis and modeling (Fig. 6&7).

To avoid asymmetries that might bias our bidirectional clustering analysis (*D_conn_* to *D_phys_* and *D_phys_* to *D_conn_*, Fig. 7), we switched to a *K*-means clustering algorithm. To determine an appropriate number of clusters for the “main” dataset, we first used affinity propagation on type-to-type *D_conn_* and *D_phys_*, yielding optimal cluster counts of 9 and 5, respectively. We applied the less stringent 5-cluster result to *K*-means clustering in *both* directions, to most fairly assess the predictive capacity of *D_conn_* and *D_phys_*. Using 9 clusters in both directions, instead of 5, did not meaningfully impact the results. Within each cluster, cell types were sorted in descending order of mean type-to-type distance. The exact same clustered and sorted order from one variable (*D_conn_* or *D_phys_*) was applied to the other to generate the matrices in Fig. 7.

#### Statistics

All statistical comparisons, except for resampled correlation coefficients (see above), were made with the *SciPy* package using non-parametric tests and *α* = 0.05. For matched sample tests of equal medians (Fig. 6C), we used the Wilcoxon signed-rank test. For unmatched sample tests of equal medians (Fig. 2B, 6A, 7C and Fig. S8A), we used the Mann-Whitney *U* (rank sum) test. Finally, for comparisons of one-dimensional probability distribution shapes (Fig. 6F and Fig. S8F), we used the Kolmogorov-Smirnov (K-S) test. When the same data was compared at multiple thresholds (Fig. S8A,F), *α* was Bonferroni corrected according to the number of thresholds tested.

### Key Resources Table

| REAGENT or RESOURCE | SOURCE | IDENTIFIER |
| --- | --- | --- |
| Fly strains |  |  |
| {20XUAS-SPARC2-S-LexA::p65}AttP40 | Isaacman-Beck et al., 2020 | RRID:BDSC_84142 |
| {20XUAS-SPARC2-I-LexA::p65}AttP40 | Isaacman-Beck et al., 2020 | RRID:BDSC_84141 |
| {nsyb-ΦC31}AttP18 | Isaacman-Beck et al., 2020 | RRID:BDSC_84150 |
| VGlut1-T2A-GAL4 | Deng et al., 2019 | RRID:BDSC_84697 |
| VGAT-T2A-GAL4 | Deng et al., 2019 | RRID:BDSC_84696 |
| ChAT-T2A-GAL4 | Deng et al., 2019 | RRID:BDSC_84618 |
| $P\{w[+mW.hs]=GawB\}ap^{md544} (ap-GAL4)$ | Calleja et al., 1996 | RRID:BDSC_3041 |
| {13xLexAop-SPARC2-I-GCaMP8m}VK05 | This paper | TBD |
| {13xLexAop-SPARC2-S-GCaMP8m}VK05 | This paper | TBD |
| {13xLexAop-SPARC2-I-GCaMP8m-P2A-myr::tdTomato}VK05 | This paper | TBD |
| {13xLexAop-SPARC2-S-GCaMP8m-P2A-myr::tdTomato}VK05 | This paper | TBD |

