## Supplementary material for "Infrequent strong connections constrain connectomic predictions of neuronal function": Figure S1

**Figure S1.** Related to Figure 1. *Visual response profiles for cell types in the main dataset*. Each page displays mean data +/- SEM for a single cell type, including the last 0.5 s and middle 40º of the blue and UV STRFs, full-field sinusoidal flicker responses, center and surround TRFs for blue and UV, mean SRFs for blue and UV, and radial spatial receptive fields for blue and UV. The difference between blue and UV TRFs for center and surround are also shown. Because blue and UV light intensities equally excite R1-R6, this difference specifically reflects signals inherited from inner photoreceptors (Methods). Details about how each plot is calculated can be found in Fig. S3. Main dataset cell types are used for most analyses.

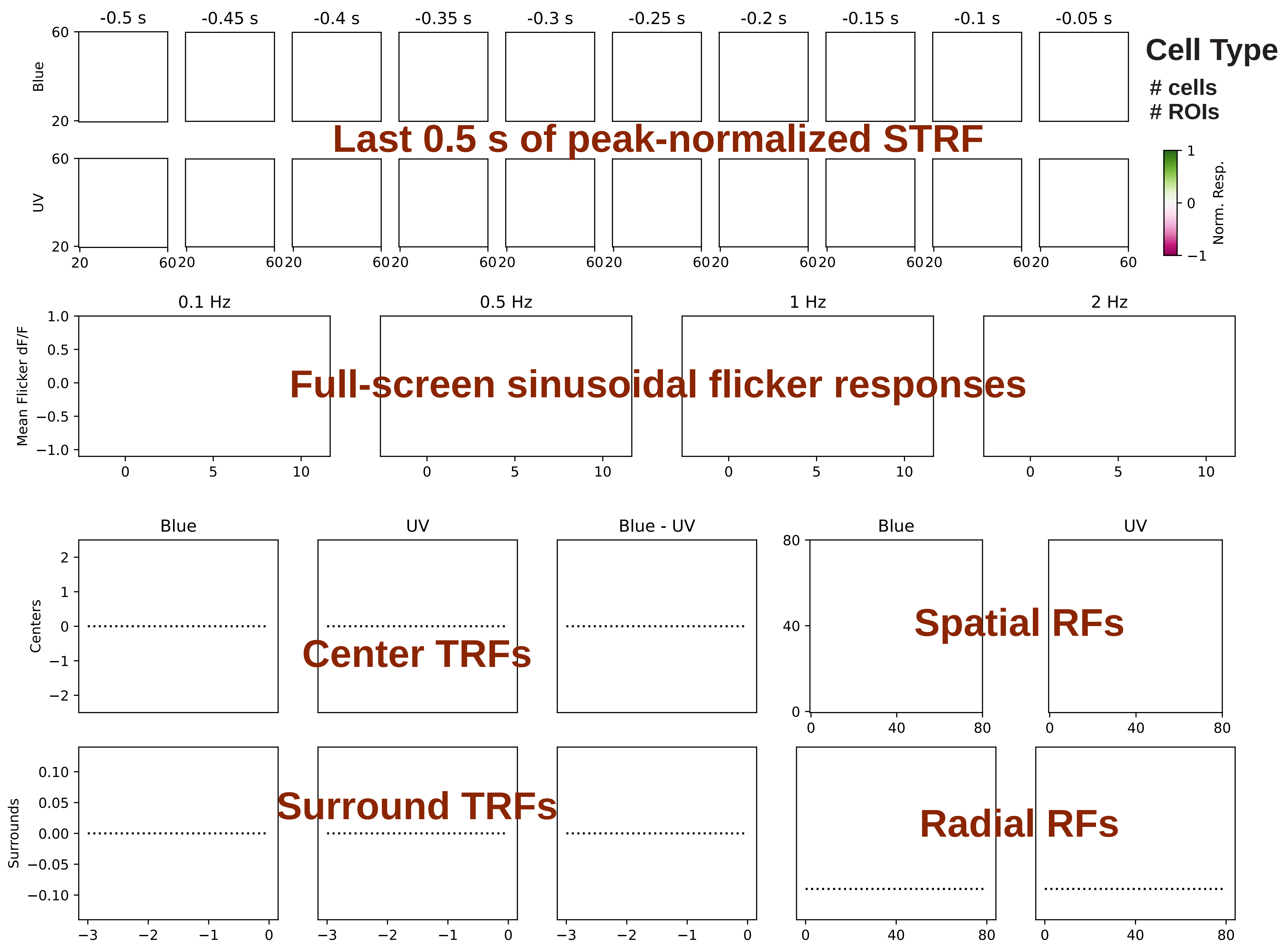

Cell Type
### cells
### ROIs
Last 0.5 s of peak-normalized STRF
-0.5 s
-0.45 s
-0.4 s
-0.35 s
-0.3 s
-0.25 s
-0.2 s
-0.15 s
-0.1 s
-0.05 s
Blue
UV
Norm. Resp.
Full-screen sinusoidal flicker responses
0.1 Hz
0.5 Hz
1 Hz
2 Hz
Mean Flicker dF/F
Center TRFs
Surround TRFs
Centers
Surrounds
Blue
UV
Blue - UV
Spatial RFs
Radial RFs

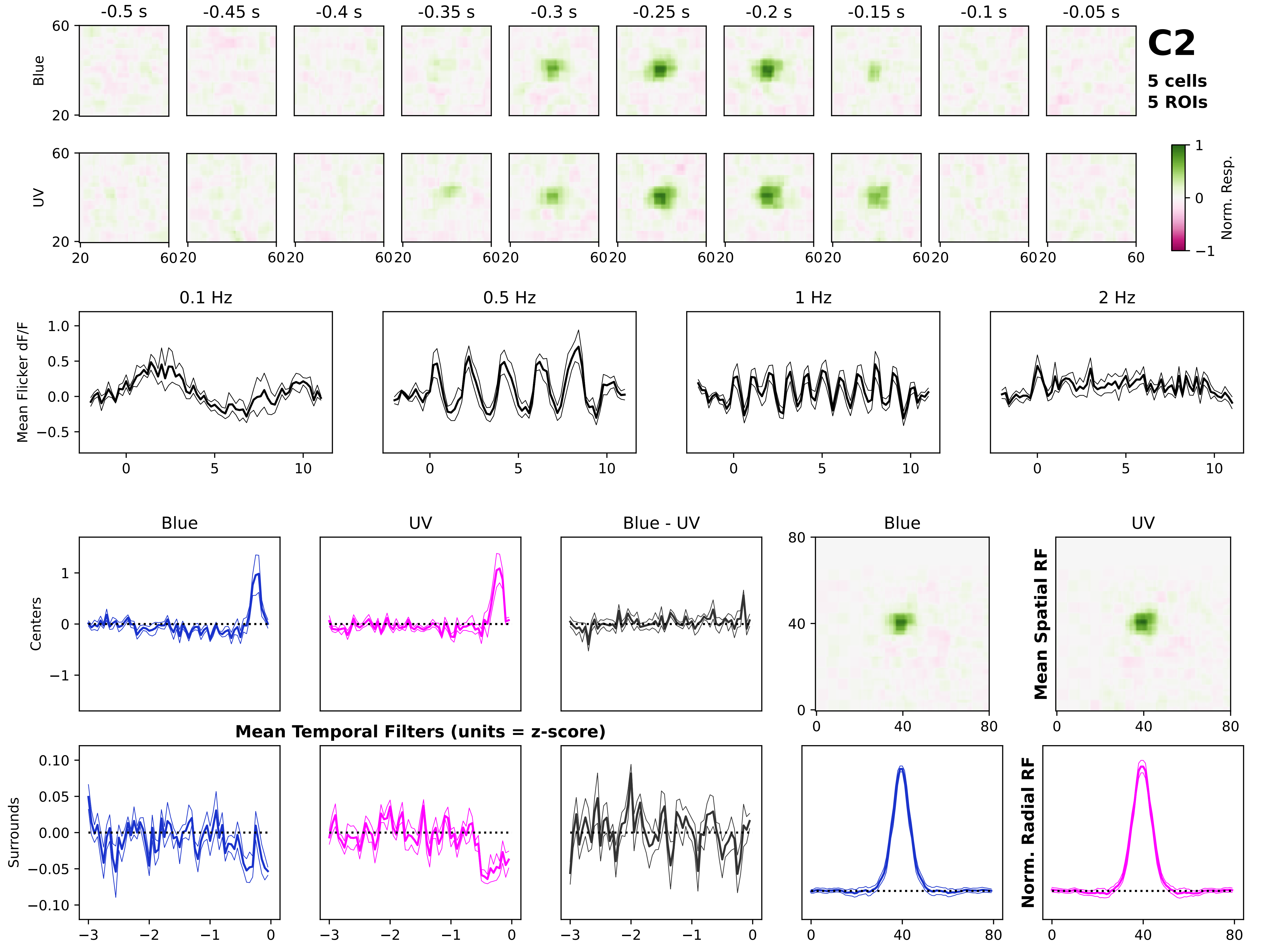

C2
5 cells
5 ROIs
-0.5 s
-0.45 s
-0.4 s
-0.35 s
-0.3 s
-0.25 s
-0.2 s
-0.15 s
-0.1 s
-0.05 s
Blue
UV
Norm. Resp.
0.1 Hz
0.5 Hz
1 Hz
2 Hz
Mean Flicker dF/F
Blue
UV
Blue - UV
Centers
Surrounds
Mean Temporal Filters (units = z-score)
Mean Spatial RF
Norm. Radial RF

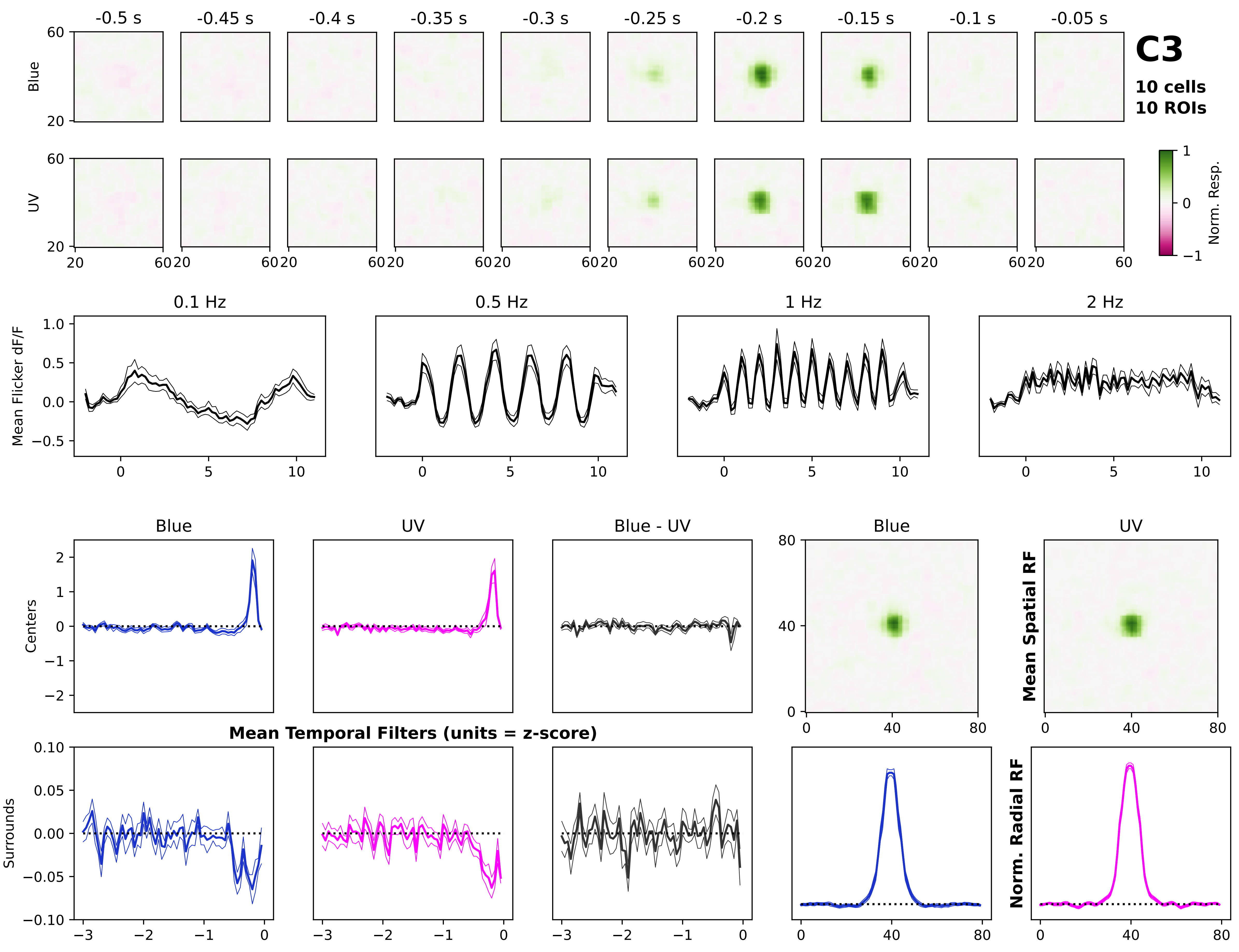
-0.5 s
-0.45 s
-0.4 s
-0.35 s
-0.3 s
-0.25 s
-0.2 s
-0.15 s
-0.1 s
-0.05 s
C3
10 cells
10 ROIs
Blue
UV
Norm. Resp.
0.1 Hz
0.5 Hz
1 Hz
2 Hz
Mean Flicker dF/F
Blue
UV
Blue - UV
Centers
Surrounds
Mean Temporal Filters (units = z-score)
Mean Spatial RF
Norm. Radial RF

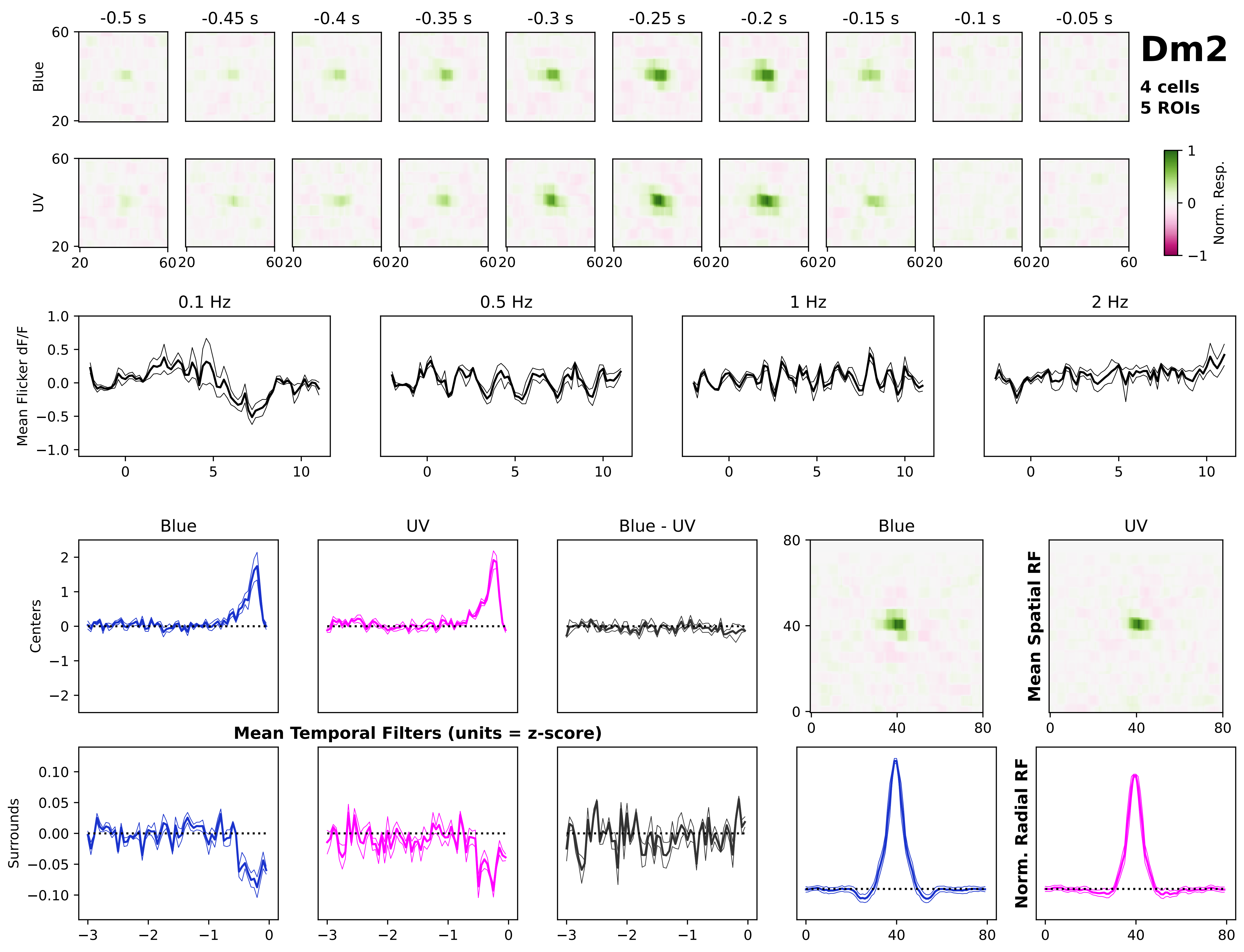
Dm2
4 cells
5 ROIs
-0.5 s
-0.45 s
-0.4 s
-0.35 s
-0.3 s
-0.25 s
-0.2 s
-0.15 s
-0.1 s
-0.05 s
Blue
UV
Norm. Resp.
0.1 Hz
0.5 Hz
1 Hz
2 Hz
Mean Flicker dF/F
Blue
UV
Blue - UV
Centers
Surrounds
Mean Temporal Filters (units = z-score)
Mean Spatial RF
Norm. Radial RF

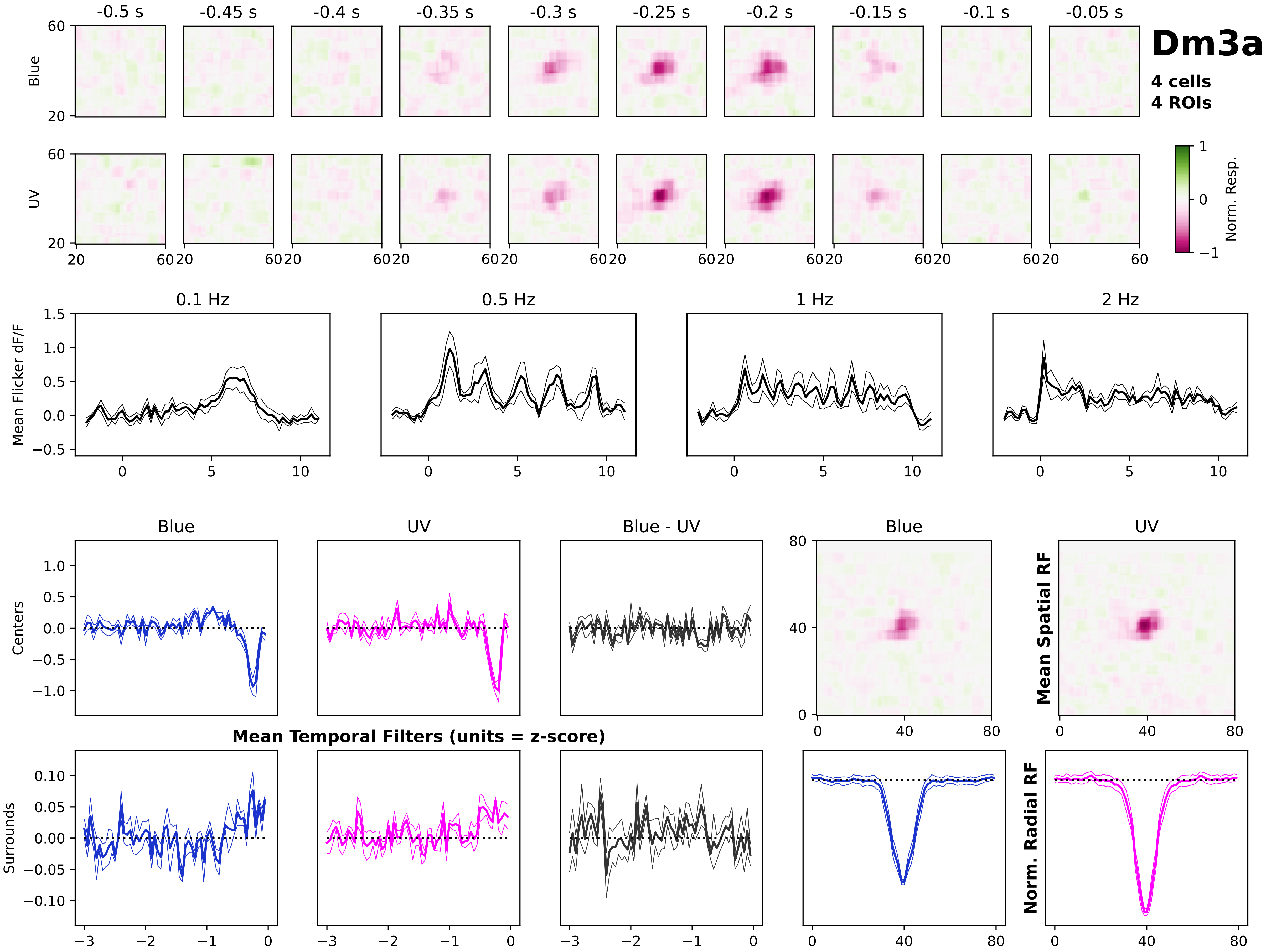
Dm3a
4 cells
4 ROIs
-0.5 s
-0.45 s
-0.4 s
-0.35 s
-0.3 s
-0.25 s
-0.2 s
-0.15 s
-0.1 s
-0.05 s
Blue
UV
Norm. Resp.
0.1 Hz
0.5 Hz
1 Hz
2 Hz
Mean Flicker dF/F
Blue
UV
Blue - UV
Centers
Surrounds
Mean Temporal Filters (units = z-score)
Mean Spatial RF
Norm. Radial RF

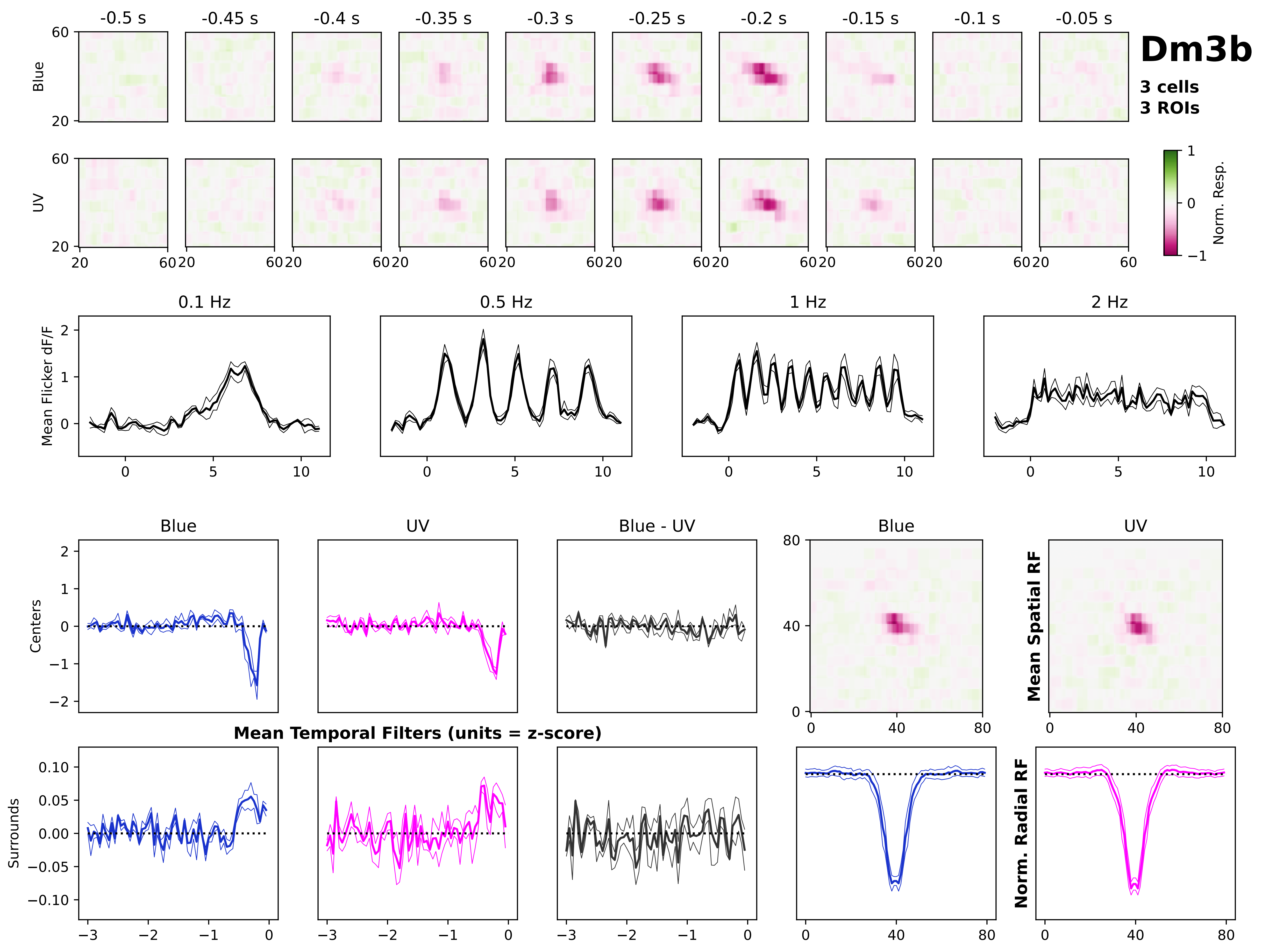

Dm3b
3 cells
3 ROIs
-0.5 s
-0.45 s
-0.4 s
-0.35 s
-0.3 s
-0.25 s
-0.2 s
-0.15 s
-0.1 s
-0.05 s
Blue
UV
Norm. Resp.
0.1 Hz
0.5 Hz
1 Hz
2 Hz
Mean Flicker dF/F
Blue
UV
Blue - UV
Centers
Surrounds
Mean Temporal Filters (units = z-score)
Mean Spatial RF
Norm. Radial RF

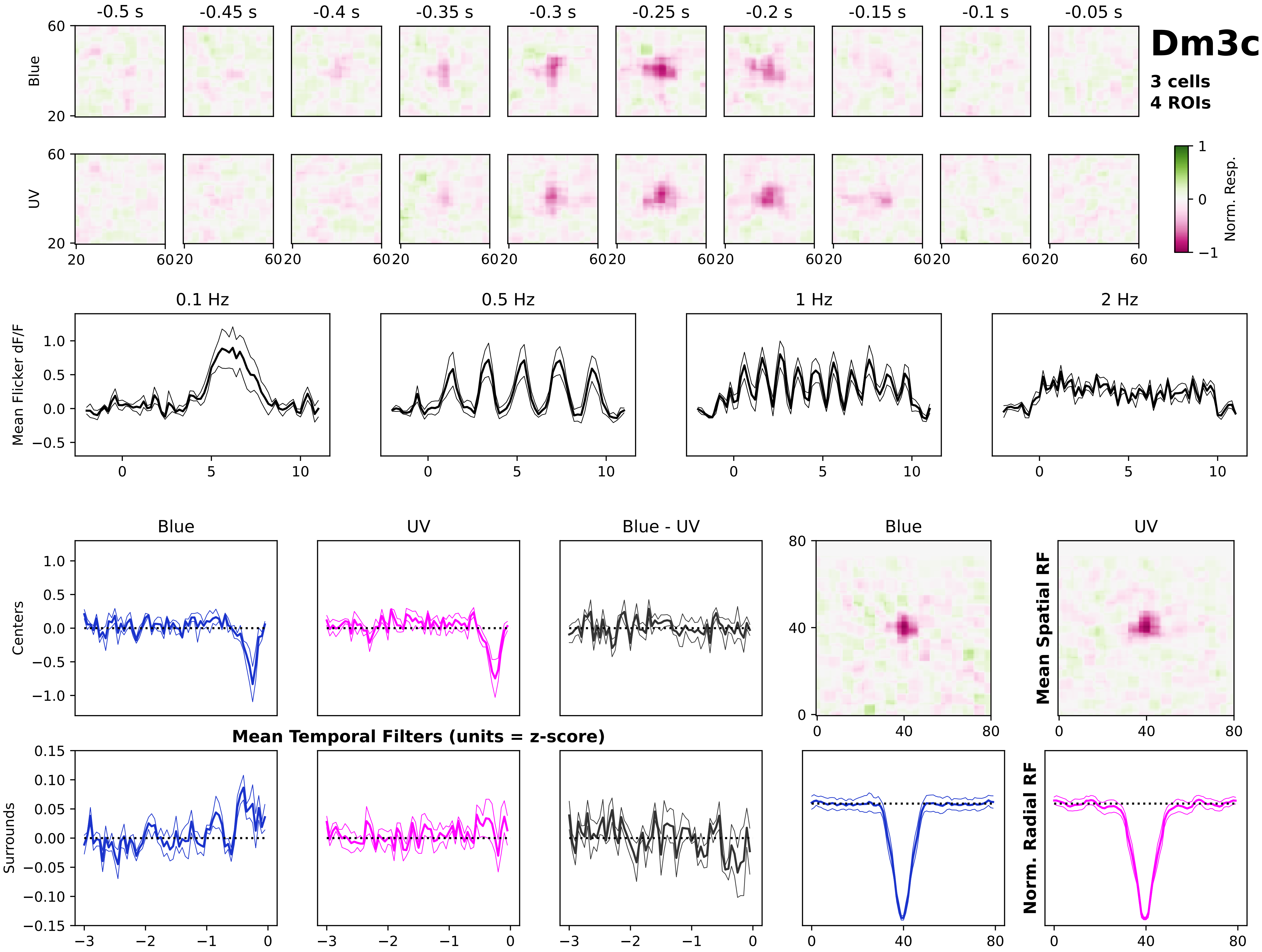
Dm3c
3 cells
4 ROIs
-0.5 s
-0.45 s
-0.4 s
-0.35 s
-0.3 s
-0.25 s
-0.2 s
-0.15 s
-0.1 s
-0.05 s
Blue
UV
Norm. Resp.
0.1 Hz
0.5 Hz
1 Hz
2 Hz
Mean Flicker dF/F
Blue
UV
Blue - UV
Centers
Surrounds
Mean Temporal Filters (units = z-score)
Mean Spatial RF
Norm. Radial RF

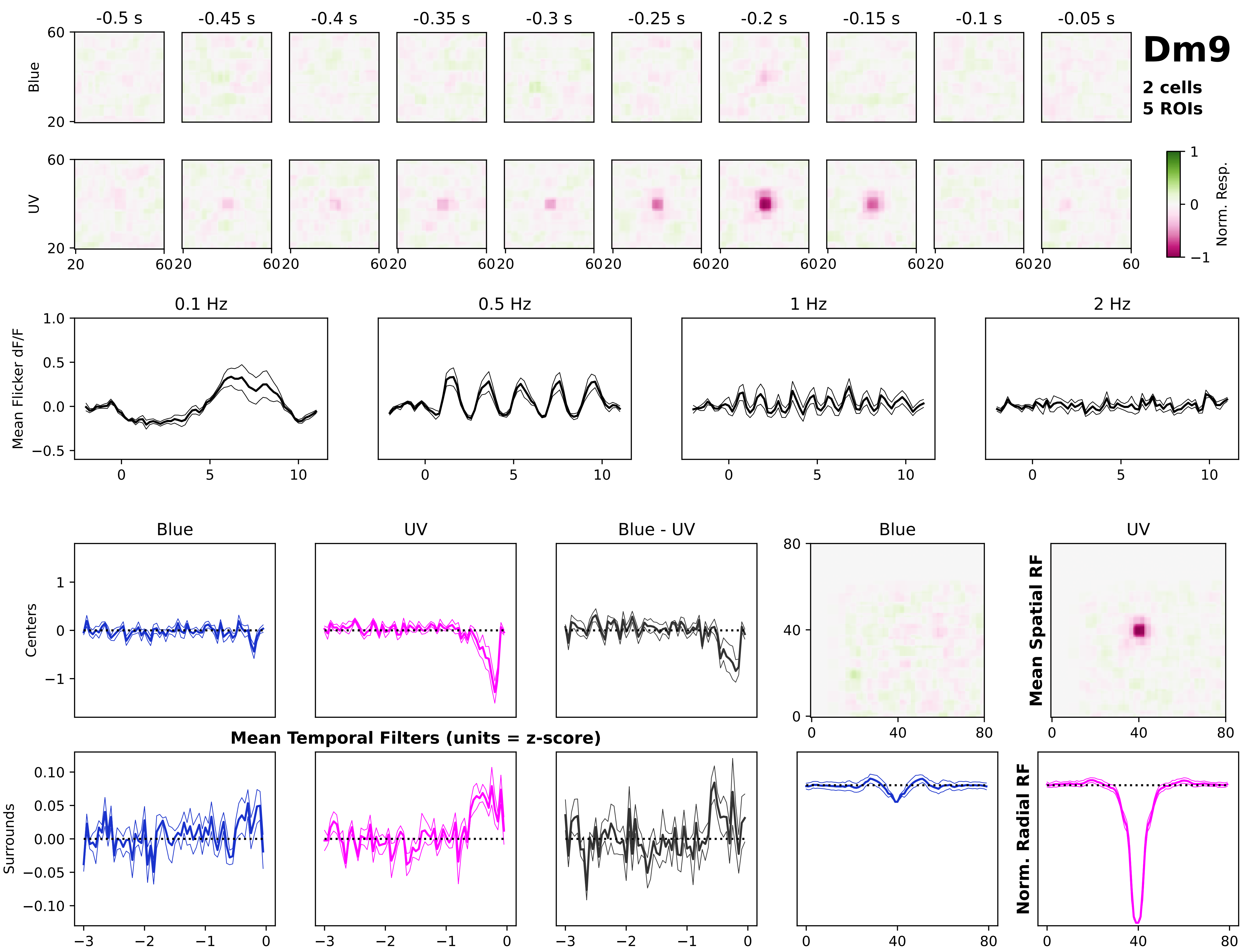

Dm9
2 cells
5 ROIs
-0.5 s
-0.45 s
-0.4 s
-0.35 s
-0.3 s
-0.25 s
-0.2 s
-0.15 s
-0.1 s
-0.05 s
Blue
UV
Norm. Resp.
0.1 Hz
0.5 Hz
1 Hz
2 Hz
Mean Flicker dF/F
Blue
UV
Blue - UV
Centers
Surrounds
Mean Temporal Filters (units = z-score)
Mean Spatial RF
Norm. Radial RF

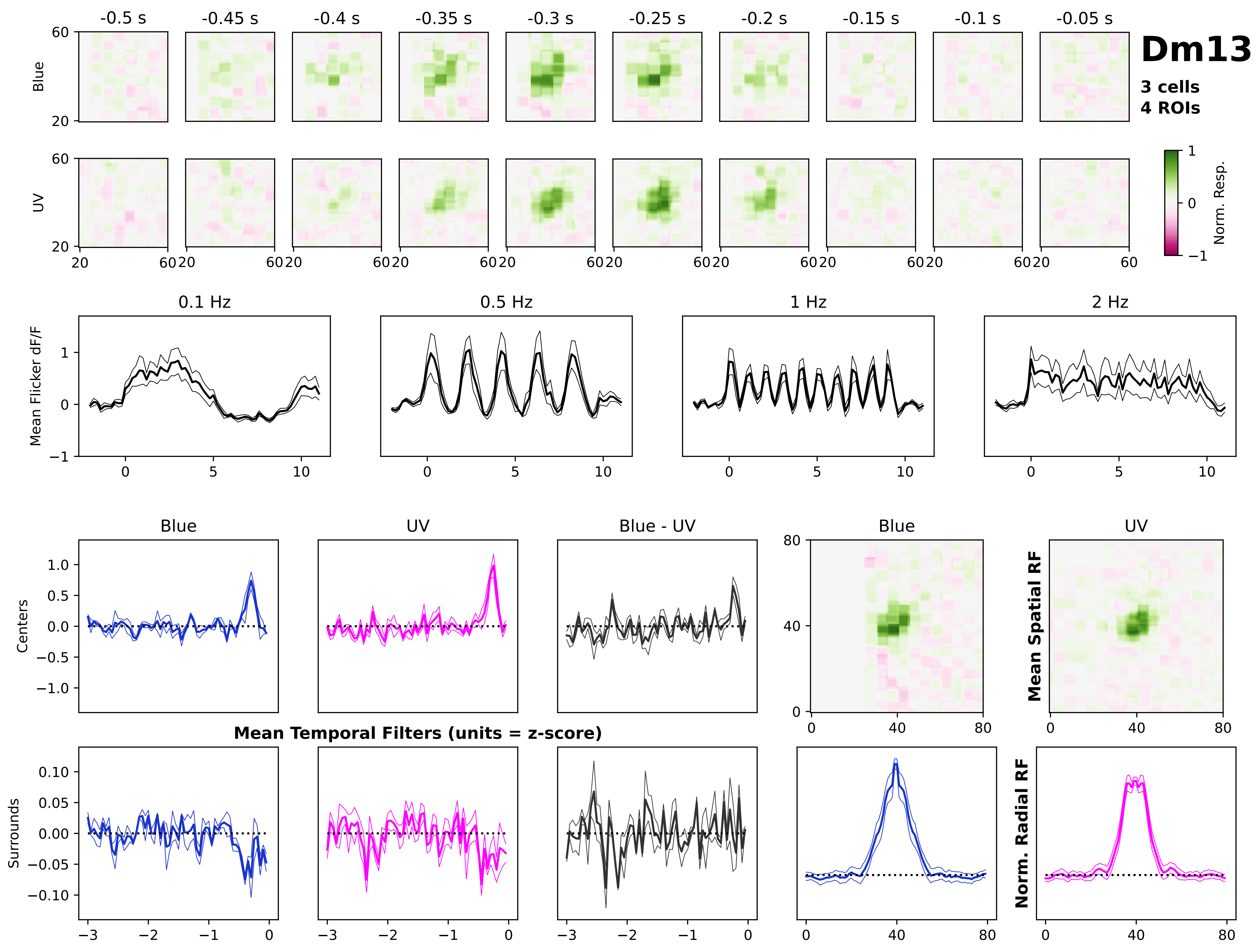
Dm13
3 cells
4 ROIs
-0.5 s
-0.45 s
-0.4 s
-0.35 s
-0.3 s
-0.25 s
-0.2 s
-0.15 s
-0.1 s
-0.05 s
Blue
UV
Norm. Resp.
0.1 Hz
0.5 Hz
1 Hz
2 Hz
Mean Flicker dF/F
Blue
UV
Blue - UV
Centers
Surrounds
Mean Temporal Filters (units = z-score)
Mean Spatial RF
Norm. Radial RF

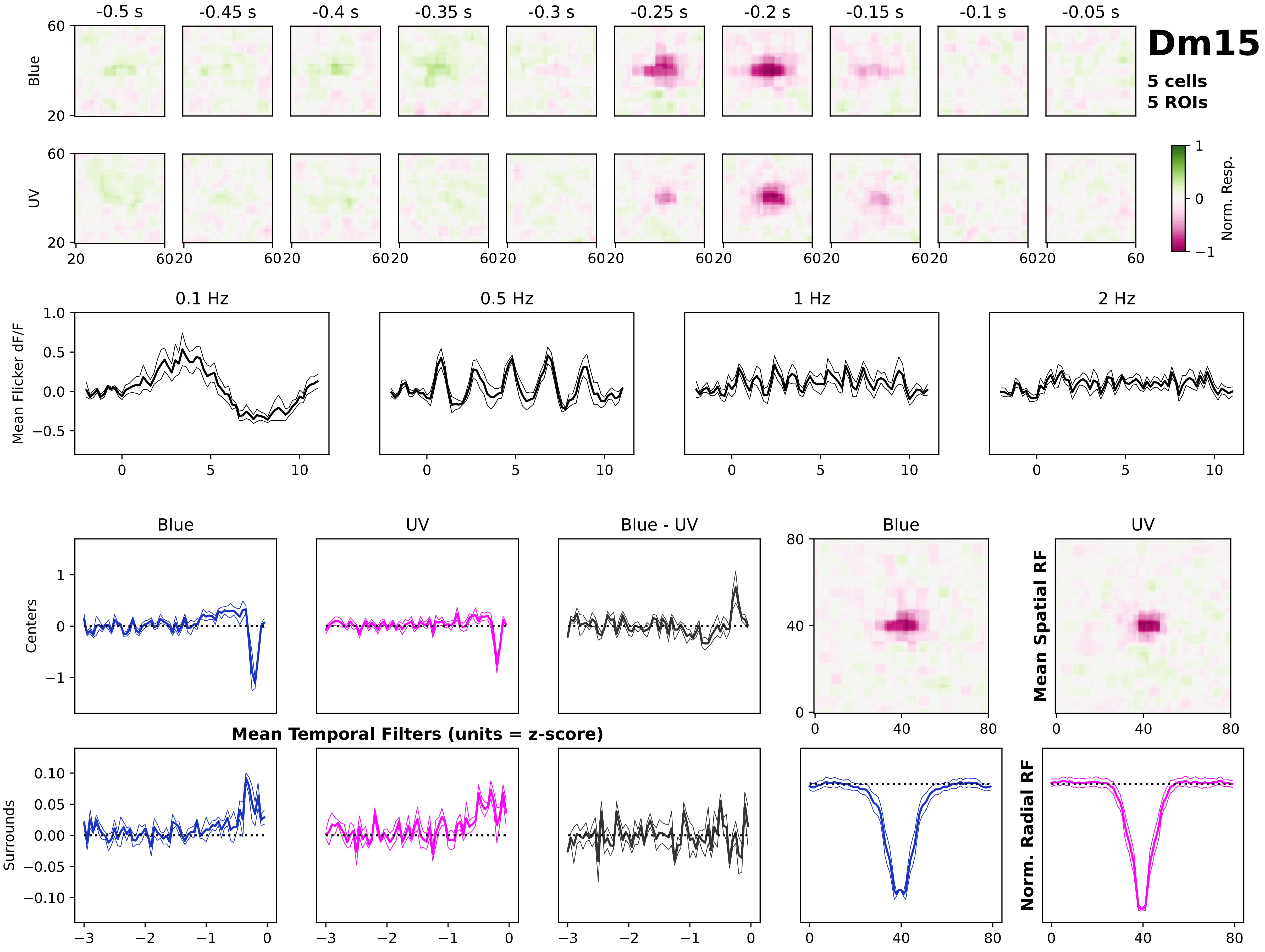

Dm15
5 cells
5 ROIs
-0.5 s
-0.45 s
-0.4 s
-0.35 s
-0.3 s
-0.25 s
-0.2 s
-0.15 s
-0.1 s
-0.05 s
Blue
UV
Norm. Resp.
0.1 Hz
0.5 Hz
1 Hz
2 Hz
Mean Flicker dF/F
Blue
UV
Blue - UV
Centers
Surrounds
Mean Temporal Filters (units = z-score)
Mean Spatial RF
Norm. Radial RF

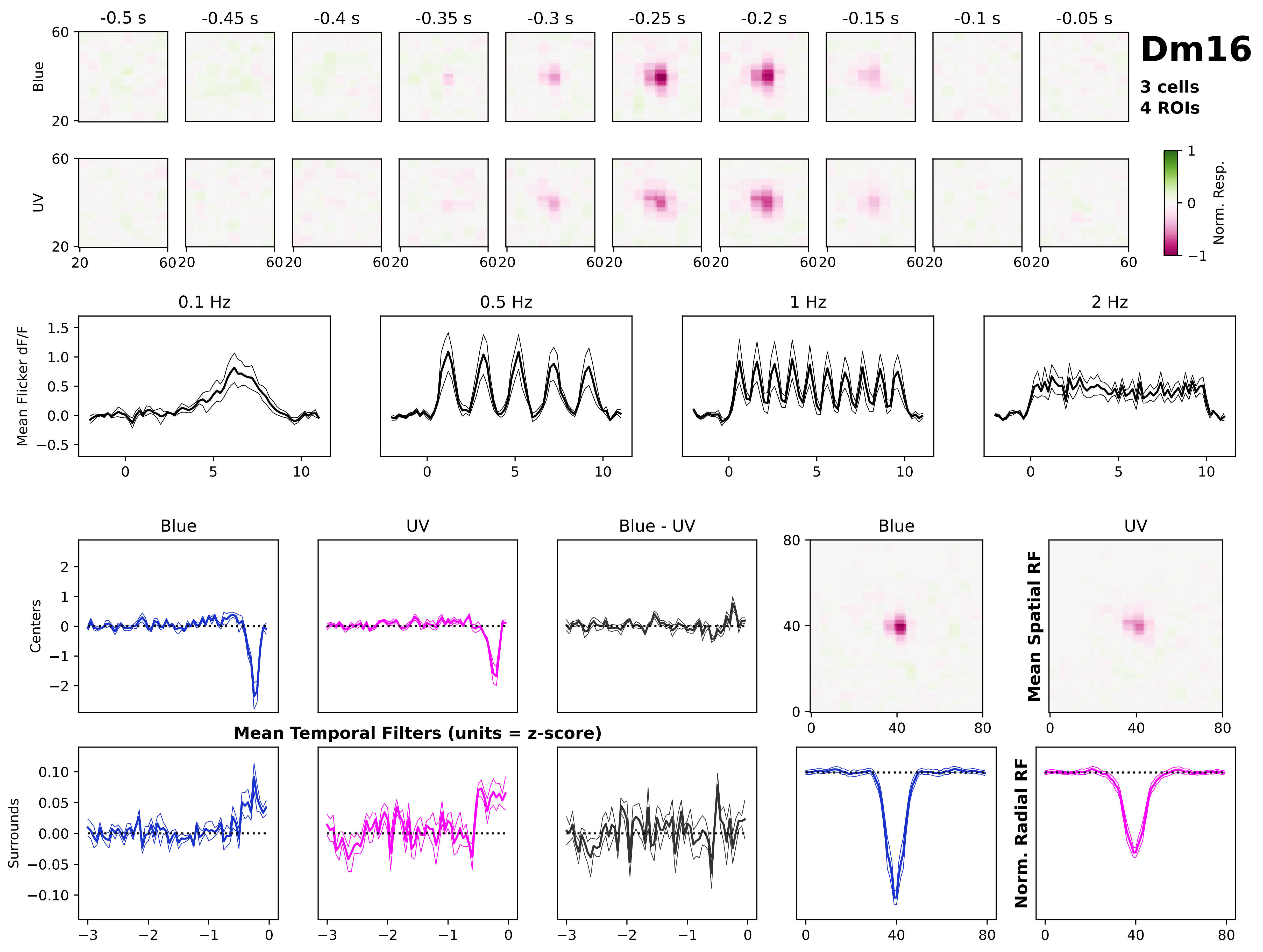
Dm16
3 cells
4 ROIs
-0.5 s
-0.45 s
-0.4 s
-0.35 s
-0.3 s
-0.25 s
-0.2 s
-0.15 s
-0.1 s
-0.05 s
Blue
UV
Norm. Resp.
0.1 Hz
0.5 Hz
1 Hz
2 Hz
Mean Flicker dF/F
Blue
UV
Blue - UV
Centers
Surrounds
Mean Temporal Filters (units = z-score)
Mean Spatial RF
Norm. Radial RF

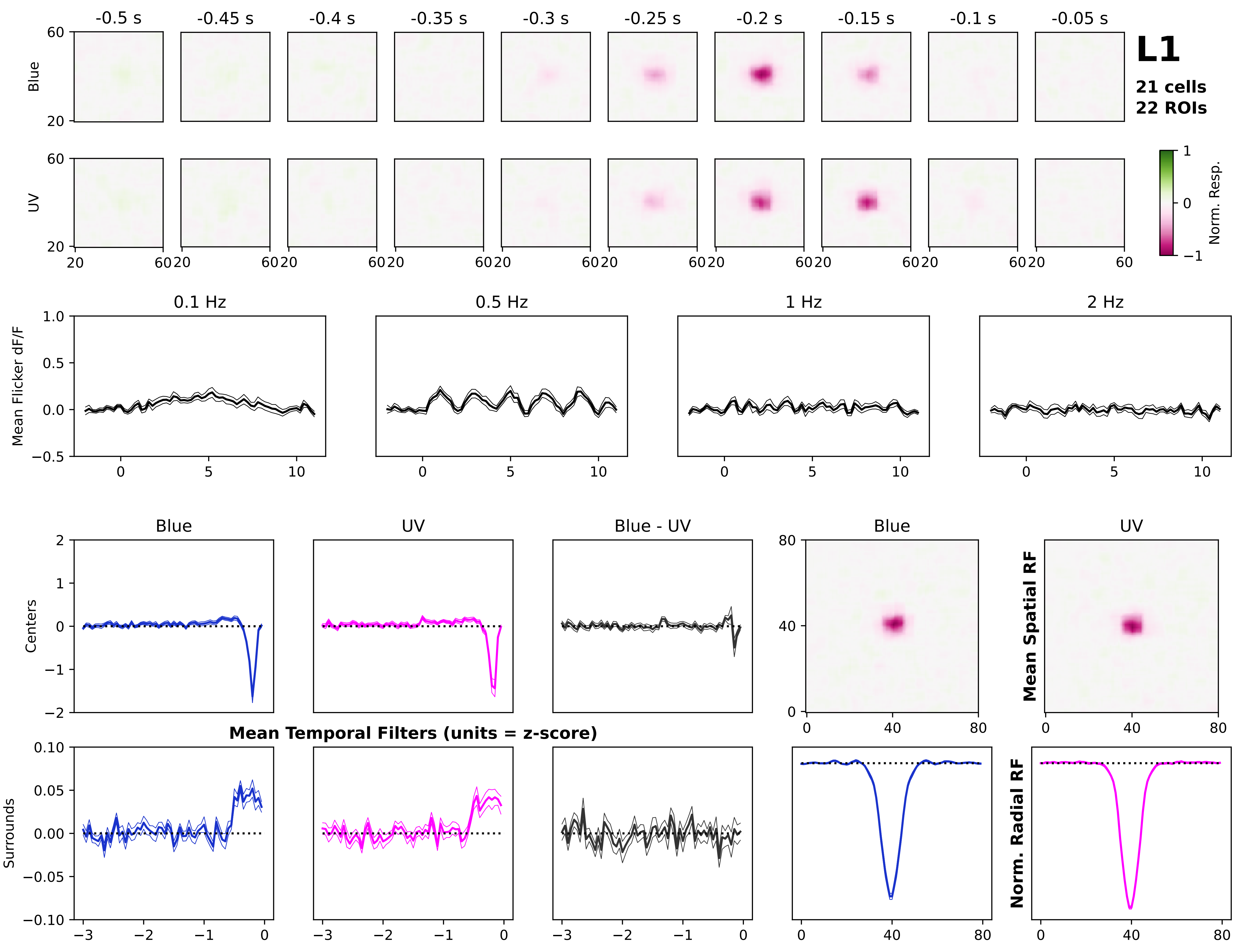

L1
21 cells
22 ROIs
-0.5 s
-0.45 s
-0.4 s
-0.35 s
-0.3 s
-0.25 s
-0.2 s
-0.15 s
-0.1 s
-0.05 s
Blue
UV
Norm. Resp.
0.1 Hz
0.5 Hz
1 Hz
2 Hz
Mean Flicker dF/F
Blue
UV
Blue - UV
Centers
Surrounds
Mean Temporal Filters (units = z-score)
Mean Spatial RF
Norm. Radial RF

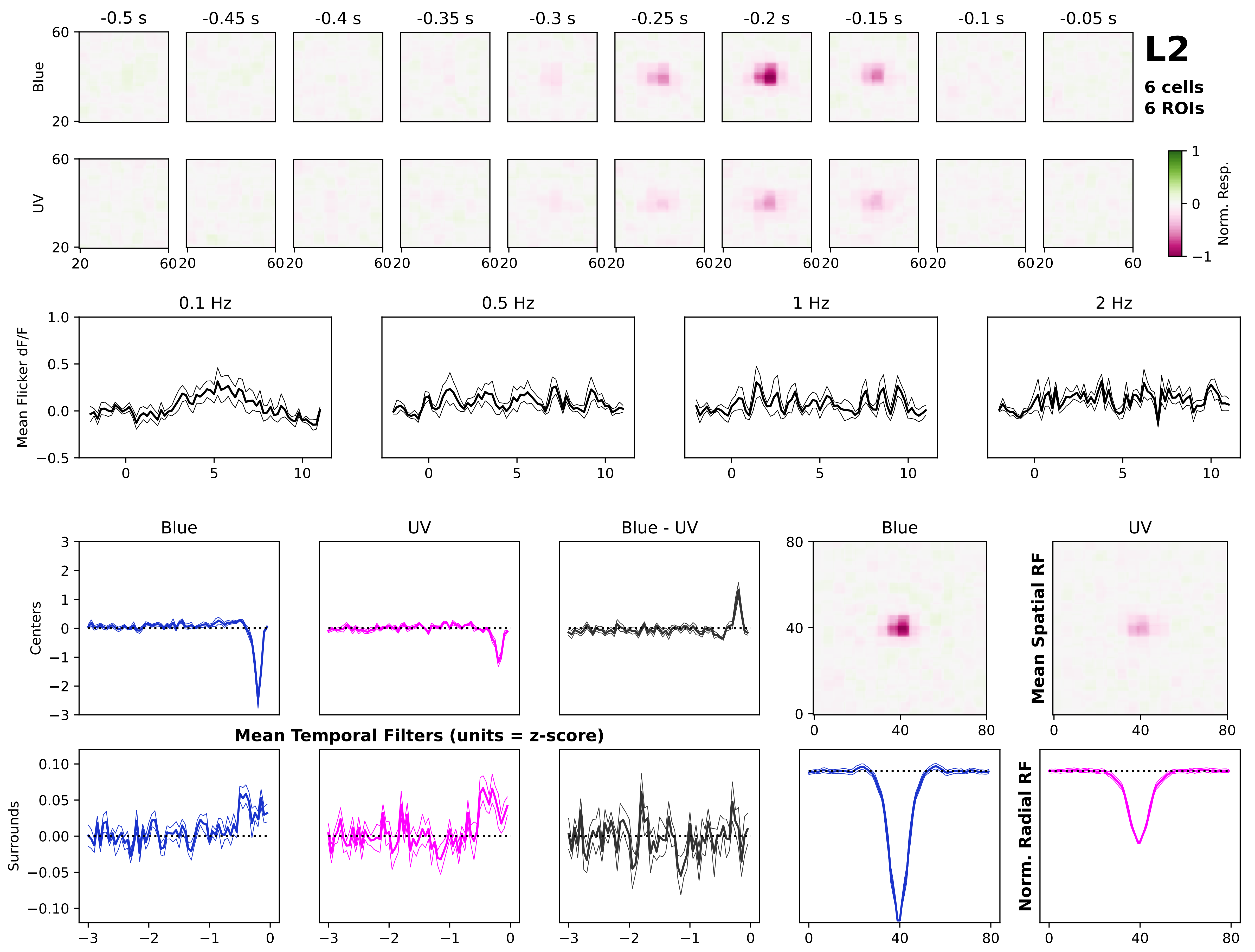

L2
6 cells
6 ROIs
-0.5 s
-0.45 s
-0.4 s
-0.35 s
-0.3 s
-0.25 s
-0.2 s
-0.15 s
-0.1 s
-0.05 s
Blue
UV
Norm. Resp.
0.1 Hz
0.5 Hz
1 Hz
2 Hz
Mean Flicker dF/F
Blue
UV
Blue - UV
Centers
Surrounds
Mean Temporal Filters (units = z-score)
Mean Spatial RF
Norm. Radial RF

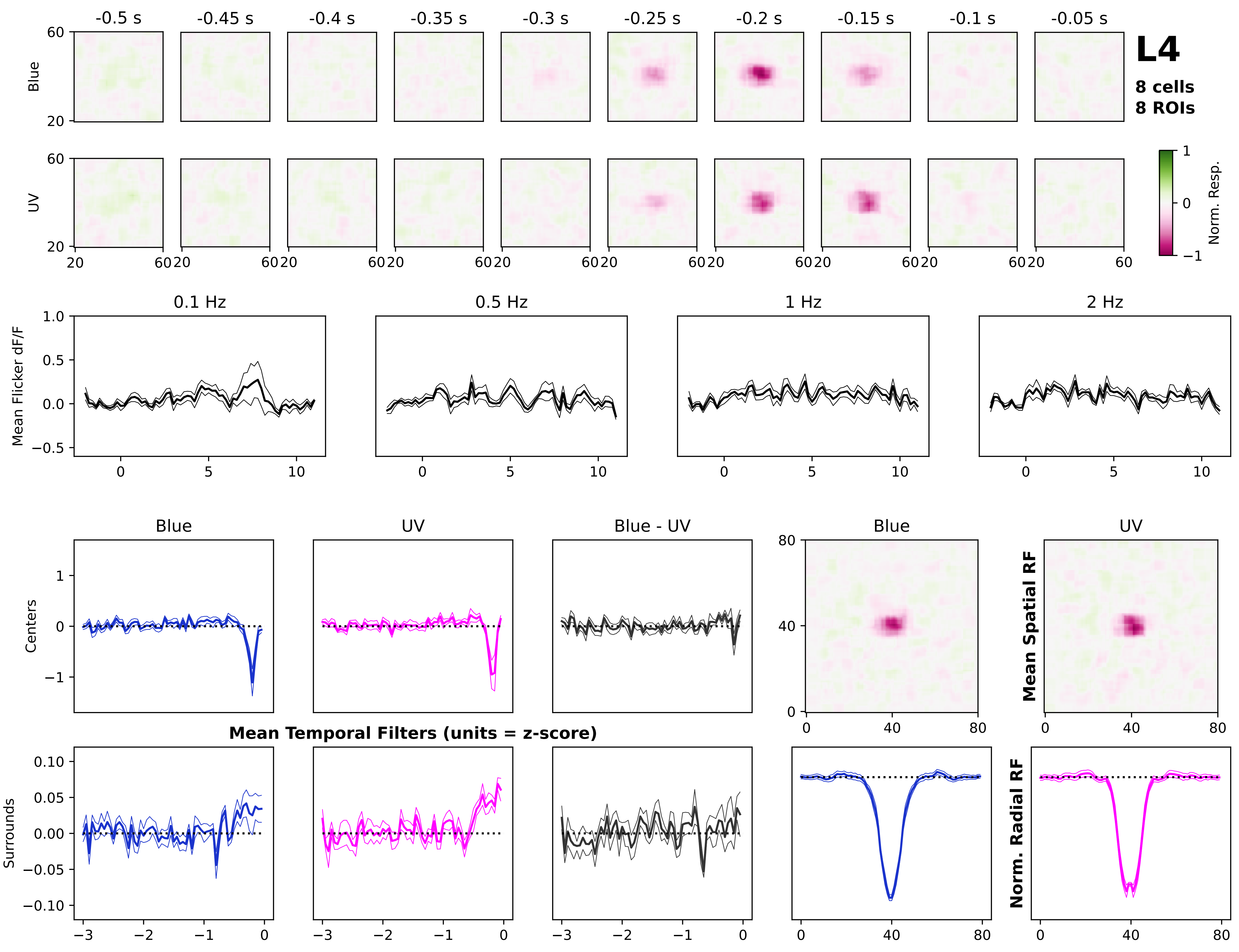

L4
8 cells
8 ROIs
-0.5 s
-0.45 s
-0.4 s
-0.35 s
-0.3 s
-0.25 s
-0.2 s
-0.15 s
-0.1 s
-0.05 s
Blue
UV
Norm. Resp.
0.1 Hz
0.5 Hz
1 Hz
2 Hz
Mean Flicker dF/F
Blue
UV
Blue - UV
Centers
Surrounds
Mean Temporal Filters (units = z-score)
Mean Spatial RF
Norm. Radial RF

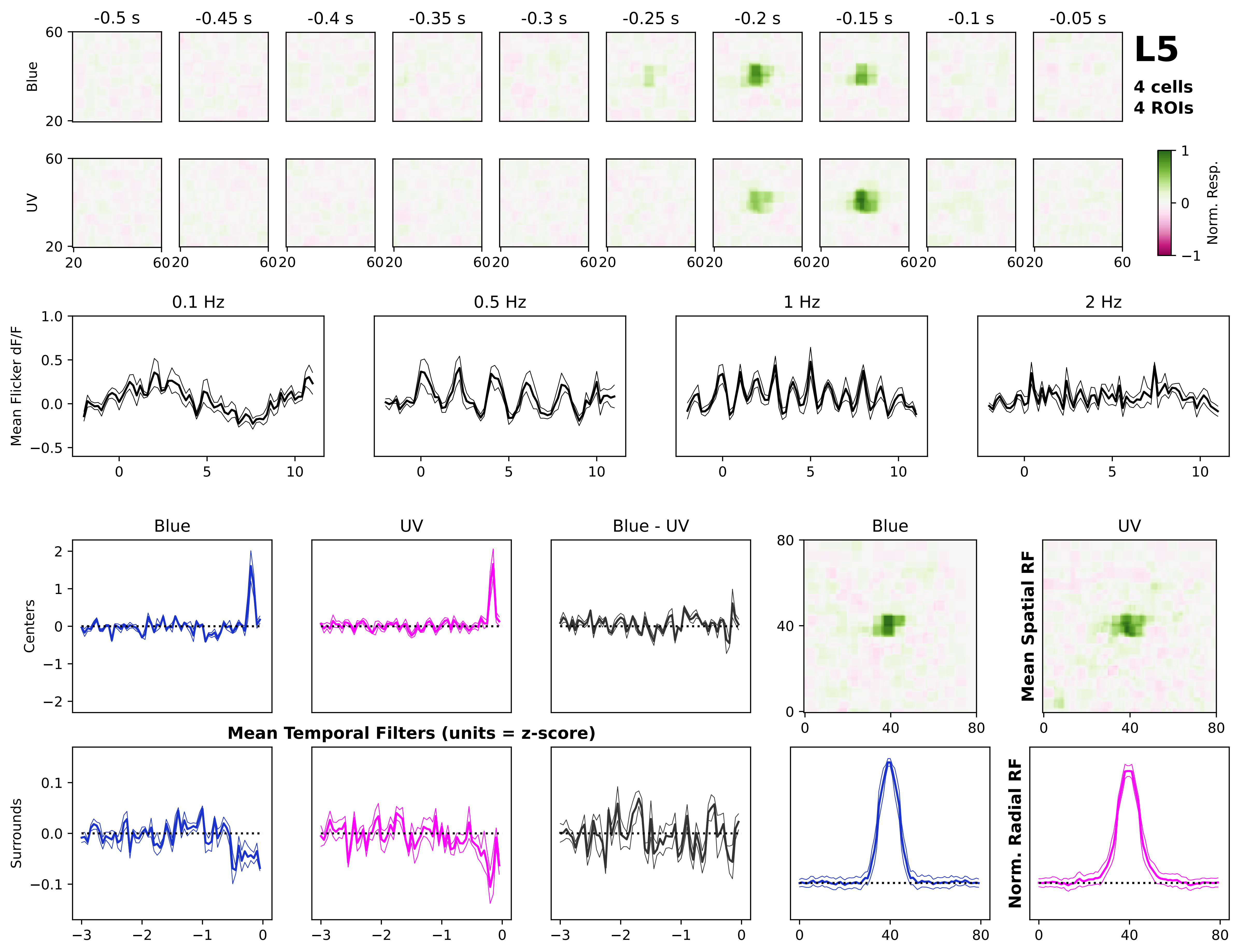
L5
4 cells
4 ROIs
-0.5 s
-0.45 s
-0.4 s
-0.35 s
-0.3 s
-0.25 s
-0.2 s
-0.15 s
-0.1 s
-0.05 s
Blue
UV
Norm. Resp.
0.1 Hz
0.5 Hz
1 Hz
2 Hz
Mean Flicker dF/F
Centers
Surrounds
Blue
UV
Blue - UV
Mean Temporal Filters (units = z-score)
Mean Spatial RF
Norm. Radial RF

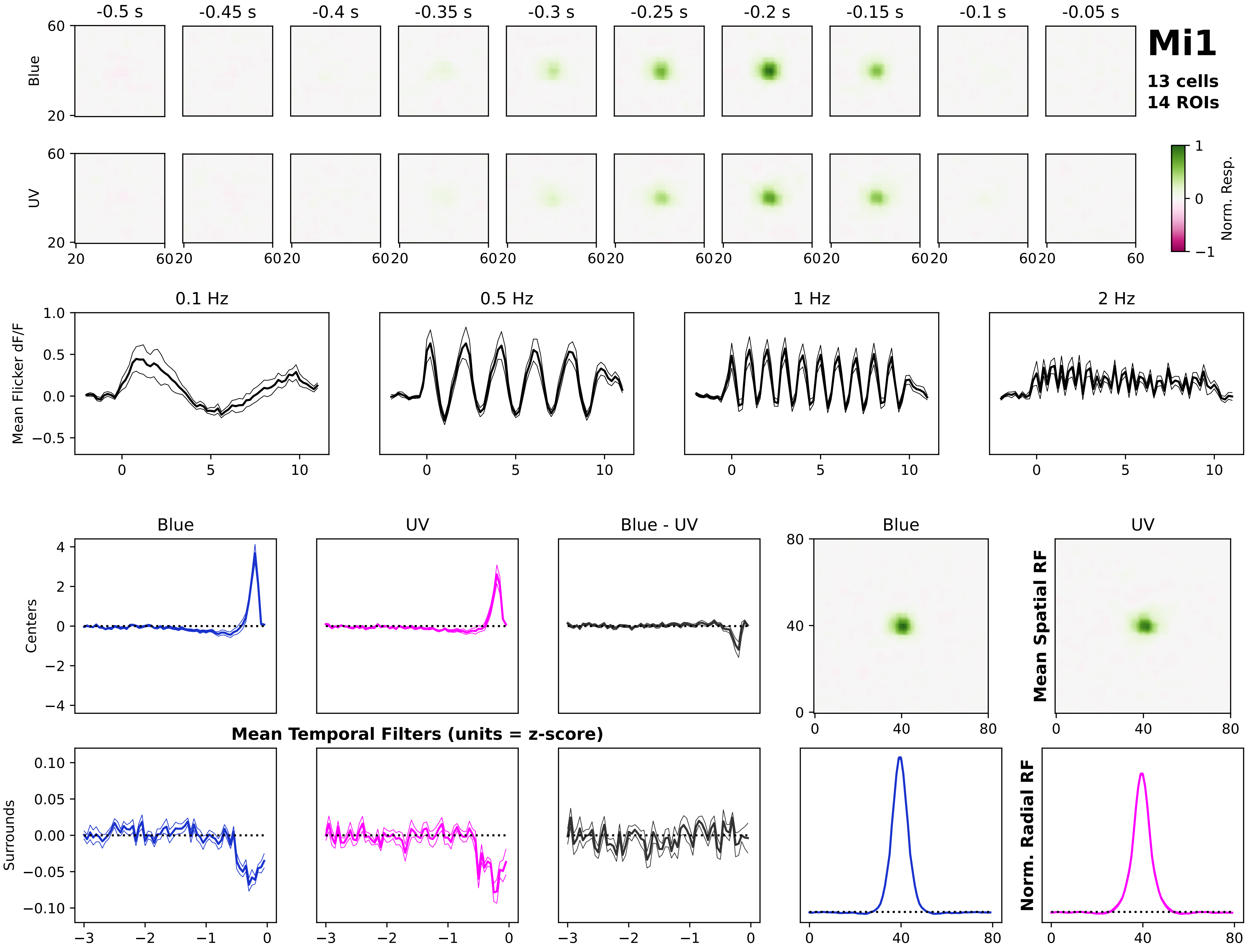

Mi1
13 cells
14 ROIs
-0.5 s
-0.45 s
-0.4 s
-0.35 s
-0.3 s
-0.25 s
-0.2 s
-0.15 s
-0.1 s
-0.05 s
Blue
UV
Norm. Resp.
0.1 Hz
0.5 Hz
1 Hz
2 Hz
Mean Flicker dF/F
Blue
UV
Blue - UV
Centers
Surrounds
Mean Temporal Filters (units = z-score)
Mean Spatial RF
Norm. Radial RF

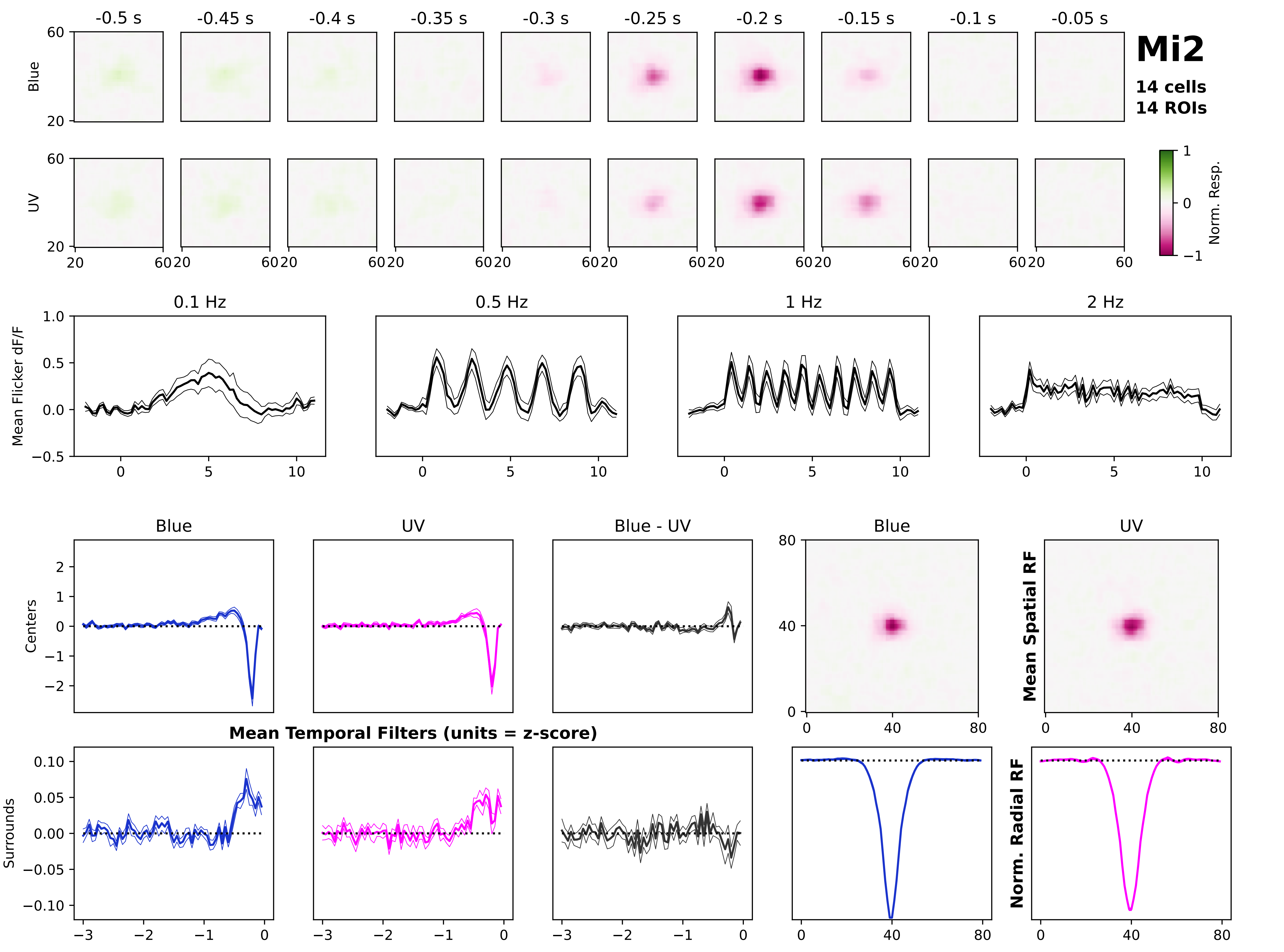

Mi2
14 cells
14 ROIs
-0.5 s
-0.45 s
-0.4 s
-0.35 s
-0.3 s
-0.25 s
-0.2 s
-0.15 s
-0.1 s
-0.05 s
Blue
UV
Norm. Resp.
0.1 Hz
0.5 Hz
1 Hz
2 Hz
Mean Flicker dF/F
Blue
UV
Blue - UV
Centers
Surrounds
Mean Temporal Filters (units = z-score)
Mean Spatial RF
Norm. Radial RF

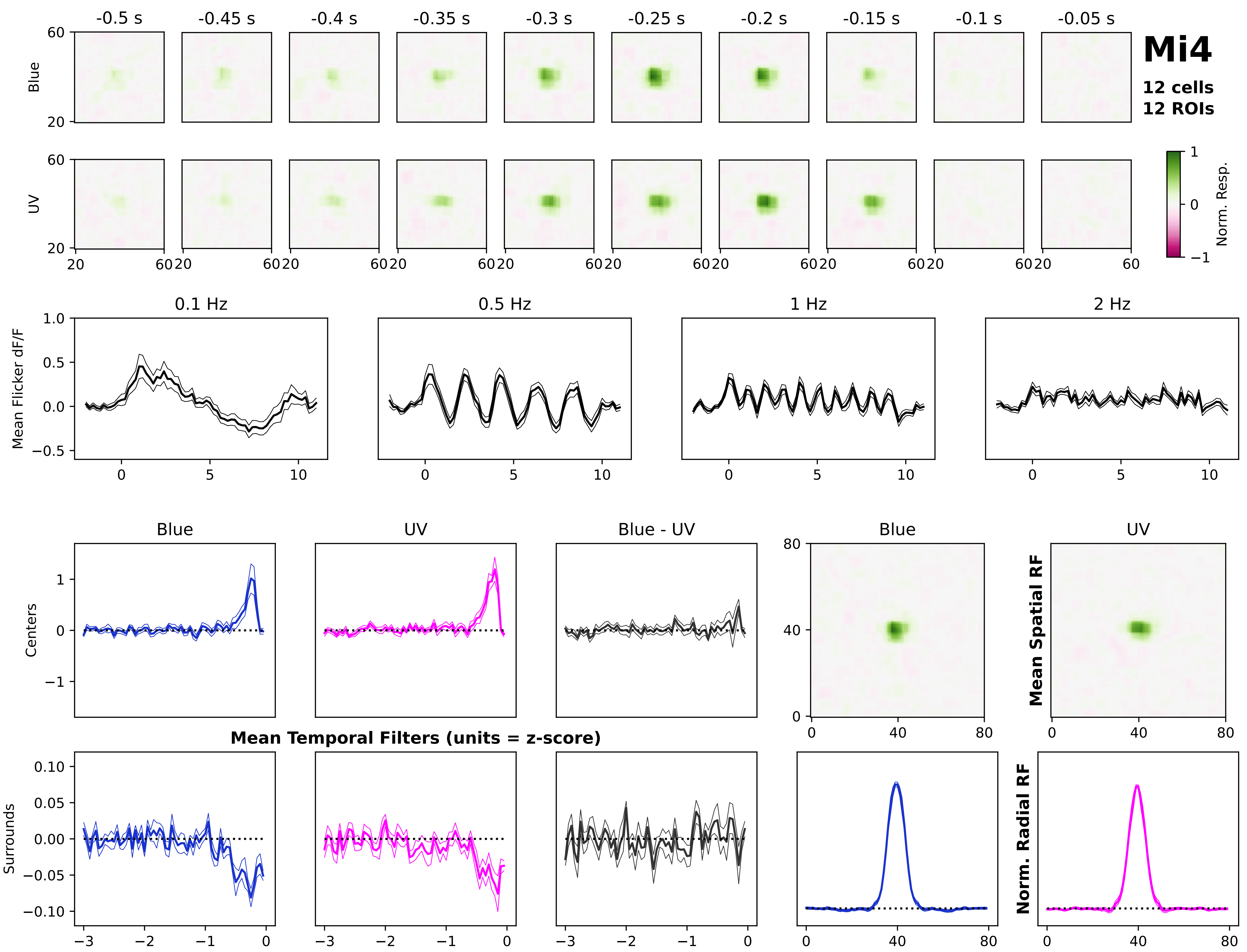

Mi4
12 cells
12 ROIs
-0.5 s
-0.45 s
-0.4 s
-0.35 s
-0.3 s
-0.25 s
-0.2 s
-0.15 s
-0.1 s
-0.05 s
Blue
UV
Norm. Resp.
0.1 Hz
0.5 Hz
1 Hz
2 Hz
Mean Flicker dF/F
Centers
Surrounds
Blue
UV
Blue - UV
Mean Temporal Filters (units = z-score)
Mean Spatial RF
Norm. Radial RF

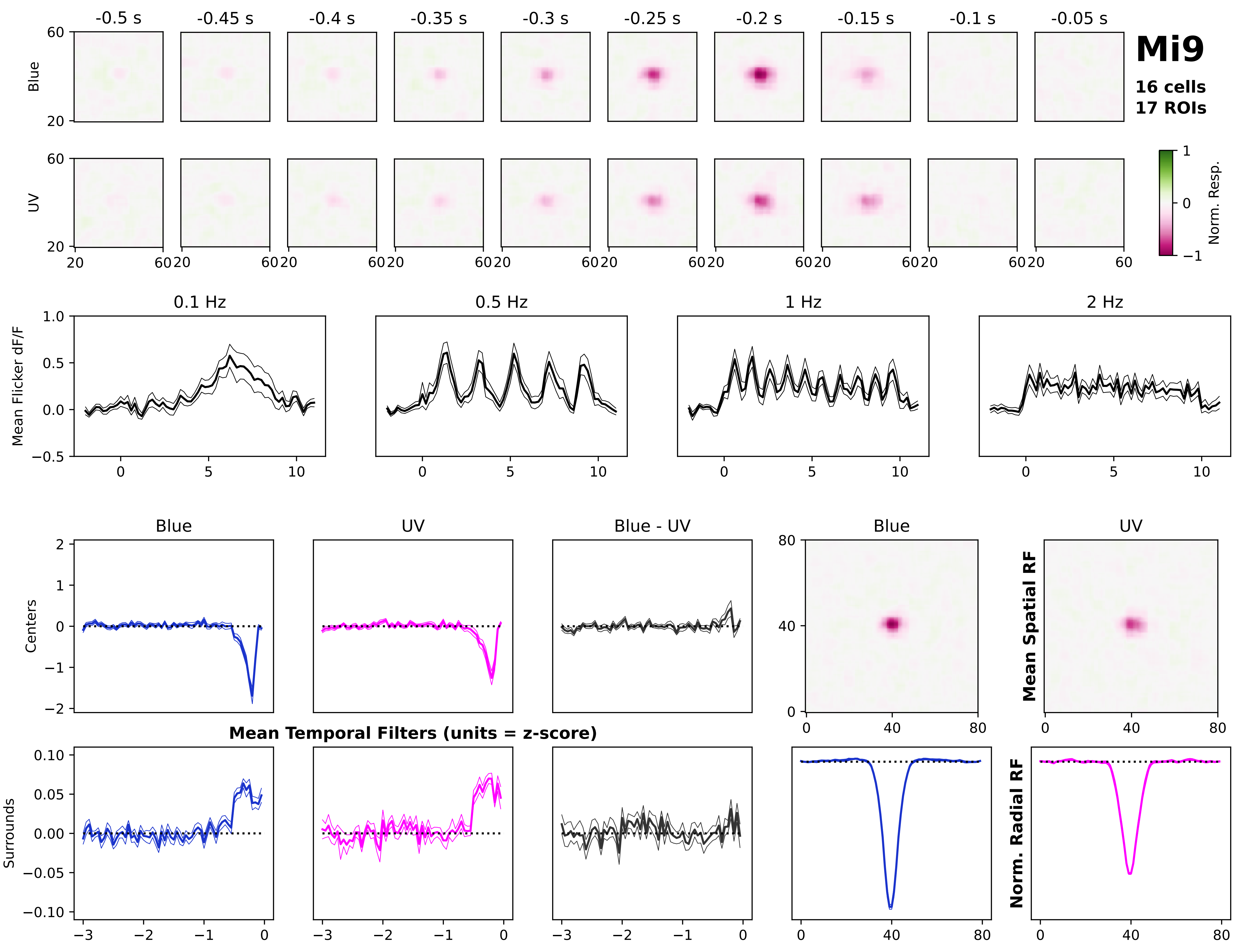

Mi9
16 cells
17 ROIs
-0.5 s
-0.45 s
-0.4 s
-0.35 s
-0.3 s
-0.25 s
-0.2 s
-0.15 s
-0.1 s
-0.05 s
Blue
UV
Norm. Resp.
0.1 Hz
0.5 Hz
1 Hz
2 Hz
Mean Flicker dF/F
Blue
UV
Blue - UV
Centers
Surrounds
Mean Temporal Filters (units = z-score)
Mean Spatial RF
Norm. Radial RF

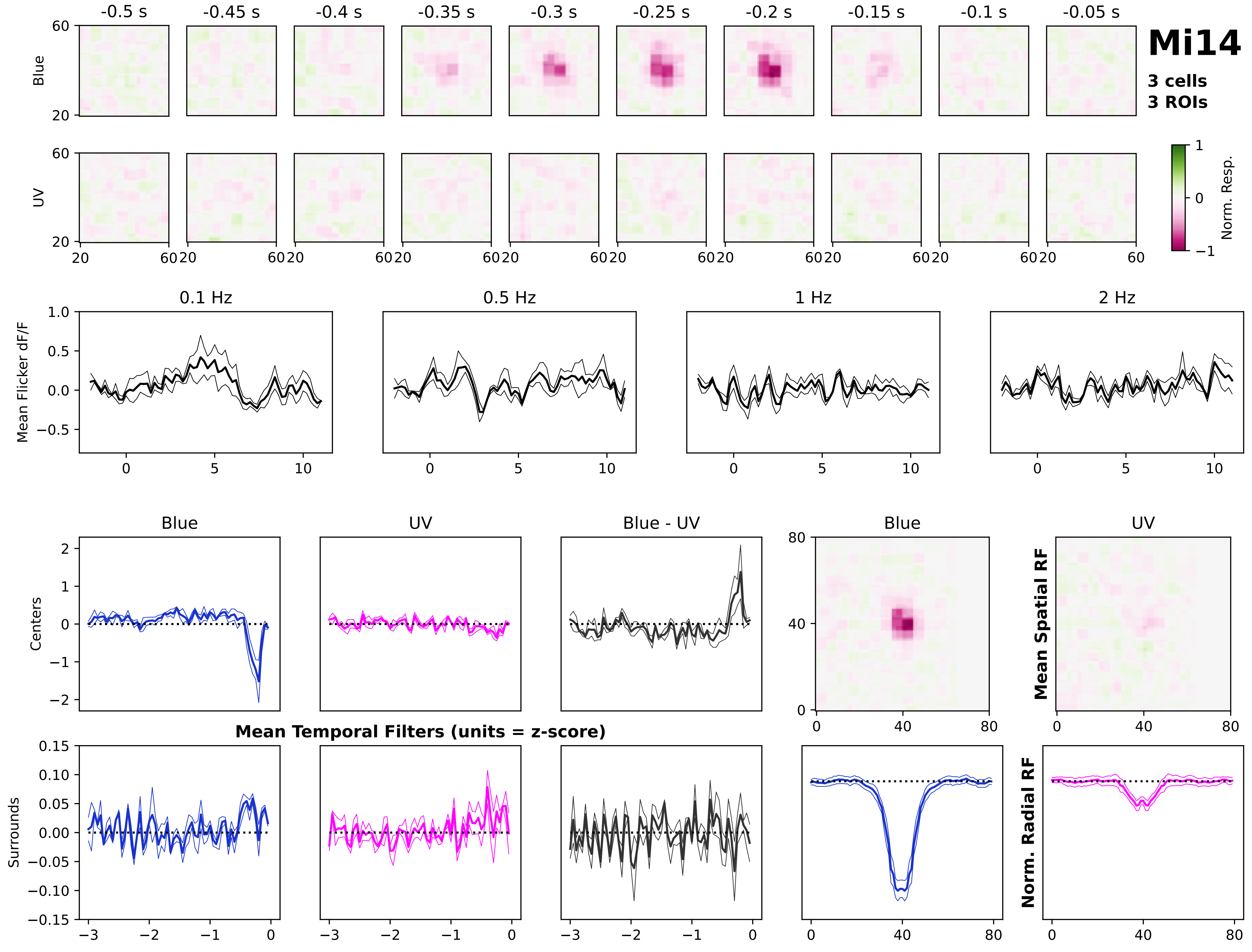

Mi14
3 cells
3 ROIs
-0.5 s
-0.45 s
-0.4 s
-0.35 s
-0.3 s
-0.25 s
-0.2 s
-0.15 s
-0.1 s
-0.05 s
Blue
UV
Norm. Resp.
0.1 Hz
0.5 Hz
1 Hz
2 Hz
Mean Flicker dF/F
Blue
UV
Blue - UV
Centers
Surrounds
Mean Temporal Filters (units = z-score)
Mean Spatial RF
Norm. Radial RF

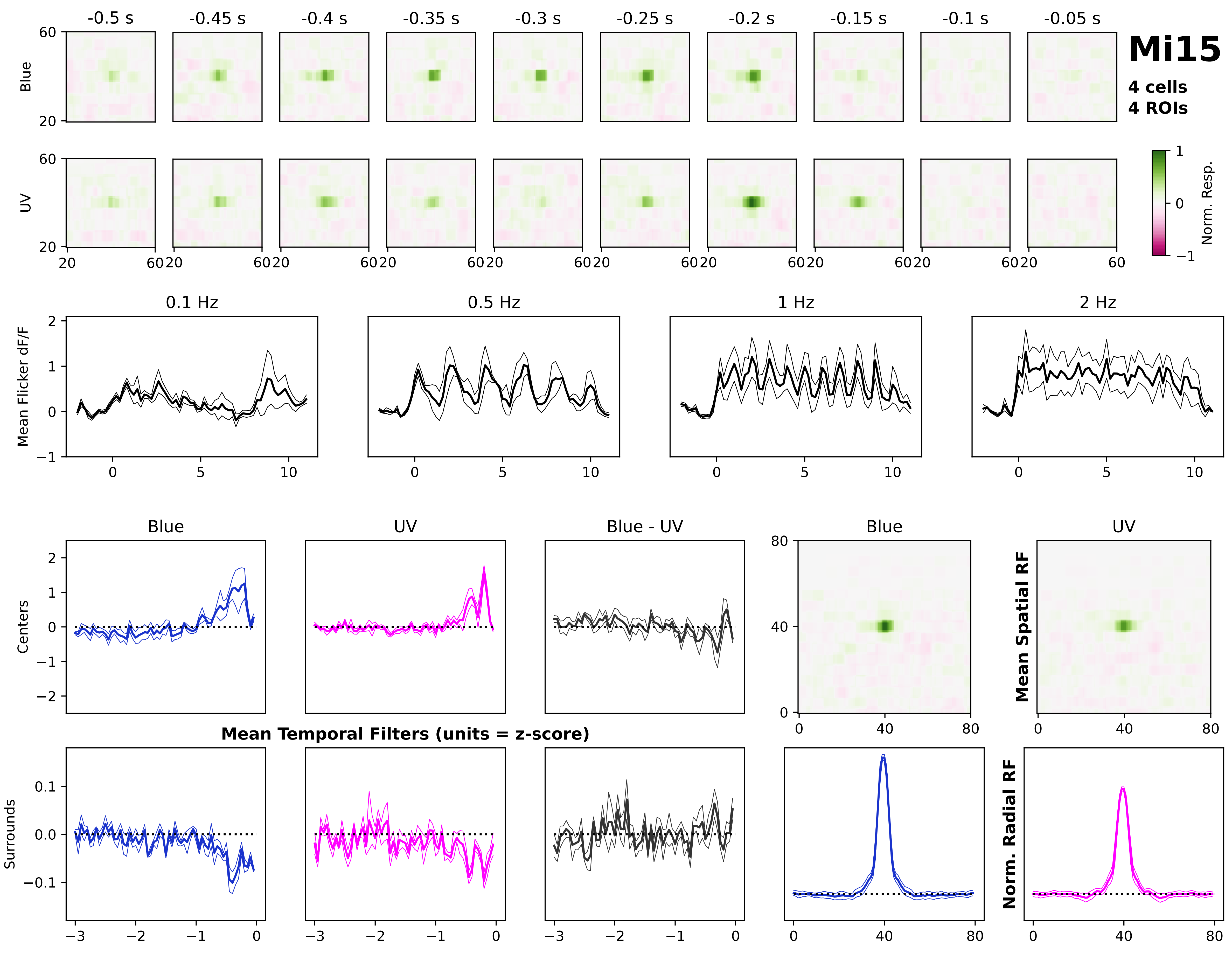

Mi15
4 cells
4 ROIs
-0.5 s
-0.45 s
-0.4 s
-0.35 s
-0.3 s
-0.25 s
-0.2 s
-0.15 s
-0.1 s
-0.05 s
Blue
UV
Norm. Resp.
0.1 Hz
0.5 Hz
1 Hz
2 Hz
Mean Flicker dF/F
Blue
UV
Blue - UV
Centers
Surrounds
Mean Temporal Filters (units = z-score)
Mean Spatial RF
Norm. Radial RF

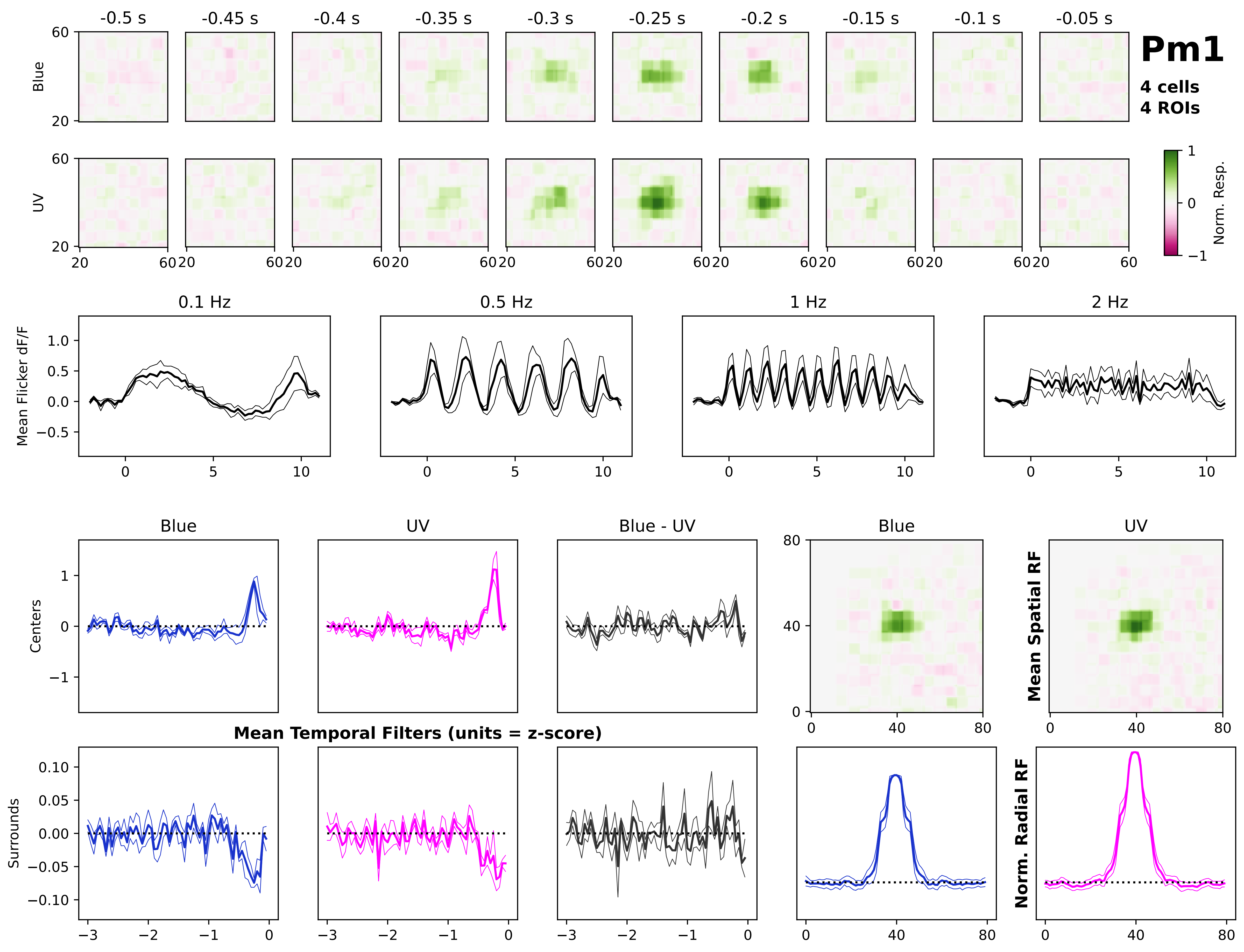

Pm1
4 cells
4 ROIs
-0.5 s
-0.45 s
-0.4 s
-0.35 s
-0.3 s
-0.25 s
-0.2 s
-0.15 s
-0.1 s
-0.05 s
Blue
UV
Norm. Resp.
0.1 Hz
0.5 Hz
1 Hz
2 Hz
Mean Flicker dF/F
Blue
UV
Blue - UV
Centers
Surrounds
Mean Temporal Filters (units = z-score)
Mean Spatial RF
Norm. Radial RF

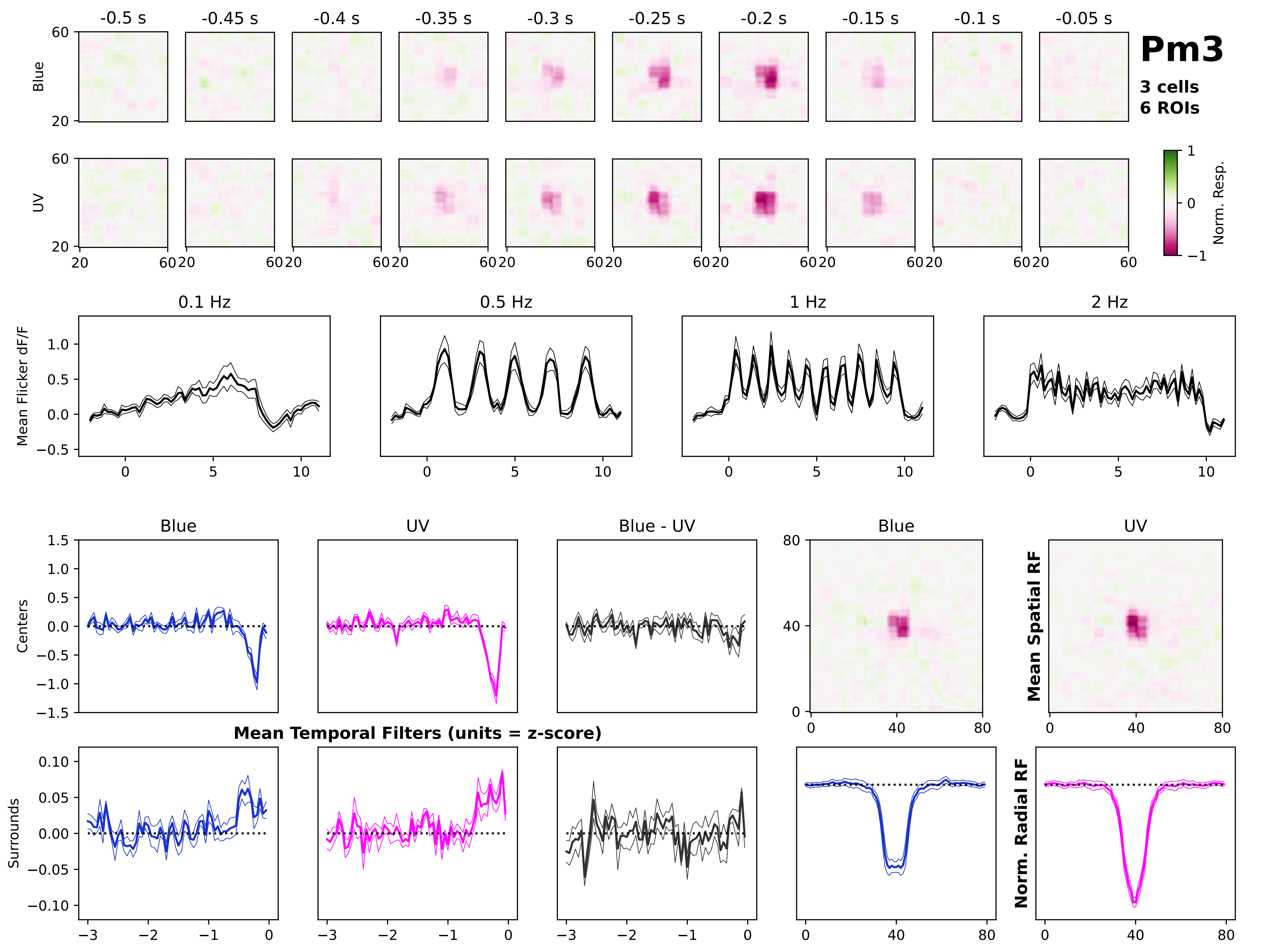
Pm3
3 cells
6 ROIs
-0.5 s
-0.45 s
-0.4 s
-0.35 s
-0.3 s
-0.25 s
-0.2 s
-0.15 s
-0.1 s
-0.05 s
Blue
UV
Norm. Resp.
0.1 Hz
0.5 Hz
1 Hz
2 Hz
Mean Flicker dF/F
Blue
UV
Blue - UV
Centers
Surrounds
Mean Temporal Filters (units = z-score)
Mean Spatial RF
Norm. Radial RF

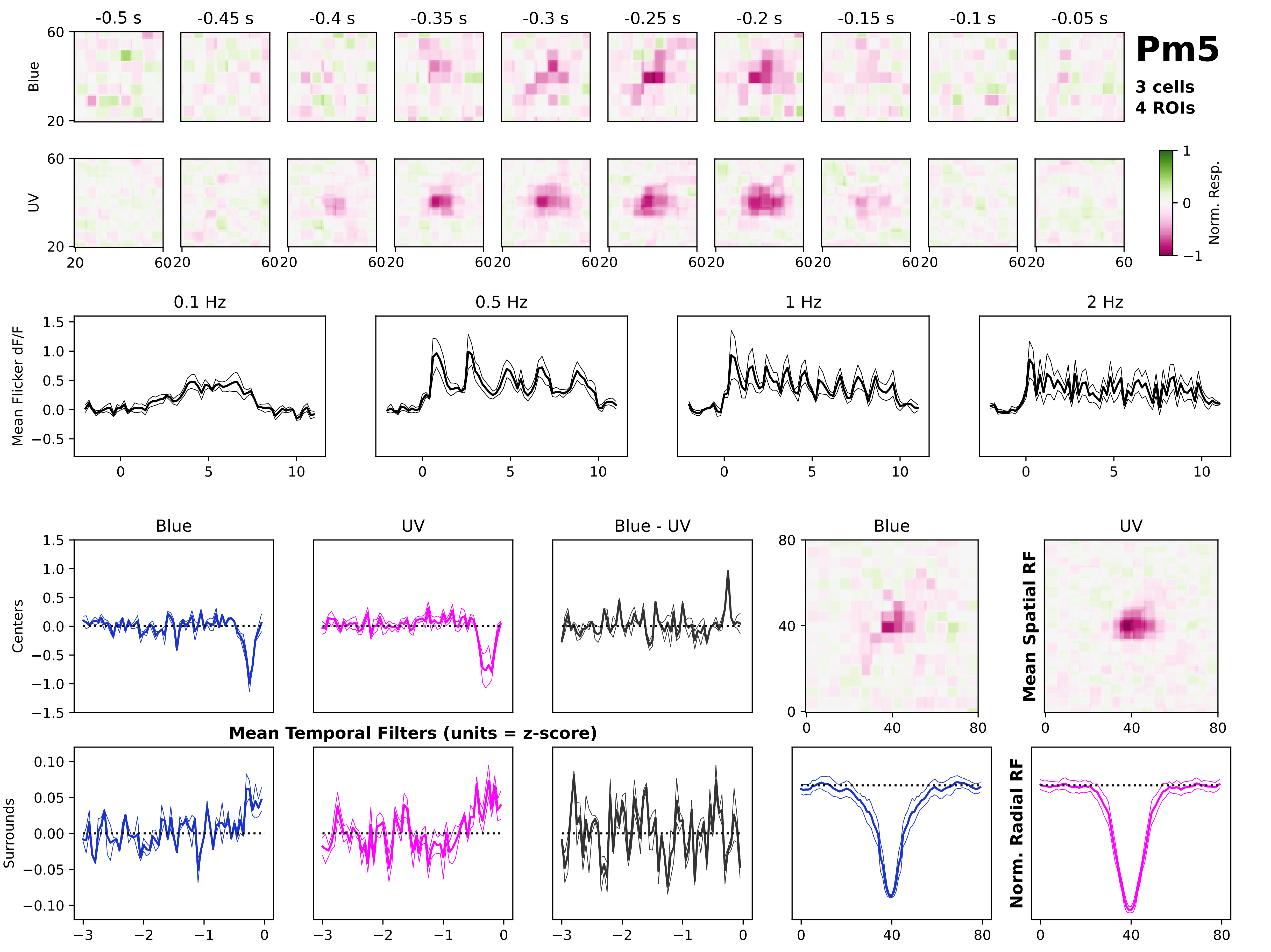
Pm5
3 cells
4 ROIs
-0.5 s
-0.45 s
-0.4 s
-0.35 s
-0.3 s
-0.25 s
-0.2 s
-0.15 s
-0.1 s
-0.05 s
Blue
UV
Norm. Resp.
0.1 Hz
0.5 Hz
1 Hz
2 Hz
Mean Flicker dF/F
Blue
UV
Blue - UV
Centers
Surrounds
Mean Temporal Filters (units = z-score)
Mean Spatial RF
Norm. Radial RF

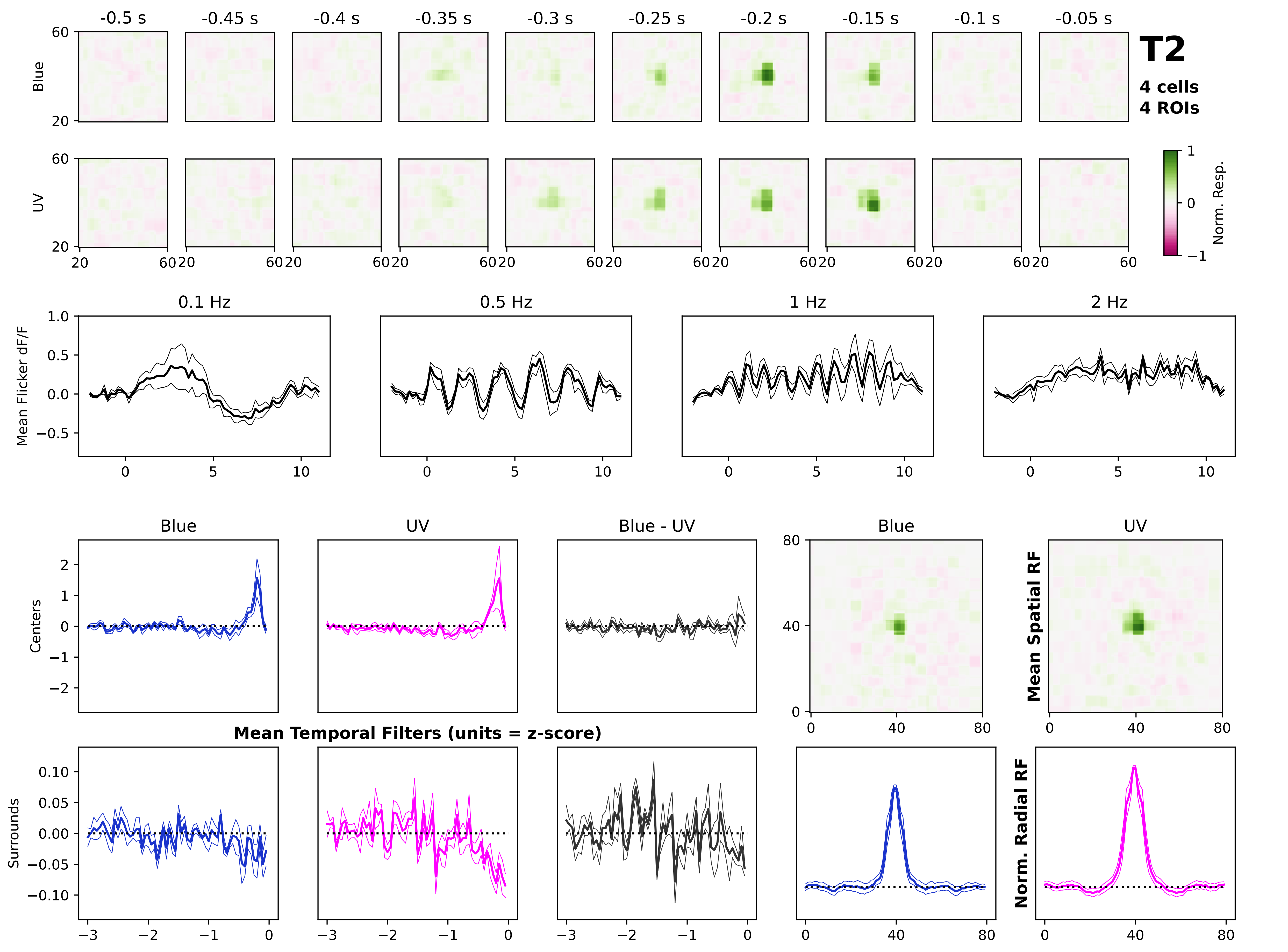
T2
4 cells
4 ROIs
-0.5 s
-0.45 s
-0.4 s
-0.35 s
-0.3 s
-0.25 s
-0.2 s
-0.15 s
-0.1 s
-0.05 s
Blue
UV
Norm. Resp.
0.1 Hz
0.5 Hz
1 Hz
2 Hz
Mean Flicker dF/F
Blue
UV
Blue - UV
Centers
Surrounds
Mean Temporal Filters (units = z-score)
Mean Spatial RF
Norm. Radial RF

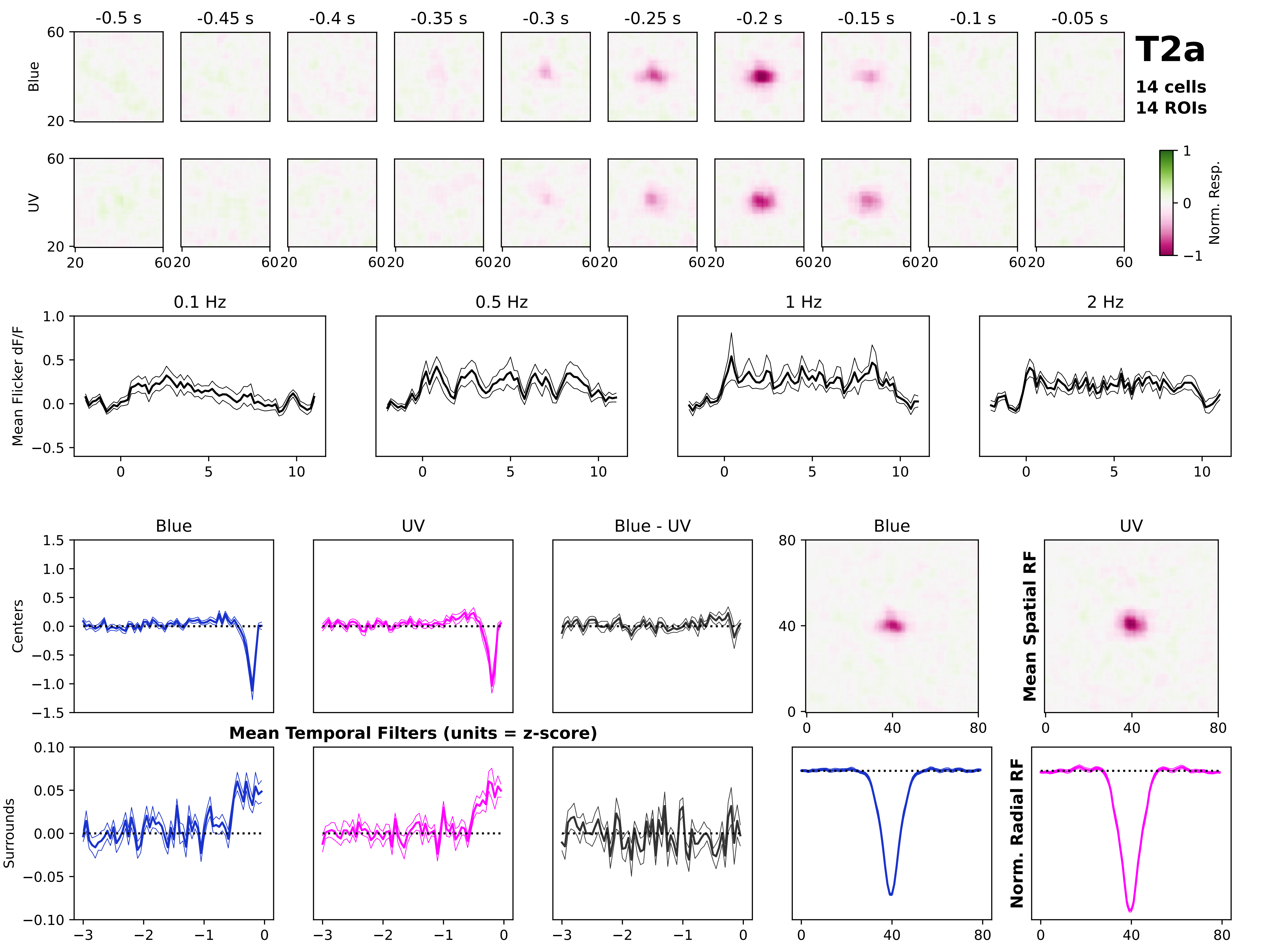
T2a
14 cells
14 ROIs
-0.5 s
-0.45 s
-0.4 s
-0.35 s
-0.3 s
-0.25 s
-0.2 s
-0.15 s
-0.1 s
-0.05 s
Blue
UV
Norm. Resp.
Mean Flicker dF/F
0.1 Hz
0.5 Hz
1 Hz
2 Hz
Centers
Surrounds
Blue
UV
Blue - UV
Mean Temporal Filters (units = z-score)
Mean Spatial RF
Norm. Radial RF

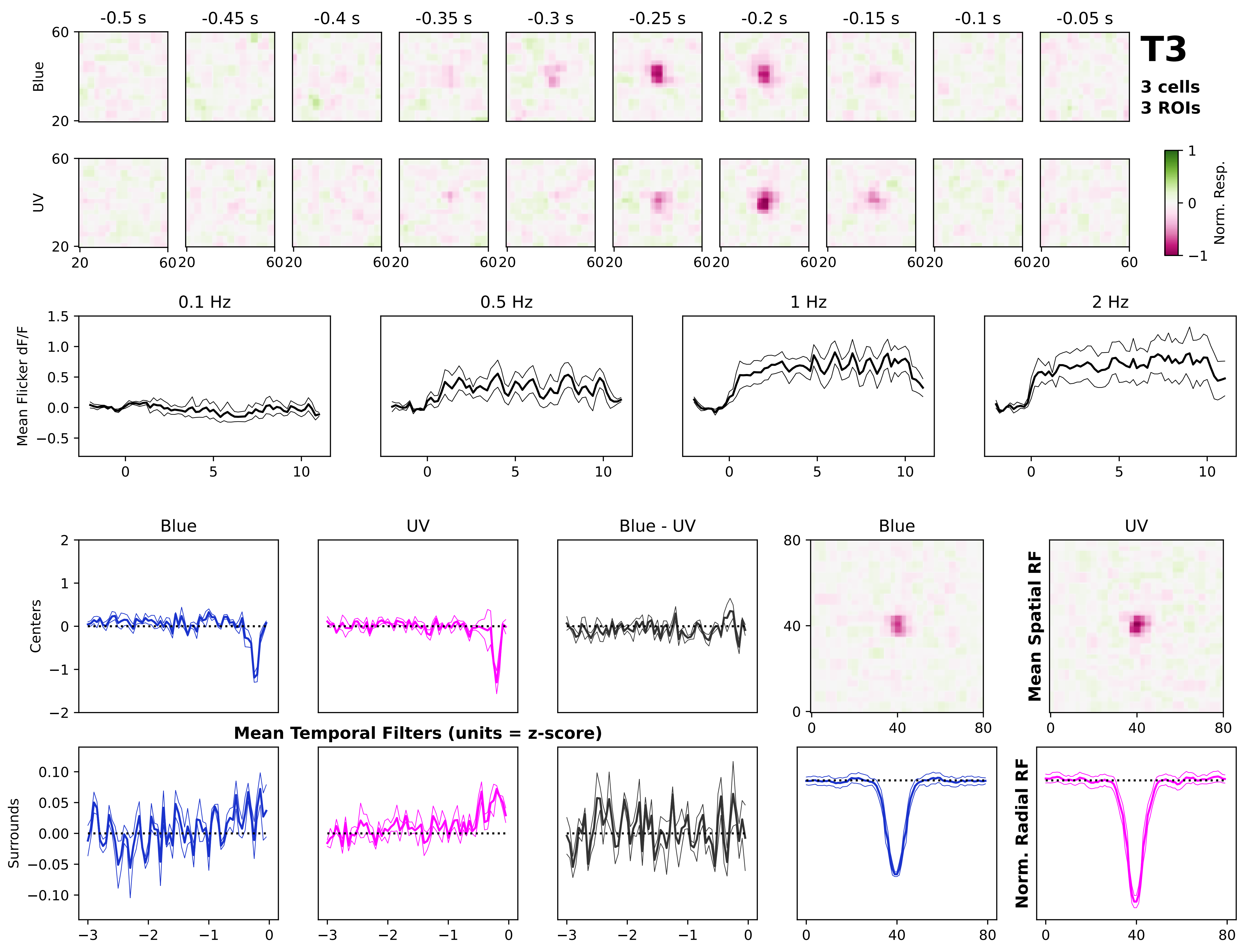

T3
3 cells
3 ROIs
-0.5 s
-0.45 s
-0.4 s
-0.35 s
-0.3 s
-0.25 s
-0.2 s
-0.15 s
-0.1 s
-0.05 s
Blue
UV
Norm. Resp.
0.1 Hz
0.5 Hz
1 Hz
2 Hz
Mean Flicker dF/F
Blue
UV
Blue - UV
Centers
Surrounds
Mean Temporal Filters (units = z-score)
Mean Spatial RF
Norm. Radial RF

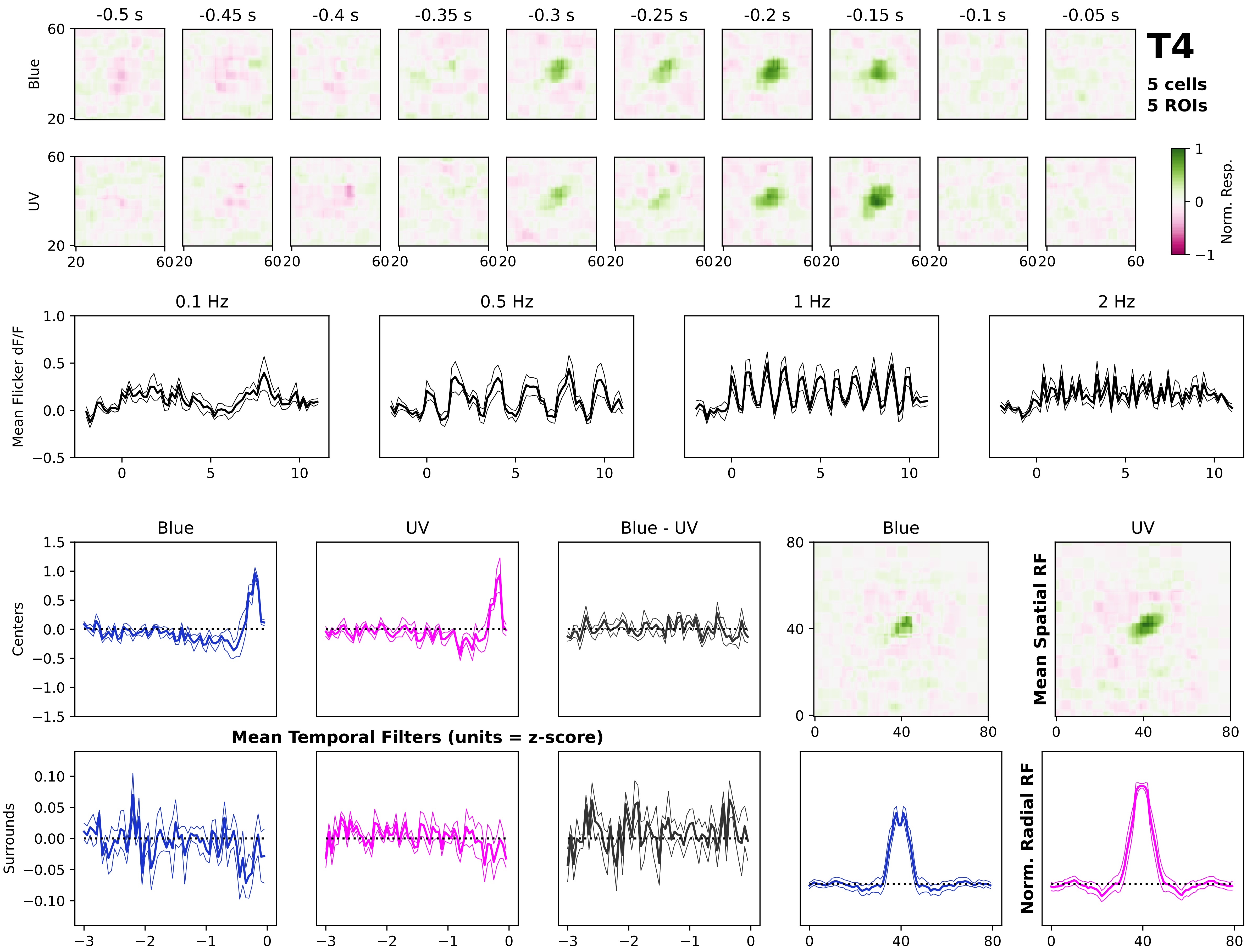
T4
5 cells
5 ROIs
-0.5 s
-0.45 s
-0.4 s
-0.35 s
-0.3 s
-0.25 s
-0.2 s
-0.15 s
-0.1 s
-0.05 s
Blue
UV
Norm. Resp.
0.1 Hz
0.5 Hz
1 Hz
2 Hz
Mean Flicker dF/F
Blue
UV
Blue - UV
Centers
Surrounds
Mean Temporal Filters (units = z-score)
Mean Spatial RF
Norm. Radial RF

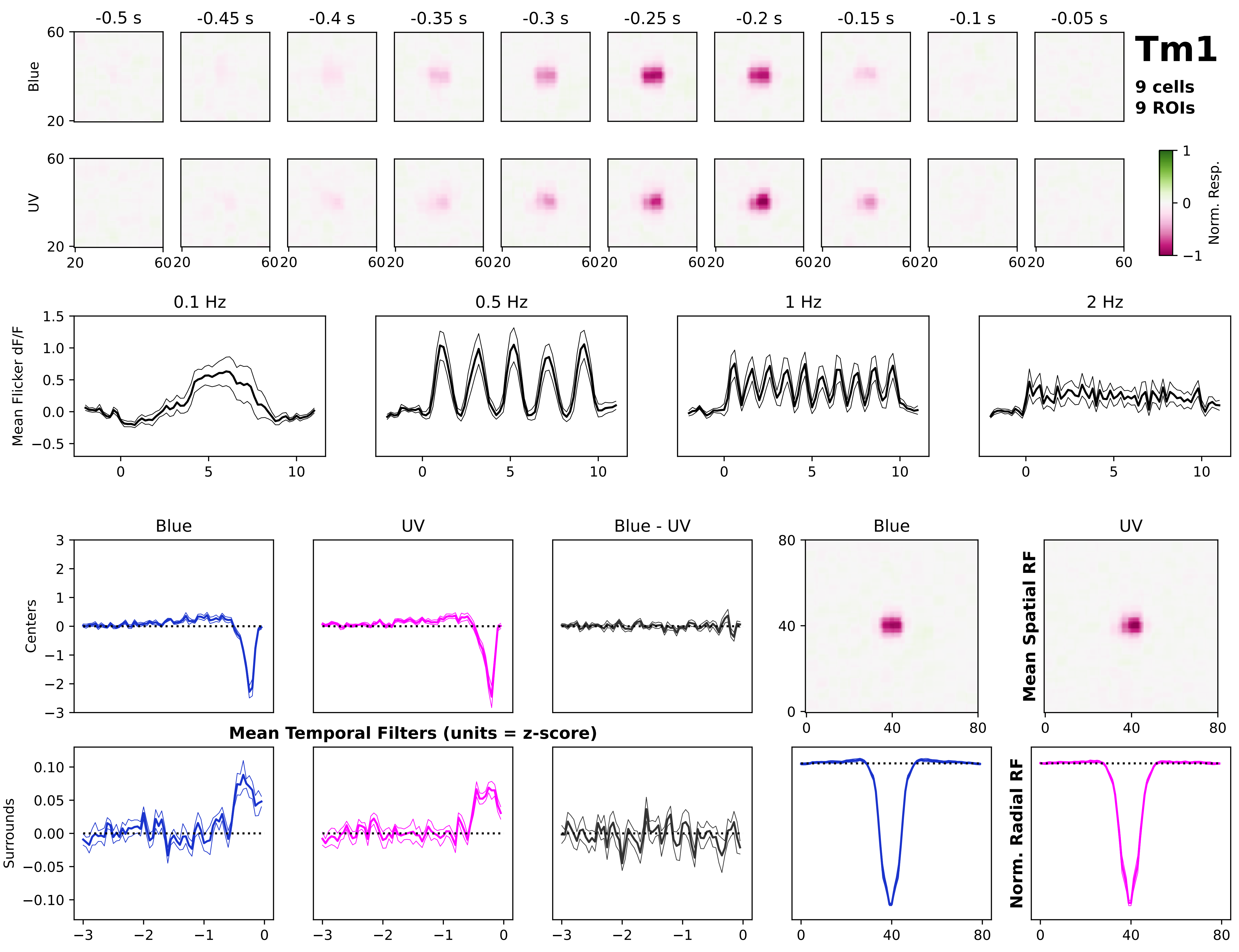
Tm1
9 cells
9 ROIs
-0.5 s
-0.45 s
-0.4 s
-0.35 s
-0.3 s
-0.25 s
-0.2 s
-0.15 s
-0.1 s
-0.05 s
Blue
UV
Norm. Resp.
0.1 Hz
0.5 Hz
1 Hz
2 Hz
Mean Flicker dF/F
Blue
UV
Blue - UV
Centers
Surrounds
Mean Temporal Filters (units = z-score)
Mean Spatial RF
Norm. Radial RF

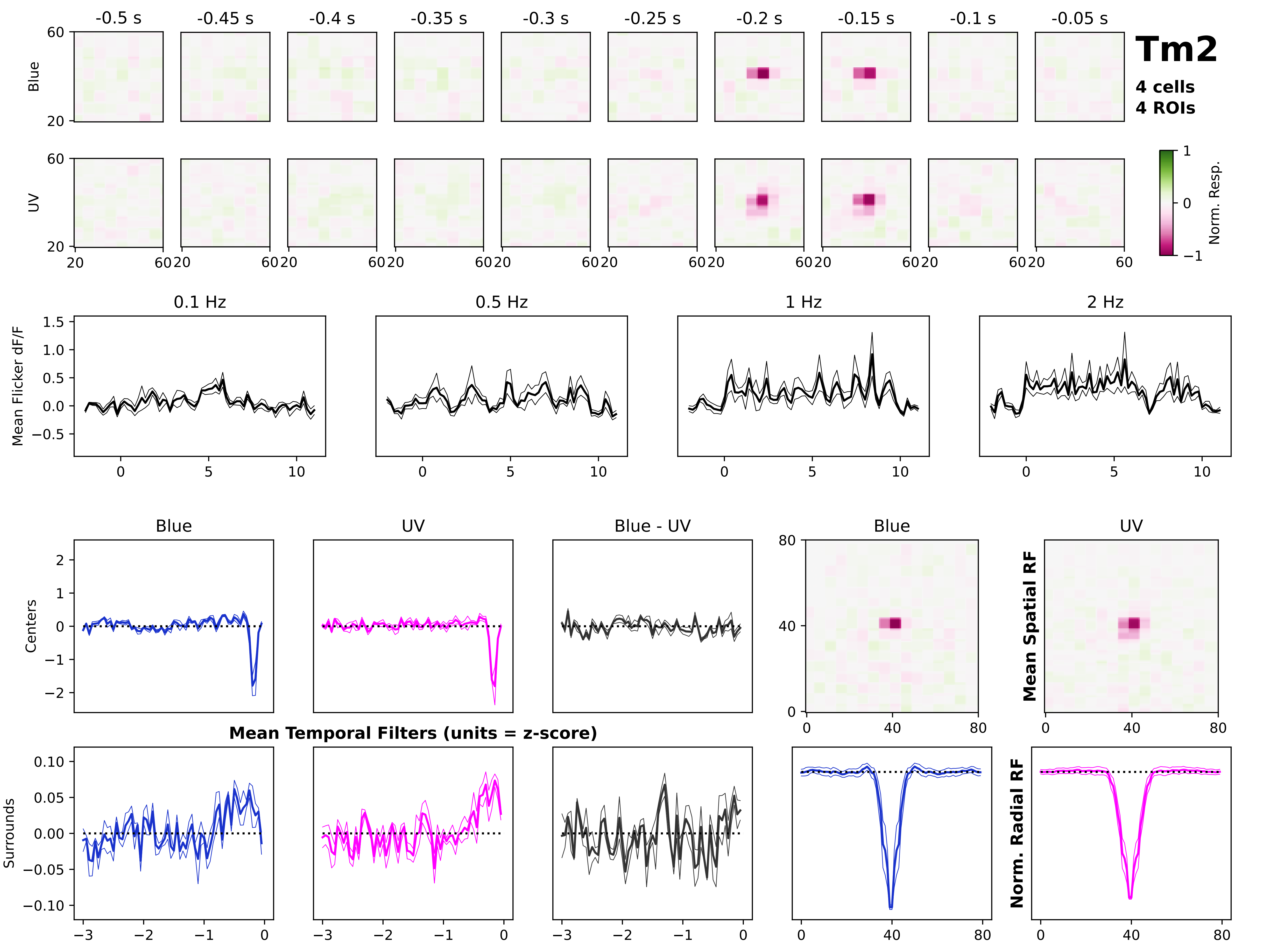

Tm2
4 cells
4 ROIs
-0.5 s
-0.45 s
-0.4 s
-0.35 s
-0.3 s
-0.25 s
-0.2 s
-0.15 s
-0.1 s
-0.05 s
Blue
UV
Norm. Resp.
0.1 Hz
0.5 Hz
1 Hz
2 Hz
Mean Flicker dF/F
Blue
UV
Blue - UV
Centers
Surrounds
Mean Temporal Filters (units = z-score)
Mean Spatial RF
Norm. Radial RF

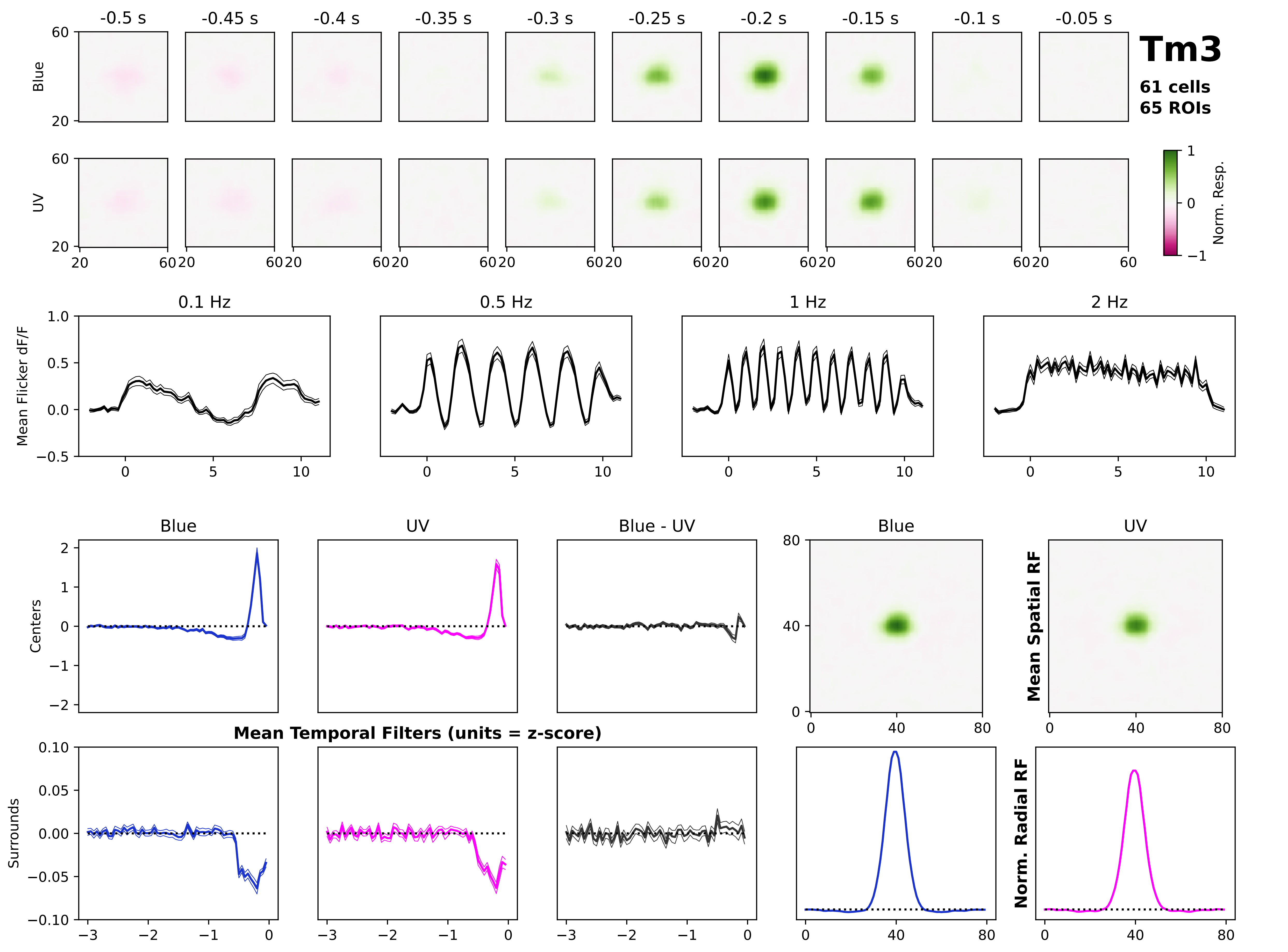

Tm3
61 cells
65 ROIs
-0.5 s
-0.45 s
-0.4 s
-0.35 s
-0.3 s
-0.25 s
-0.2 s
-0.15 s
-0.1 s
-0.05 s
Blue
UV
Norm. Resp.
0.1 Hz
0.5 Hz
1 Hz
2 Hz
Mean Flicker dF/F
Blue
UV
Blue - UV
Centers
Surrounds
Mean Temporal Filters (units = z-score)
Mean Spatial RF
Norm. Radial RF

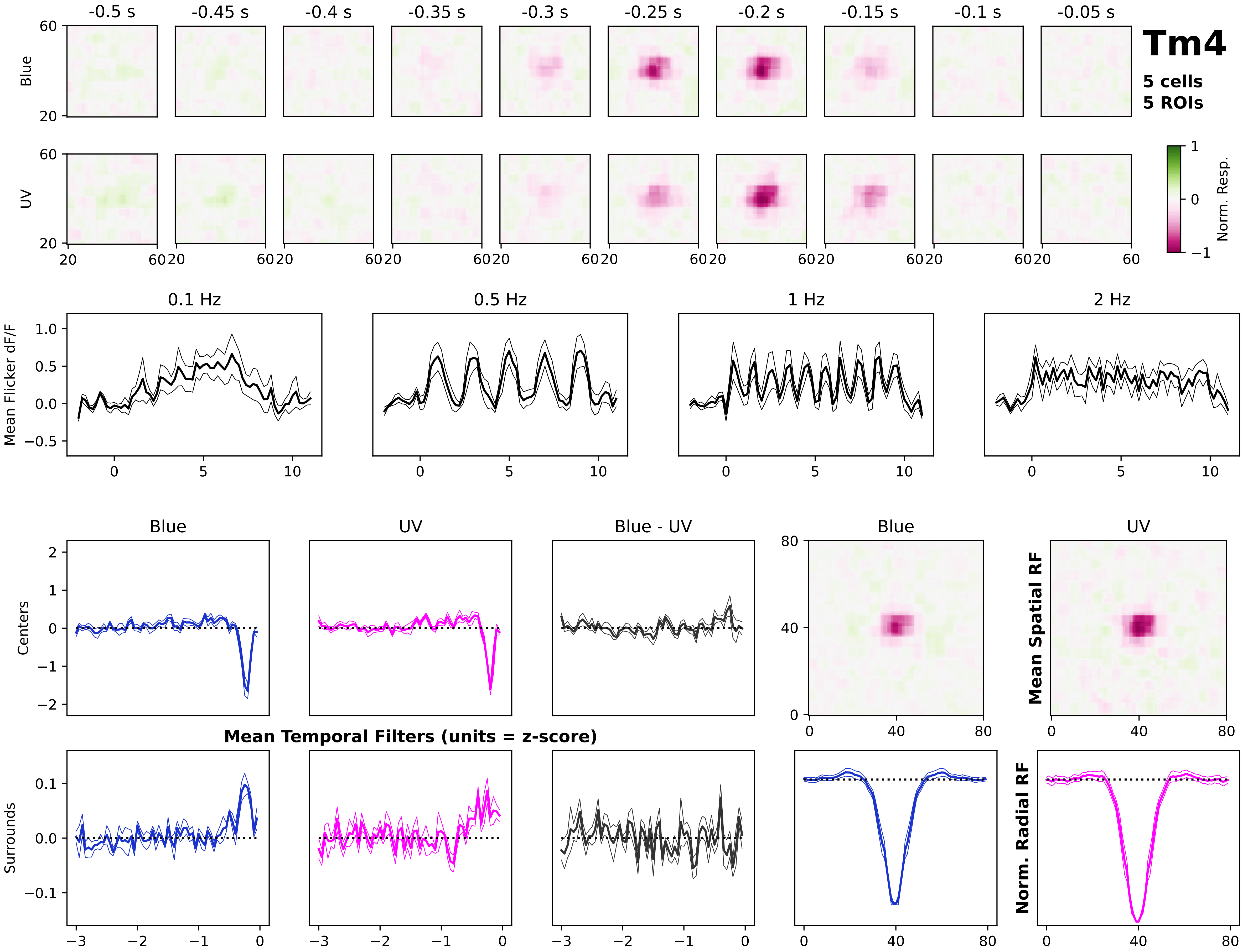
Tm4
5 cells
5 ROIs
-0.5 s
-0.45 s
-0.4 s
-0.35 s
-0.3 s
-0.25 s
-0.2 s
-0.15 s
-0.1 s
-0.05 s
Blue
UV
Norm. Resp.
0.1 Hz
0.5 Hz
1 Hz
2 Hz
Mean Flicker dF/F
Centers
Surrounds
Blue
UV
Blue - UV
Mean Temporal Filters (units = z-score)
Mean Spatial RF
Norm. Radial RF

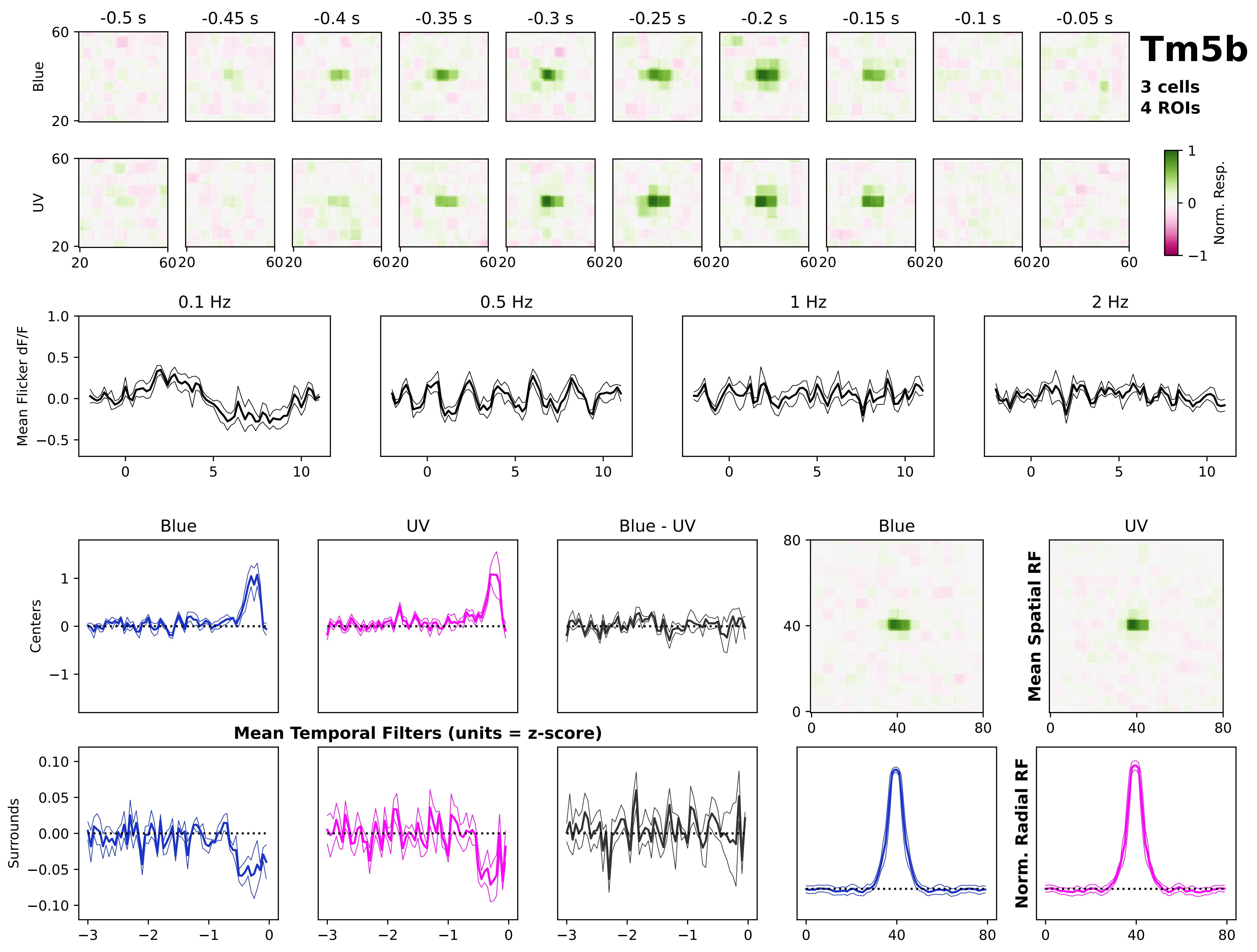

Tm5b
3 cells
4 ROIs
-0.5 s
-0.45 s
-0.4 s
-0.35 s
-0.3 s
-0.25 s
-0.2 s
-0.15 s
-0.1 s
-0.05 s
Blue
UV
Norm. Resp.
0.1 Hz
0.5 Hz
1 Hz
2 Hz
Mean Flicker dF/F
Blue
UV
Blue - UV
Centers
Surrounds
Mean Temporal Filters (units = z-score)
Mean Spatial RF
Norm. Radial RF

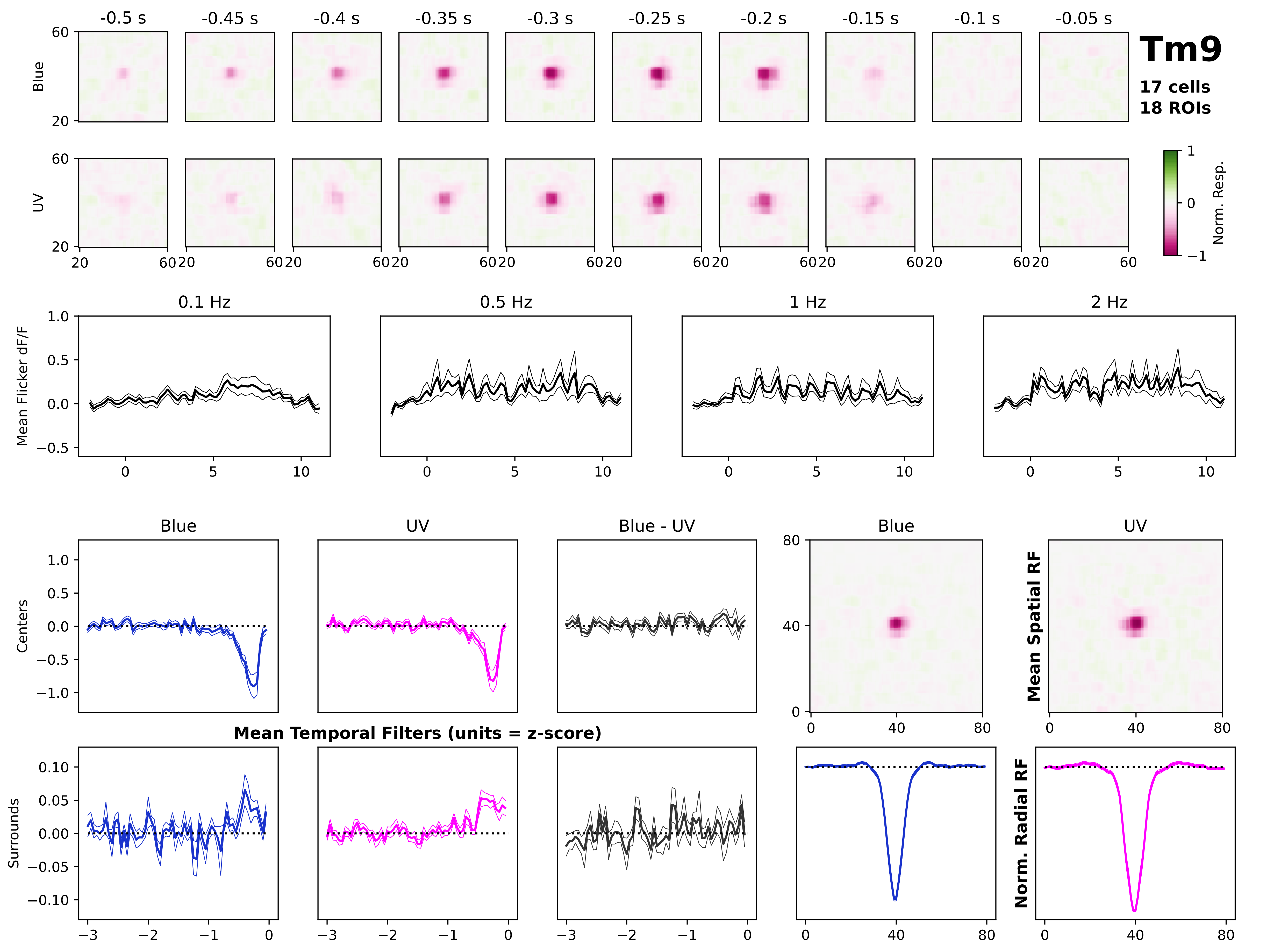

Tm9
17 cells
18 ROIs
-0.5 s
-0.45 s
-0.4 s
-0.35 s
-0.3 s
-0.25 s
-0.2 s
-0.15 s
-0.1 s
-0.05 s
Blue
UV
Norm. Resp.
0.1 Hz
0.5 Hz
1 Hz
2 Hz
Mean Flicker dF/F
Blue
UV
Blue - UV
Centers
Surrounds
Mean Temporal Filters (units = z-score)
Mean Spatial RF
Norm. Radial RF

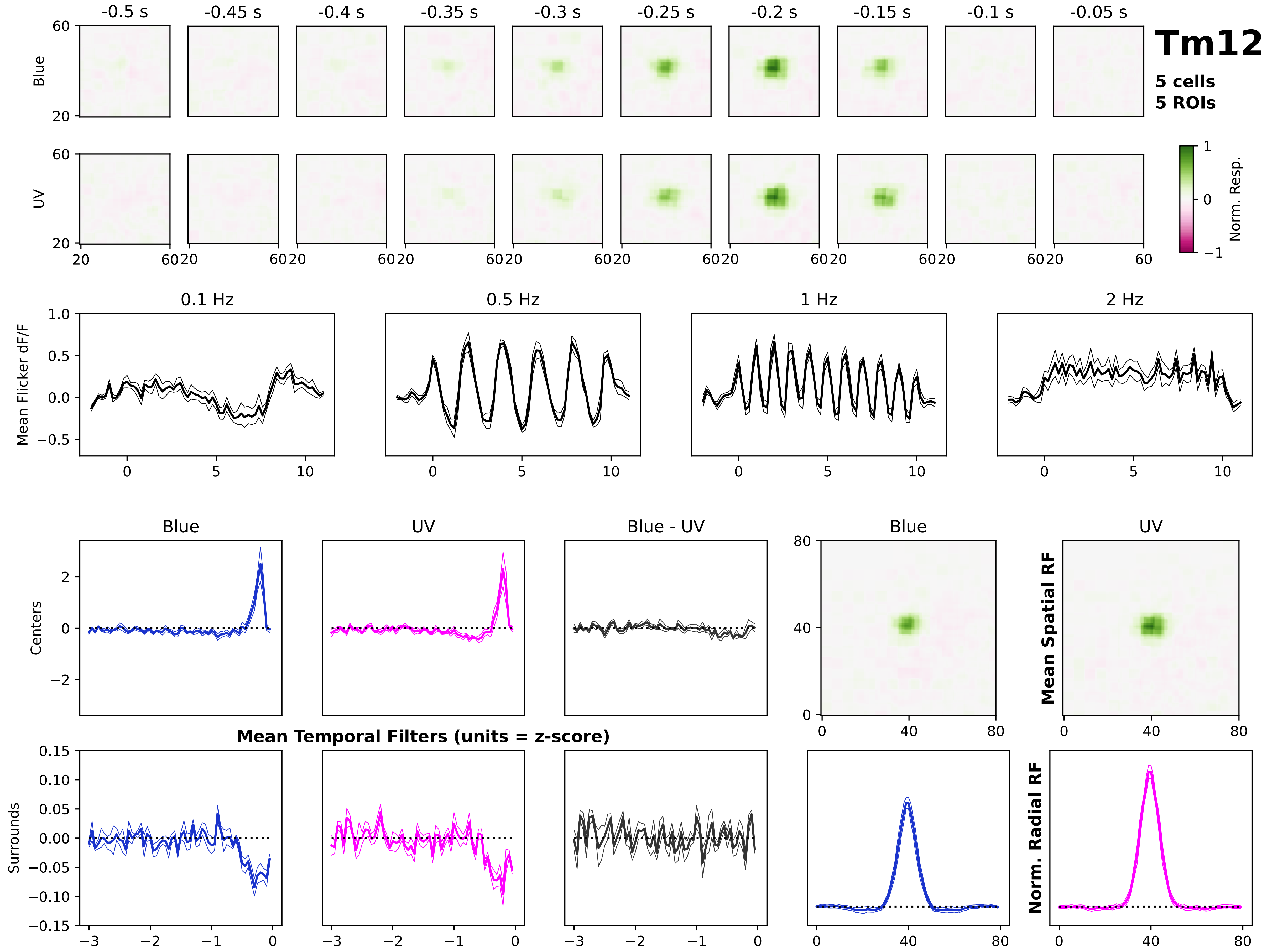
Tm12
5 cells
5 ROIs
-0.5 s
-0.45 s
-0.4 s
-0.35 s
-0.3 s
-0.25 s
-0.2 s
-0.15 s
-0.1 s
-0.05 s
Blue
UV
Norm. Resp.
0.1 Hz
0.5 Hz
1 Hz
2 Hz
Mean Flicker dF/F
Blue
UV
Blue - UV
Centers
Surrounds
Mean Temporal Filters (units = z-score)
Mean Spatial RF
Norm. Radial RF

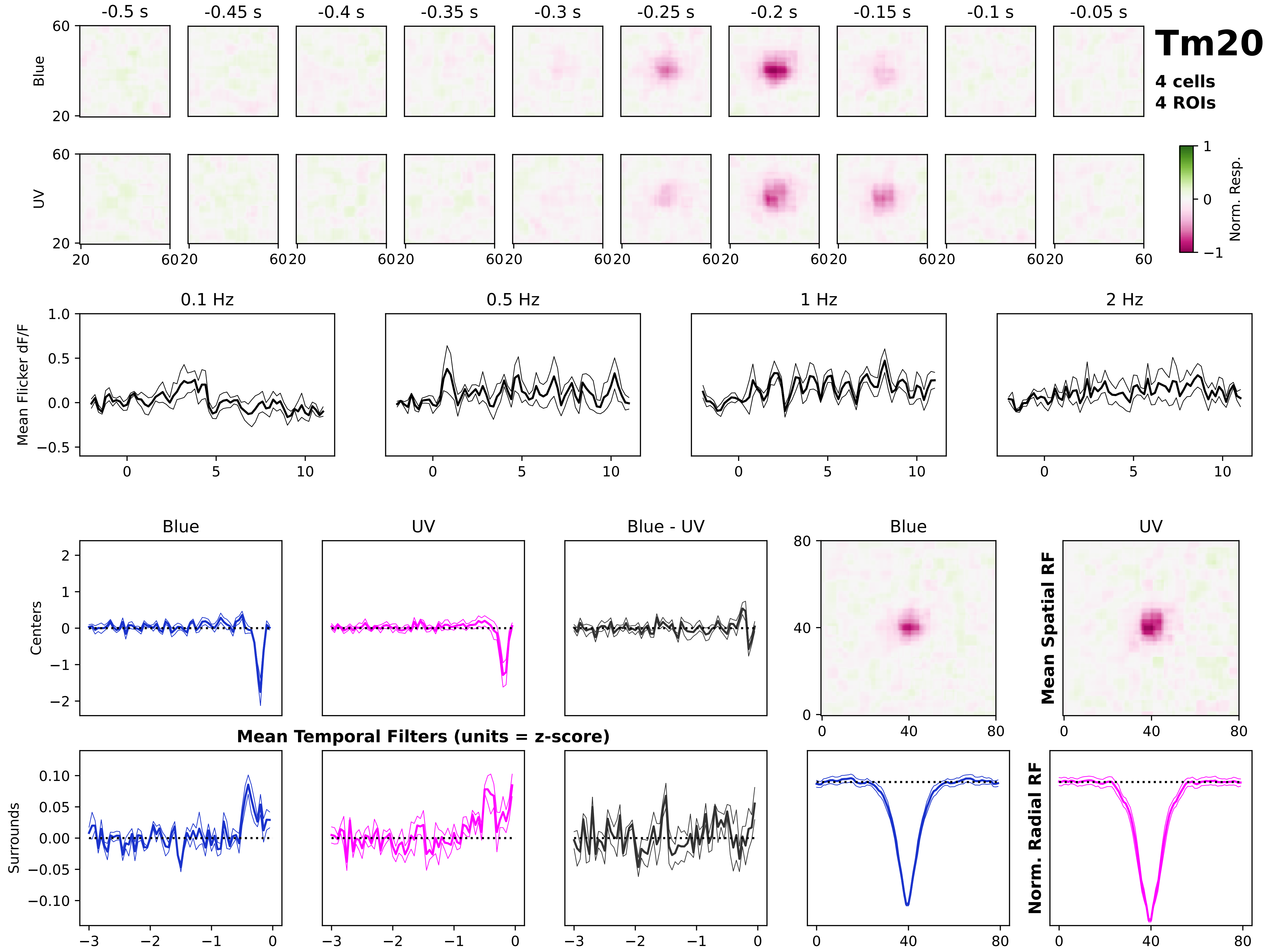
Tm20
4 cells
4 ROIs
-0.5 s
-0.45 s
-0.4 s
-0.35 s
-0.3 s
-0.25 s
-0.2 s
-0.15 s
-0.1 s
-0.05 s
Blue
UV
Norm. Resp.
0.1 Hz
0.5 Hz
1 Hz
2 Hz
Mean Flicker dF/F
Blue
UV
Blue - UV
Centers
Surrounds
Mean Temporal Filters (units = z-score)
Mean Spatial RF
Norm. Radial RF

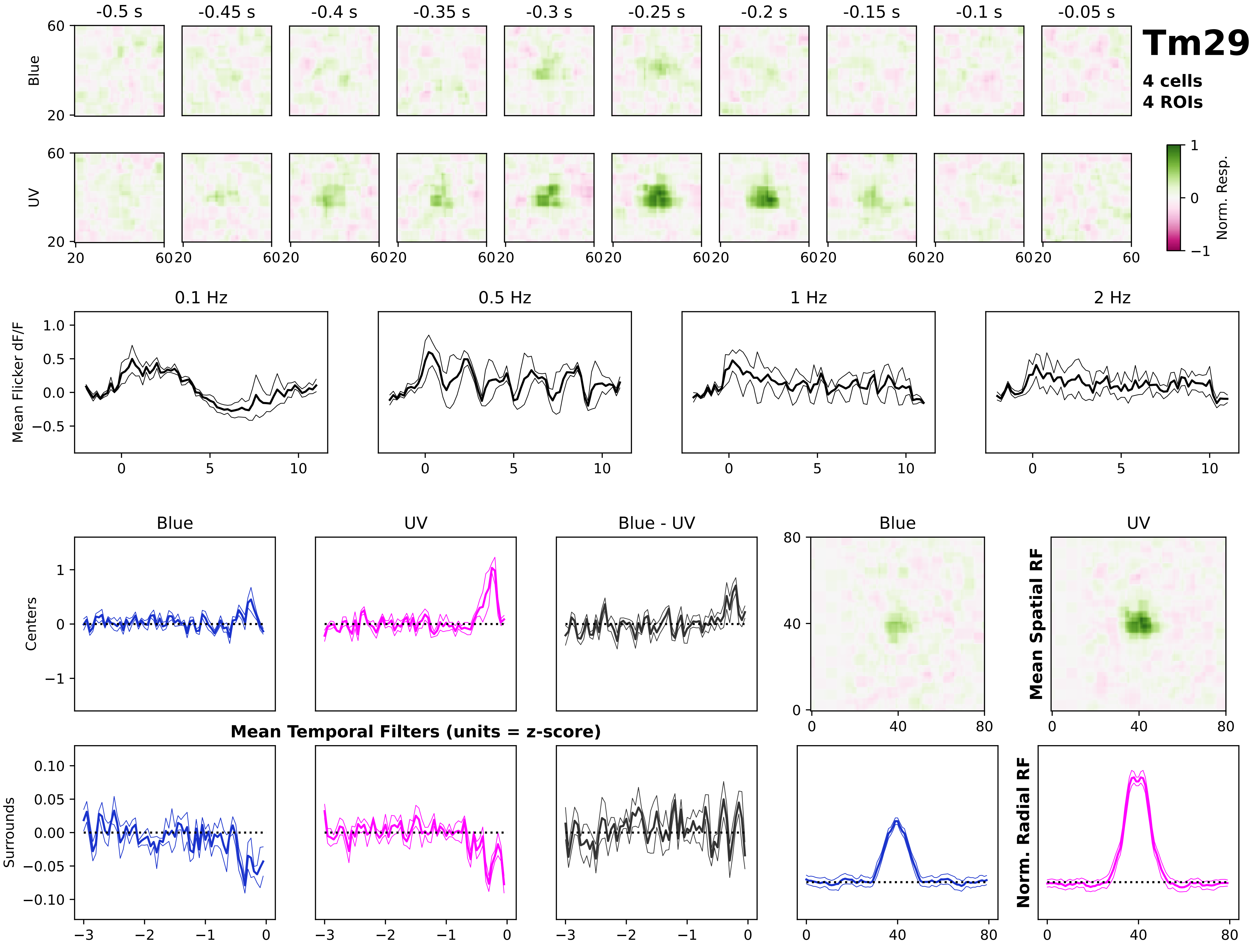
Tm29
4 cells
4 ROIs
-0.5 s
-0.45 s
-0.4 s
-0.35 s
-0.3 s
-0.25 s
-0.2 s
-0.15 s
-0.1 s
-0.05 s
Blue
UV
Norm. Resp.
0.1 Hz
0.5 Hz
1 Hz
2 Hz
Mean Flicker dF/F
Blue
UV
Blue - UV
Centers
Surrounds
Mean Temporal Filters (units = z-score)
Mean Spatial RF
Norm. Radial RF

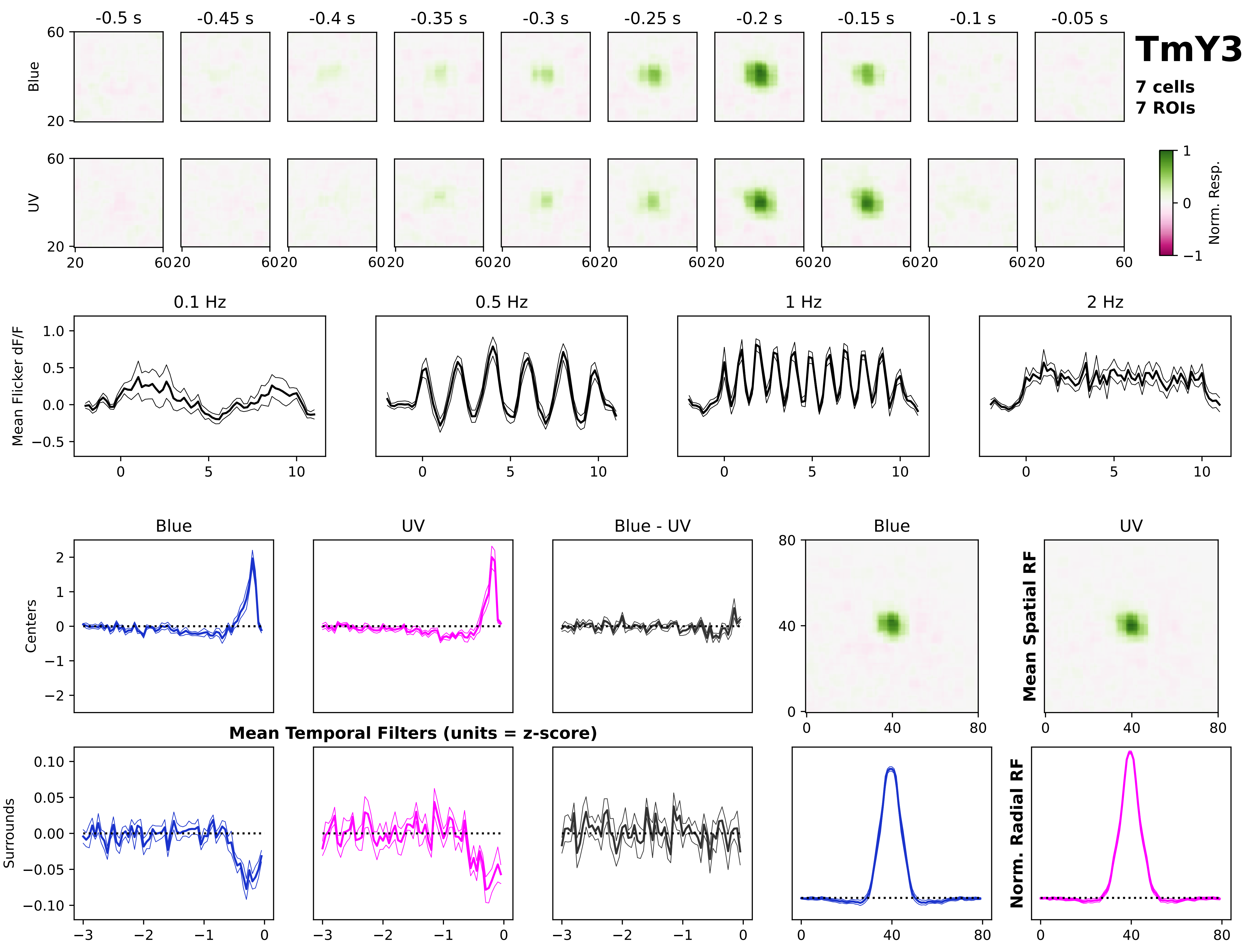

TmY3
7 cells
7 ROIs
-0.5 s
-0.45 s
-0.4 s
-0.35 s
-0.3 s
-0.25 s
-0.2 s
-0.15 s
-0.1 s
-0.05 s
Blue
UV
Norm. Resp.
0.1 Hz
0.5 Hz
1 Hz
2 Hz
Mean Flicker dF/F
Blue
UV
Blue - UV
Centers
Surrounds
Mean Temporal Filters (units = z-score)
Mean Spatial RF
Norm. Radial RF

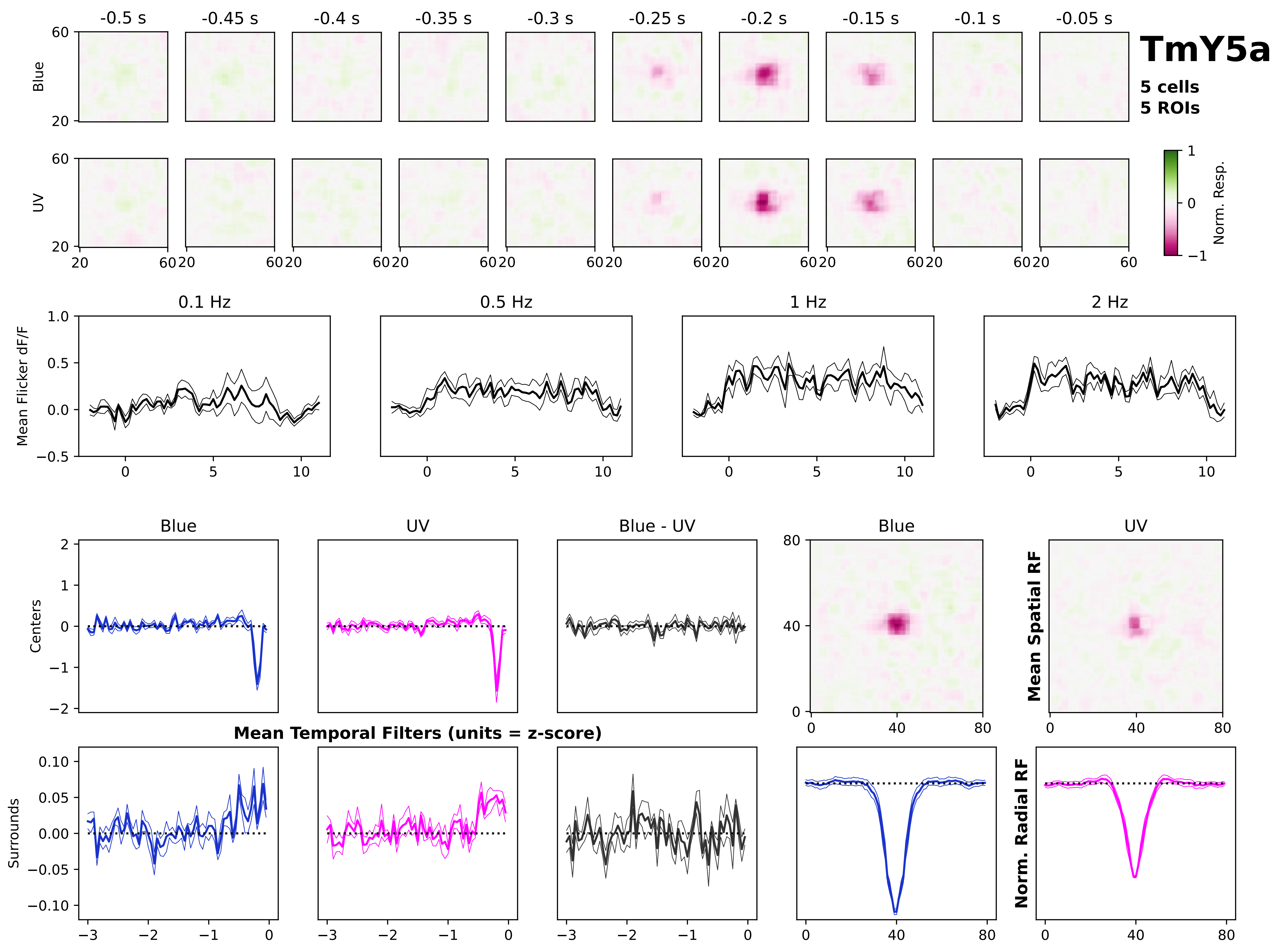
TmY5a
5 cells
5 ROIs
-0.5 s
-0.45 s
-0.4 s
-0.35 s
-0.3 s
-0.25 s
-0.2 s
-0.15 s
-0.1 s
-0.05 s
Blue
UV
Norm. Resp.
0.1 Hz
0.5 Hz
1 Hz
2 Hz
Mean Flicker dF/F
Blue
UV
Blue - UV
Centers
Surrounds
Mean Temporal Filters (units = z-score)
Mean Spatial RF
Norm. Radial RF

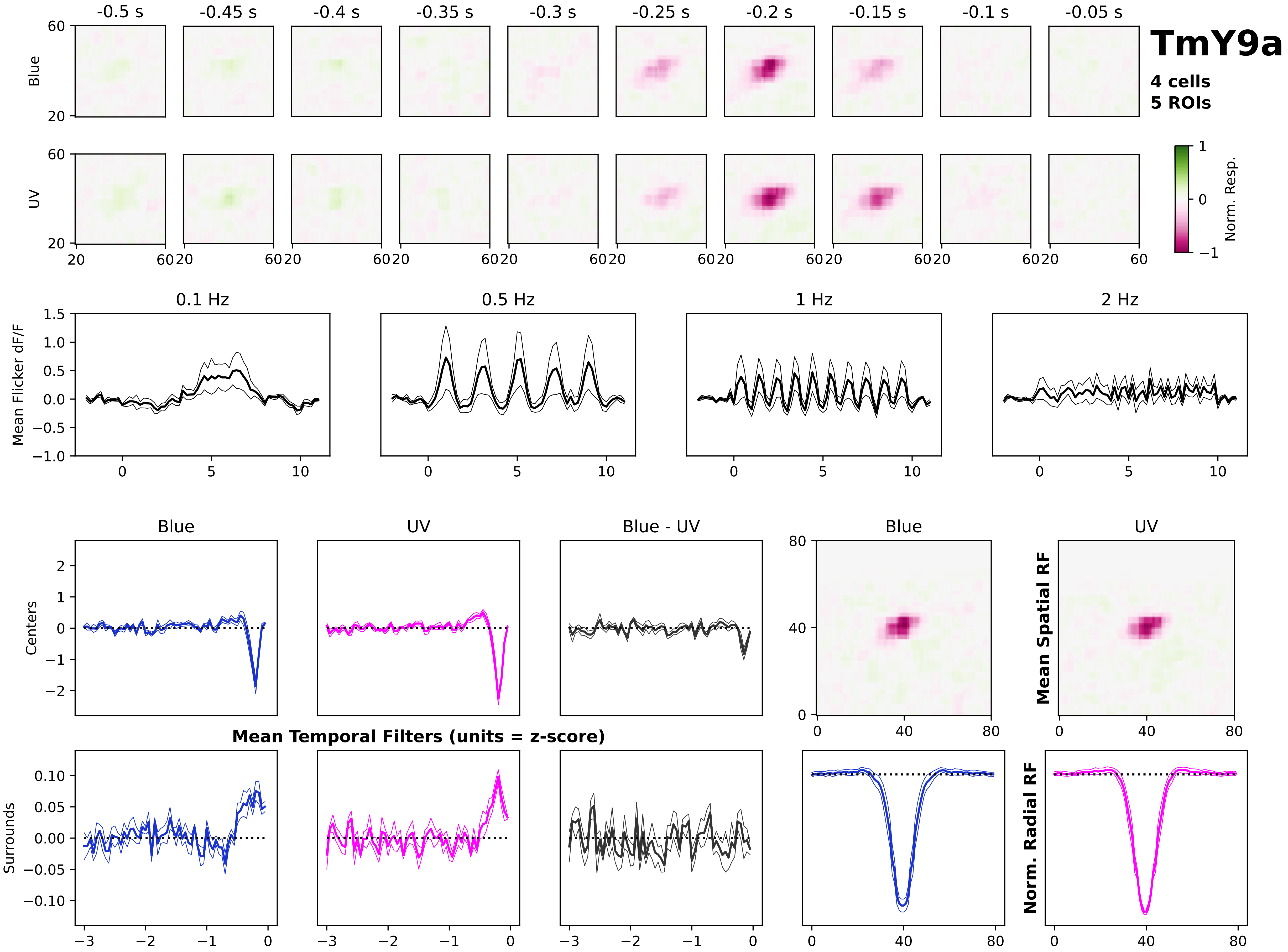

TmY9a
4 cells
5 ROIs
-0.5 s
-0.45 s
-0.4 s
-0.35 s
-0.3 s
-0.25 s
-0.2 s
-0.15 s
-0.1 s
-0.05 s
Blue
UV
Norm. Resp.
0.1 Hz
0.5 Hz
1 Hz
2 Hz
Mean Flicker dF/F
Centers
Surrounds
Blue - UV
Mean Temporal Filters (units = z-score)
Mean Spatial RF
Norm. Radial RF

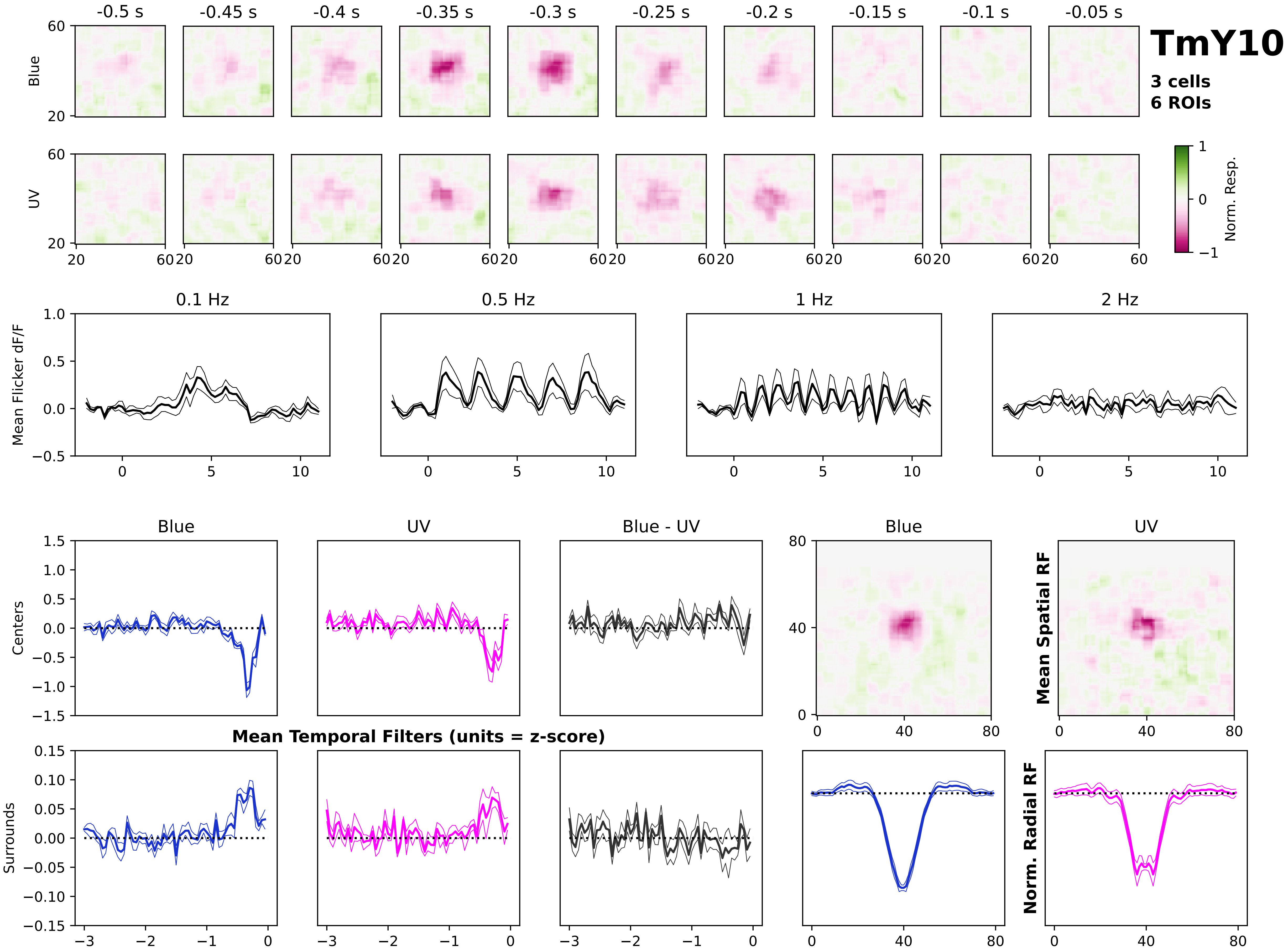
TmY10
3 cells
6 ROIs
-0.5 s
-0.45 s
-0.4 s
-0.35 s
-0.3 s
-0.25 s
-0.2 s
-0.15 s
-0.1 s
-0.05 s
Blue
UV
Norm. Resp.
0.1 Hz
0.5 Hz
1 Hz
2 Hz
Mean Flicker dF/F
Blue
UV
Blue - UV
Centers
Surrounds
Mean Temporal Filters (units = z-score)
Mean Spatial RF
Norm. Radial RF

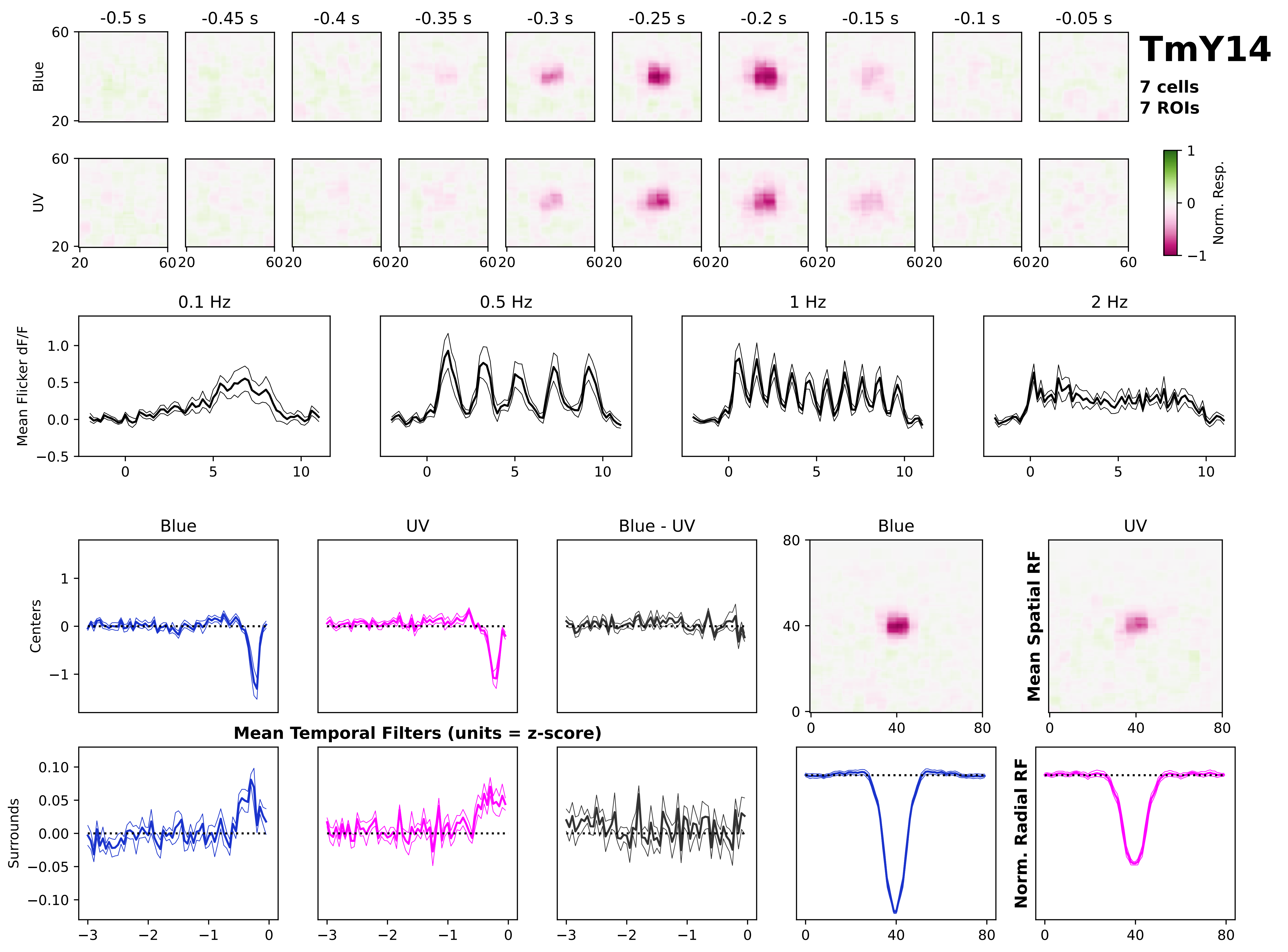
TmY14
7 cells
7 ROIs
-0.5 s
-0.45 s
-0.4 s
-0.35 s
-0.3 s
-0.25 s
-0.2 s
-0.15 s
-0.1 s
-0.05 s
Blue
UV
Norm. Resp.
0.1 Hz
0.5 Hz
1 Hz
2 Hz
Mean Flicker dF/F
Blue
UV
Blue - UV
Centers
Surrounds
Mean Temporal Filters (units = z-score)
Mean Spatial RF
Norm. Radial RF

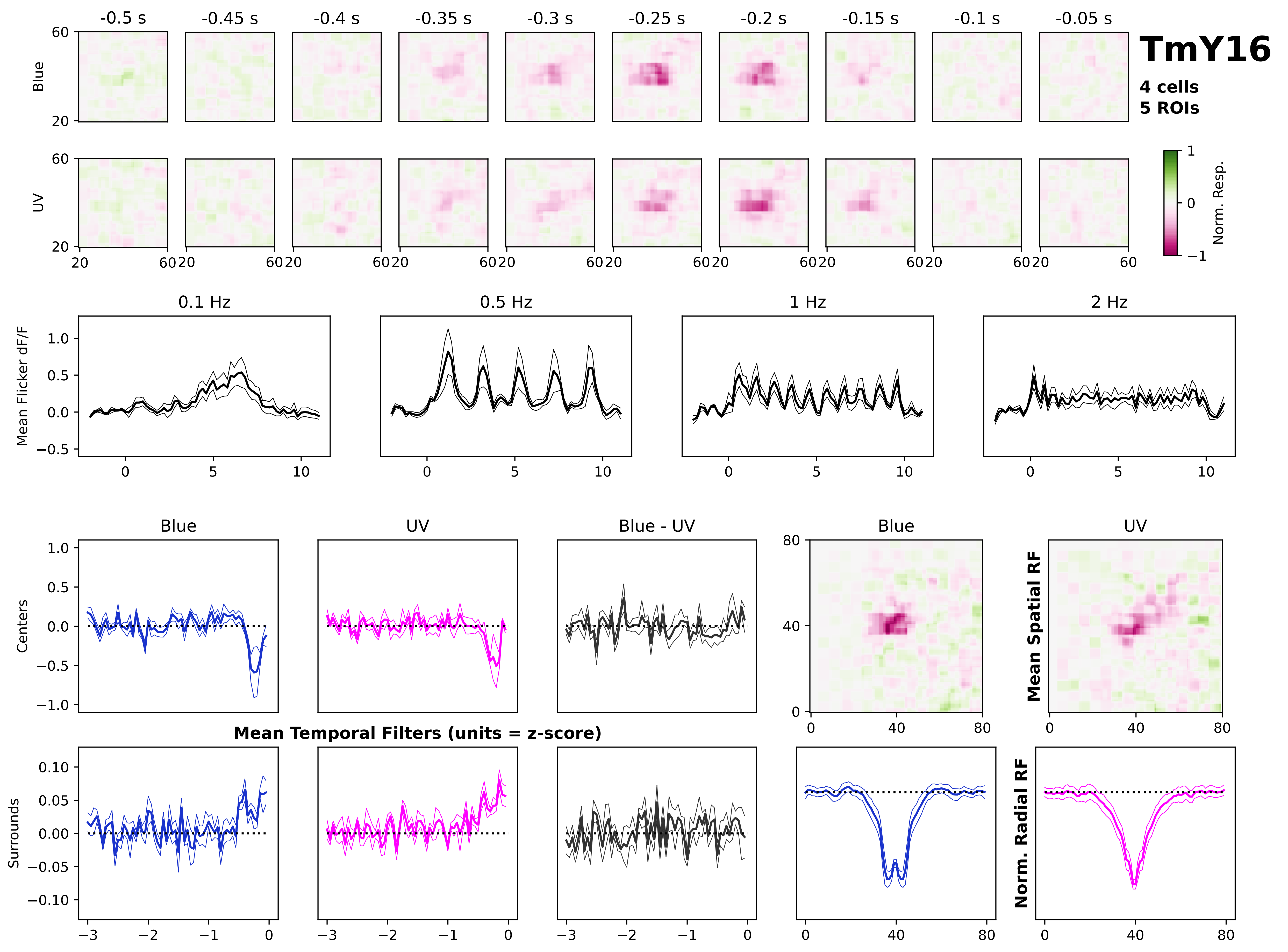
TmY16
4 cells
5 ROIs
-0.5 s
-0.45 s
-0.4 s
-0.35 s
-0.3 s
-0.25 s
-0.2 s
-0.15 s
-0.1 s
-0.05 s
Blue
UV
Norm. Resp.
0.1 Hz
0.5 Hz
1 Hz
2 Hz
Mean Flicker dF/F
Blue
UV
Blue - UV
Centers
Surrounds
Mean Temporal Filters (units = z-score)
Mean Spatial RF
Norm. Radial RF

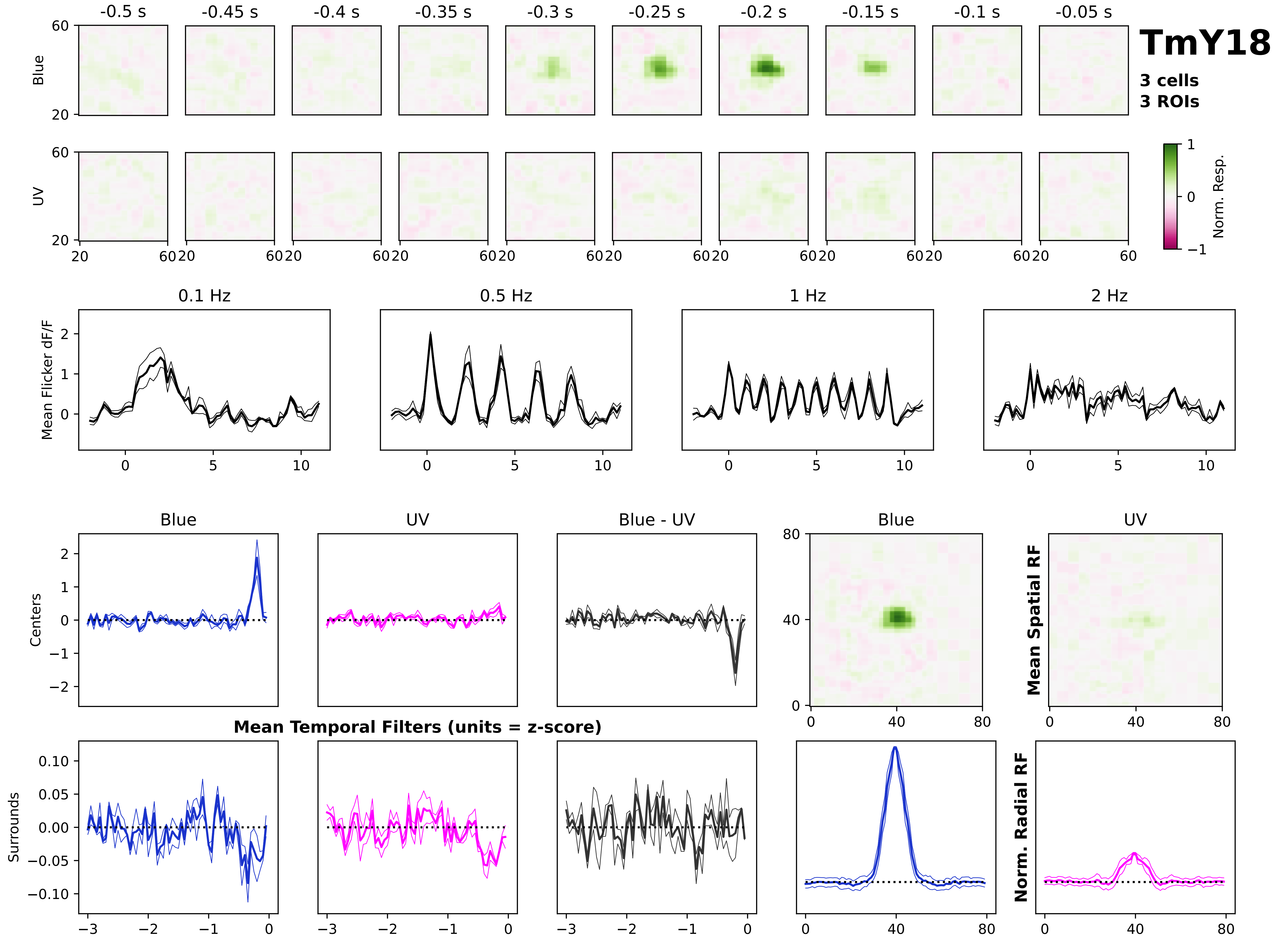

TmY18
3 cells
3 ROIs
-0.5 s
-0.45 s
-0.4 s
-0.35 s
-0.3 s
-0.25 s
-0.2 s
-0.15 s
-0.1 s
-0.05 s
Blue
UV
Norm. Resp.
0.1 Hz
0.5 Hz
1 Hz
2 Hz
Mean Flicker dF/F
Blue
UV
Blue - UV
Centers
Surrounds
Mean Temporal Filters (units = z-score)
Mean Spatial RF
Norm. Radial RF

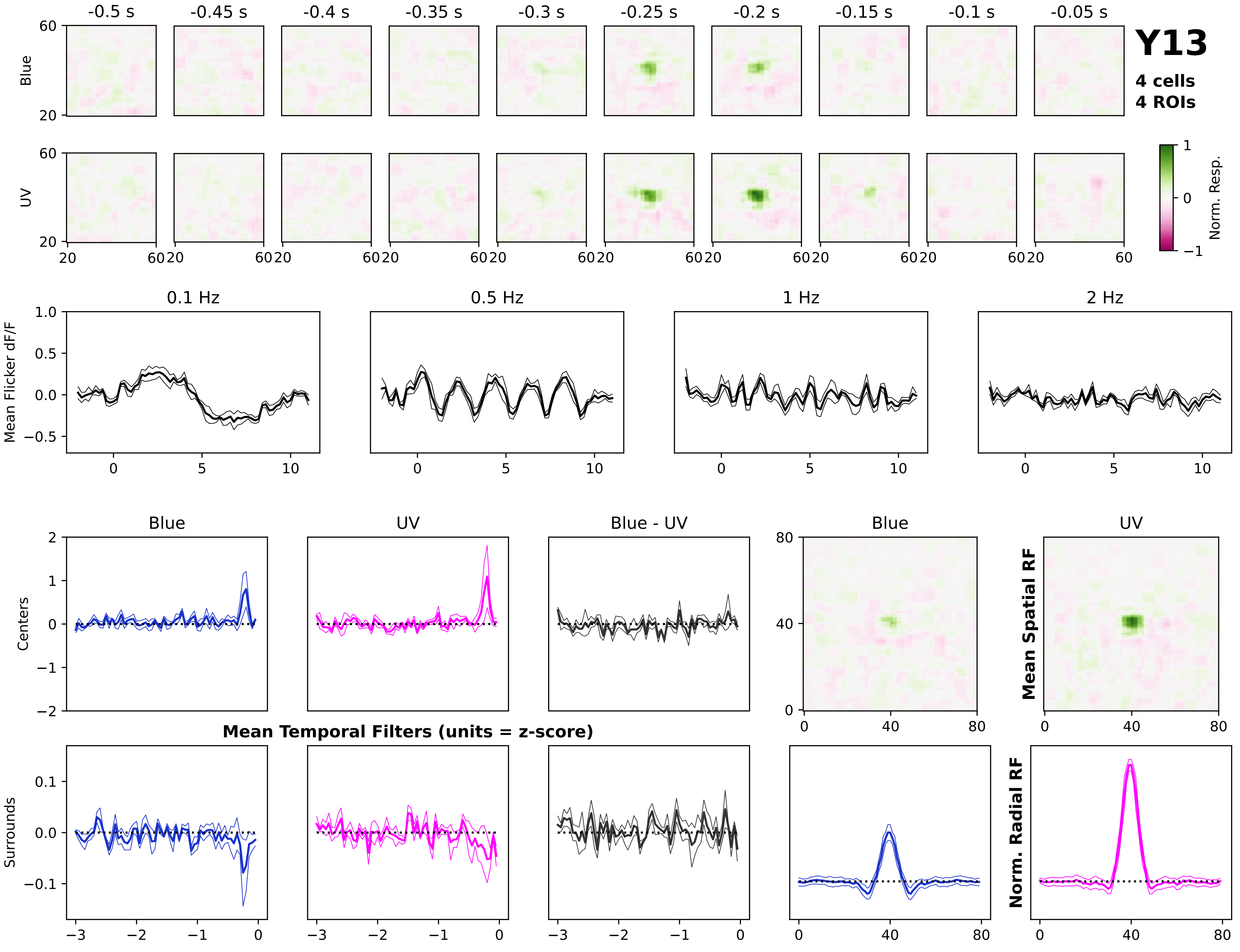

Y13
4 cells
4 ROIs
-0.5 s
-0.45 s
-0.4 s
-0.35 s
-0.3 s
-0.25 s
-0.2 s
-0.15 s
-0.1 s
-0.05 s
Blue
UV
Norm. Resp.
0.1 Hz
0.5 Hz
1 Hz
2 Hz
Mean Flicker dF/F
Blue
UV
Blue - UV
Centers
Surrounds
Mean Temporal Filters (units = z-score)
Mean Spatial RF
Norm. Radial RF
