## Supplementary material for "Infrequent strong connections constrain connectomic predictions of neuronal function": Figure S2

**Figure S2.** Related to Figure 1. *Visual response profiles for cell types not in the main dataset*. Same as Fig. S1, but for cell types sampled fewer than 3 times. These cell types are only considered in the analyses associated with Figs. 2 and 6.

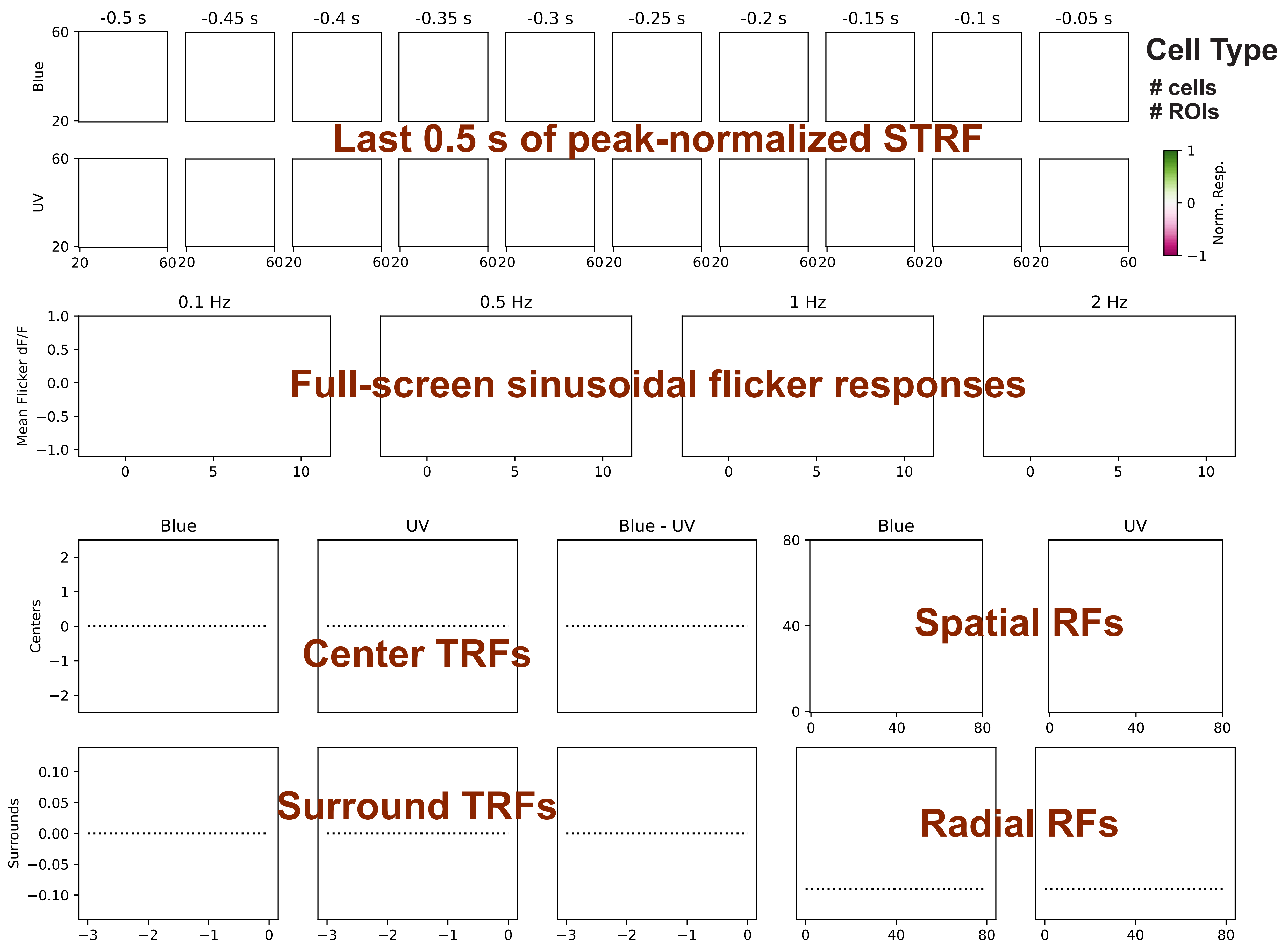
Cell Type
### cells
### ROIs
Last 0.5 s of peak-normalized STRF
-0.5 s
-0.45 s
-0.4 s
-0.35 s
-0.3 s
-0.25 s
-0.2 s
-0.15 s
-0.1 s
-0.05 s
Blue
UV
Norm. Resp.
Full-screen sinusoidal flicker responses
0.1 Hz
0.5 Hz
1 Hz
2 Hz
Mean Flicker dF/F
Center TRFs
Surround TRFs
Spatial RFs
Radial RFs
Blue
UV
Blue - UV
Centers
Surrounds

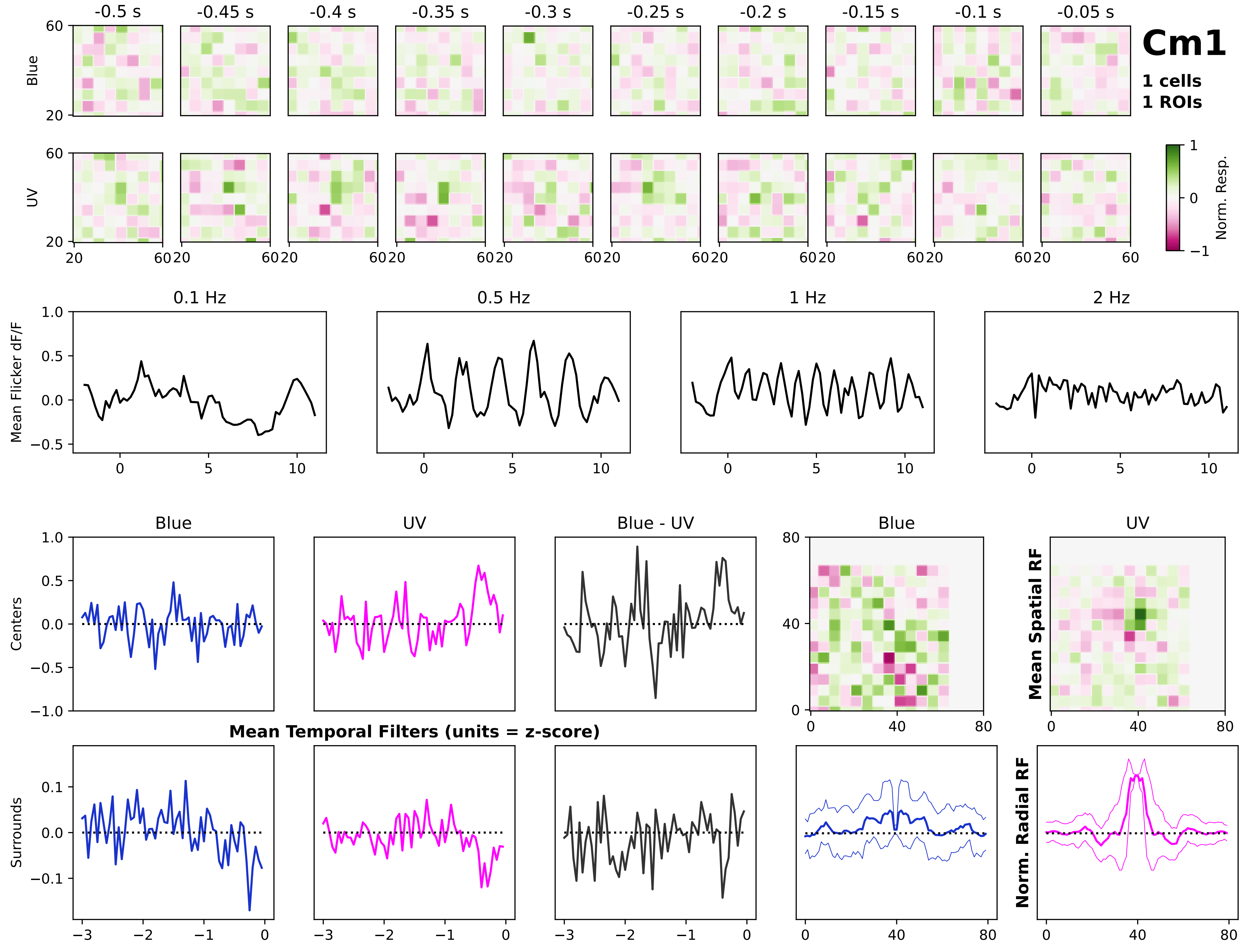
Cm1
1 cells
1 ROIs
-0.5 s
-0.45 s
-0.4 s
-0.35 s
-0.3 s
-0.25 s
-0.2 s
-0.15 s
-0.1 s
-0.05 s
Blue
UV
Norm. Resp.
0.1 Hz
0.5 Hz
1 Hz
2 Hz
Mean Flicker dF/F
Blue
UV
Blue - UV
Centers
Surrounds
Mean Temporal Filters (units = z-score)
Mean Spatial RF
Norm. Radial RF

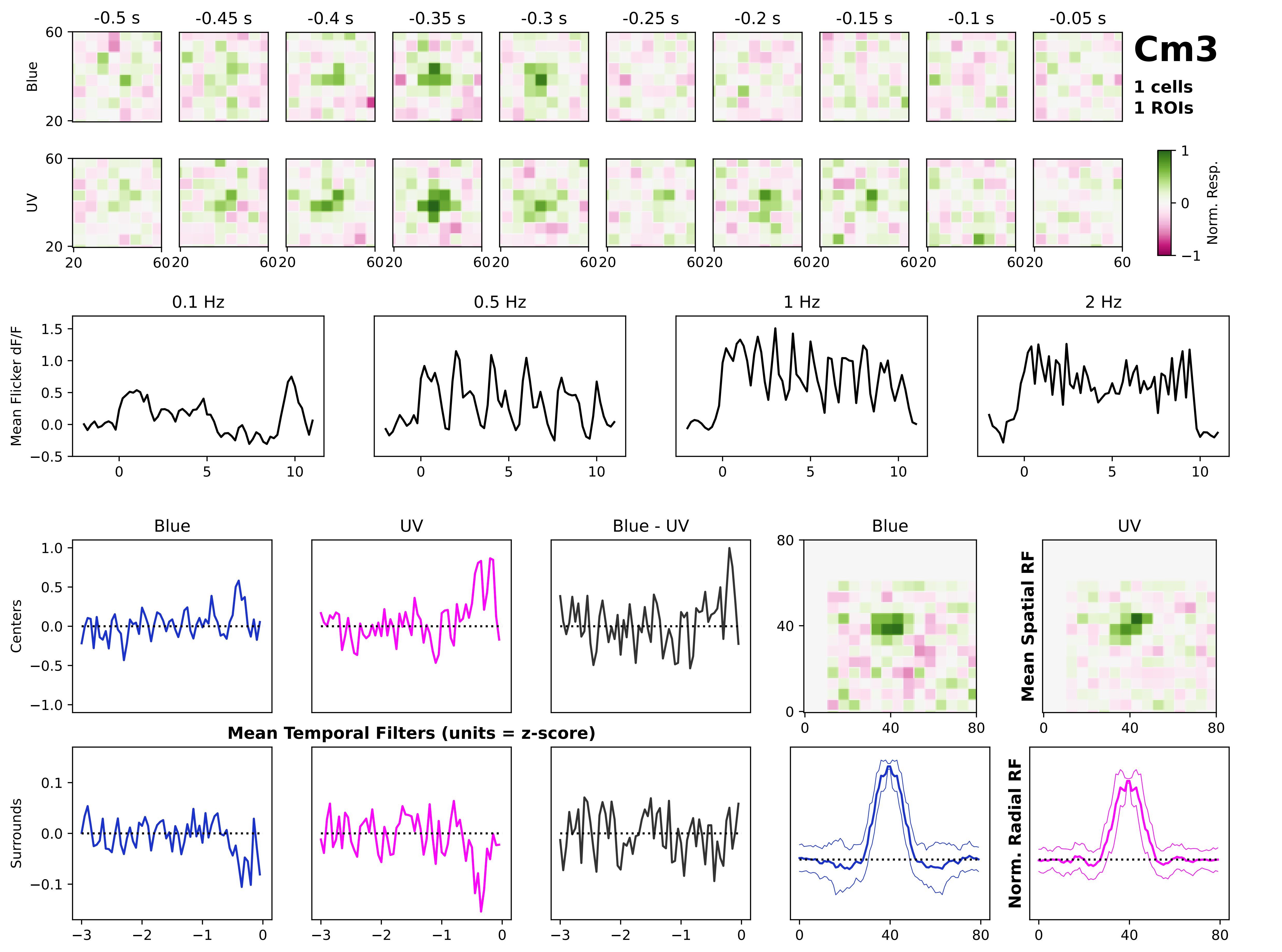
-0.5 s
-0.45 s
-0.4 s
-0.35 s
-0.3 s
-0.25 s
-0.2 s
-0.15 s
-0.1 s
-0.05 s
Cm3
1 cells
1 ROIs
Blue
UV
Norm. Resp.
0.1 Hz
0.5 Hz
1 Hz
2 Hz
Mean Flicker dF/F
Blue
UV
Blue - UV
Centers
Surrounds
Mean Temporal Filters (units = z-score)
Mean Spatial RF
Norm. Radial RF

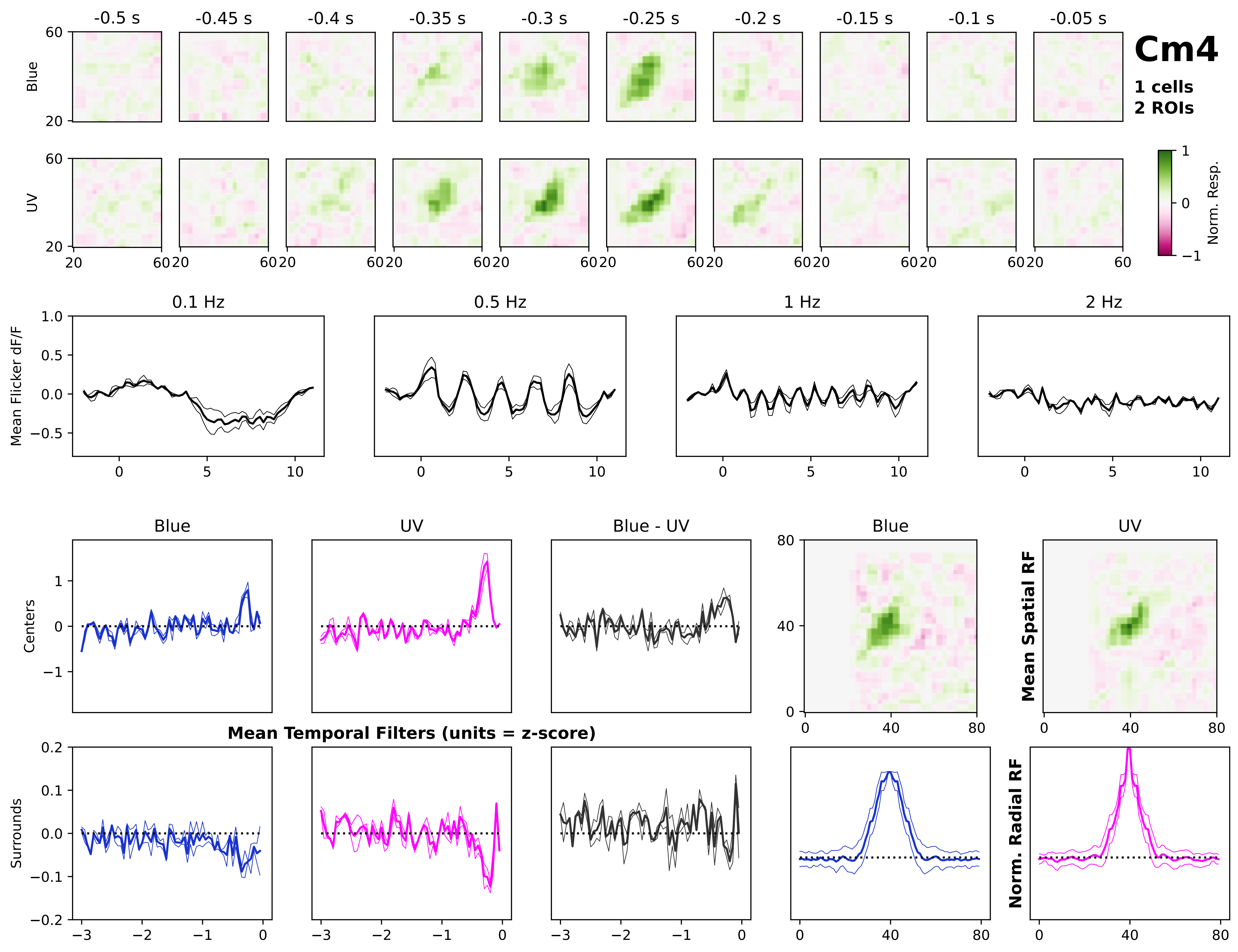
Cm4
1 cells
2 ROIs
-0.5 s
-0.45 s
-0.4 s
-0.35 s
-0.3 s
-0.25 s
-0.2 s
-0.15 s
-0.1 s
-0.05 s
Blue
UV
Norm. Resp.
0.1 Hz
0.5 Hz
1 Hz
2 Hz
Mean Flicker dF/F
Centers
Surrounds
Blue
UV
Blue - UV
Mean Temporal Filters (units = z-score)
Mean Spatial RF
Norm. Radial RF

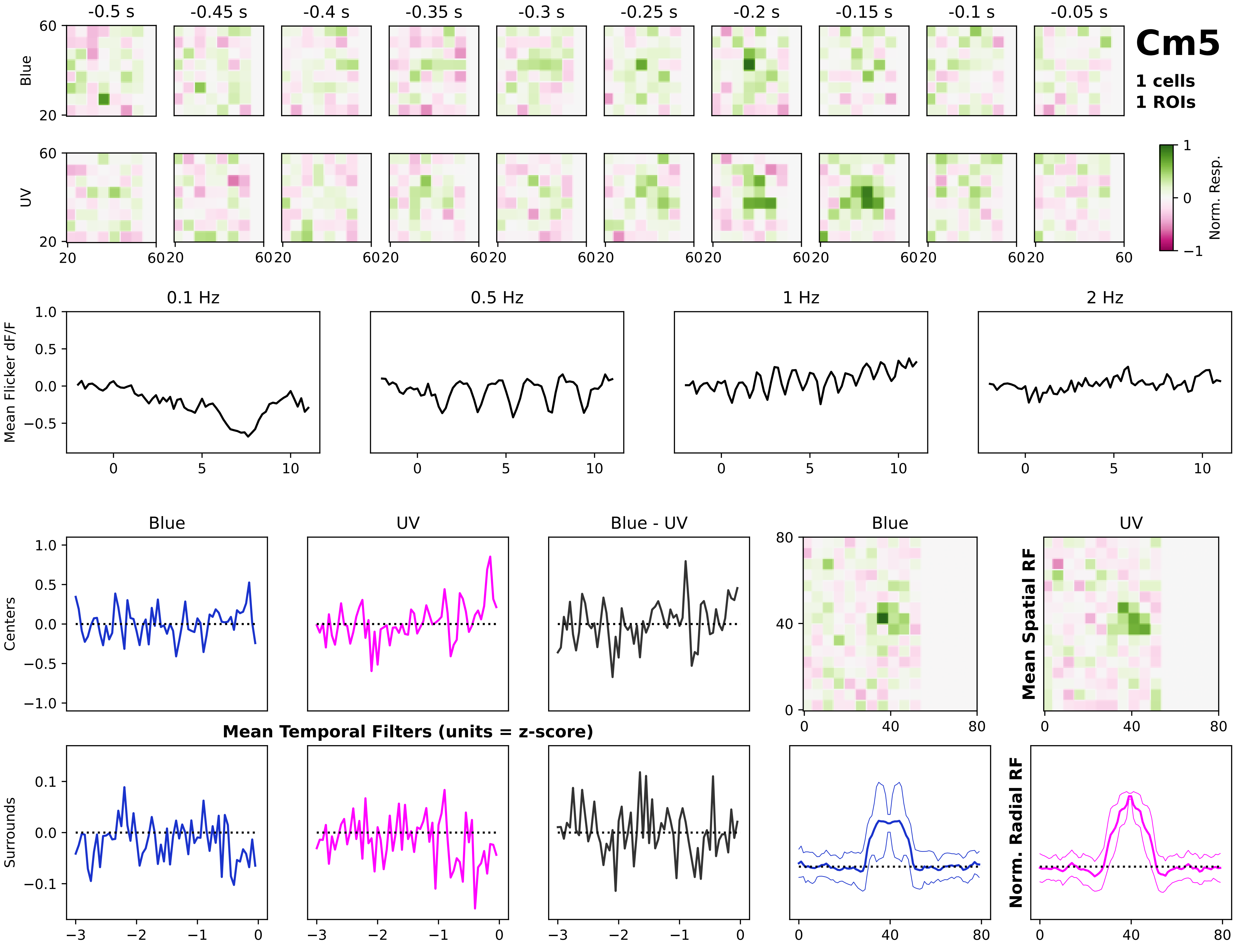
Cm5
1 cells
1 ROIs
-0.5 s
-0.45 s
-0.4 s
-0.35 s
-0.3 s
-0.25 s
-0.2 s
-0.15 s
-0.1 s
-0.05 s
Blue
UV
Norm. Resp.
0.1 Hz
0.5 Hz
1 Hz
2 Hz
Mean Flicker dF/F
Blue
UV
Blue - UV
Centers
Surrounds
Mean Temporal Filters (units = z-score)
Mean Spatial RF
Norm. Radial RF

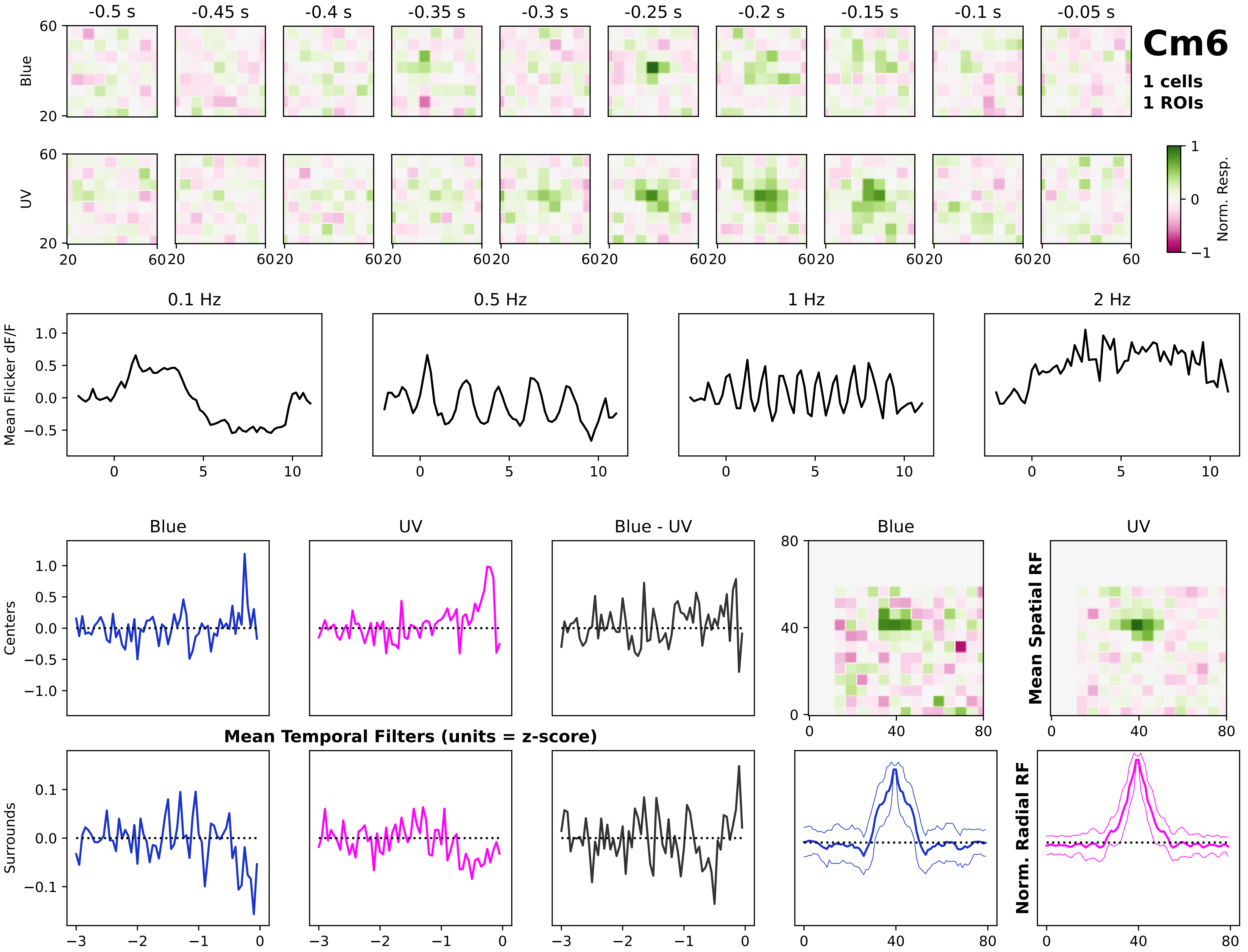
Cm6
1 cells
1 ROIs
-0.5 s
-0.45 s
-0.4 s
-0.35 s
-0.3 s
-0.25 s
-0.2 s
-0.15 s
-0.1 s
-0.05 s
Blue
UV
Norm. Resp.
0.1 Hz
0.5 Hz
1 Hz
2 Hz
Mean Flicker dF/F
Blue
UV
Blue - UV
Centers
Surrounds
Mean Temporal Filters (units = z-score)
Mean Spatial RF
Norm. Radial RF

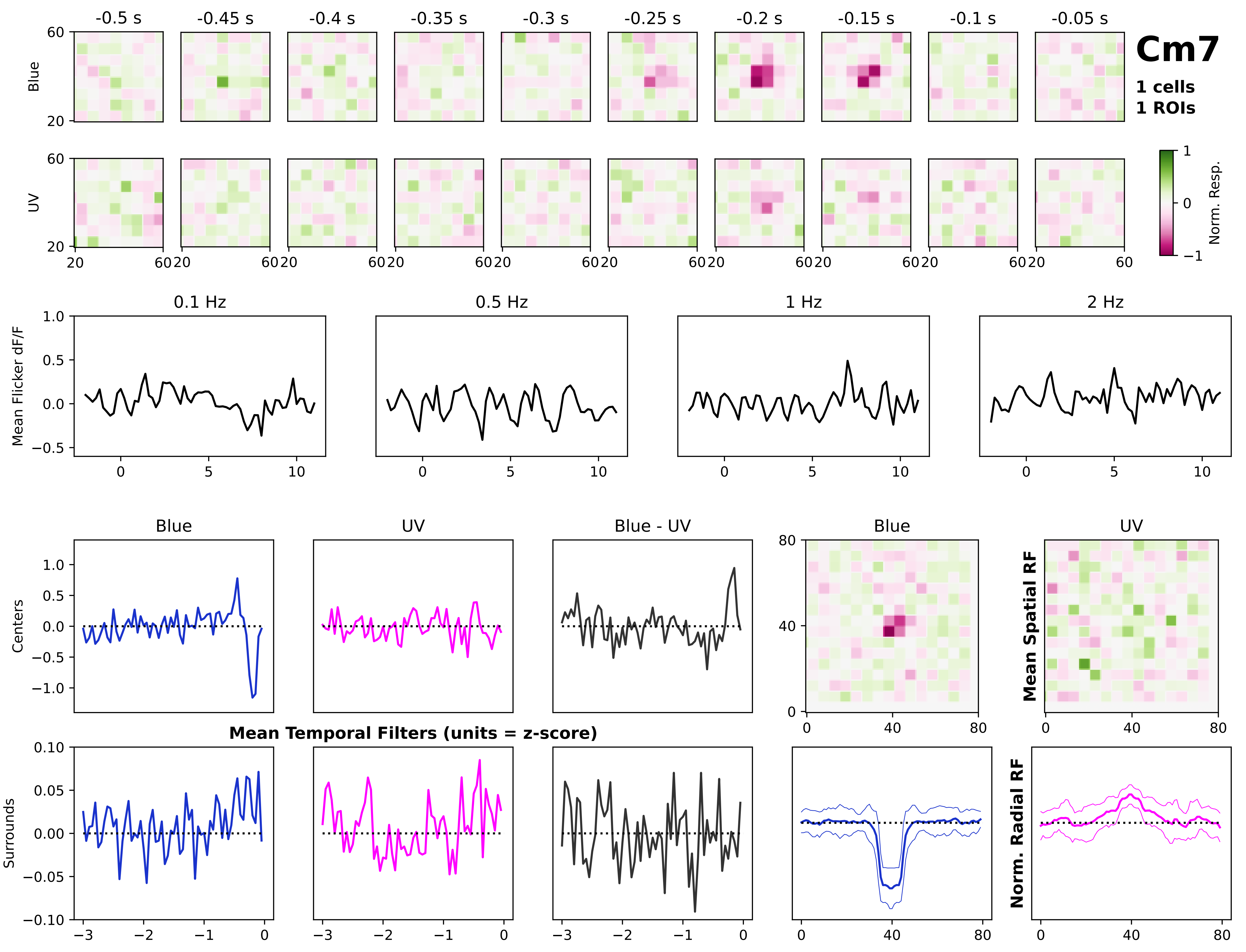
Cm7
1 cells
1 ROIs
-0.5 s
-0.45 s
-0.4 s
-0.35 s
-0.3 s
-0.25 s
-0.2 s
-0.15 s
-0.1 s
-0.05 s
Blue
UV
Norm. Resp.
0.1 Hz
0.5 Hz
1 Hz
2 Hz
Mean Flicker dF/F
Blue
UV
Blue - UV
Centers
Surrounds
Mean Temporal Filters (units = z-score)
Mean Spatial RF
Norm. Radial RF

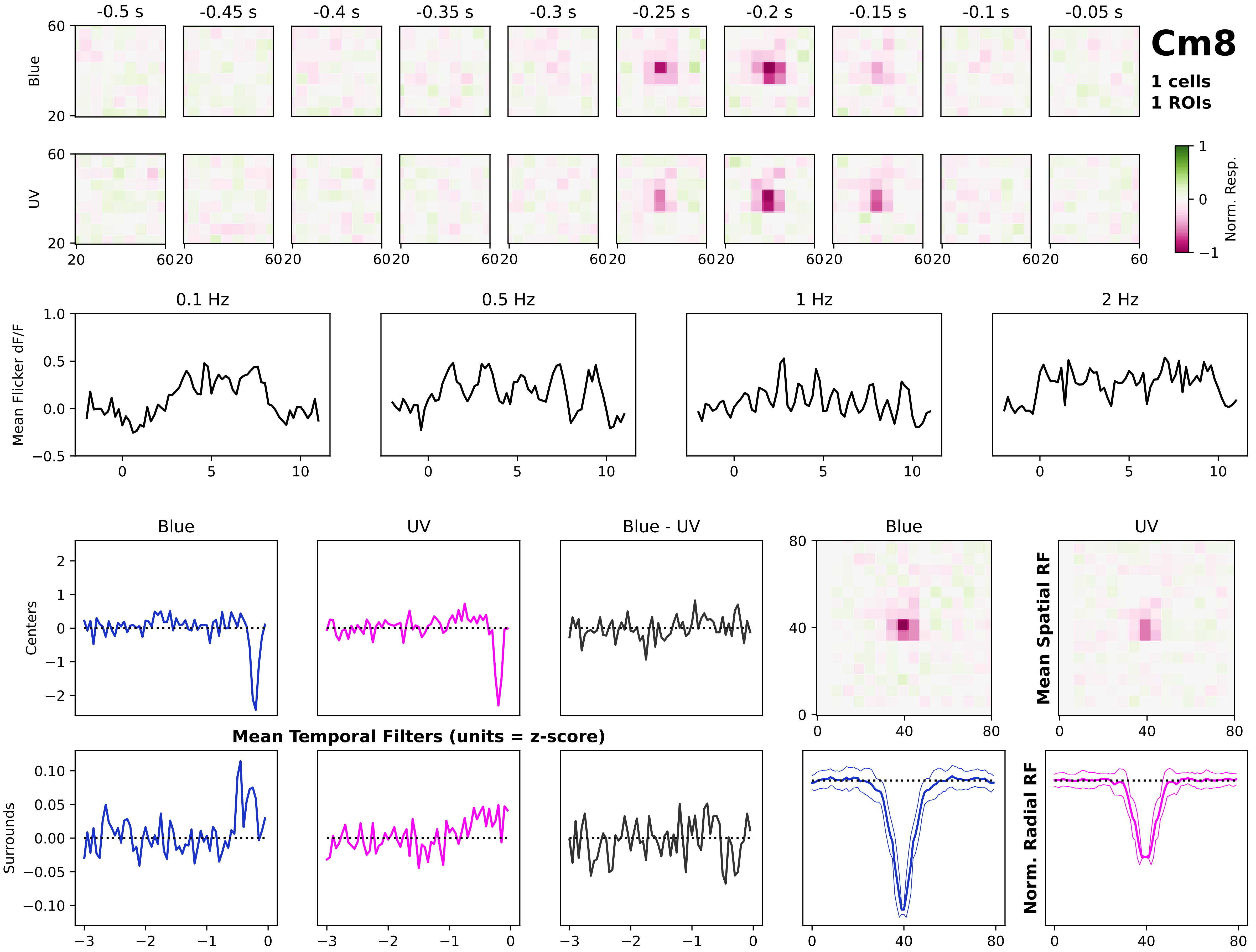
Cm8
1 cells
1 ROIs
-0.5 s
-0.45 s
-0.4 s
-0.35 s
-0.3 s
-0.25 s
-0.2 s
-0.15 s
-0.1 s
-0.05 s
Blue
UV
Norm. Resp.
0.1 Hz
0.5 Hz
1 Hz
2 Hz
Mean Flicker dF/F
Blue
UV
Blue - UV
Centers
Surrounds
Mean Temporal Filters (units = z-score)
Mean Spatial RF
Norm. Radial RF

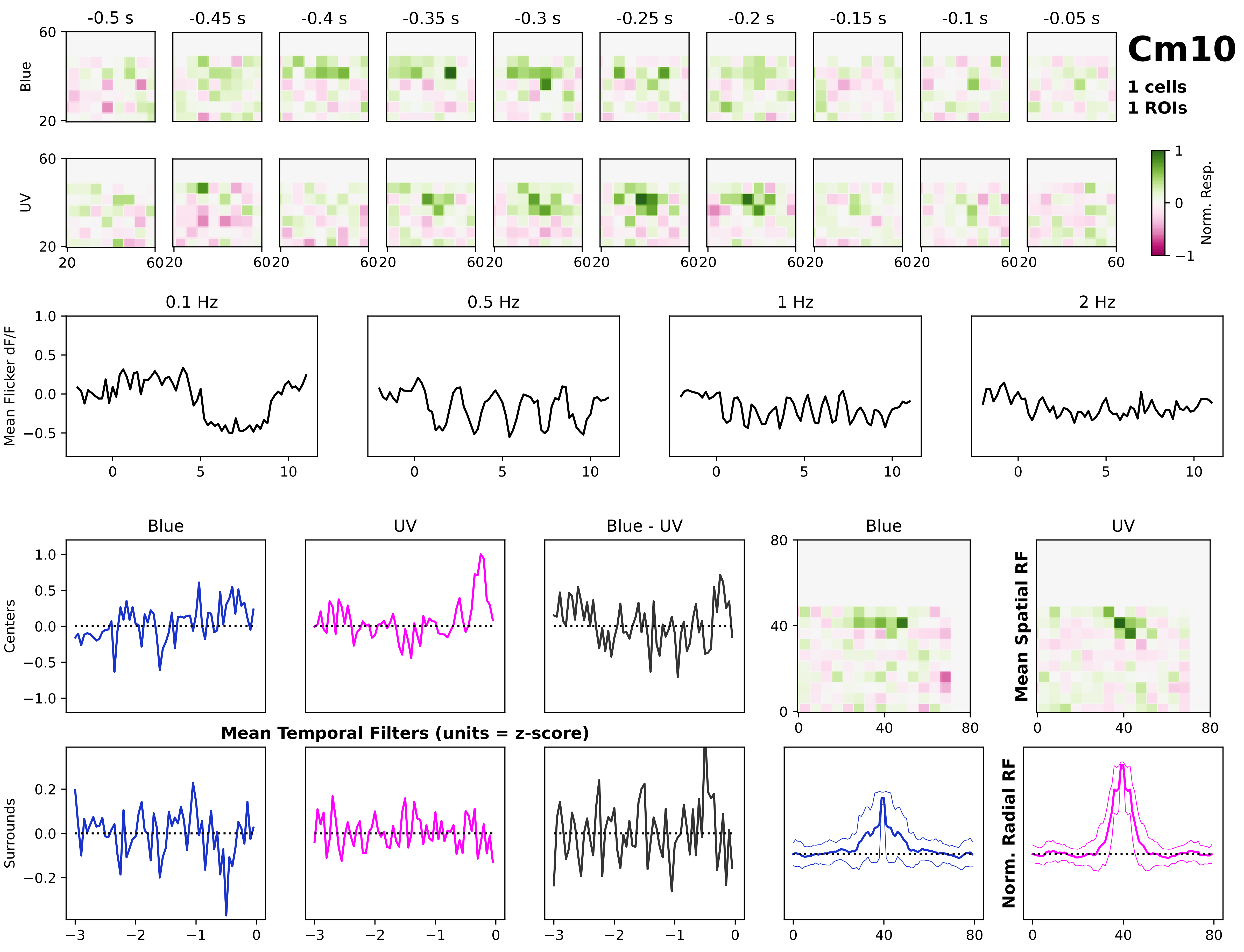
Cm10
1 cells
1 ROIs
-0.5 s
-0.45 s
-0.4 s
-0.35 s
-0.3 s
-0.25 s
-0.2 s
-0.15 s
-0.1 s
-0.05 s
Blue
UV
Norm. Resp.
0.1 Hz
0.5 Hz
1 Hz
2 Hz
Mean Flicker dF/F
Blue
UV
Blue - UV
Centers
Surrounds
Mean Temporal Filters (units = z-score)
Mean Spatial RF
Norm. Radial RF

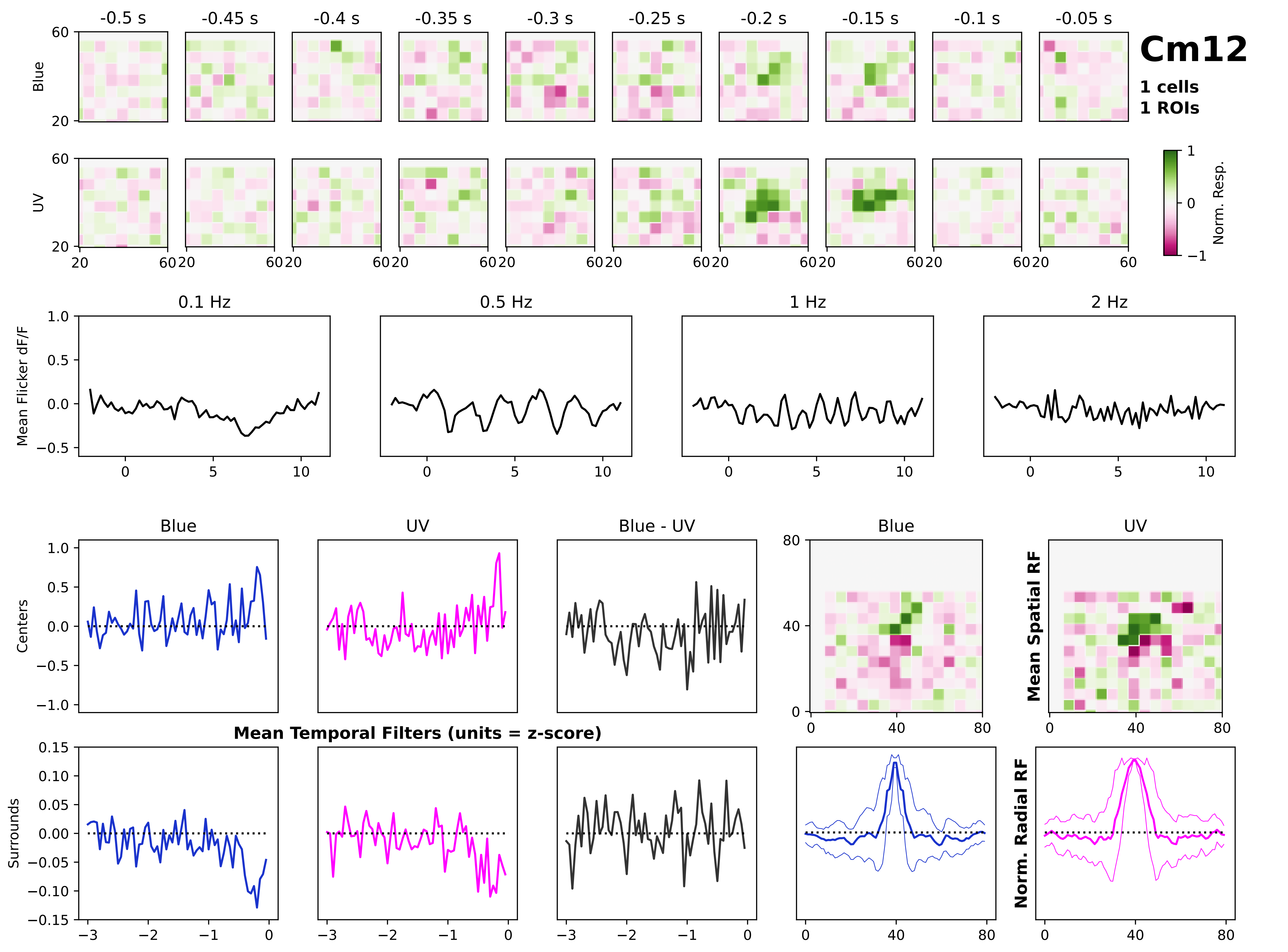
Cm12
1 cells
1 ROIs
-0.5 s
-0.45 s
-0.4 s
-0.35 s
-0.3 s
-0.25 s
-0.2 s
-0.15 s
-0.1 s
-0.05 s
Blue
UV
Norm. Resp.
0.1 Hz
0.5 Hz
1 Hz
2 Hz
Mean Flicker dF/F
Blue
UV
Blue - UV
Centers
Surrounds
Mean Temporal Filters (units = z-score)
Mean Spatial RF
Norm. Radial RF

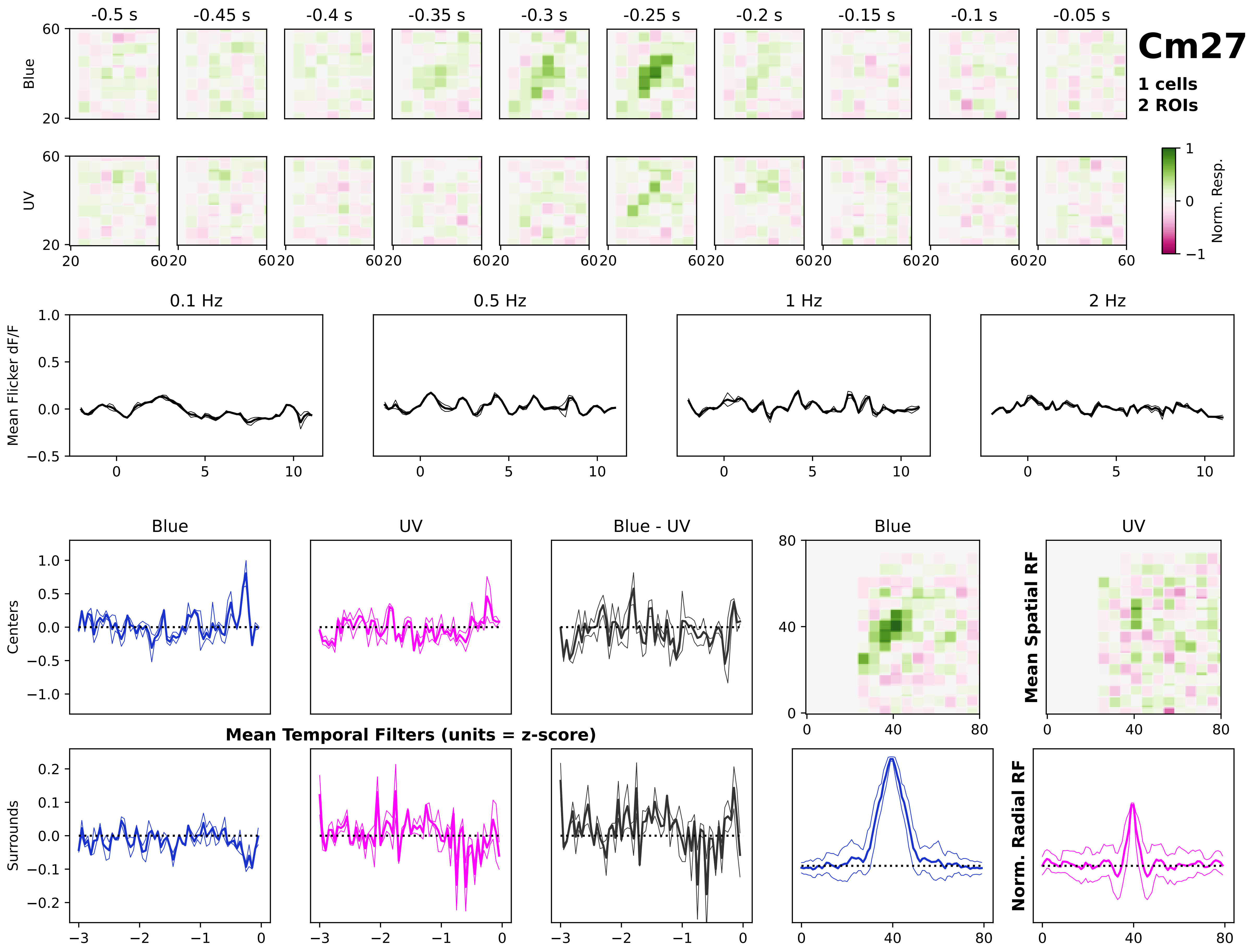
Cm27
1 cells
2 ROIs
-0.5 s
-0.45 s
-0.4 s
-0.35 s
-0.3 s
-0.25 s
-0.2 s
-0.15 s
-0.1 s
-0.05 s
Blue
UV
Norm. Resp.
0.1 Hz
0.5 Hz
1 Hz
2 Hz
Mean Flicker dF/F
Blue
UV
Blue - UV
Centers
Surrounds
Mean Temporal Filters (units = z-score)
Mean Spatial RF
Norm. Radial RF

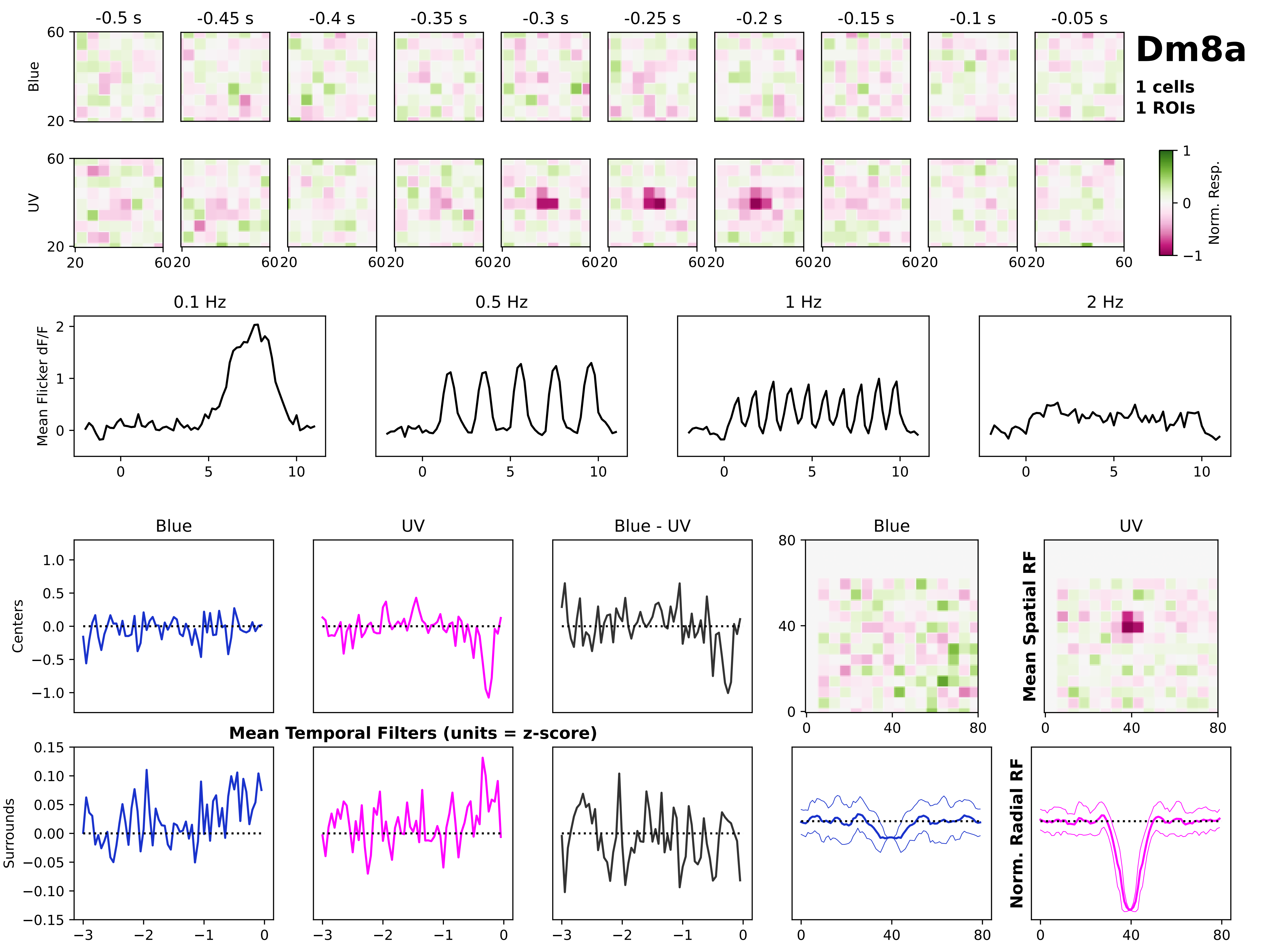

Dm8a
1 cells
1 ROIs
-0.5 s
-0.45 s
-0.4 s
-0.35 s
-0.3 s
-0.25 s
-0.2 s
-0.15 s
-0.1 s
-0.05 s
Blue
UV
Norm. Resp.
0.1 Hz
0.5 Hz
1 Hz
2 Hz
Mean Flicker dF/F
Centers
Surrounds
Blue - UV
Mean Temporal Filters (units = z-score)
Mean Spatial RF
Norm. Radial RF

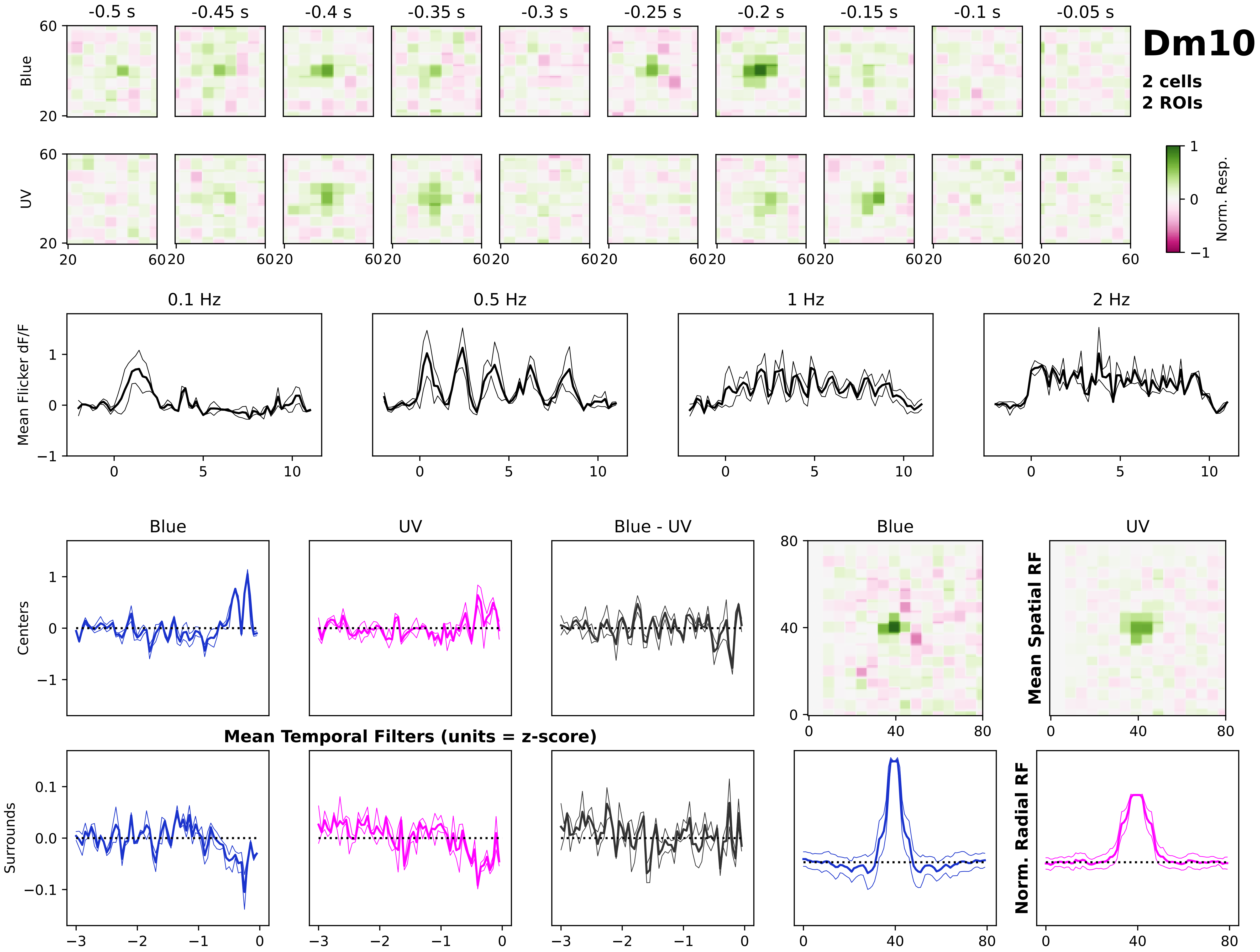
Dm10
2 cells
2 ROIs
-0.5 s
-0.45 s
-0.4 s
-0.35 s
-0.3 s
-0.25 s
-0.2 s
-0.15 s
-0.1 s
-0.05 s
Blue
UV
Norm. Resp.
0.1 Hz
0.5 Hz
1 Hz
2 Hz
Mean Flicker dF/F
Blue
UV
Blue - UV
Centers
Surrounds
Mean Temporal Filters (units = z-score)
Mean Spatial RF
Norm. Radial RF

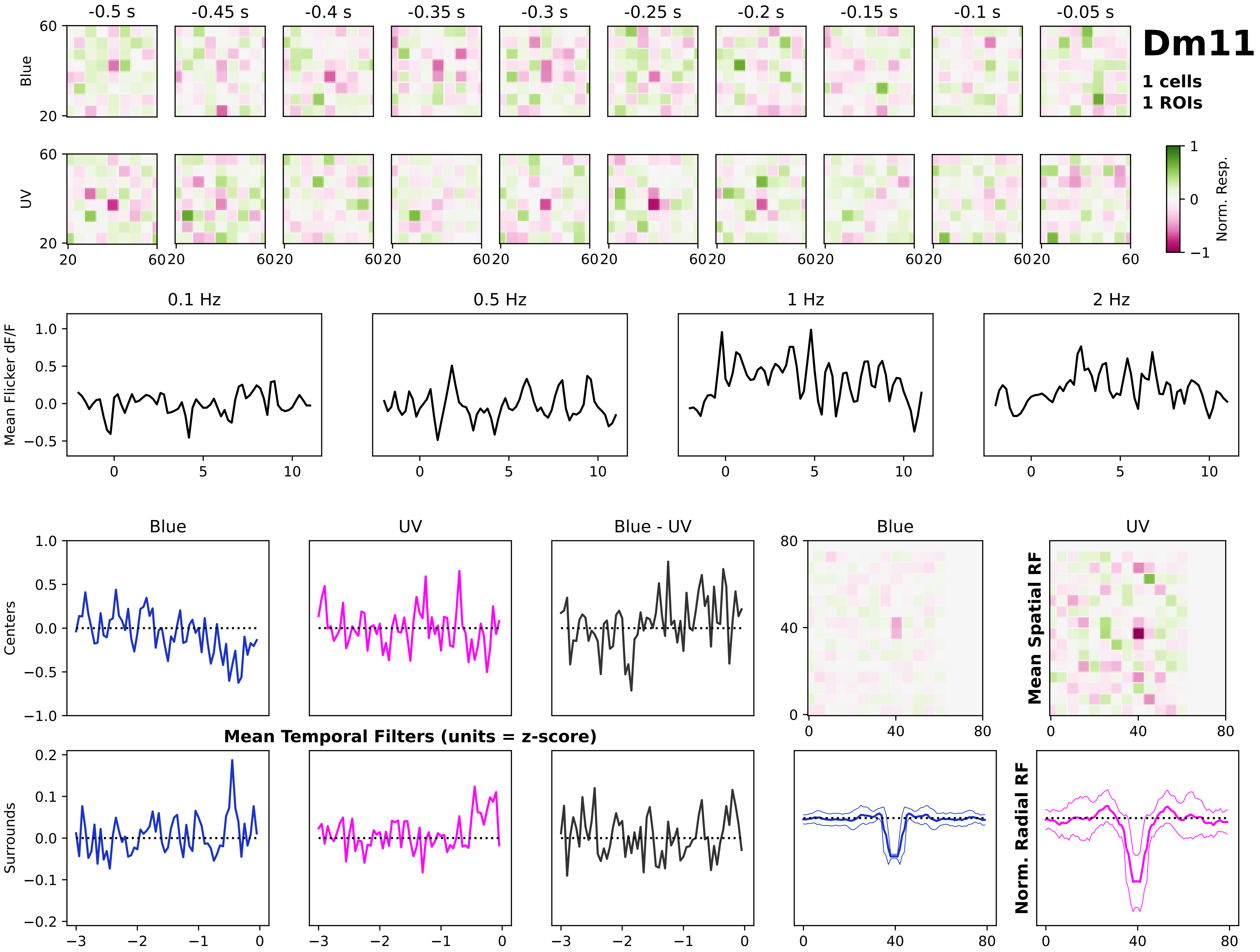

Dm11
1 cells
1 ROIs
-0.5 s
-0.45 s
-0.4 s
-0.35 s
-0.3 s
-0.25 s
-0.2 s
-0.15 s
-0.1 s
-0.05 s
Blue
UV
Norm. Resp.
0.1 Hz
0.5 Hz
1 Hz
2 Hz
Mean Flicker dF/F
Blue
UV
Blue - UV
Centers
Surrounds
Mean Temporal Filters (units = z-score)
Mean Spatial RF
Norm. Radial RF

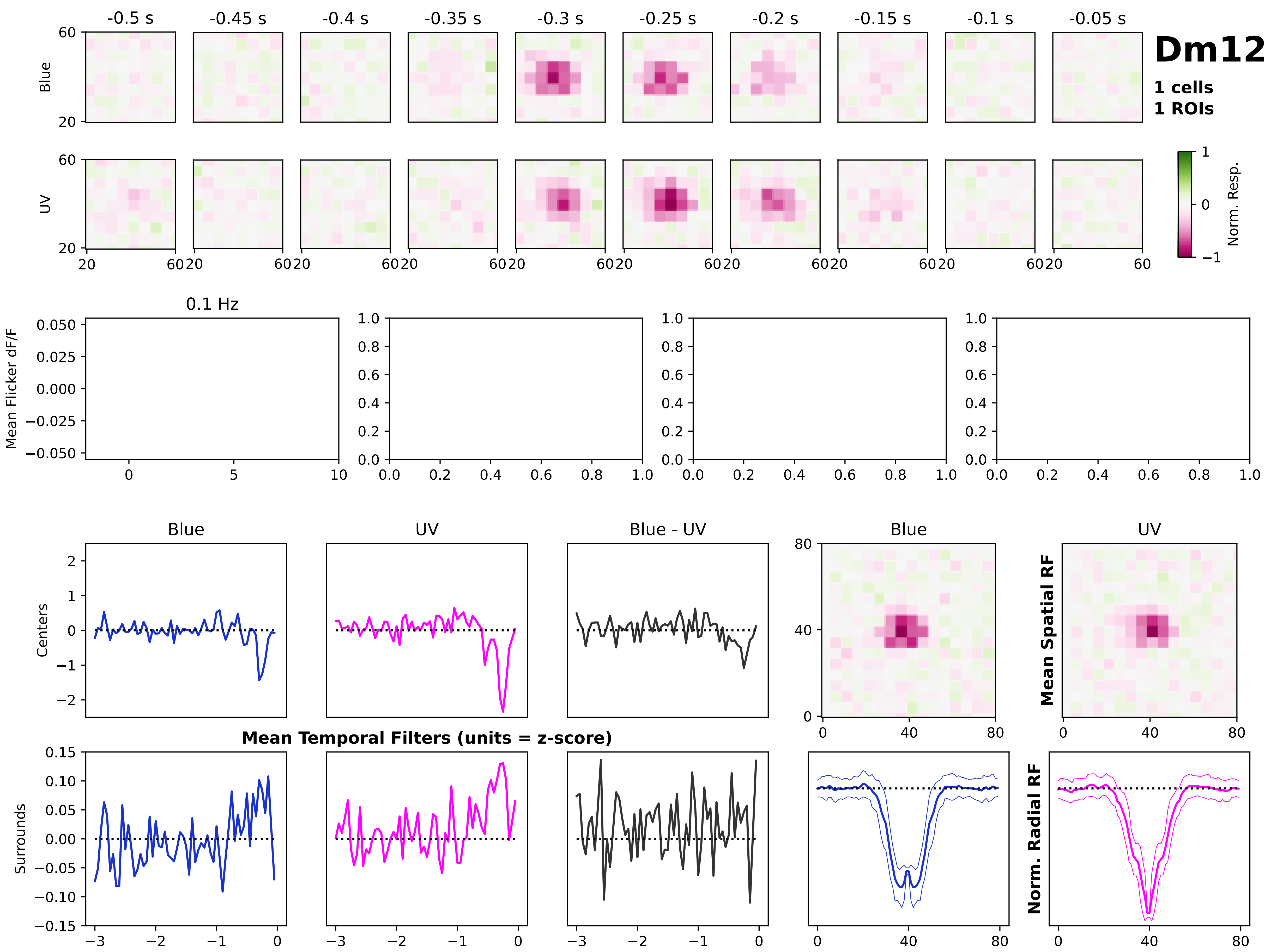

Dm12
1 cells
1 ROIs
-0.5 s
-0.45 s
-0.4 s
-0.35 s
-0.3 s
-0.25 s
-0.2 s
-0.15 s
-0.1 s
-0.05 s
Blue
UV
Norm. Resp.
0.1 Hz
Mean Flicker dF/F
Blue
UV
Blue - UV
Centers
Surrounds
Mean Temporal Filters (units = z-score)
Mean Spatial RF
Norm. Radial RF

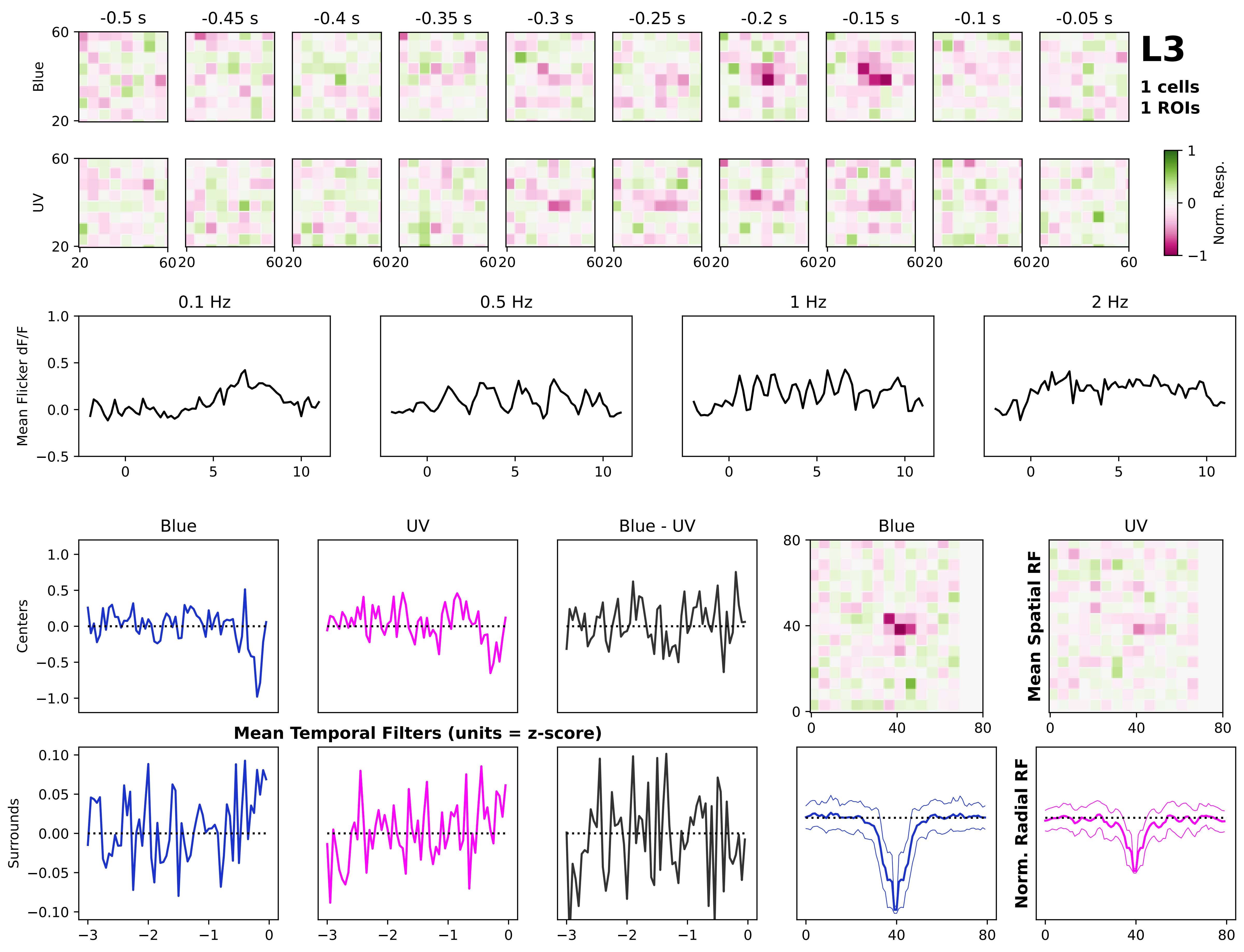
L3
1 cells
1 ROIs
-0.5 s
-0.45 s
-0.4 s
-0.35 s
-0.3 s
-0.25 s
-0.2 s
-0.15 s
-0.1 s
-0.05 s
Blue
UV
Norm. Resp.
0.1 Hz
0.5 Hz
1 Hz
2 Hz
Mean Flicker dF/F
Blue
UV
Blue - UV
Centers
Surrounds
Mean Temporal Filters (units = z-score)
Mean Spatial RF
Norm. Radial RF

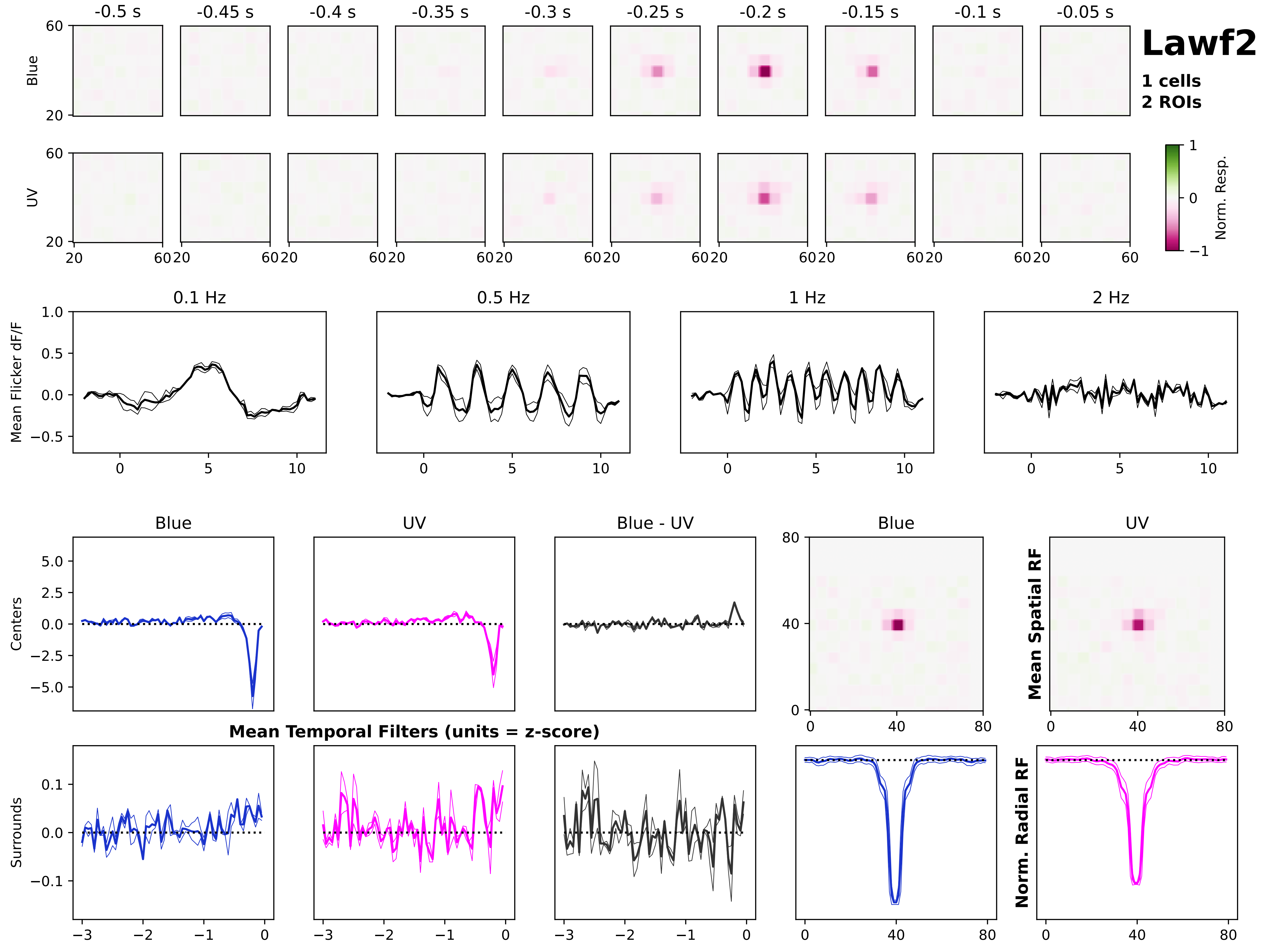

Lawf2
1 cells
2 ROIs
-0.5 s
-0.45 s
-0.4 s
-0.35 s
-0.3 s
-0.25 s
-0.2 s
-0.15 s
-0.1 s
-0.05 s
Blue
UV
Norm. Resp.
0.1 Hz
0.5 Hz
1 Hz
2 Hz
Mean Flicker dF/F
Blue
UV
Blue - UV
Centers
Surrounds
Mean Temporal Filters (units = z-score)
Mean Spatial RF
Norm. Radial RF

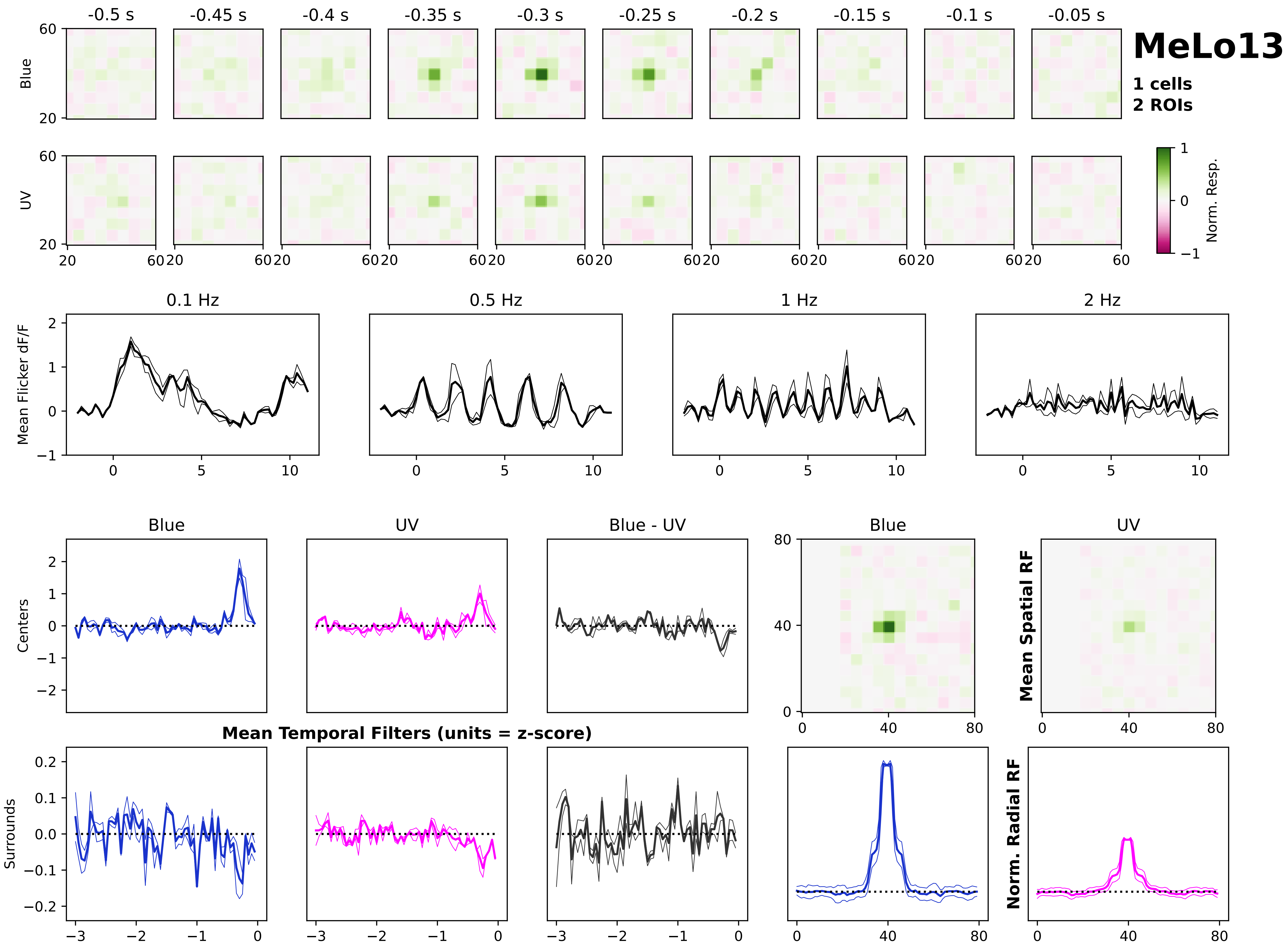
MeLo13
1 cells
2 ROIs
-0.5 s
-0.45 s
-0.4 s
-0.35 s
-0.3 s
-0.25 s
-0.2 s
-0.15 s
-0.1 s
-0.05 s
Blue
UV
Norm. Resp.
0.1 Hz
0.5 Hz
1 Hz
2 Hz
Mean Flicker dF/F
Blue
UV
Blue - UV
Centers
Surrounds
Mean Temporal Filters (units = z-score)
Mean Spatial RF
Norm. Radial RF

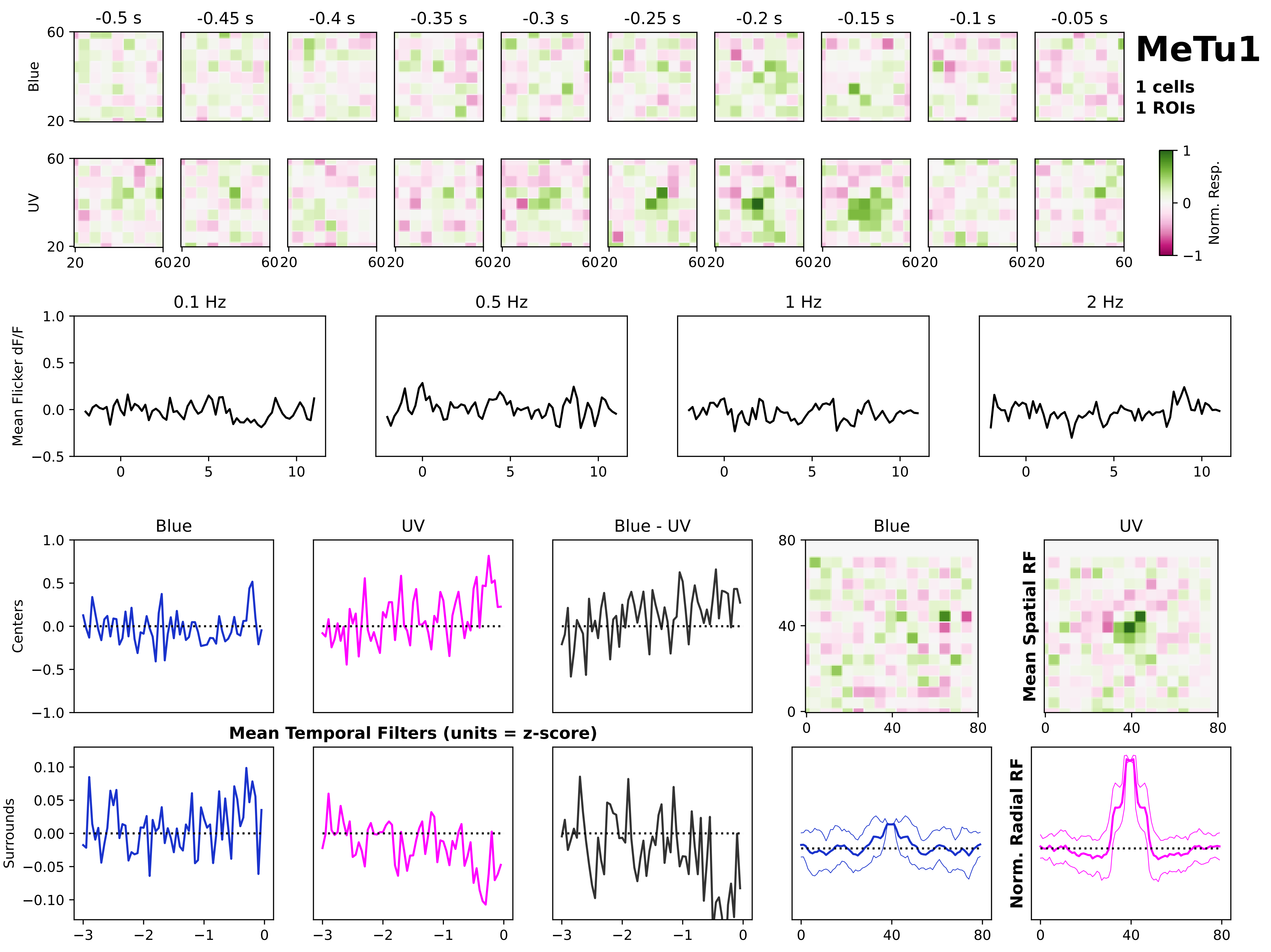
MeTu1
1 cells
1 ROIs
-0.5 s
-0.45 s
-0.4 s
-0.35 s
-0.3 s
-0.25 s
-0.2 s
-0.15 s
-0.1 s
-0.05 s
Blue
UV
Norm. Resp.
0.1 Hz
0.5 Hz
1 Hz
2 Hz
Mean Flicker dF/F
Blue
UV
Blue - UV
Centers
Surrounds
Mean Temporal Filters (units = z-score)
Mean Spatial RF
Norm. Radial RF

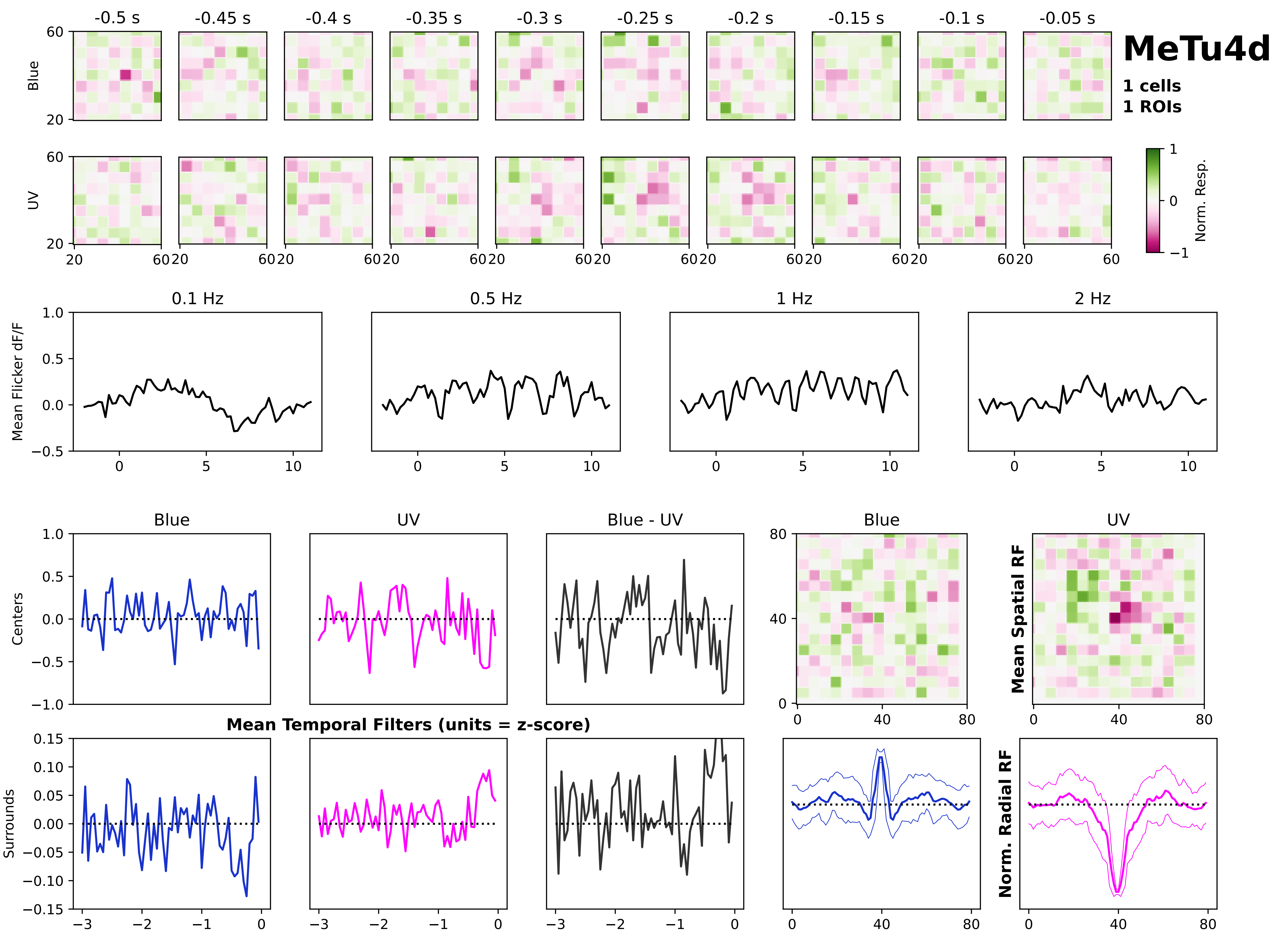
MeTu4d
1 cells
1 ROIs
-0.5 s
-0.45 s
-0.4 s
-0.35 s
-0.3 s
-0.25 s
-0.2 s
-0.15 s
-0.1 s
-0.05 s
Blue
UV
Norm. Resp.
0.1 Hz
0.5 Hz
1 Hz
2 Hz
Mean Flicker dF/F
Mean Temporal Filters (units = z-score)
Centers
Surrounds
Blue - UV
Mean Spatial RF
Norm. Radial RF

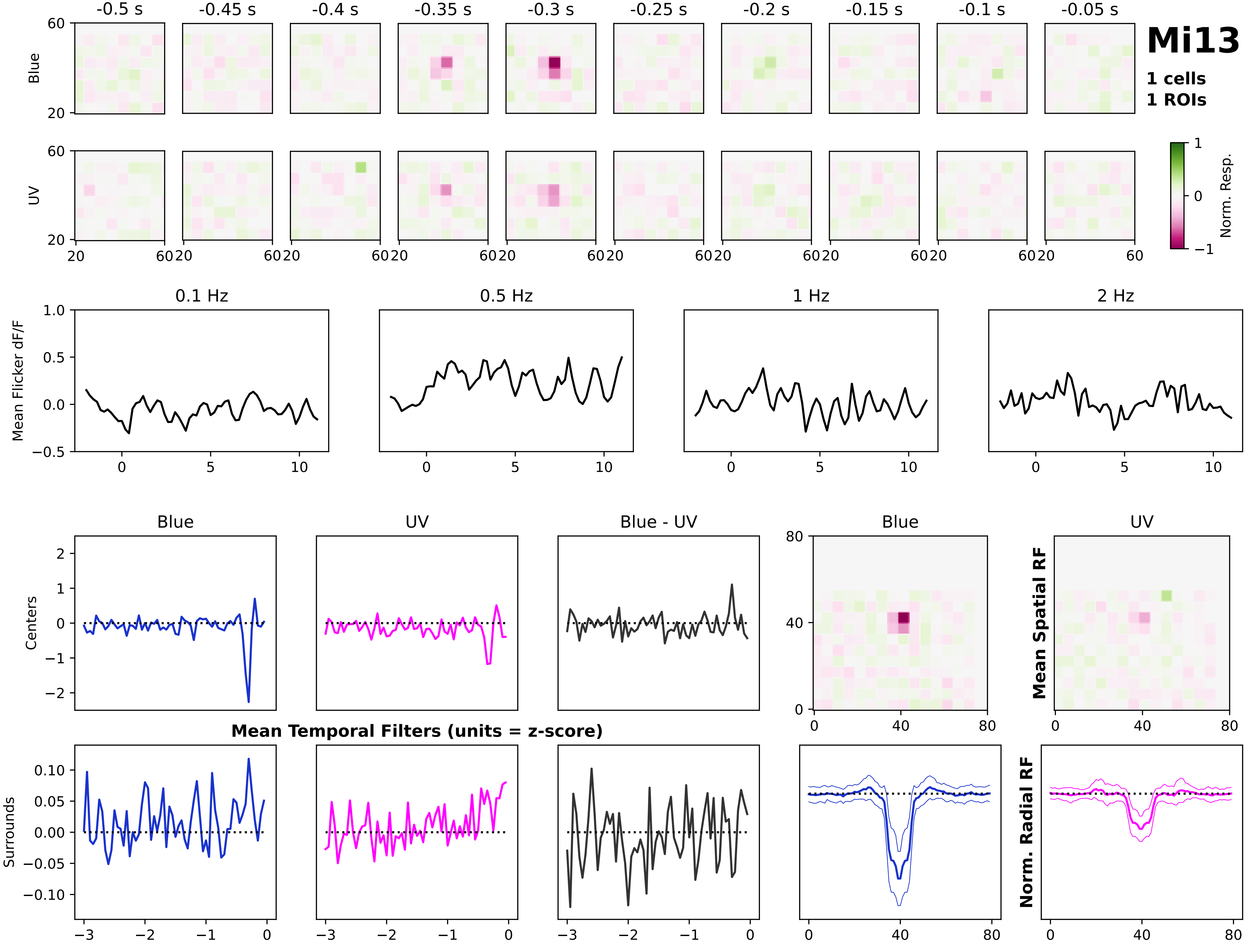

Mi13
1 cells
1 ROIs
-0.5 s
-0.45 s
-0.4 s
-0.35 s
-0.3 s
-0.25 s
-0.2 s
-0.15 s
-0.1 s
-0.05 s
Blue
UV
Norm. Resp.
0.1 Hz
0.5 Hz
1 Hz
2 Hz
Mean Flicker dF/F
Blue
UV
Blue - UV
Centers
Surrounds
Mean Temporal Filters (units = z-score)
Mean Spatial RF
Norm. Radial RF

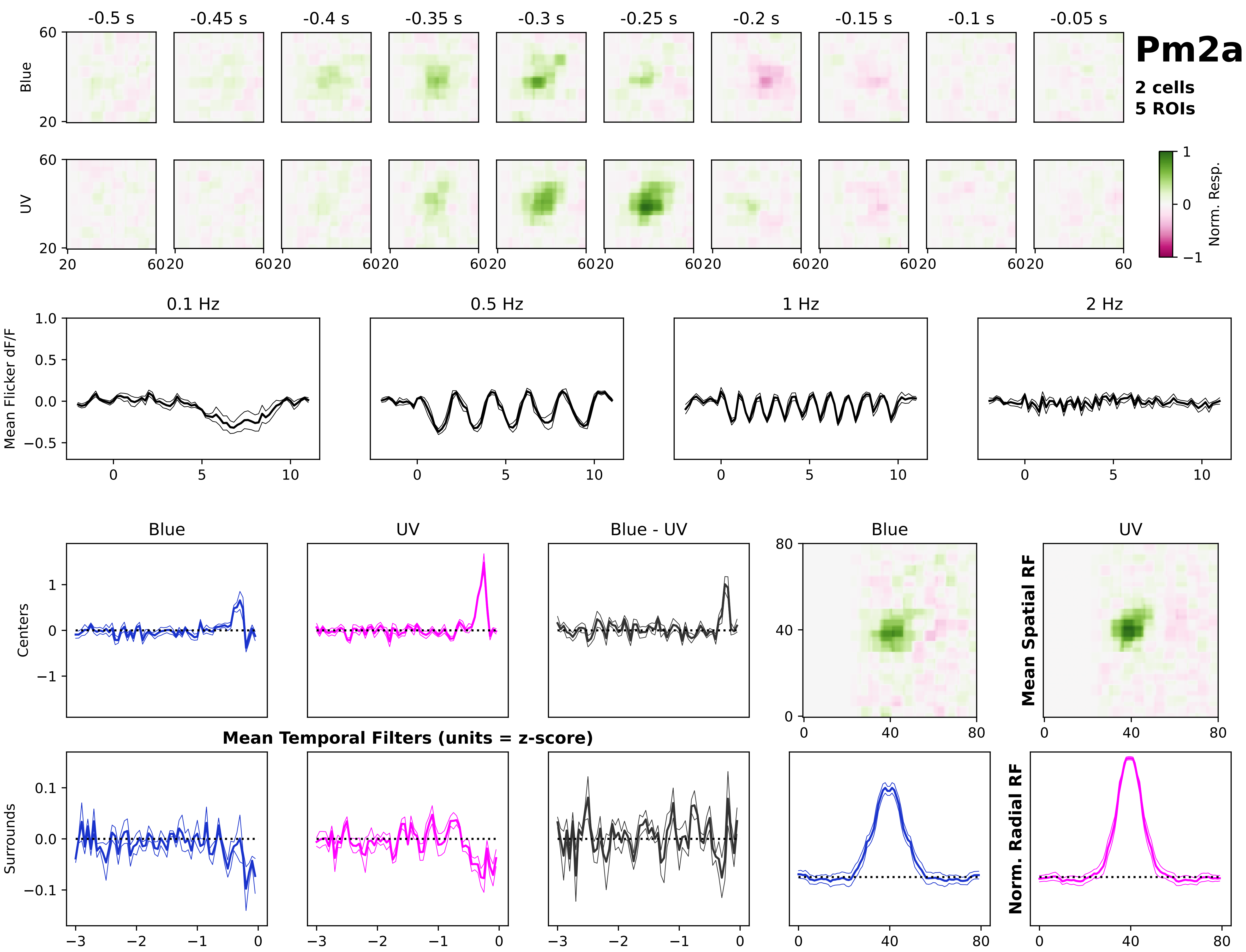

Pm2a
2 cells
5 ROIs
-0.5 s
-0.45 s
-0.4 s
-0.35 s
-0.3 s
-0.25 s
-0.2 s
-0.15 s
-0.1 s
-0.05 s
Blue
UV
Norm. Resp.
0.1 Hz
0.5 Hz
1 Hz
2 Hz
Mean Flicker dF/F
Blue
UV
Blue - UV
Centers
Surrounds
Mean Temporal Filters (units = z-score)
Mean Spatial RF
Norm. Radial RF

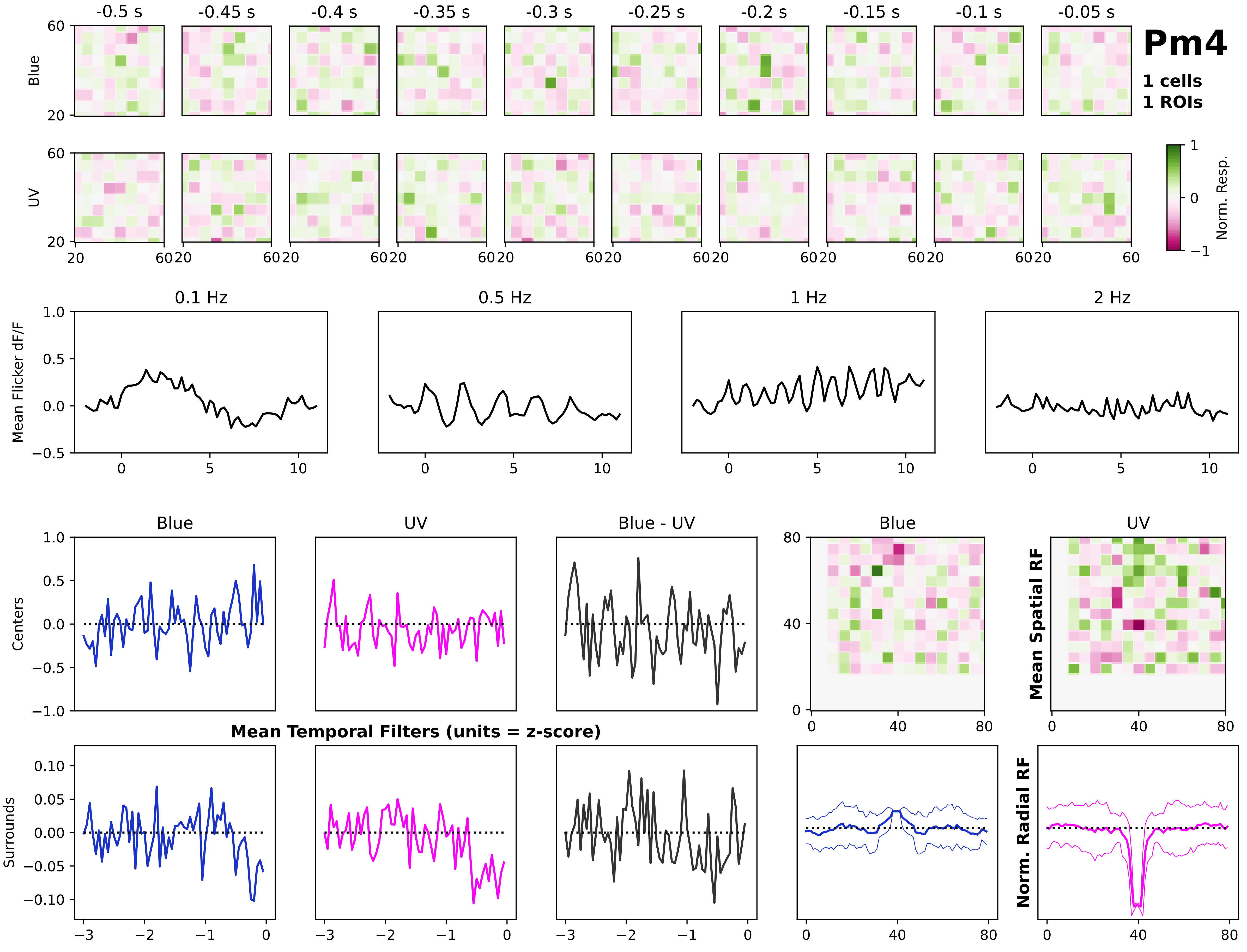

Pm4
1 cells
1 ROIs
-0.5 s
-0.45 s
-0.4 s
-0.35 s
-0.3 s
-0.25 s
-0.2 s
-0.15 s
-0.1 s
-0.05 s
Blue
UV
Norm. Resp.
0.1 Hz
0.5 Hz
1 Hz
2 Hz
Mean Flicker dF/F
Blue
UV
Blue - UV
Centers
Surrounds
Mean Temporal Filters (units = z-score)
Mean Spatial RF
Norm. Radial RF

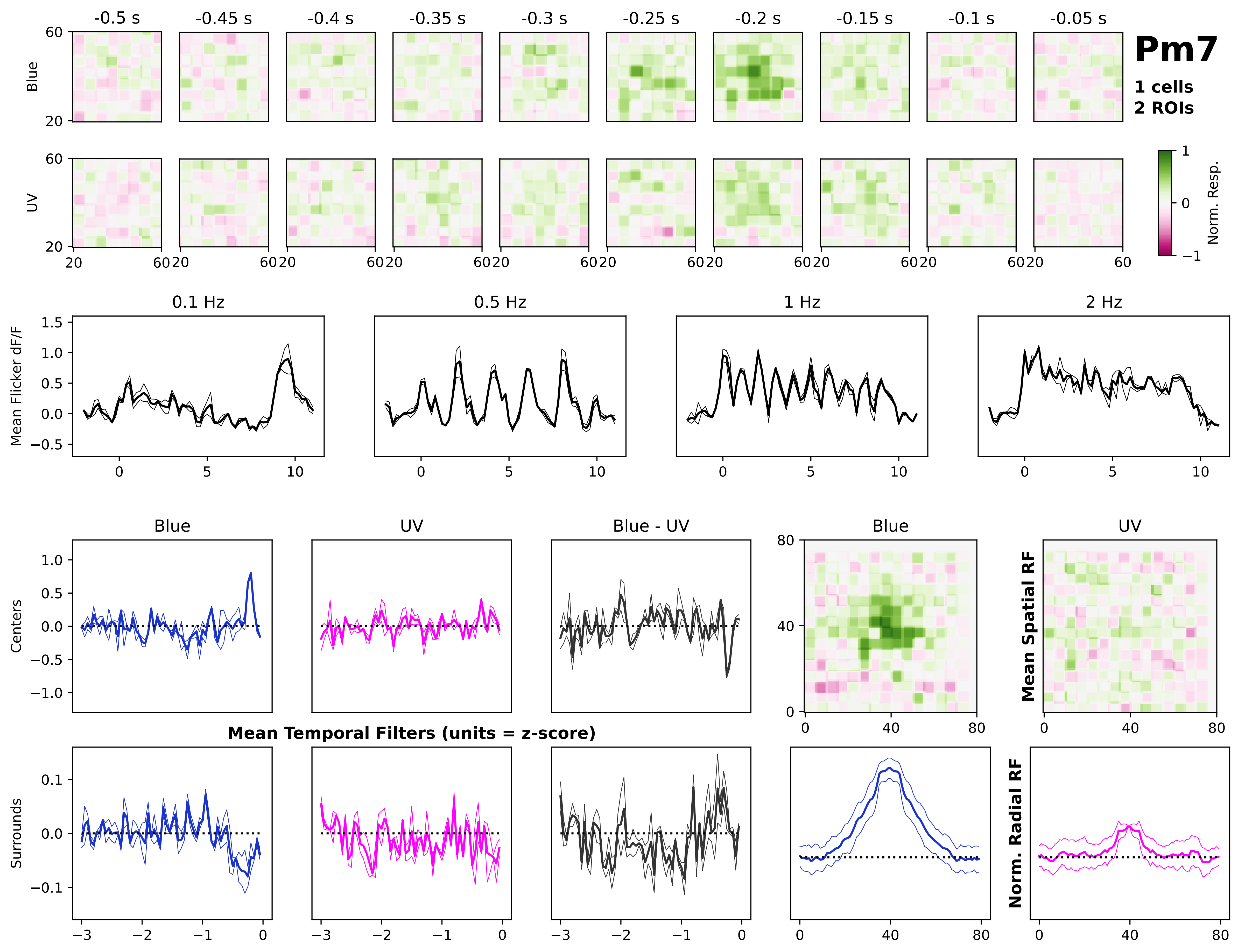
Pm7
1 cells
2 ROIs
-0.5 s
-0.45 s
-0.4 s
-0.35 s
-0.3 s
-0.25 s
-0.2 s
-0.15 s
-0.1 s
-0.05 s
Blue
UV
Norm. Resp.
0.1 Hz
0.5 Hz
1 Hz
2 Hz
Mean Flicker dF/F
Blue
UV
Blue - UV
Centers
Surrounds
Mean Temporal Filters (units = z-score)
Mean Spatial RF
Norm. Radial RF

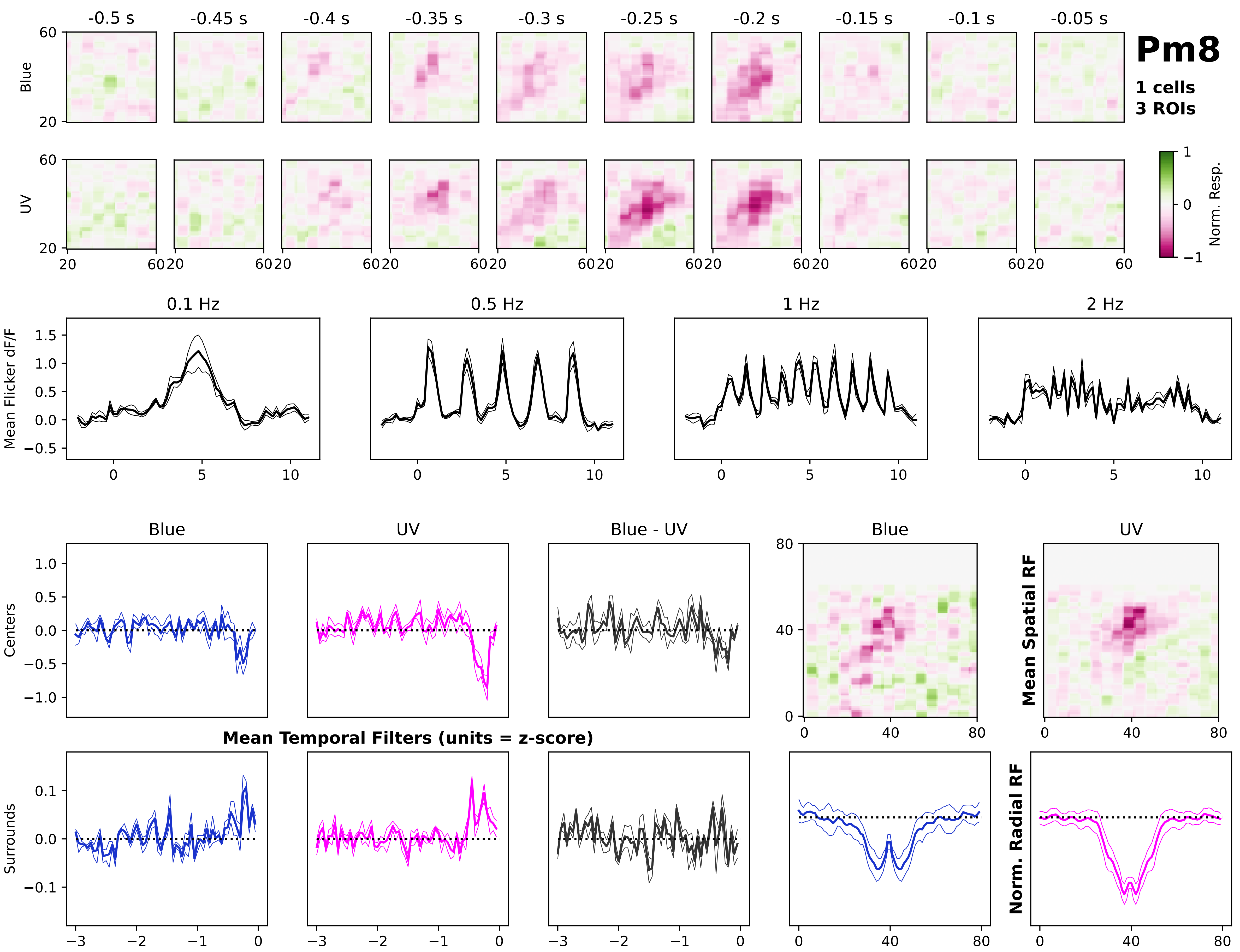
Pm8
1 cells
3 ROIs
-0.5 s
-0.45 s
-0.4 s
-0.35 s
-0.3 s
-0.25 s
-0.2 s
-0.15 s
-0.1 s
-0.05 s
Blue
UV
Norm. Resp.
0.1 Hz
0.5 Hz
1 Hz
2 Hz
Mean Flicker dF/F
Blue
UV
Blue - UV
Centers
Surrounds
Mean Temporal Filters (units = z-score)
Mean Spatial RF
Norm. Radial RF

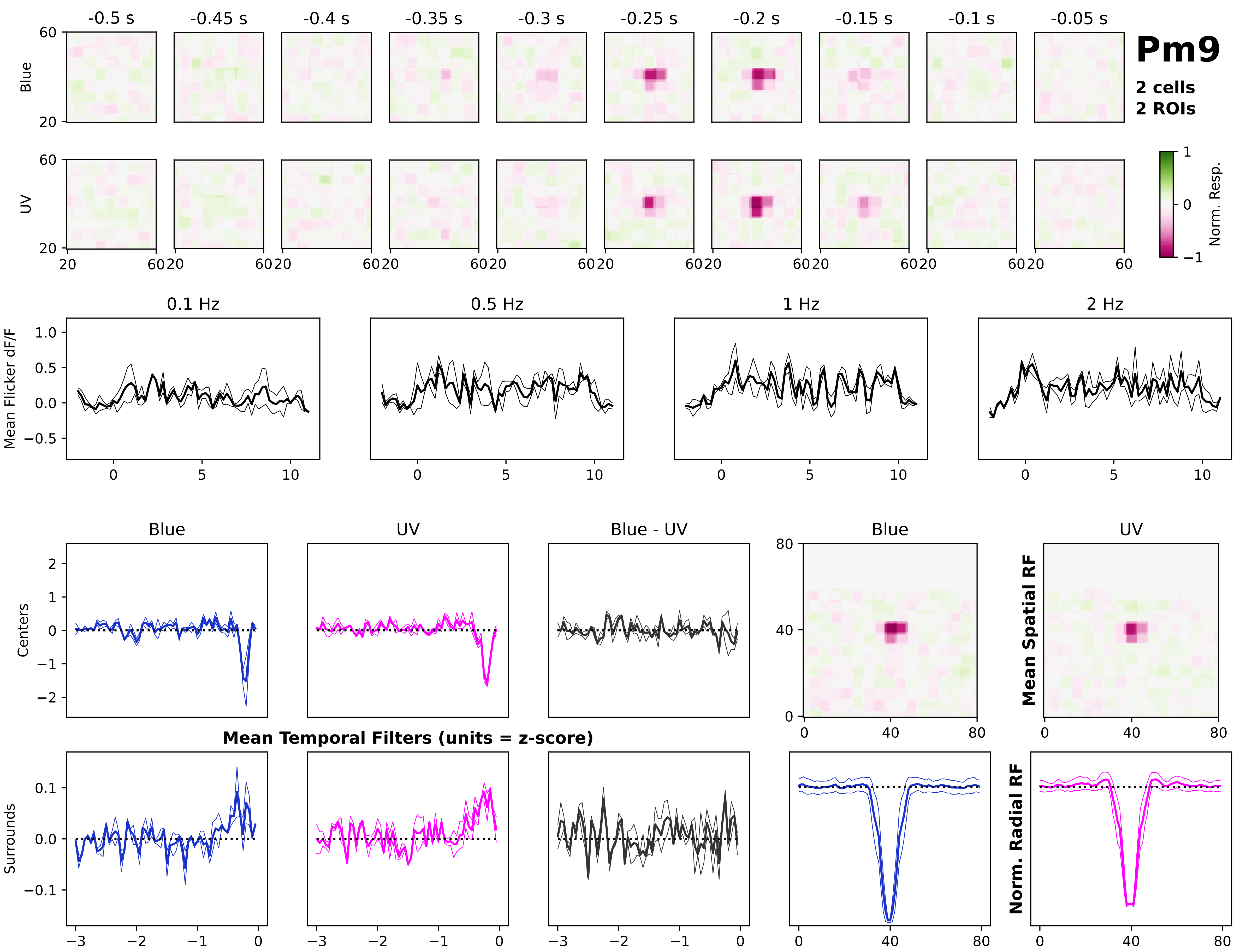
Pm9
2 cells
2 ROIs
-0.5 s
-0.45 s
-0.4 s
-0.35 s
-0.3 s
-0.25 s
-0.2 s
-0.15 s
-0.1 s
-0.05 s
Blue
UV
Norm. Resp.
0.1 Hz
0.5 Hz
1 Hz
2 Hz
Mean Flicker dF/F
Blue
UV
Blue - UV
Centers
Surrounds
Mean Temporal Filters (units = z-score)
Mean Spatial RF
Norm. Radial RF

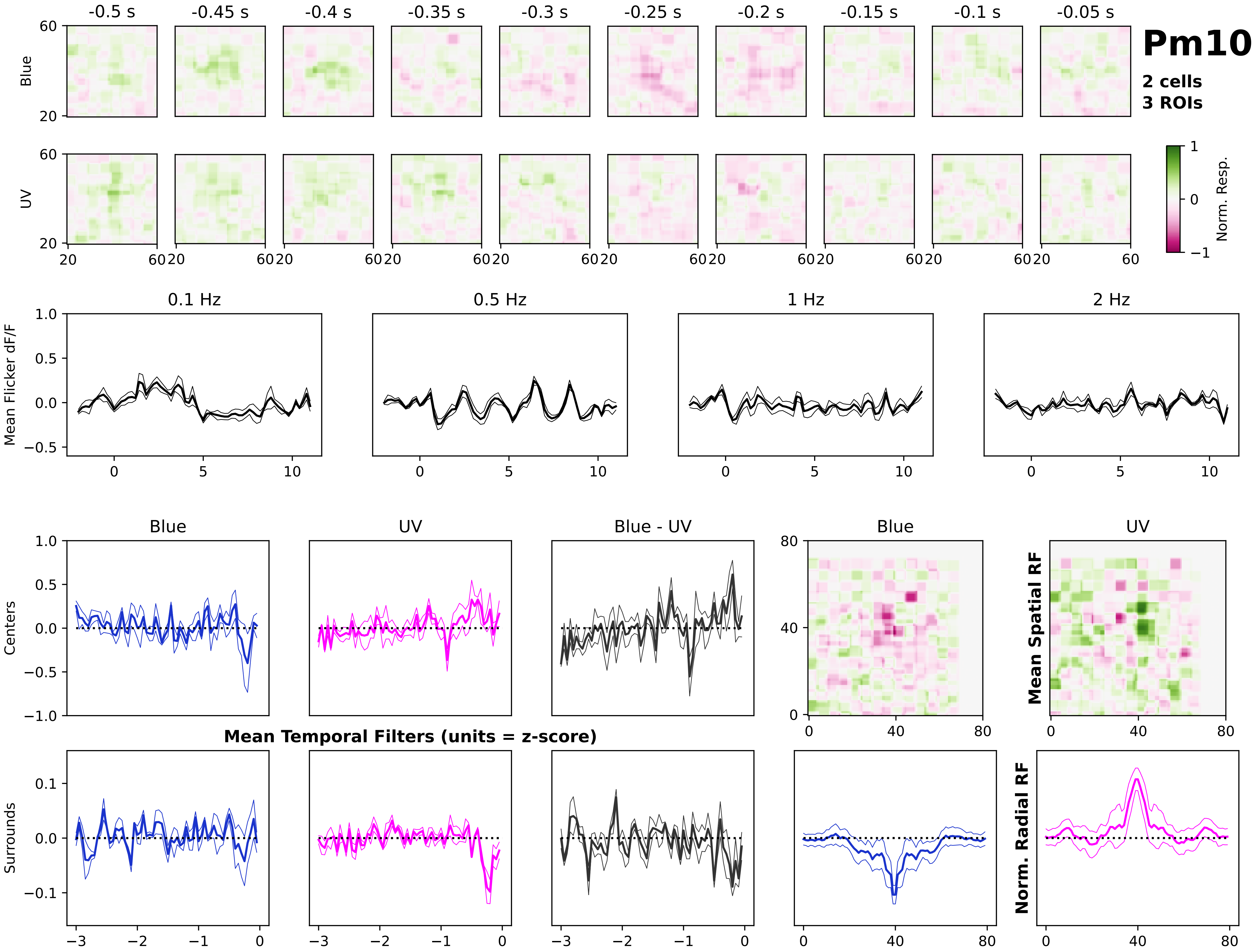

Pm10
2 cells
3 ROIs
-0.5 s
-0.45 s
-0.4 s
-0.35 s
-0.3 s
-0.25 s
-0.2 s
-0.15 s
-0.1 s
-0.05 s
Blue
UV
Norm. Resp.
0.1 Hz
0.5 Hz
1 Hz
2 Hz
Mean Flicker dF/F
Blue
UV
Blue - UV
Centers
Surrounds
Mean Temporal Filters (units = z-score)
Mean Spatial RF
Norm. Radial RF

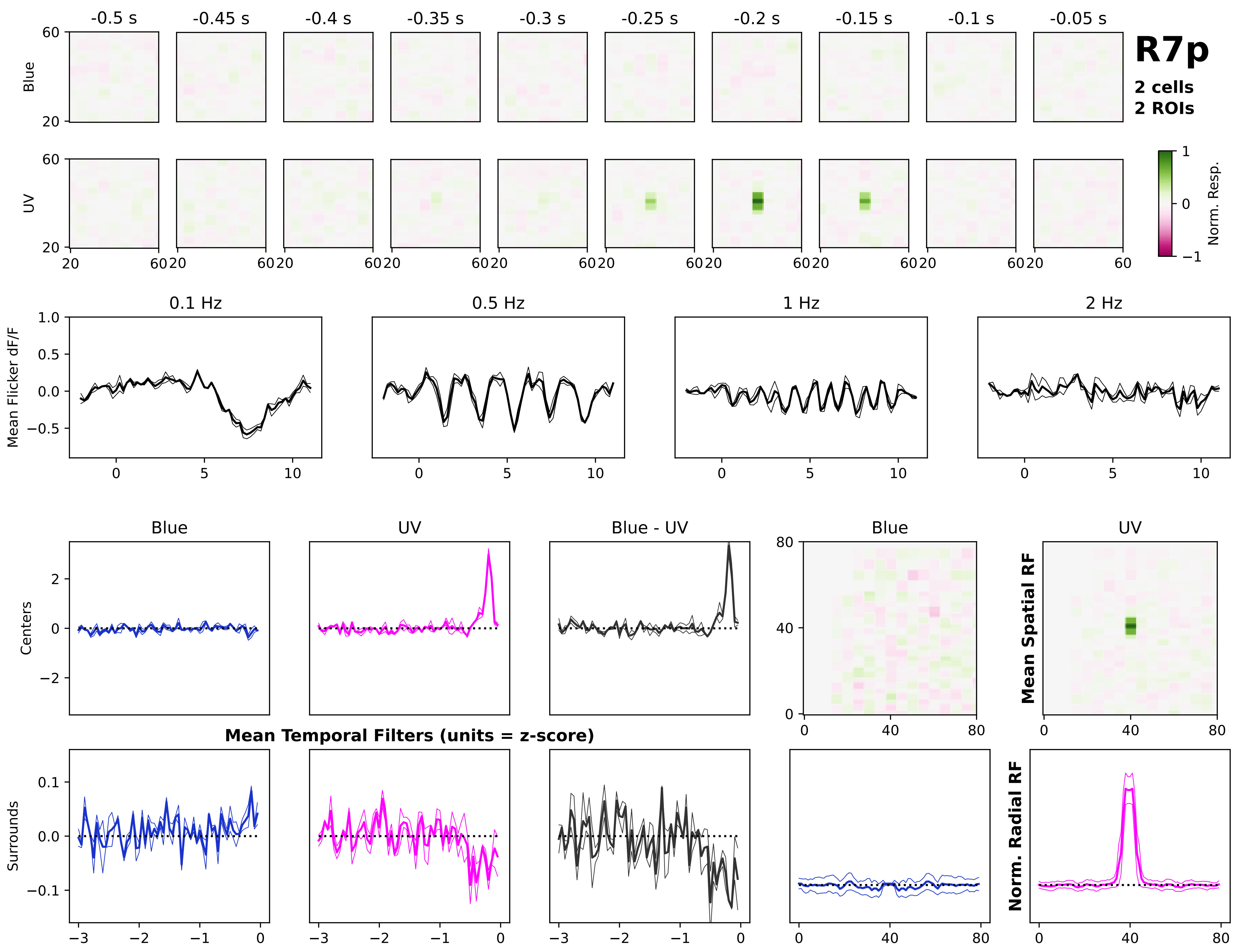

R7p
2 cells
2 ROIs
-0.5 s
-0.45 s
-0.4 s
-0.35 s
-0.3 s
-0.25 s
-0.2 s
-0.15 s
-0.1 s
-0.05 s
Blue
UV
Norm. Resp.
0.1 Hz
0.5 Hz
1 Hz
2 Hz
Mean Flicker dF/F
Blue
UV
Blue - UV
Centers
Surrounds
Mean Temporal Filters (units = z-score)
Mean Spatial RF
Norm. Radial RF

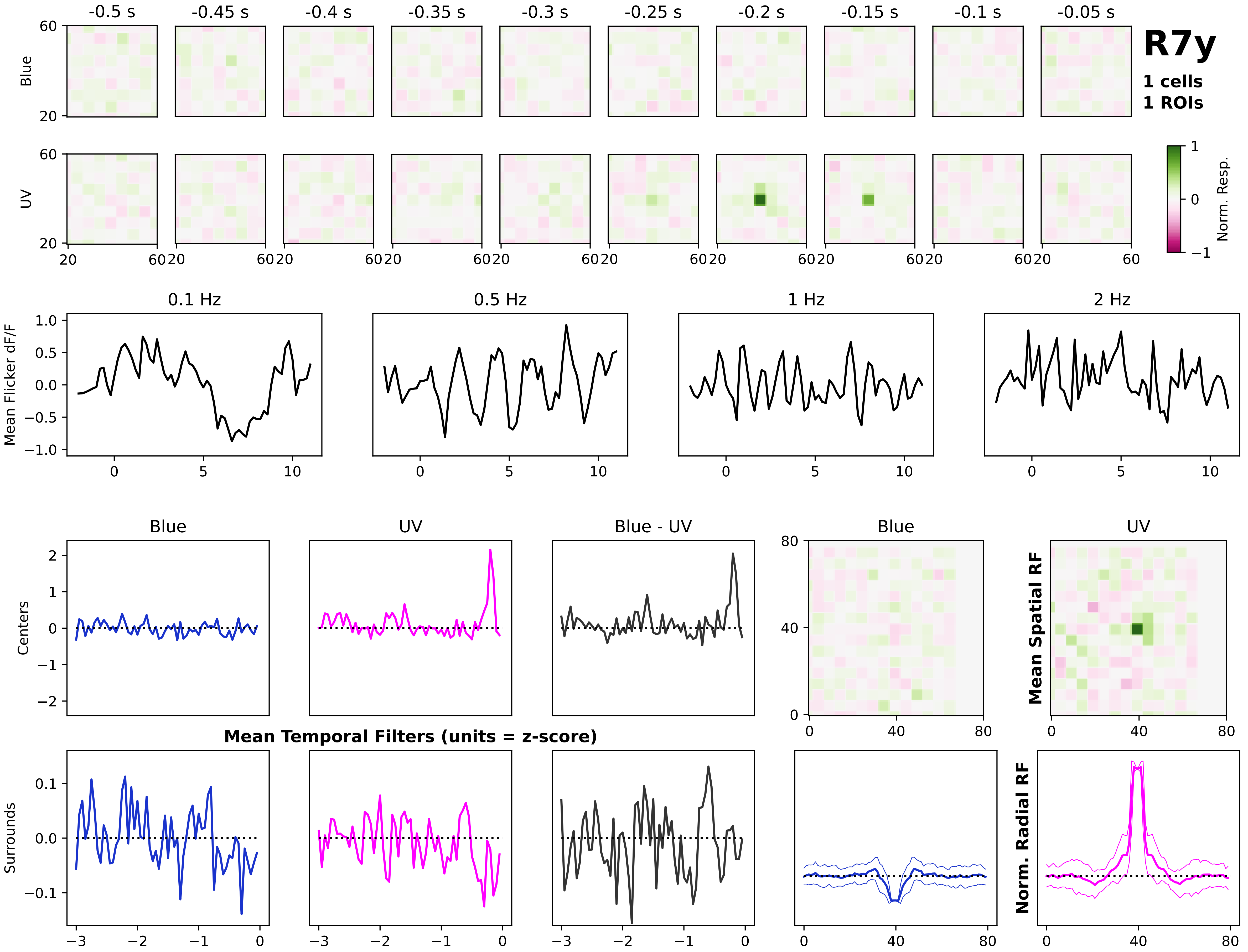
R7y
1 cells
1 ROIs
-0.5 s
-0.45 s
-0.4 s
-0.35 s
-0.3 s
-0.25 s
-0.2 s
-0.15 s
-0.1 s
-0.05 s
Blue
UV
Norm. Resp.
0.1 Hz
0.5 Hz
1 Hz
2 Hz
Mean Flicker dF/F
Blue
UV
Blue - UV
Centers
Surrounds
Mean Temporal Filters (units = z-score)
Mean Spatial RF
Norm. Radial RF

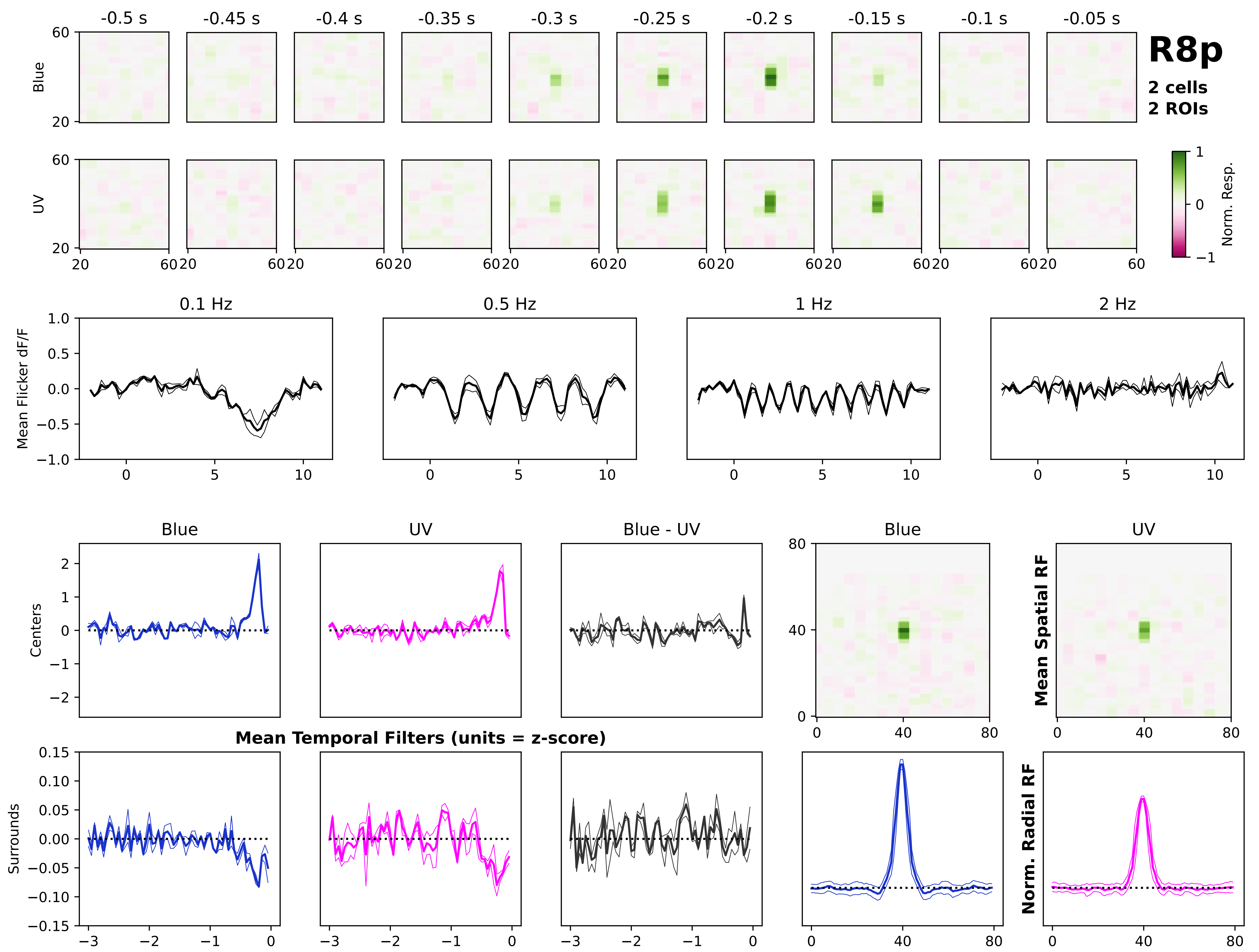
R8p
2 cells
2 ROIs
-0.5 s
-0.45 s
-0.4 s
-0.35 s
-0.3 s
-0.25 s
-0.2 s
-0.15 s
-0.1 s
-0.05 s
Blue
UV
Norm. Resp.
0.1 Hz
0.5 Hz
1 Hz
2 Hz
Mean Flicker dF/F
Blue
UV
Blue - UV
Centers
Surrounds
Mean Temporal Filters (units = z-score)
Mean Spatial RF
Norm. Radial RF

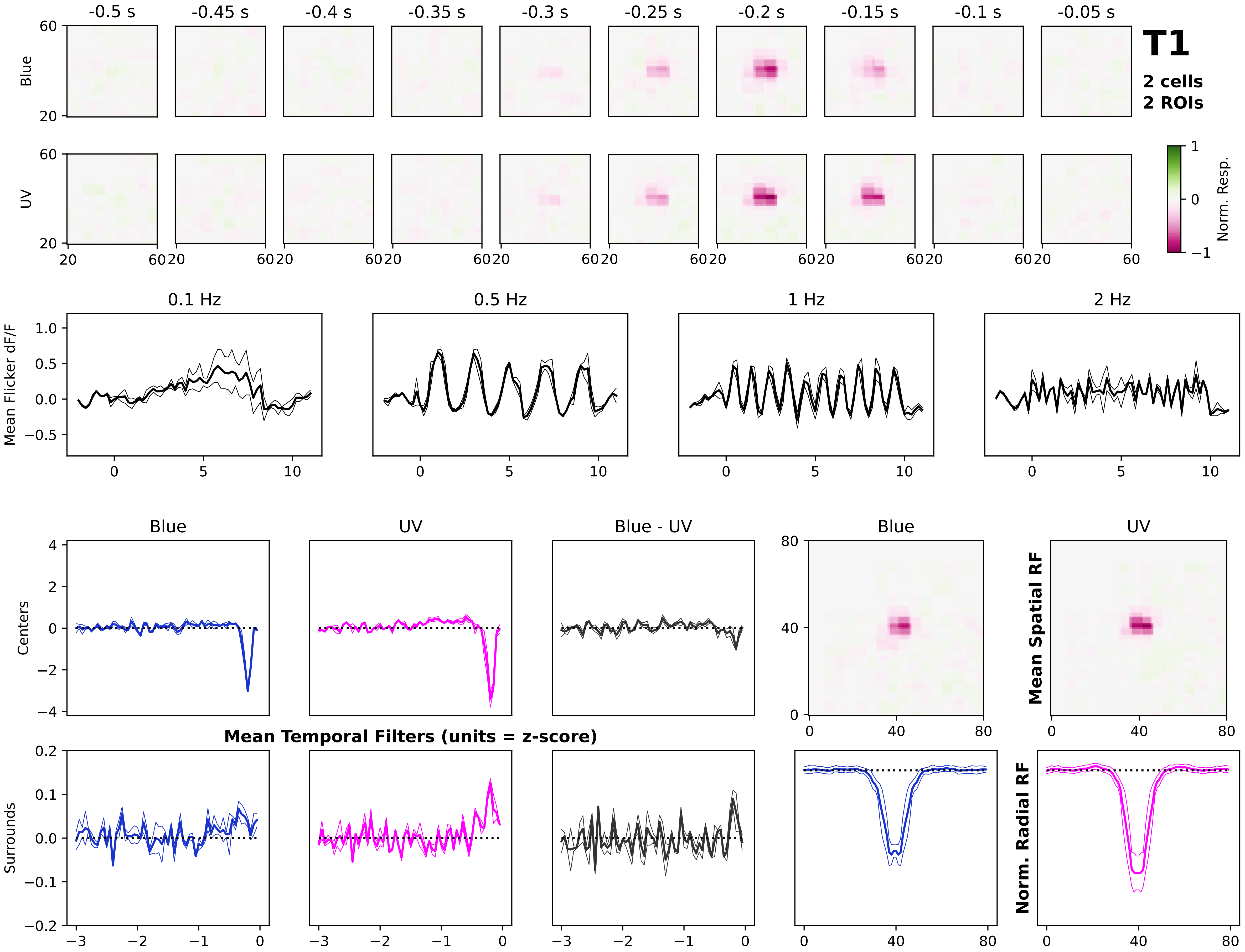
T1
2 cells
2 ROIs
-0.5 s
-0.45 s
-0.4 s
-0.35 s
-0.3 s
-0.25 s
-0.2 s
-0.15 s
-0.1 s
-0.05 s
Blue
UV
Norm. Resp.
0.1 Hz
0.5 Hz
1 Hz
2 Hz
Mean Flicker dF/F
Centers
Surrounds
Blue
UV
Blue - UV
Mean Temporal Filters (units = z-score)
Mean Spatial RF
Norm. Radial RF

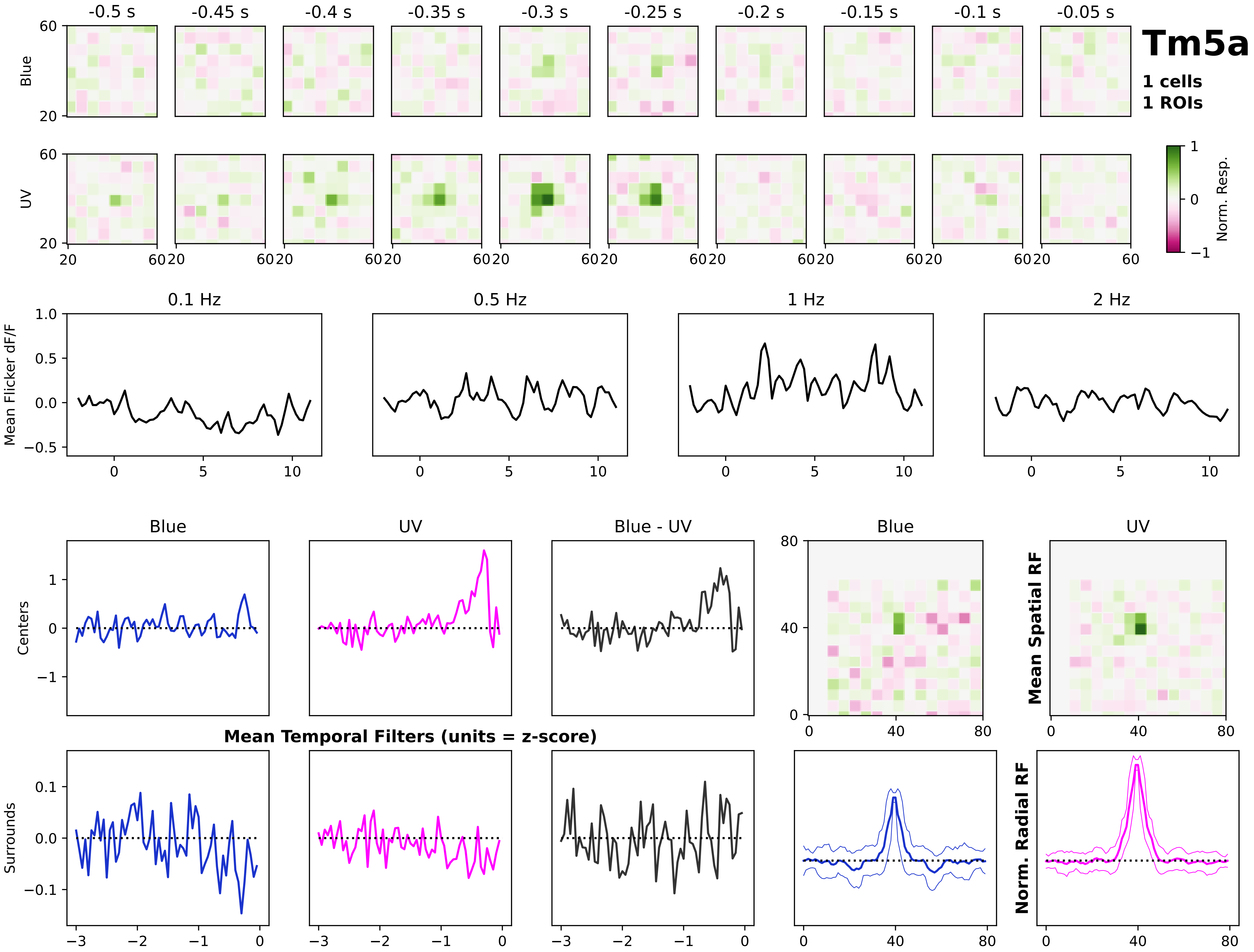

Tm5a
1 cells
1 ROIs
-0.5 s
-0.45 s
-0.4 s
-0.35 s
-0.3 s
-0.25 s
-0.2 s
-0.15 s
-0.1 s
-0.05 s
Blue
UV
Norm. Resp.
0.1 Hz
0.5 Hz
1 Hz
2 Hz
Mean Flicker dF/F
Blue
UV
Blue - UV
Centers
Surrounds
Mean Temporal Filters (units = z-score)
Mean Spatial RF
Norm. Radial RF

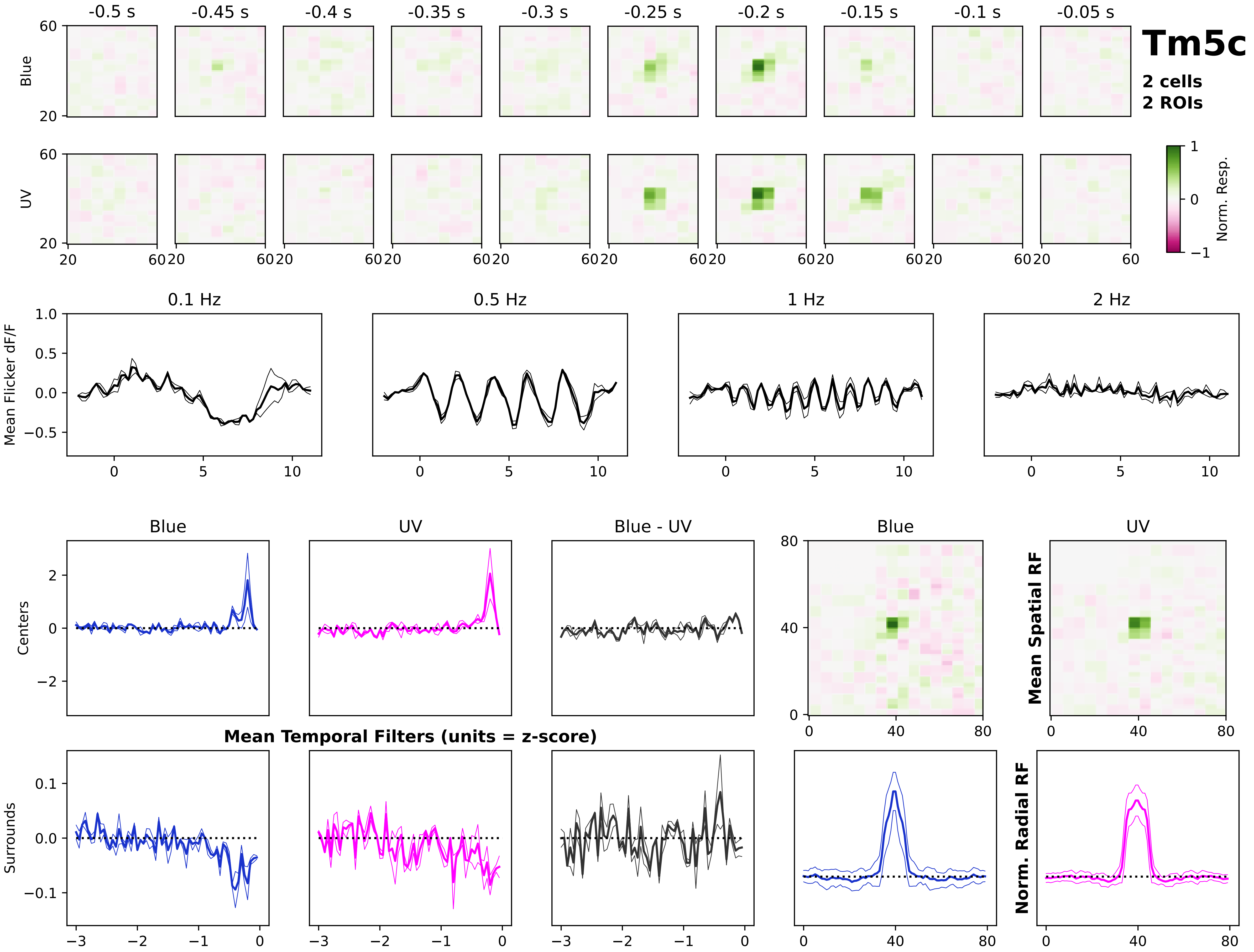
Tm5c
2 cells
2 ROIs
-0.5 s
-0.45 s
-0.4 s
-0.35 s
-0.3 s
-0.25 s
-0.2 s
-0.15 s
-0.1 s
-0.05 s
Blue
UV
Norm. Resp.
0.1 Hz
0.5 Hz
1 Hz
2 Hz
Mean Flicker dF/F
Blue
UV
Blue - UV
Centers
Surrounds
Mean Temporal Filters (units = z-score)
Mean Spatial RF
Norm. Radial RF

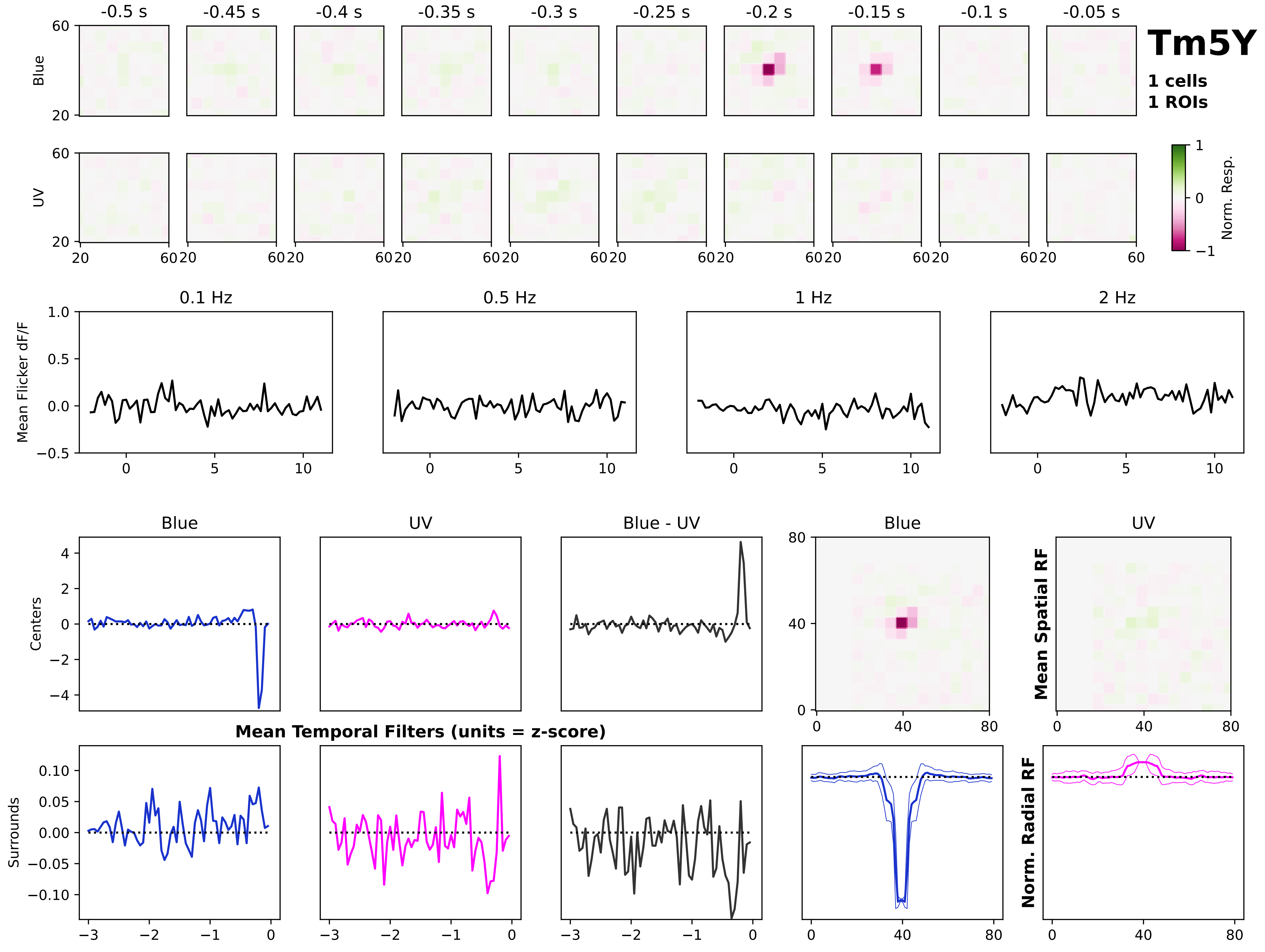
Tm5Y
1 cells
1 ROIs
-0.5 s
-0.45 s
-0.4 s
-0.35 s
-0.3 s
-0.25 s
-0.2 s
-0.15 s
-0.1 s
-0.05 s
Blue
UV
Norm. Resp.
0.1 Hz
0.5 Hz
1 Hz
2 Hz
Mean Flicker dF/F
Blue
UV
Blue - UV
Centers
Surrounds
Mean Temporal Filters (units = z-score)
Mean Spatial RF
Norm. Radial RF

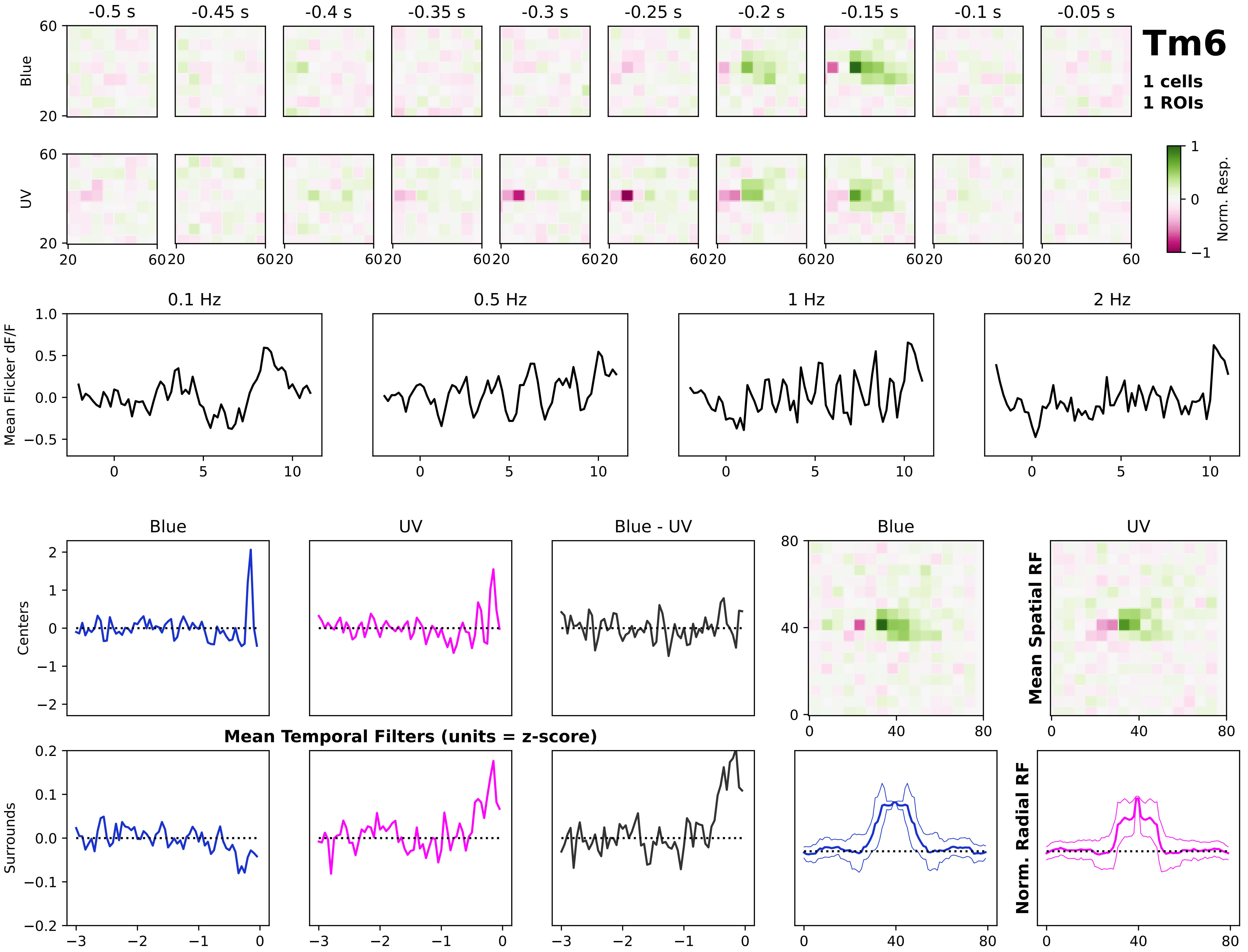

Tm6
1 cells
1 ROIs
-0.5 s
-0.45 s
-0.4 s
-0.35 s
-0.3 s
-0.25 s
-0.2 s
-0.15 s
-0.1 s
-0.05 s
Blue
UV
Norm. Resp.
0.1 Hz
0.5 Hz
1 Hz
2 Hz
Mean Flicker dF/F
Blue
UV
Blue - UV
Centers
Surrounds
Mean Temporal Filters (units = z-score)
Mean Spatial RF
Norm. Radial RF

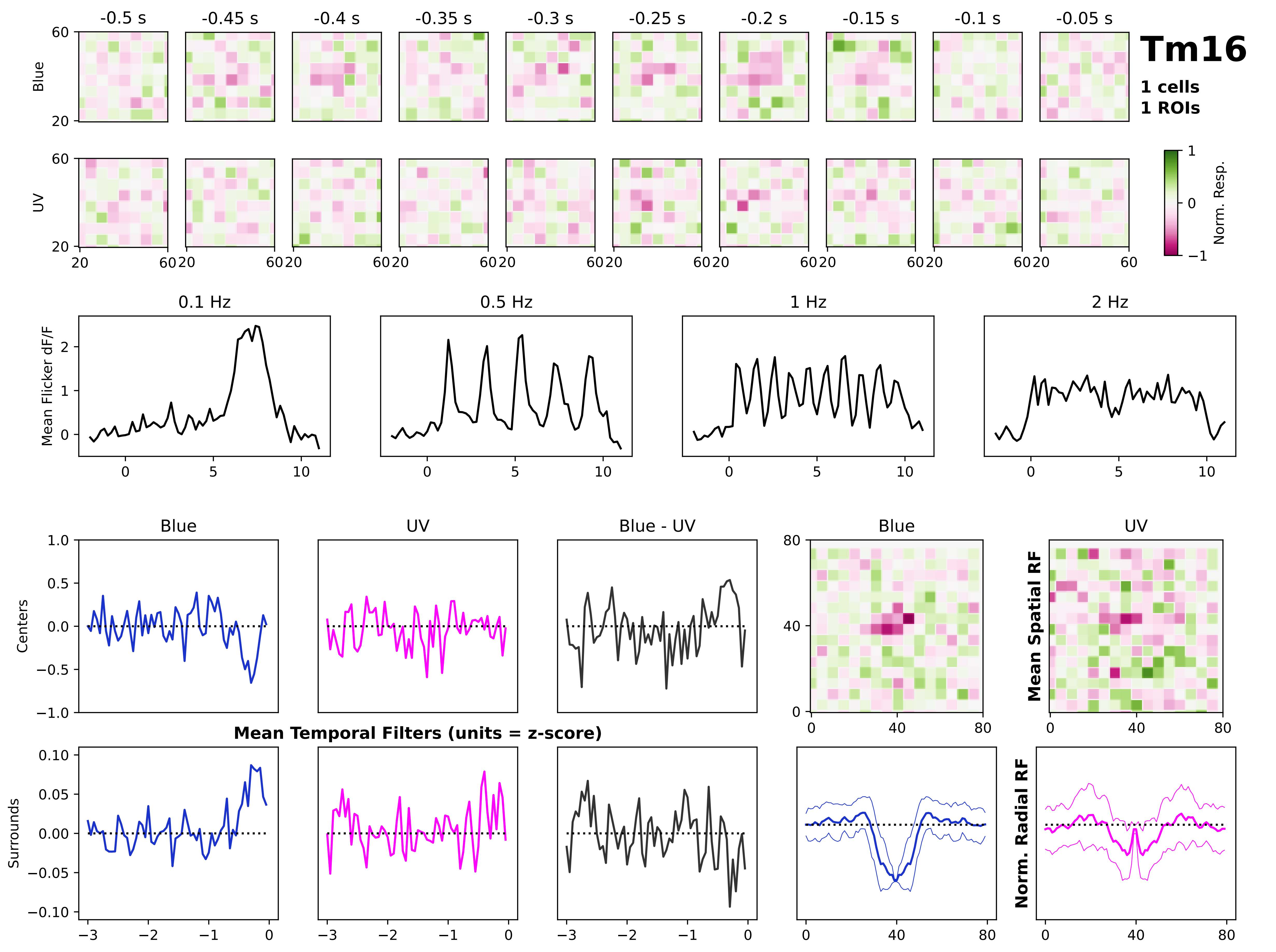

Tm16
1 cells
1 ROIs
-0.5 s
-0.45 s
-0.4 s
-0.35 s
-0.3 s
-0.25 s
-0.2 s
-0.15 s
-0.1 s
-0.05 s
Blue
UV
Norm. Resp.
0.1 Hz
0.5 Hz
1 Hz
2 Hz
Mean Flicker dF/F
Centers
Surrounds
Blue
UV
Blue - UV
Mean Temporal Filters (units = z-score)
Mean Spatial RF
Norm. Radial RF

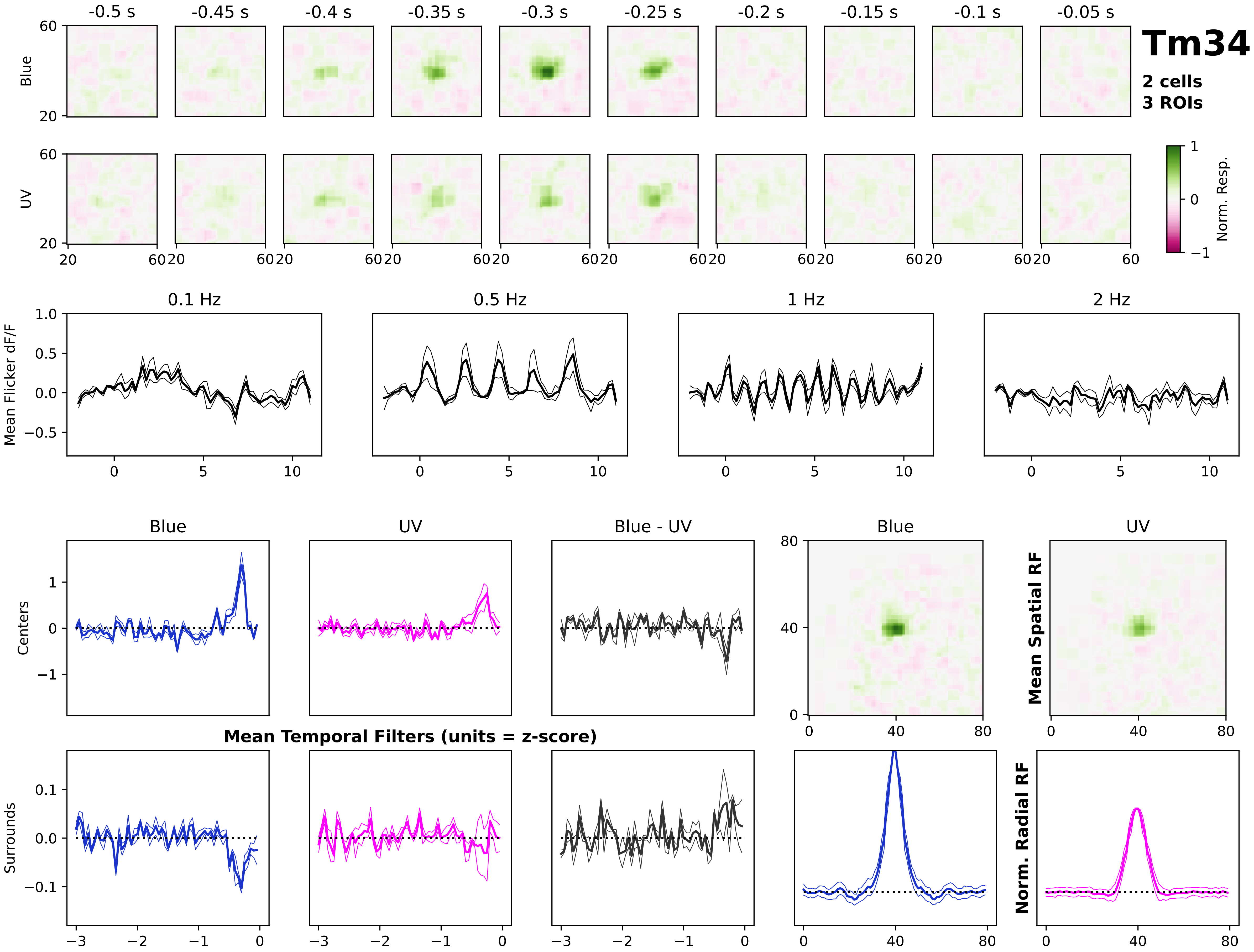

Tm34
2 cells
3 ROIs
-0.5 s
-0.45 s
-0.4 s
-0.35 s
-0.3 s
-0.25 s
-0.2 s
-0.15 s
-0.1 s
-0.05 s
Blue
UV
Norm. Resp.
0.1 Hz
0.5 Hz
1 Hz
2 Hz
Mean Flicker dF/F
Blue
UV
Blue - UV
Centers
Surrounds
Mean Temporal Filters (units = z-score)
Mean Spatial RF
Norm. Radial RF

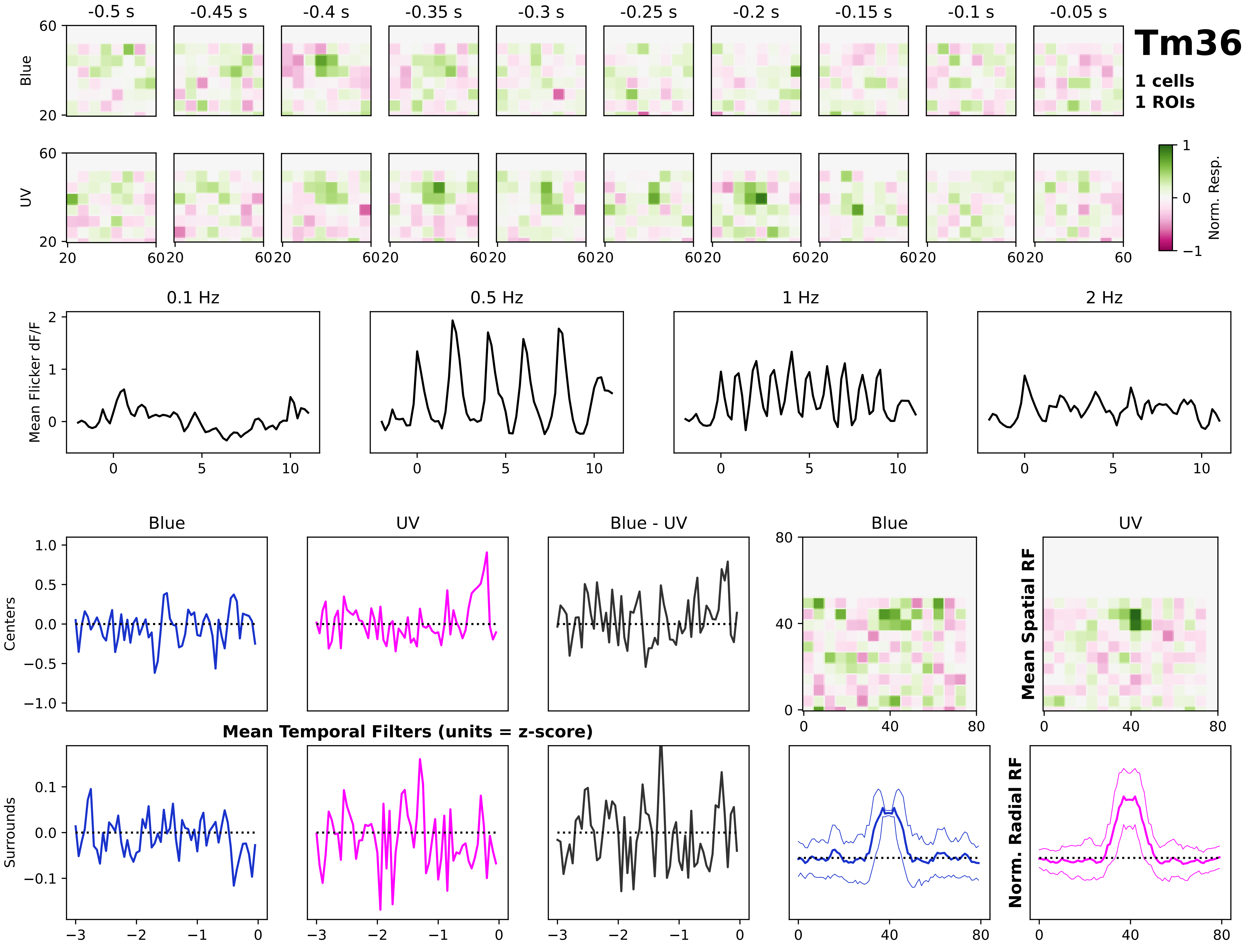
Tm36
1 cells
1 ROIs
-0.5 s
-0.45 s
-0.4 s
-0.35 s
-0.3 s
-0.25 s
-0.2 s
-0.15 s
-0.1 s
-0.05 s
Blue
UV
Norm. Resp.
0.1 Hz
0.5 Hz
1 Hz
2 Hz
Mean Flicker dF/F
Blue
UV
Blue - UV
Centers
Surrounds
Mean Temporal Filters (units = z-score)
Mean Spatial RF
Norm. Radial RF

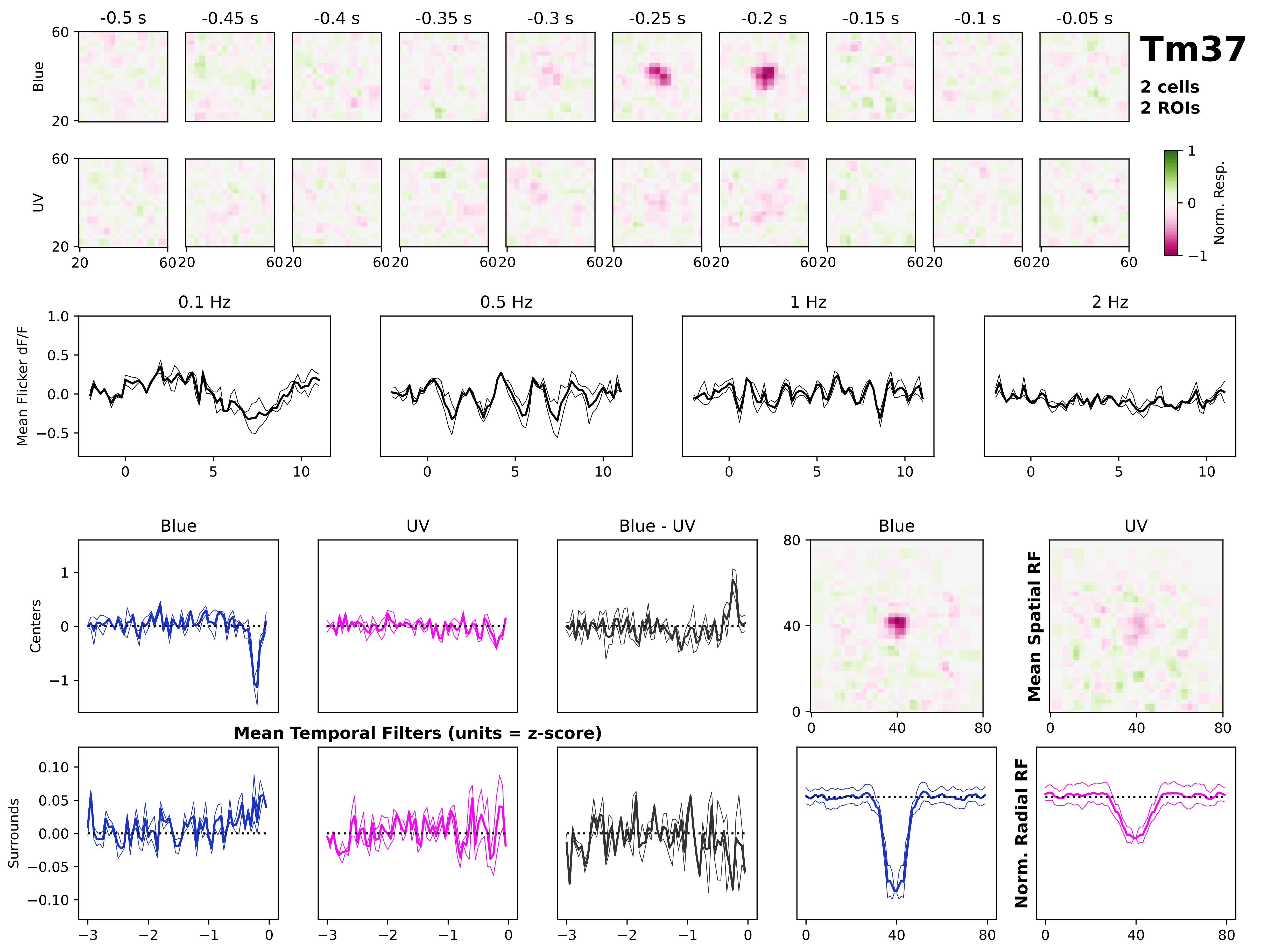

Tm37
2 cells
2 ROIs
-0.5 s
-0.45 s
-0.4 s
-0.35 s
-0.3 s
-0.25 s
-0.2 s
-0.15 s
-0.1 s
-0.05 s
Blue
UV
Norm. Resp.
0.1 Hz
0.5 Hz
1 Hz
2 Hz
Mean Flicker dF/F
Blue
UV
Blue - UV
Centers
Surrounds
Mean Temporal Filters (units = z-score)
Mean Spatial RF
Norm. Radial RF

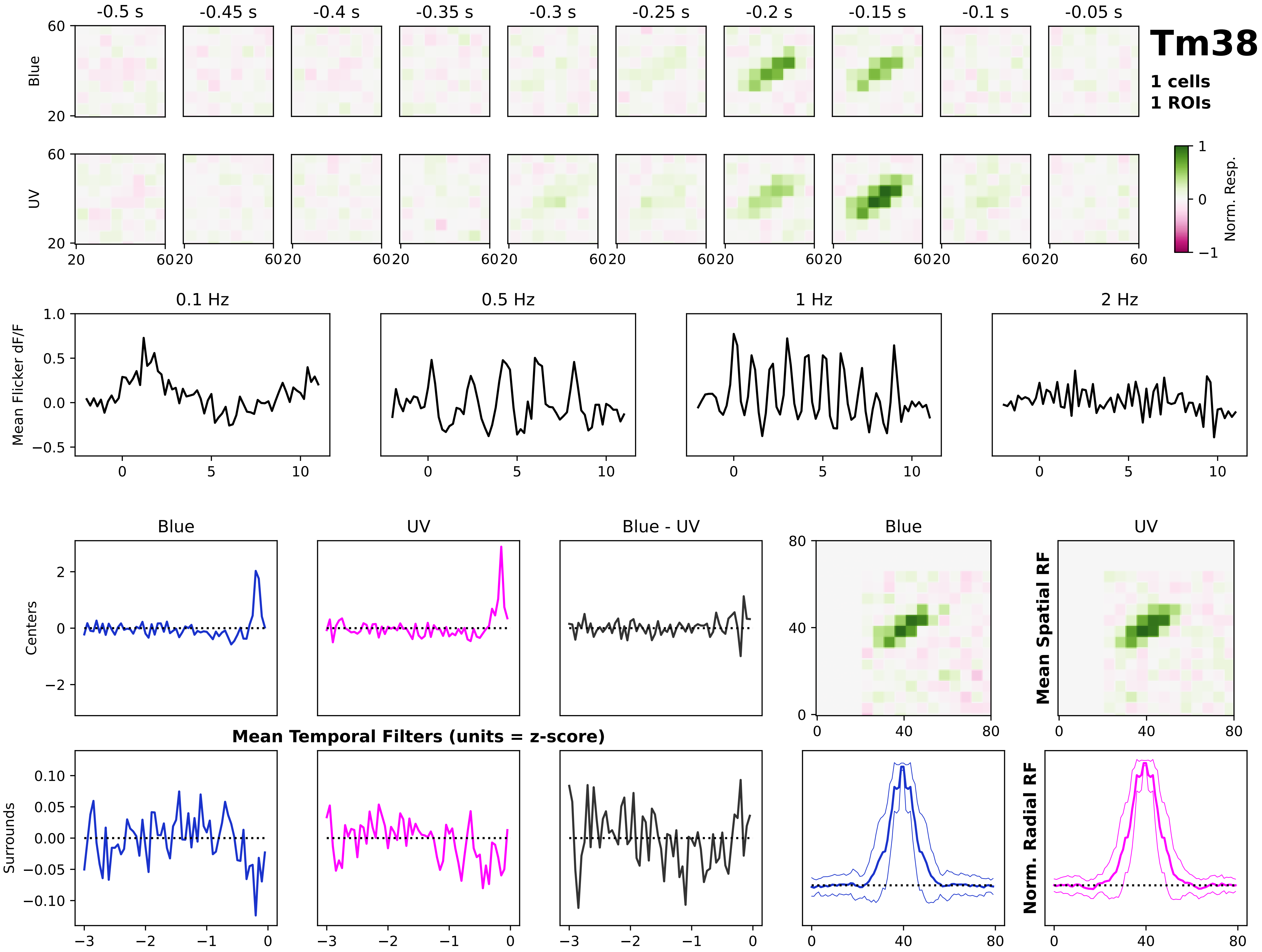
Tm38
1 cells
1 ROIs
-0.5 s
-0.45 s
-0.4 s
-0.35 s
-0.3 s
-0.25 s
-0.2 s
-0.15 s
-0.1 s
-0.05 s
Blue
UV
Norm. Resp.
0.1 Hz
0.5 Hz
1 Hz
2 Hz
Mean Flicker dF/F
Blue
UV
Blue - UV
Centers
Surrounds
Mean Temporal Filters (units = z-score)
Mean Spatial RF
Norm. Radial RF

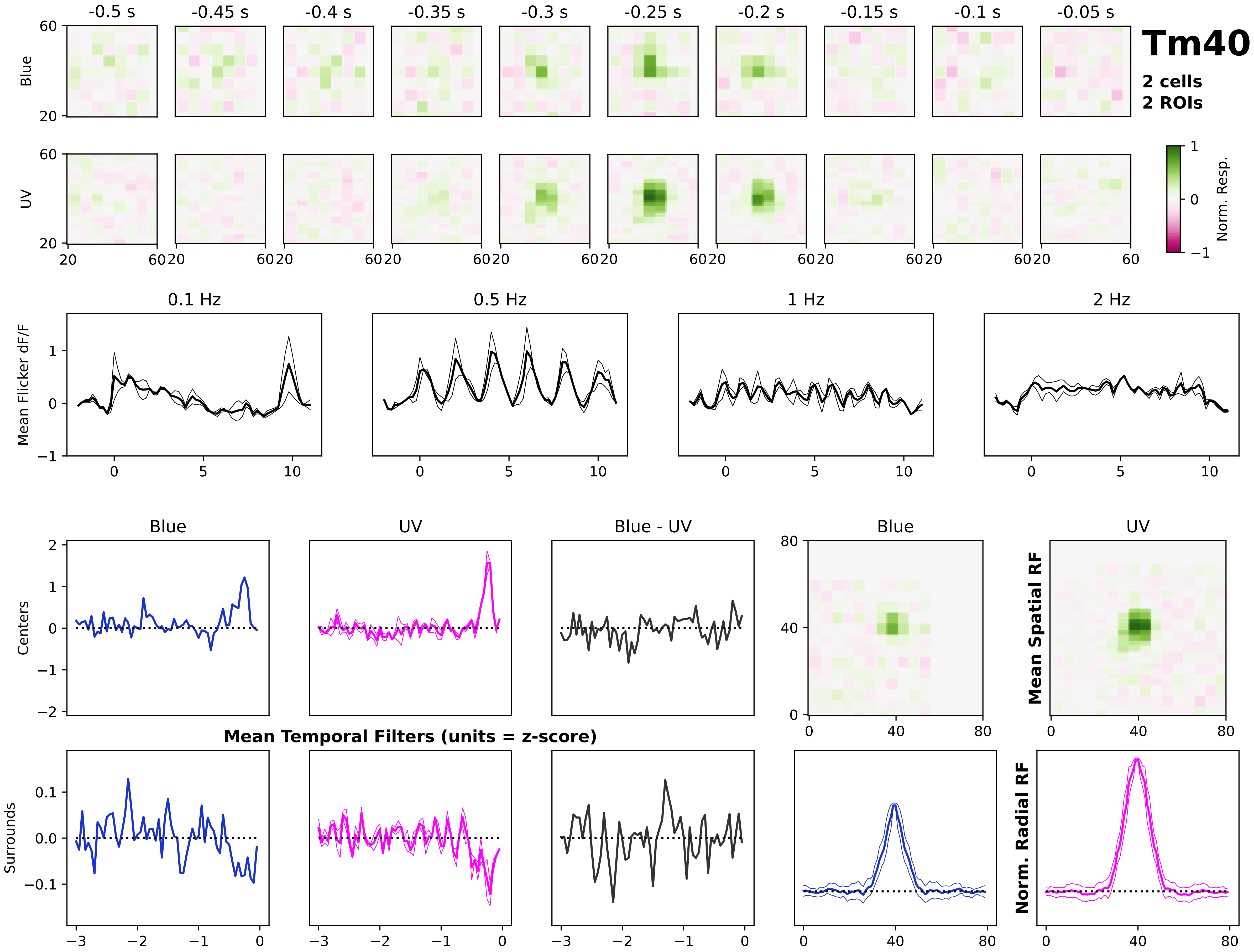
Tm40
2 cells
2 ROIs
-0.5 s
-0.45 s
-0.4 s
-0.35 s
-0.3 s
-0.25 s
-0.2 s
-0.15 s
-0.1 s
-0.05 s
Blue
UV
Norm. Resp.
0.1 Hz
0.5 Hz
1 Hz
2 Hz
Mean Flicker dF/F
Blue
UV
Blue - UV
Centers
Surrounds
Mean Temporal Filters (units = z-score)
Mean Spatial RF
Norm. Radial RF

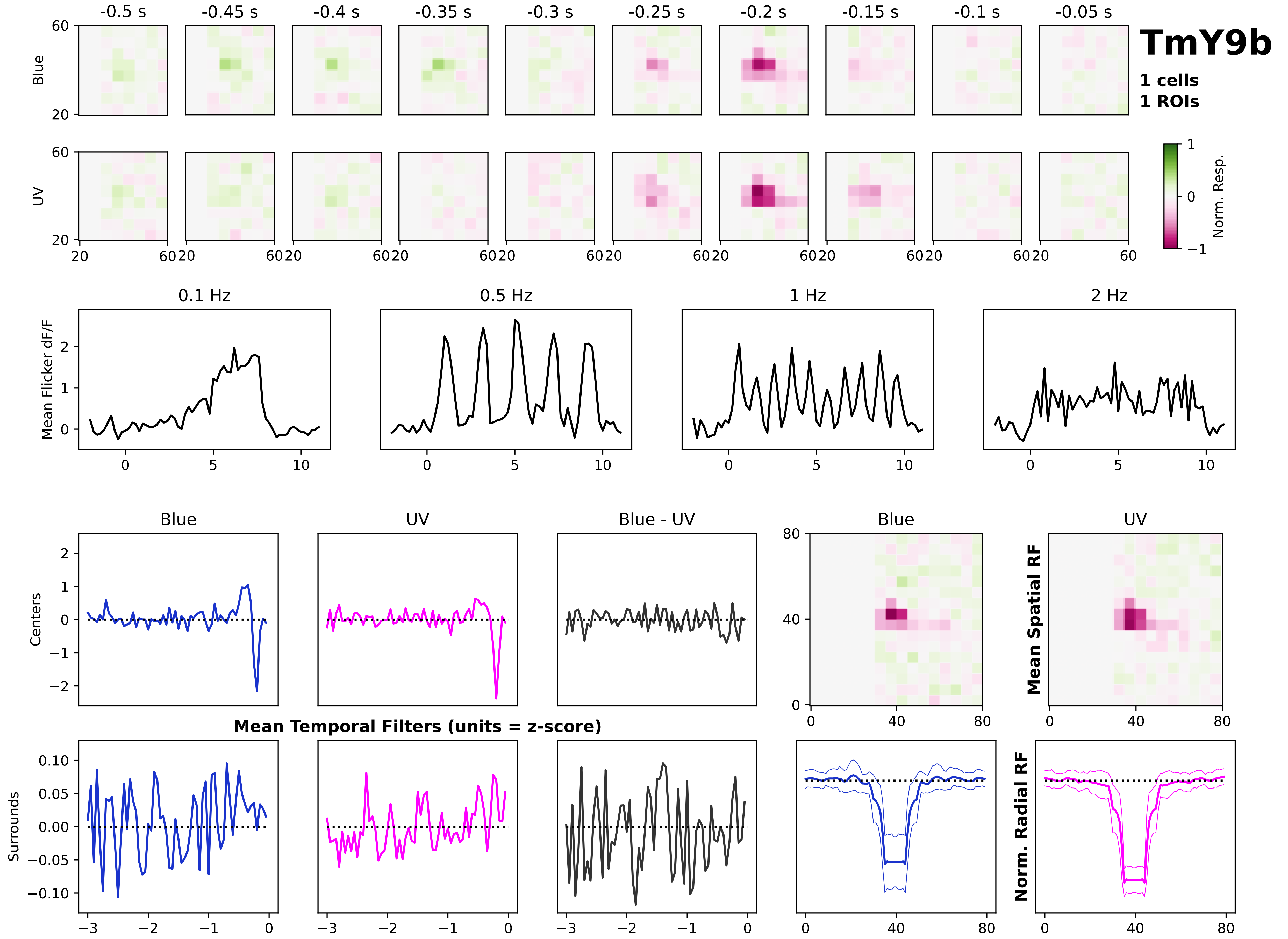

TmY9b
1 cells
1 ROIs
-0.5 s
-0.45 s
-0.4 s
-0.35 s
-0.3 s
-0.25 s
-0.2 s
-0.15 s
-0.1 s
-0.05 s
Blue
UV
Norm. Resp.
0.1 Hz
0.5 Hz
1 Hz
2 Hz
Mean Flicker dF/F
Blue
UV
Blue - UV
Centers
Surrounds
Mean Temporal Filters (units = z-score)
Mean Spatial RF
Norm. Radial RF

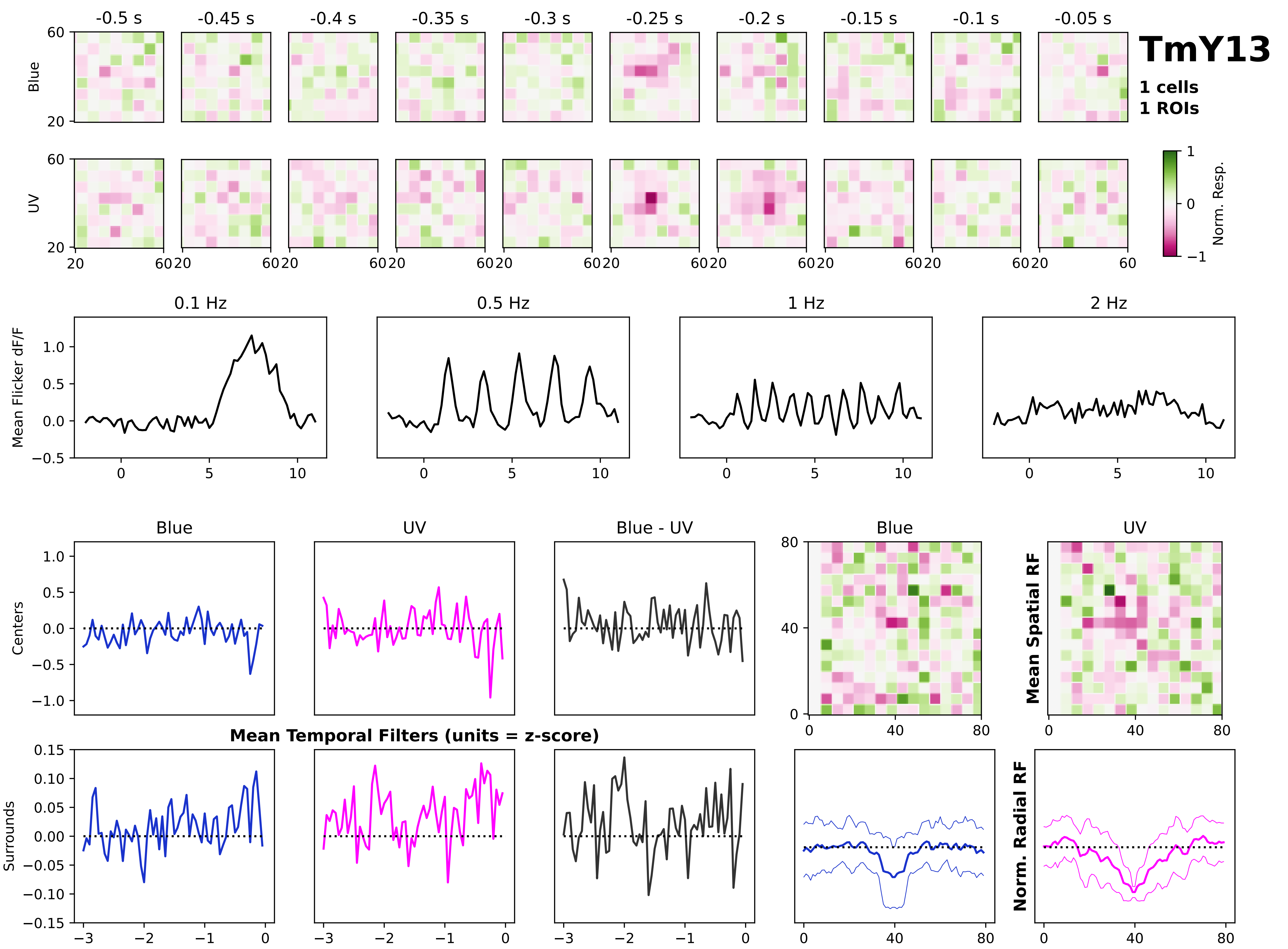
TmY13
1 cells
1 ROIs
-0.5 s
-0.45 s
-0.4 s
-0.35 s
-0.3 s
-0.25 s
-0.2 s
-0.15 s
-0.1 s
-0.05 s
Blue
UV
Norm. Resp.
0.1 Hz
0.5 Hz
1 Hz
2 Hz
Mean Flicker dF/F
Blue
UV
Blue - UV
Centers
Surrounds
Mean Temporal Filters (units = z-score)
Mean Spatial RF
Norm. Radial RF

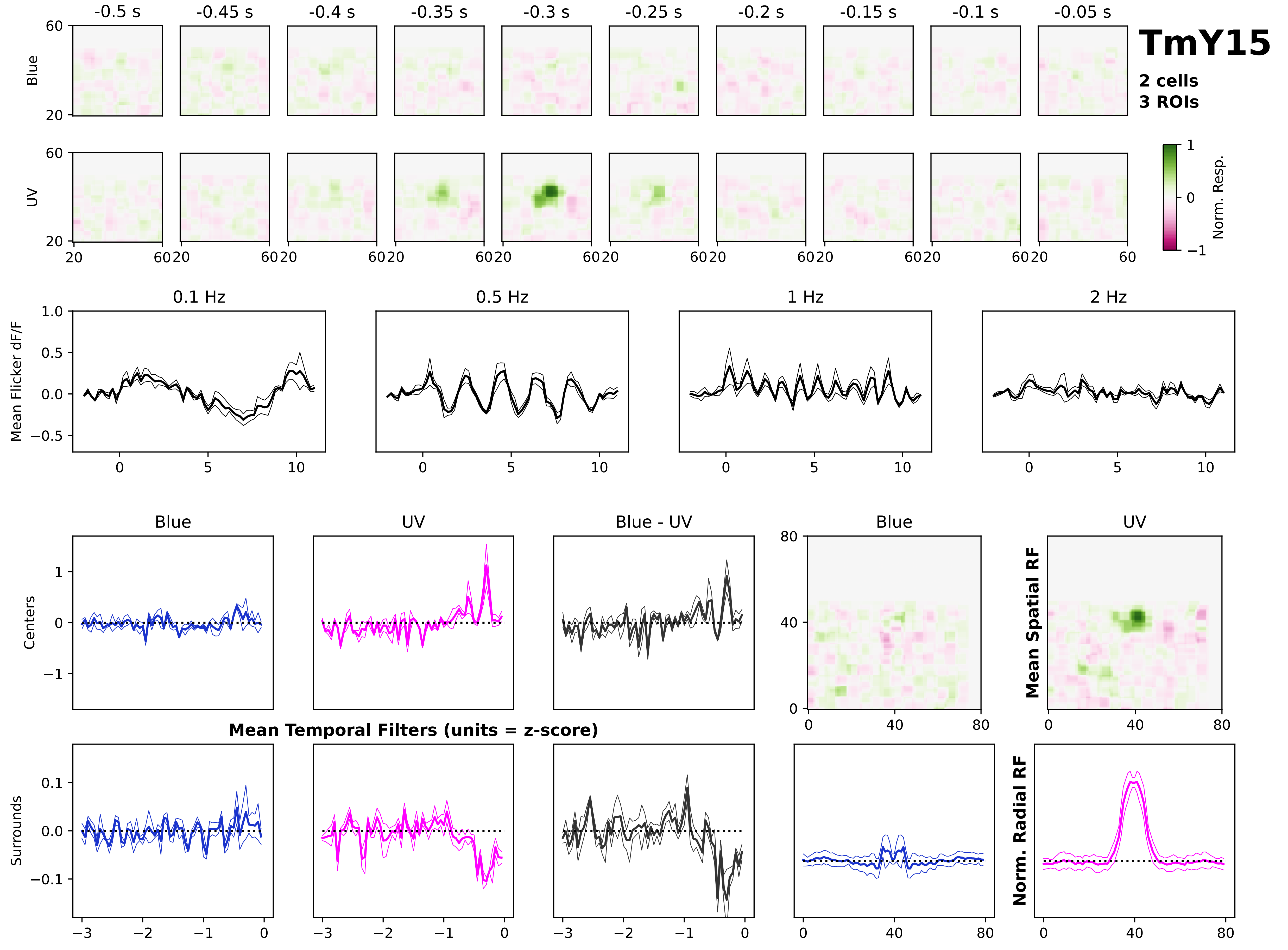
TmY15
2 cells
3 ROIs
-0.5 s
-0.45 s
-0.4 s
-0.35 s
-0.3 s
-0.25 s
-0.2 s
-0.15 s
-0.1 s
-0.05 s
Blue
UV
Norm. Resp.
0.1 Hz
0.5 Hz
1 Hz
2 Hz
Mean Flicker dF/F
Blue
UV
Blue - UV
Centers
Surrounds
Mean Temporal Filters (units = z-score)
Mean Spatial RF
Norm. Radial RF

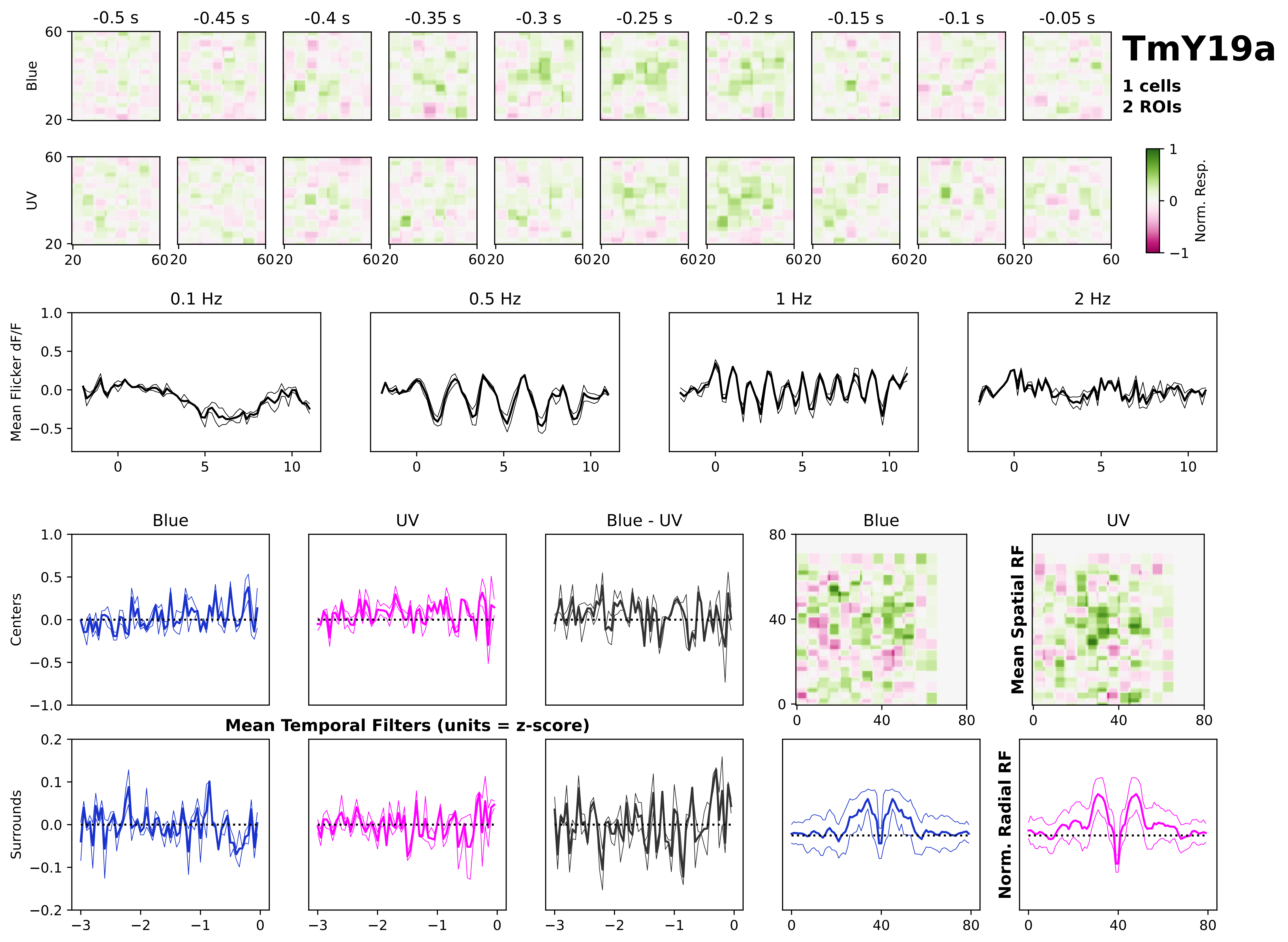
TmY19a
1 cells
2 ROIs
-0.5 s
-0.45 s
-0.4 s
-0.35 s
-0.3 s
-0.25 s
-0.2 s
-0.15 s
-0.1 s
-0.05 s
Blue
UV
Norm. Resp.
0.1 Hz
0.5 Hz
1 Hz
2 Hz
Mean Flicker dF/F
Blue
UV
Blue - UV
Centers
Surrounds
Mean Temporal Filters (units = z-score)
Mean Spatial RF
Norm. Radial RF

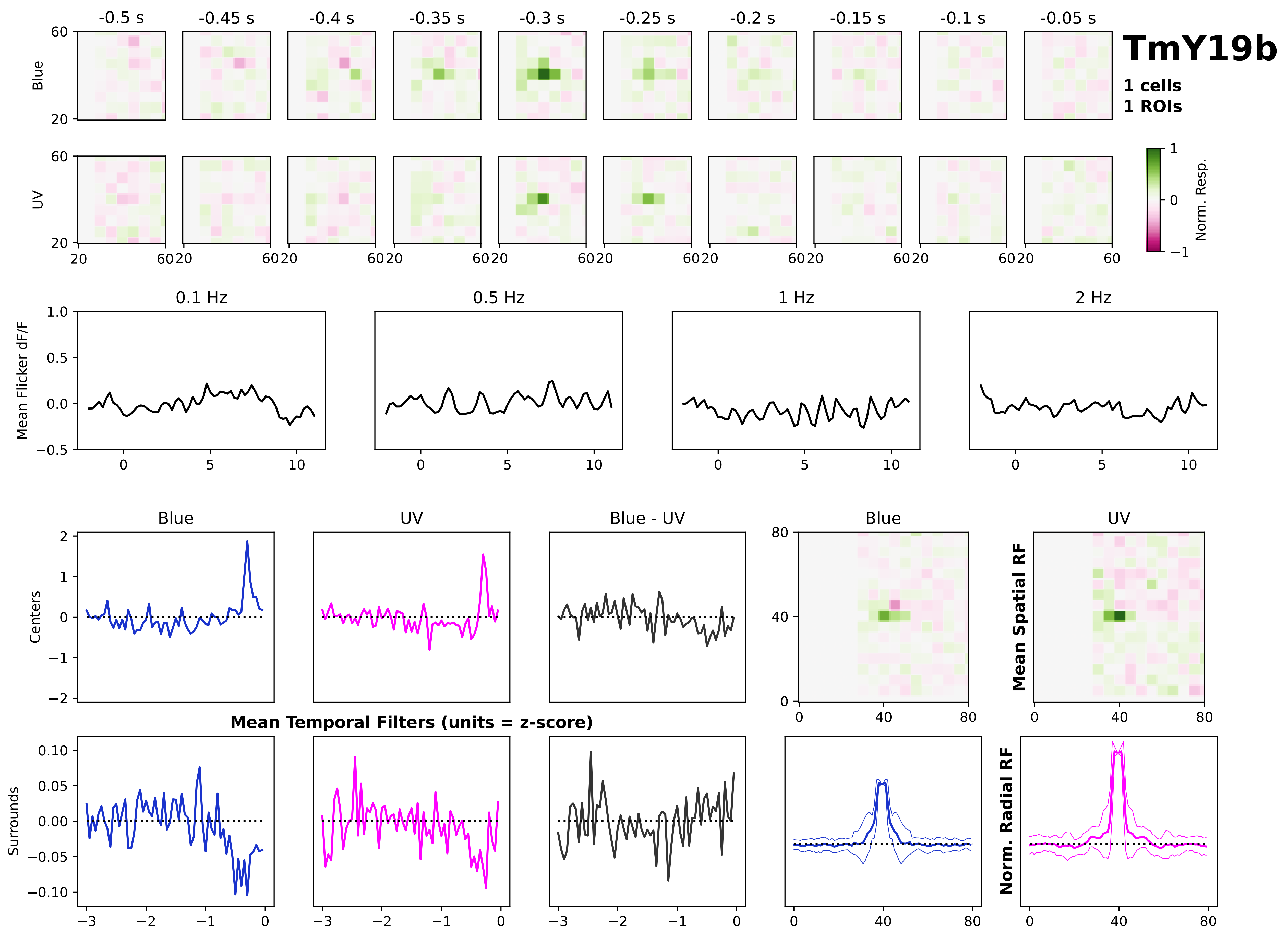

TmY19b
1 cells
1 ROIs
-0.5 s
-0.45 s
-0.4 s
-0.35 s
-0.3 s
-0.25 s
-0.2 s
-0.15 s
-0.1 s
-0.05 s
Blue
UV
Norm. Resp.
0.1 Hz
0.5 Hz
1 Hz
2 Hz
Mean Flicker dF/F
Blue
UV
Blue - UV
Centers
Surrounds
Mean Temporal Filters (units = z-score)
Mean Spatial RF
Norm. Radial RF

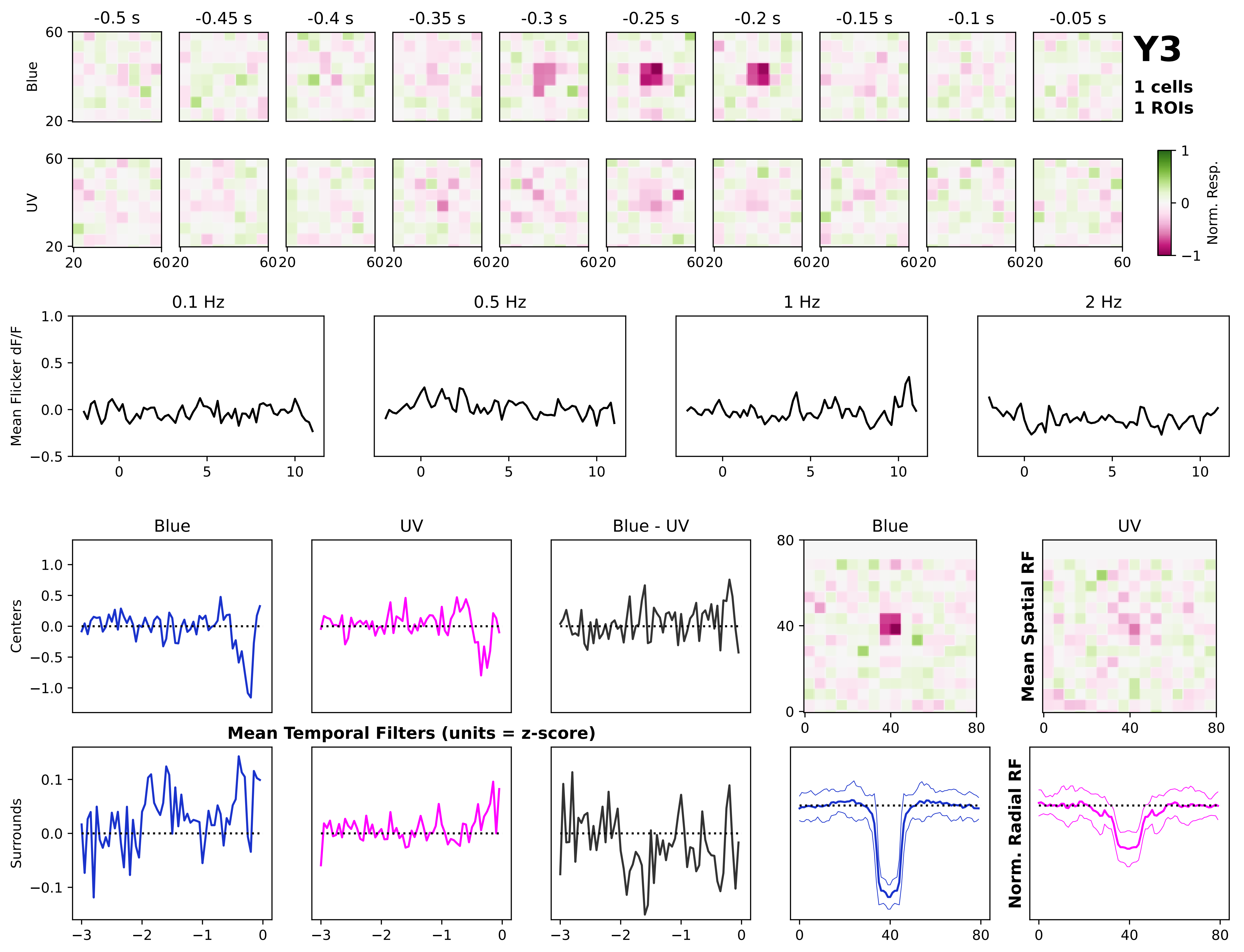
Y3
1 cells
1 ROIs
-0.5 s
-0.45 s
-0.4 s
-0.35 s
-0.3 s
-0.25 s
-0.2 s
-0.15 s
-0.1 s
-0.05 s
Blue
UV
Norm. Resp.
0.1 Hz
0.5 Hz
1 Hz
2 Hz
Mean Flicker dF/F
Blue
UV
Blue - UV
Centers
Surrounds
Mean Temporal Filters (units = z-score)
Mean Spatial RF
Norm. Radial RF

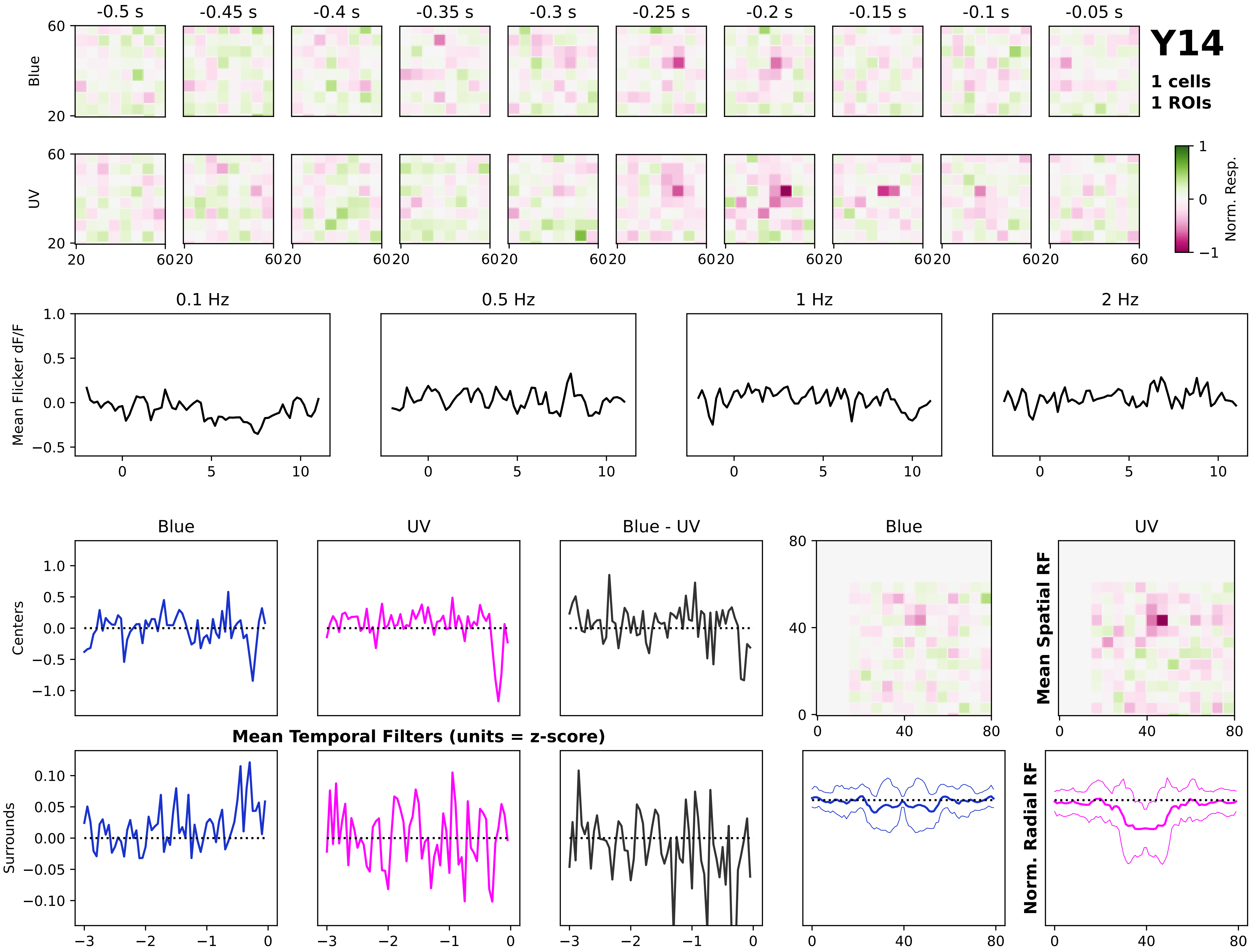
Y14
1 cells
1 ROIs
-0.5 s
-0.45 s
-0.4 s
-0.35 s
-0.3 s
-0.25 s
-0.2 s
-0.15 s
-0.1 s
-0.05 s
Blue
UV
Norm. Resp.
0.1 Hz
0.5 Hz
1 Hz
2 Hz
Mean Flicker dF/F
Blue
UV
Blue - UV
Centers
Surrounds
Mean Temporal Filters (units = z-score)
Mean Spatial RF
Norm. Radial RF
