## Supplementary material for "Infrequent strong connections constrain connectomic predictions of neuronal function": Table S2 and Figures S3-S8

| Cell Type | Figure(s) | Flywire cell ID # |
|---|---|---|
| Dm2 | 1 | 720575940618018817 |
| L1 | 4 | 720575940628546759 |
| Mi2 | 4 | 720575940624931516 |
| Pm5 | 4 | 720575940612554407 |
| Dm13 | 4, 5 | 720575940648360580 |
| Pm2a | 5 | 720575940621422597 |
| Mi15 | 5 | 720575940621430789 |

**Table S2.** Related to Figures 1, 4, and 5. Flywire cell IDs for 3D reconstructions shown in the Figures. All IDs refer to FAFB version 783 (Dorkenwald et al., 2024; Matsliah et al., 2024). On Flywire, Pm5 is called “Pm02” and Pm2a is called “Pm03” (Nern et al., 2024).

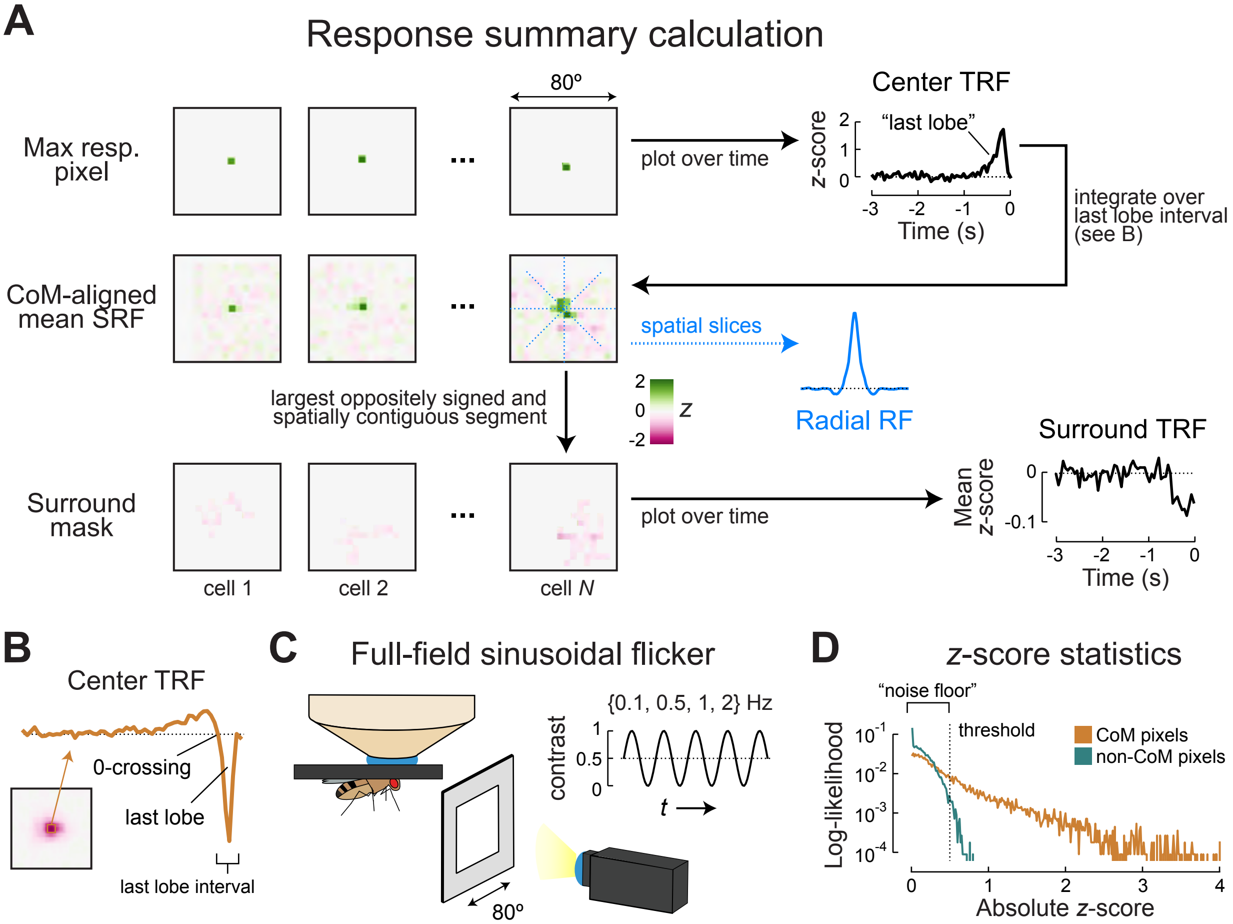

A
Response summary calculation
Max resp. pixel
80º
plot over time
Center TRF
z-score
"last lobe"
Time (s)
integrate over last lobe interval (see B)
CoM-aligned mean SRF
spatial slices
Radial RF
largest oppositely signed and spatially contiguous segment
z
Surround mask
plot over time
Surround TRF
Mean z-score
Time (s)
cell 1
cell 2
cell N
B
Center TRF
0-crossing
last lobe
last lobe interval
C
Full-field sinusoidal flicker
{0.1, 0.5, 1, 2} Hz
contrast
t
80º
D
z-score statistics
"noise floor"
threshold
CoM pixels
non-CoM pixels
Log-likelihood
Absolute z-score

**Figure S3.** Related to Figure 1. *Description of visual response profiles*.

(A) Procedure for summarizing visual responses, illustrated for the Dm2 data from Fig. 1F-G. For each neuron, we first identify the max responding patch and define the time-course of this patch's response as the center TRF. Integrating the full STRF over the interval defined by the center TRF's last lobe yields a mean SRF (which is used for CoM alignment, as shown in Fig. 1F). The largest, oppositely signed, and spatially contiguous portion of the mean SRF is defined as the surround for each neuron. We calculate the surround TRF as the mean response of surround patches over time. Finally, radial slices through the mean SRF are averaged to produce the radial RF (blue).

(B) Additional clarification of last lobe definition for mean SRF calculations. An example time-course of a max responding screen patch ("Center TRF") is shown. The last lobe interval is defined as the time from the last 0-crossing to the end of the TRF. Integrating the whole STRF over this window produces the mean SRF (as in Fig. S3A).

(C) A whole-screen (80º x 80º) sinusoidal flicker stimulus was also presented to better drive responses in neurons with large, slow receptive fields. The stimulus begins at mean gray (contrast = 0.5), increases to full on (1), then decreases to full off (0) before returning to mean gray again. Each stimulus is 10 s in duration, and flashes at the frequencies shown.

(D) Log-likelihood of absolute *z*-score values at the STRF center of mass (orange) or away from the center of mass (teal). Dotted line indicates the *z*-score (0.5) used for thresholding, as illustrated in Fig. 3A. Response elements with *z*-score values below this threshold cannot be distinguished from noise.

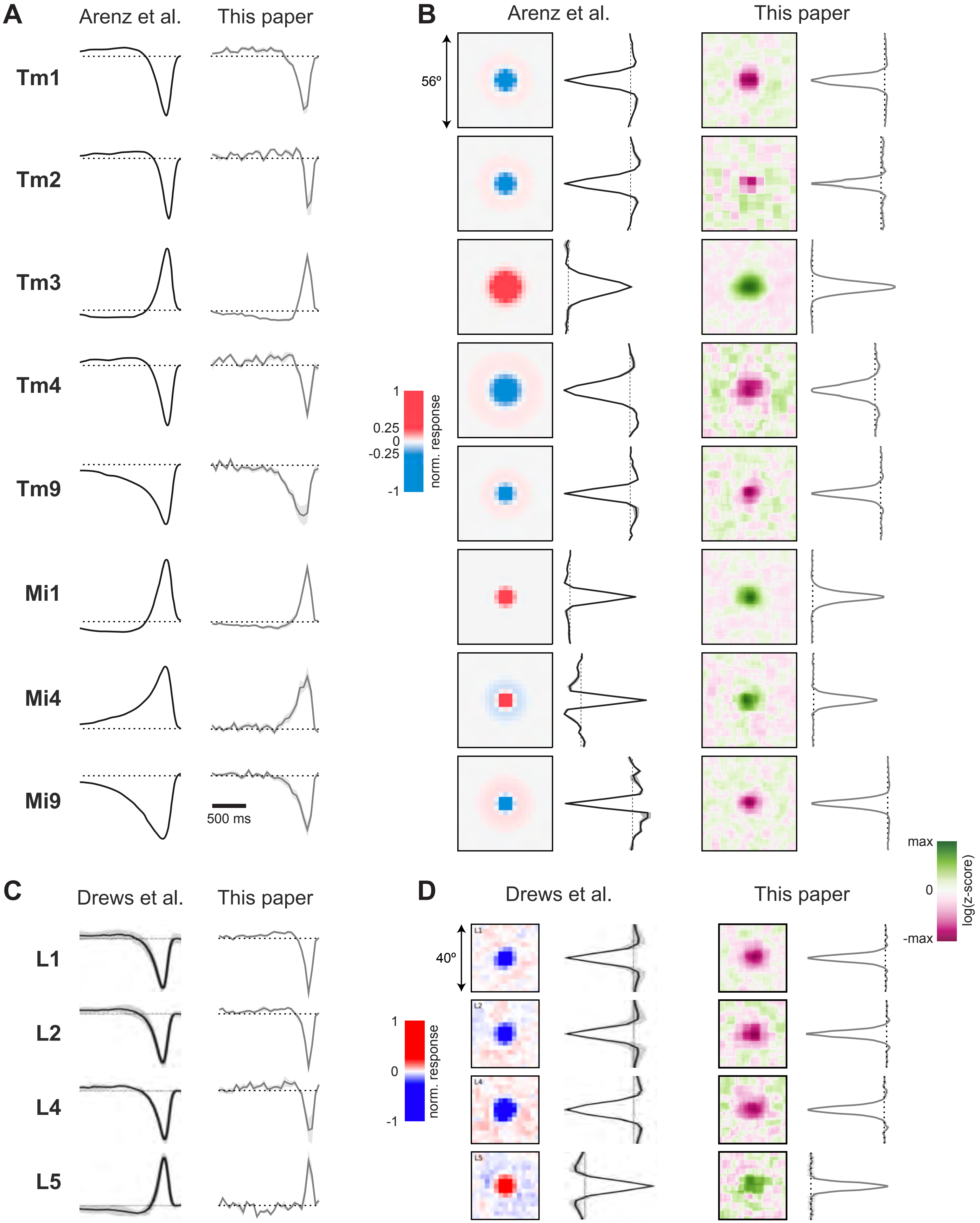

A
Arenz et al.
This paper
Tm1
Tm2
Tm3
Tm4
Tm9
Mi1
Mi4
Mi9
500 ms
B
Arenz et al.
This paper
56º
1
0.25
0
-0.25
-1
norm. response
max
0
-max
log(z-score)
C
Drews et al.
This paper
L1
L2
L4
L5
D
Drews et al.
This paper
40º
L1
L2
L4
L5
1
0
-1
norm. response

**Figure S4.** Related to Figure 1. *Verifying anatomy-based cell type identification by comparing responses to existing measurements of the same cell types*.
Temporal filters (A,C) and spatial receptive fields (B,D) for eight medulla cell types in the motion detection pathway (A,B) and four lamina monopolar cell types (C,D). All responses are peak-normalized and plotted as mean +/- SEM. Our measurements largely agree with the findings of Arenz et al., 2017 (A,B) and Drews et al., 2020 (C,D) with two primary differences. Our responses are faster, likely due to differences in indicator kinetics (GCaMP8m (in our data) vs. GCaMP6f (in previous work)), and our 2D ternary noise stimulus does not drive spatial surrounds as strongly as the 1D noise used previously. To illustrate that spatial surrounds do exist in our data, we use a logarithmic spatial color map, following similar strategies used previously (Arenz et al., 2017; Drews et al., 2020). Full response details and linear spatial colormaps for all cell types can be found in Fig. S1.

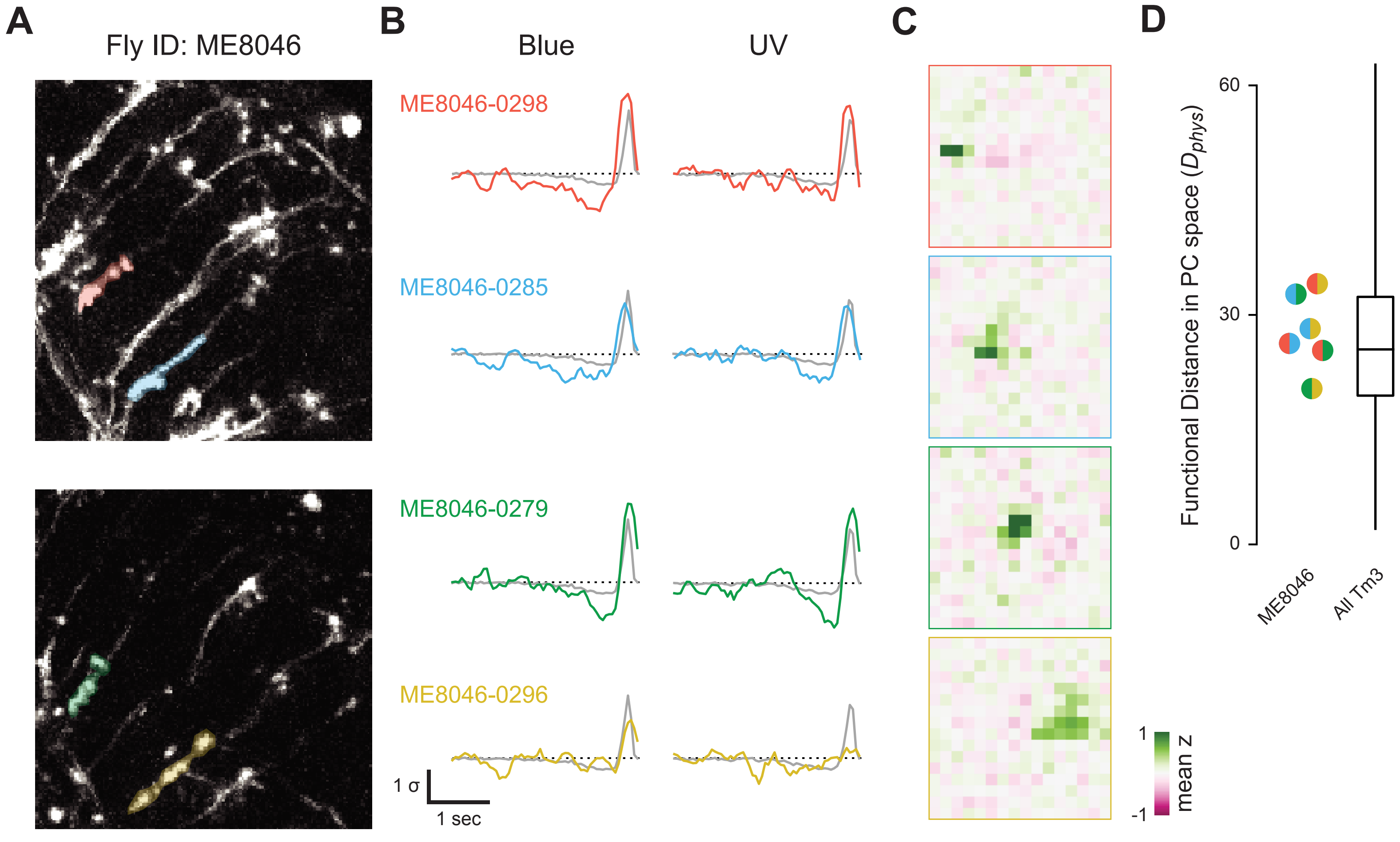
A
Fly ID: ME8046
B
Blue
UV
ME8046-0298
ME8046-0285
ME8046-0279
ME8046-0296
1 σ
1 sec
C
1
-1
mean z
D
Functional Distance in PC space ($D_{phys}$)
60
30
0
ME8046
All Tm3

**Figure S5.** Related to Figure 1. *Within-cell type variance in a single fly*.
(A) Two slices through the imaging volume of fly ME8046, showing GCaMP8m expression. ROIs for four responsive Tm3 neurons are shown as colored masks. Note that each ROI covers a similar portion of each Tm3.
(B) Center temporal filters to blue (left) and UV (right) noise for the four ROIs shown in (A). Responses are drawn in color, and the mean Tm3 response is shown in gray on each plot. Note the variation in temporal dynamics across cells.
(C) Mean spatial receptive fields for each ROI in (A) prior to centering. These four Tm3 neurons "look" at different parts of the screen, but also vary in size and surround strength.
(D) Euclidean distance in PC space ($D_{phys}$, see Fig. 2) for each pair of neurons described in (A-C), plotted next to a box-and-whisker plot of the full distribution of Tm3 within-type pairwise $D_{phys}$. Each point is colored according to the two neurons being considered. The box indicates the middle quartiles, the whiskers indicate the full range of the data, and the horizontal line is the distribution median. Different Tm3 neurons recorded in the same fly can vary in function by as much as 75% of the total within-type variance. Tm3 pairs with the largest $D_{phys}$ occur across flies. See Fig. S7C for within-type variance for all cell types in the main dataset.

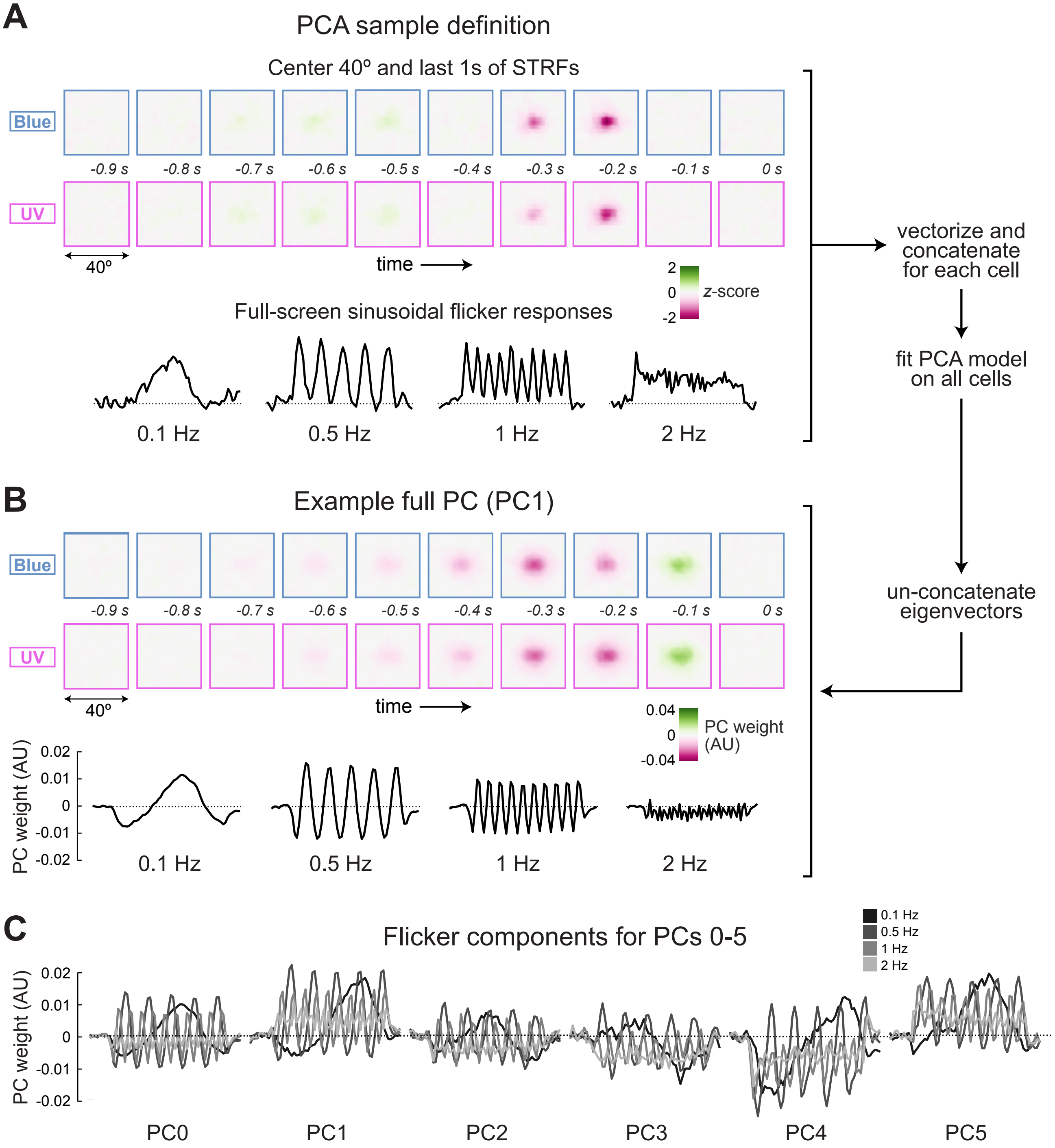
A
PCA sample definition
Center 40º and last 1s of STRFs
Blue
UV
-0.9 s
-0.8 s
-0.7 s
-0.6 s
-0.5 s
-0.4 s
-0.3 s
-0.2 s
-0.1 s
0 s
40º
time
2
0
-2
z-score
Full-screen sinusoidal flicker responses
0.1 Hz
0.5 Hz
1 Hz
2 Hz
vectorize and concatenate for each cell
fit PCA model on all cells
un-concatenate eigenvectors
B
Example full PC (PC1)
0.04
0
-0.04
PC weight (AU)
PC weight (AU)
0.02
0.01
0
-0.01
-0.02
C
Flicker components for PCs 0-5
0.1 Hz
0.5 Hz
1 Hz
2 Hz
PC0
PC1
PC2
PC3
PC4
PC5

**Figure S6.** Related to Figure 2. *Additional details of principal components analysis*.
(A) Sample construction for PCA model fitting. For each neuron, we vectorize (flatten) and concatenate the center 40º and last 1 s of the blue and UV STRFs. This spacetime period contains almost all of the signal across cells. We also concatenate the dF/F response to each flicker stimulus. The full flattened and concatenated STRFs and flicker responses define one sample for the PCA.
(B) Each resulting eigenvector has this flattened and concatenated structure, so we de-vectorize and de-concatenate eigenvectors for visualization (as shown in Fig. 2A). Each PC therefore has a blue STRF portion, a UV STRF portion, and a set of flicker portions. All of these elements for a single example PC (PC1) are plotted.
(C) Flicker portions of the first six PCs. Subsets of the STRF portions of these same PCs are shown in Fig. 2A.

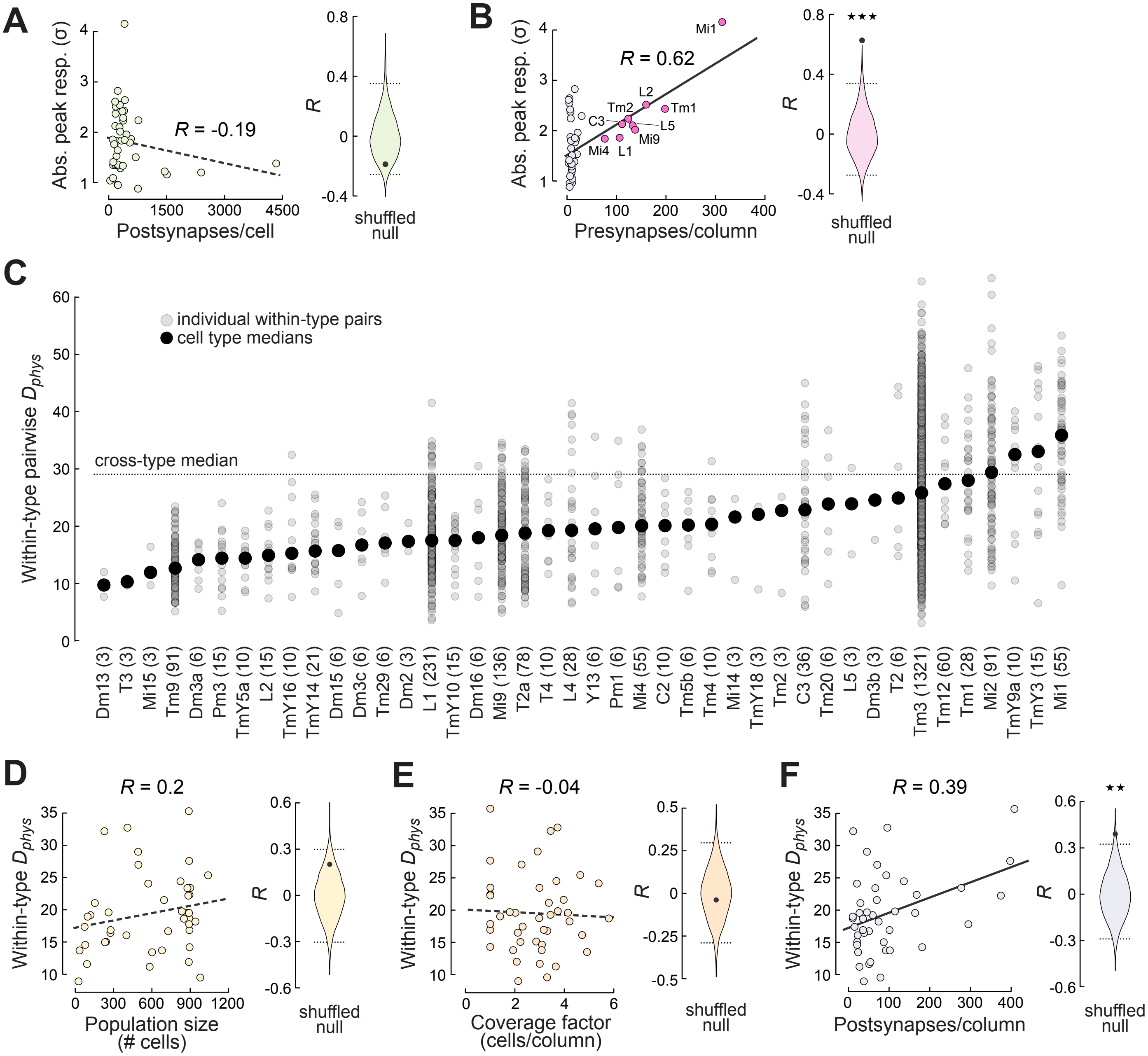

A
Abs. peak resp. (σ)
R = -0.19
Postsynapses/cell
R
shuffled null
B
Abs. peak resp. (σ)
R = 0.62
Mi1
L2
Tm2
Tm1
C3
L5
Mi9
Mi4
L1
Presynapses/column
★★★
shuffled null
C
individual within-type pairs
cell type medians
cross-type median
Within-type pairwise $D_{phys}$
Dm13 (3)
T3 (3)
Mi15 (3)
Tm9 (91)
Dm3a (6)
Pm3 (15)
TmY5a (10)
L2 (15)
TmY16 (10)
TmY14 (21)
Dm15 (6)
Dm3c (6)
Tm29 (6)
Dm2 (3)
L1 (231)
TmY10 (15)
Dm16 (6)
Mi9 (136)
T2a (78)
T4 (10)
L4 (28)
Y13 (6)
Pm1 (6)
Mi4 (55)
C2 (10)
Tm5b (6)
Tm4 (10)
Mi14 (3)
TmY18 (3)
Tm2 (3)
C3 (36)
Tm20 (6)
L5 (3)
Dm3b (3)
T2 (6)
Tm3 (1321)
Tm12 (60)
Tm1 (28)
Mi2 (91)
TmY9a (10)
TmY3 (15)
Mi1 (55)
D
R = 0.2
Within-type $D_{phys}$
Population size (# cells)
shuffled null
E
R = -0.04
Within-type $D_{phys}$
Coverage factor (cells/column)
shuffled null
F
R = 0.39
Within-type $D_{phys}$
Postsynapses/column
★★
shuffled null

**Figure S7.** Related to Figure 5. *Additional structure-function correlations*.

(A) Absolute peak response as a function of postsynapse number, plotted as in Fig. 5. The correlation coefficient of -0.19 does not cross the 95% CI of the shuffled null distribution.

(B) Absolute peak response as a function of presynapse density, plotted as in Fig. 5. The significant correlation coefficient of 0.62 is driven exclusively by a handful of highly presynaptic unicolumnar cell types, shown in pink.

(C) Within-type pairwise Euclidean distance in PC space ($D_{phys}$) for each cell type in the main dataset. Each within-type pair is plotted as a semi-transparent gray dot, with larger black dots indicating the median for each cell type. Dotted line indicates the median cross-type $D_{phys}$, as shown in Fig. 2B. The number of pairs for each cell type is written in parentheses next to the cell type name. The distribution for Tm3 is replotted from Fig. S5D.

(D-F) Within-type median $D_{phys}$ for each cell type plotted as a function of population size (D), coverage factor (E), and postsynapse (input) density (F), with the corresponding shuffled $R$ distributions for each plot, as in Fig. 5. Only postsynapse density significantly predicts within-type functional variance ($R$ = 0.39).

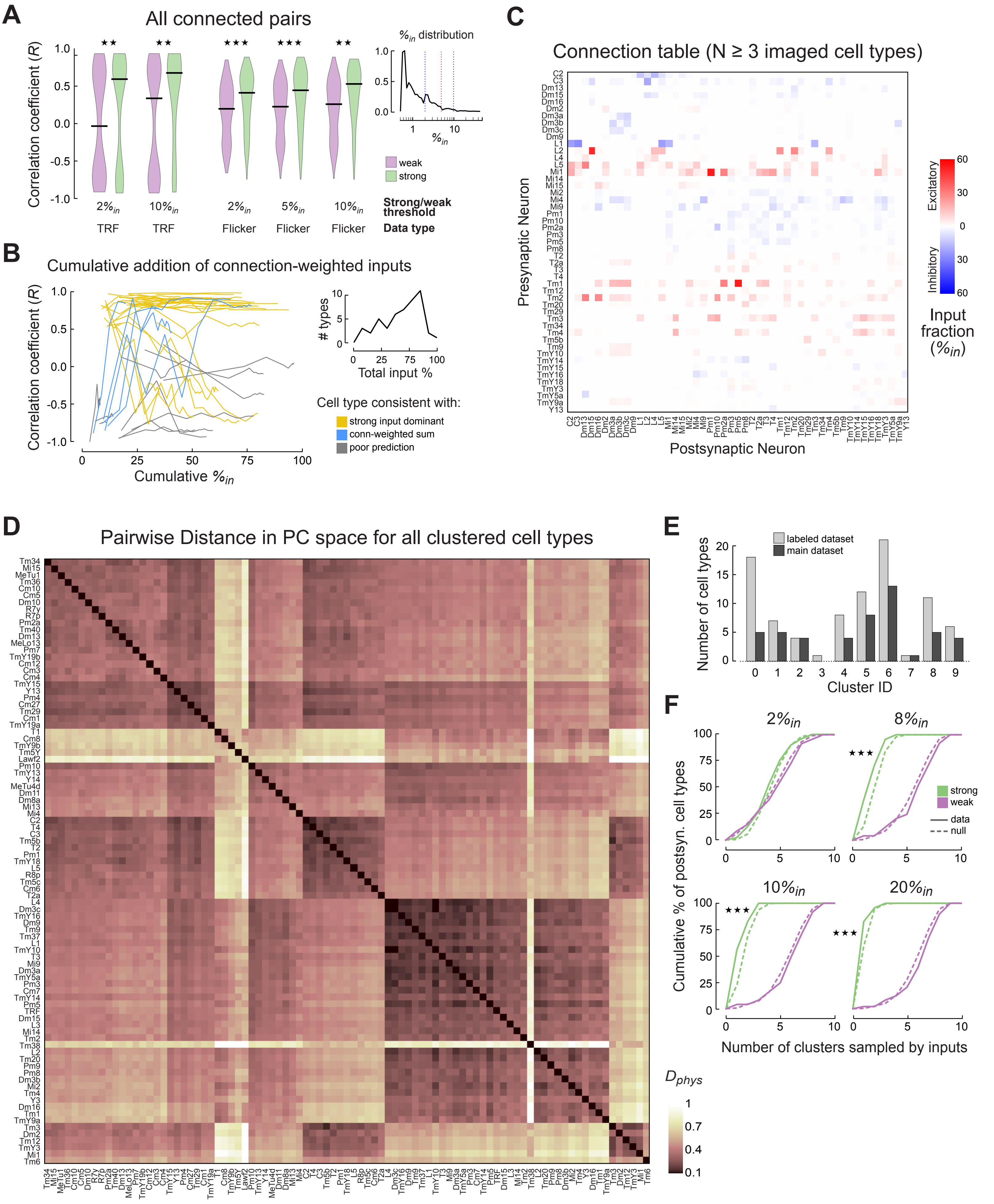

A
All connected pairs
Correlation coefficient (R)
%in distribution
weak
strong
2%in TRF
10%in TRF
2%in Flicker
5%in Flicker
10%in Flicker
Strong/weak threshold
Data type
B
Cumulative addition of connection-weighted inputs
Correlation coefficient (R)
Cumulative %in
### types
Total input %
Cell type consistent with:
strong input dominant
conn-weighted sum
poor prediction
C
Connection table (N ≥ 3 imaged cell types)
Presynaptic Neuron
Postsynaptic Neuron
Excitatory
Inhibitory
Input fraction (%in)
D
Pairwise Distance in PC space for all clustered cell types
Dphys
E
labeled dataset
main dataset
Number of cell types
Cluster ID
F
2%in
8%in
10%in
20%in
strong
weak
data
null
Cumulative % of postsyn. cell types
Number of clusters sampled by inputs

**Figure S8.** Related to Figure 6. *Additional evidence supporting the dominance and homogeneity of strong inputs*.

(A) Distributions of cross-correlation coefficients between the center TRFs (left) or full-screen flicker responses (right) of all connected pairs of cell types. More information about flicker responses can be found in Figs. S1-S3 and S7. Connections are grouped according to the fraction of total input onto the postsynaptic partner that is provided by the presynaptic partner. For each pair of distributions, one input fraction ($\%_{in}$) cutoff is considered, with connections stronger (green) or weaker (purple) than that threshold grouped together. Pre- and postsynaptic partner responses are better correlated in strong connections than weak ones, across a range of cutoff values, for both center TRF and flicker data. All comparisons are significant by rank-sum test (**, $p < 0.01$; ***, $p < 0.001$). The inset shows the overall distribution of $\%_{in}$ for connections between main dataset cell types, plotted on a log scale. Red dashed line is the 5% cutoff used in Fig. 6. Blue dashed lines are the 2% and 10% cutoffs also considered here.

(B) Correlation coefficients between measured responses and a linear connection-weighted sum prediction that considers different subsets of inputs. Each line is one cell type, colored by the model that cell type is most consistent with: yellow for the nonlinear “strong inputs dominant” model, and blue for the linear “connection-weighted sum of inputs” model. Gray lines are not consistent with either model. For each cell type, inputs are added into the connection-weighted sum prediction one at a time, from strongest to weakest. Each line plots the correlation coefficients for this family of predictions against the cumulative $\%_{in}$ as each new input is added to the prediction. The end point of each line illustrates the percent of total input accounted for by the “all inputs” data in Fig. 6C. Inset shows the distribution of $\%_{in}$ values at these end points, describing the percent of total input accounted for by main dataset cell types. A high proportion of total input is captured for most cell types.

(C) Connectivity matrix for cell types in the main dataset, used for connection-weighted sum predictions and the functional redundancy analysis in Fig. 6. Data is from Nern et al., 2024. Connections are expressed as the percentage of total input onto the postsynaptic neuron that is provided by the presynaptic neuron.

(D) Same clustered type-to-type normalized $D_{phys}$ matrix for all 91 recorded cell types from Fig. 6D, this time with each cell type labeled.

(E) Number of cell types in each functional cluster for the full dataset (light gray) and the main dataset (dark gray). Main dataset cluster counts are replotted from Fig. 6E. The general ratio of cell types in each functional cluster is largely unchanged when only main dataset cell types are considered.

(F) Cumulative probability density functions describing the number of functional clusters represented by strong (green) and weak (purple) inputs across real (solid lines) and synthetic (dashed lines) postsynaptic cell types. These plots are the same as Fig. 6F, but here we consider different strong/weak cutoff values, as in (A). Strong inputs to real neurons come from significantly fewer functional clusters than expected by chance, over a range of thresholds. All comparisons marked with *** are significant by K-S test at $p < 0.001$.
